# AlphaFold 3 Prediction Accuracy and Reliability across RNA Structures and Protein–RNA Interfaces

**DOI:** 10.64898/2026.11.02.763690

**Authors:** Iris K. Tennie, Michael Uhl, Jörg K.H. Franke, Rolf Backofen, Frederic Runge

## Abstract

AlphaFold 3 (AF3) models RNA and protein–RNA complexes, but when its predictions and confidence metrics can be trusted is unclear. We evaluated AF3 on more than 1,300 protein–RNA complexes and about 270 RNA-only systems, and compared post-cutoff RNAs with pre-cutoff candidates selected by sequence and fold similarity. On structurally novel RNAs, AF3 largely preserves the protein fold, while RNA-fold and interface accuracy drop sharply with little change in predicted RNA confidence score. Input ablations show that protein alignments and templates from RNA-bound proteins govern complex assembly, whereas RNA alignments matter most for single-chain and long RNAs. The most frequent failed interface retains protein-side contacts but places an often-misfolded RNA in the wrong pose. RNA interface confidence only partly distinguishes this error from correct placements. Against CLIP-seq, SELEX and RNAcompete motifs, AF3 usually places the protein against the experimentally observed motif-bearing RNA segment, even where the interface fails, but often not in the native pose. In protein–RNA complexes released in 2026, a score combining AF3 confidence with RNA familiarity and template use ranked RNA chains with accurate folds and correctly placed interfaces better than AF3’s ranking score.

**Graphical abstract:** 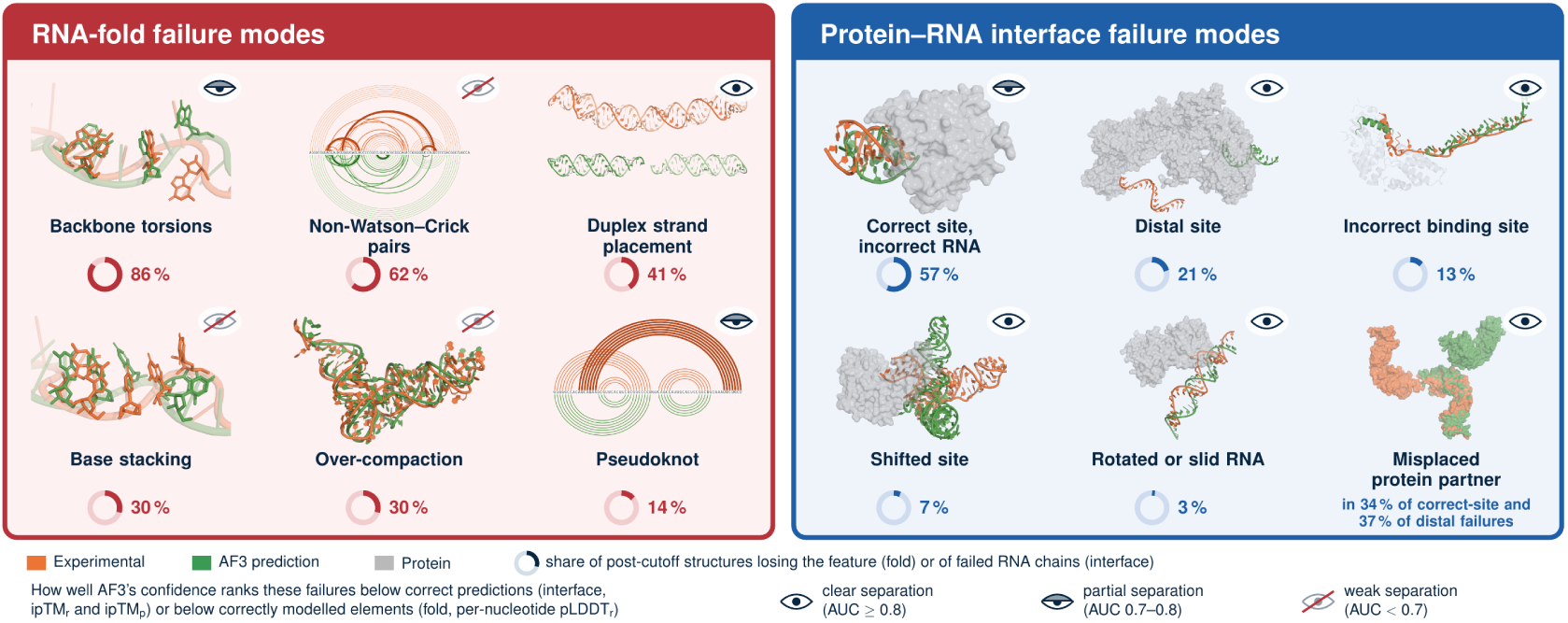

## 1. Introduction

RNA molecules and the ribonucleoprotein complexes they form act at nearly every step of gene expression, and their specificity is encoded in three-dimensional structure rather than in sequence alone [1–4]. Experimentally determined RNA and protein–RNA structures cover only a small fraction of functional RNAs and their interactions [5, 6]. Reliable prediction could therefore have broad impact. RNA is, however, a harder target than protein. It is conformationally flexible, its tertiary fold depends on metal ions and on solution conditions [7, 8], its sequence alignments are often shallow, and the small number of solved structures constrains both training and benchmarking [9–11].

Deep learning has transformed protein structure prediction [12], and a family of methods now targets RNA tertiary structure, from restraint-based folding (trRosettaRNA [13], DeepFoldRNA [14]) to end-to-end coordinate prediction (NuFold [15], RhoFold+ [16], DRfold [17]) [18]. AlphaFold 3 (AF3) predicts the structures of proteins, nucleic acids, selected ligands and ions, and their complexes in one all-atom model [19]. Related systems model protein–nucleic acid and other biomolecular complexes [20, 21]. AF3 also reports confidence metrics (per-atom pLDDT, the predicted aligned error PAE, and predicted and interface TM-scores) that are routinely used to judge a prediction when no experimental structure exists. AF3 is therefore applied to RNA-containing systems at scale, and the practical question is when its predictions and its confidence metrics can be trusted.

Part of the answer is known. Community assessments documented the difficulty of RNA structure prediction before AF3 [22, 23], and benchmarks have since ranked AF3 against other predictors on RNAs and complexes [24–26]. A recent benchmark by Ludaic and Elofsson [27] compared ten methods on 79 RNAs and four methods on 158 RNA complexes with protein, RNA or DNA partners released after the respective training cutoffs. Accuracy depended on structural similarity to RNAs released before the cutoff, which could have been available for training, so recurring folds were predicted well and divergent ones were not. Deeper RNA alignments went with higher accuracy, global and interface accuracy of a complex diverged, and several RNAs were bound to the wrong region of a correctly folded protein. The authors concluded that better model selection and confidence estimation are needed. Such a memory effect has long been documented for RNA deep-learning models [28–30]. Case studies report a bias toward canonical pairing and unreliable protein–RNA interfaces [31–33], and on training-set complexes AF3 docks a consensus-motif or poly(U) RNA at an acceptable interface about as often as the native one, although high-quality interfaces are much rarer without the native sequence [34].

These benchmarks place AF3 among the field’s leading RNA predictors, with Boltz-1 a close competitor, whereby the leading method depends on the task and metric [24, 27]. We therefore do not ask which predictor performs best. We ask, for AF3, what exactly fails when it fails, which inputs govern each failure, why the model’s own confidence misses it, and whether the missed error can be recognised from information available without an experimental structure. Where earlier work reports associations, such as that between alignment depth and accuracy [27], we test them by intervention, predicting each target of our larger benchmark with and without the input in question.

We address these questions on 1,307 protein–RNA complexes, 165 single-chain RNAs and 106 multi-chain RNA complexes from the Protein Data Bank (PDB), released before or after the AF3 training cutoff. We compare post-cutoff RNAs with pre-cutoff candidates selected separately by sequence and fold similarity to test whether the loss of accuracy tracks RNA novelty. Controlled input ablations, residue-level tracking of protein–RNA interfaces and comparison with RNA-binding protein (RBP) motif atlases identify which inputs matter and how AF3’s interfaces fail. Finally, we use these findings to build a reliability score for RNA folds and protein–RNA interfaces, which we test on complexes released in 2026. Our source code, as well as all databases, predictions and analyses are released for reuse.

## 2. Materials and Methods

This section summarises the benchmark, the accuracy metrics and the analyses. Explicit formulas, third-party tools and the feature-extraction pipeline are given in Appendix A, dataset statistics in Appendix A.1 (Supplementary Figs. S1 and S2) and the statistical procedures in Appendix B.7.

### 2.1. Benchmark and system classes

Experimental structures of protein–RNA complexes and of RNA-only systems were obtained from the PDB [35] at a resolution below 4.5 Å, downloaded between 8 February and 6 May 2025, with proteins of at least 20 amino acids and RNAs of at least 4 nucleotides. Complexes whose deposited RNA chains are broken (consecutive phosphates more than about 10 Å apart) were excluded, as were duplexes deposited as a single strand and entries whose chain identifiers could not be matched to the prediction (Appendix A.2). Protein and RNA families were annotated from UniProt [36] and Rfam [37]. Most deposited RNAs could not be assigned to an Rfam family (90% of the RNA chains in the protein–RNA complexes), because many are synthetic constructs or short fragments. Each structure was predicted with AF3 (model parameters trained with the 30 September 2021 cutoff) and paired with its prediction through a shared database key. Accuracy was computed per RNA chain, per protein–RNA interface and per complex. Because accuracy hardly changed with the random seed or with the choice among the five sampled models on 49 post-cutoff protein–RNA complexes (Appendix C), all analyses use seed 1 and model 0 unless stated otherwise. After the quality-control exclusions of Appendix A.2, the analysed sets comprise 1,307 protein–RNA complexes (923 released before and 384 after the AF3 training cutoff), 165 single-chain RNAs (99 before, 66 after) and 106 multi-chain RNA complexes (86 before, 20 after). RNA-binding domains of the deposited protein chains were annotated with InterProScan (Appendix O).

We distinguish three system classes by chain composition. Single-chain RNA systems contain one RNA chain and no protein. Single protein–RNA (1v1) complexes contain exactly one protein and one RNA chain, and multi-chain complexes are any larger protein–RNA assembly. Multi-chain RNA complexes, two or more interacting RNA chains without protein, are analysed separately (Appendix D). At the level of individual RNA chains, the regression working set contains 165 single-chain RNA, 344 1v1 and 1,996 multi-chain RNA chains. The classes are analysed separately because their determinants of accuracy differ (Section 3.5).

### 2.2. Novelty controls by training cutoff and similarity matching

Structures are split by release date relative to the AF3 training cutoff into a pre-cutoff set (b2021, available during training) and a post-cutoff set (a2021, released after training). A pre/post difference is evidence of poor generalisation only if the two sets are otherwise comparable. We therefore assess their comparability in three ways (Appendix E). First, the sets are compared on RNA length, sequence entropy, folding free energy and base composition, within the shared tRNA family, within fixed RNA-length bands and within each experimental method. Second, every post-cutoff target is paired with a pre-cutoff candidate selected by RNA sequence similarity. Pairs are grouped by RNA identity (*≥* 80%, 50–80%, *<* 50%) and, for protein–RNA complexes, by whether the twin carries the same protein (*≥* 80% identity). Third, an RNA without a similar pre-cutoff sequence (*<* 50% identity) can still adopt a familiar fold. The longest RNA chain of every post-cutoff target is therefore aligned with US-align [38] against every pre-cutoff RNA chain of all three sets, and the target is graded by the best TM-score as structurally redundant (*≥* 0.7), similar fold (0.5–0.7) or novel fold (*<* 0.5). RNA chains shorter than 20 nt were assessed by sequence similarity alone. At these lengths, the RNA TM-score distance scale *d*_0_ is at most 0.5 Å, making the score highly sensitive to small positional differences [39]. Within each group, accuracy is compared as the paired difference between the post-cutoff target and its twin, summarised by the median with a bootstrap 95% confidence interval (5,000 resamples).

### 2.3. Accuracy metrics and classification

A subscript denotes the level at which a metric is evaluated, with *_c_* for the whole complex, *_r_* for an RNA chain and *_p_* for the protein. TM*_c_* is the complex TM-score and TM*_r_* the stand-alone TM-score of the longest RNA chain of a target, or of each RNA chain in the per-chain analyses. TM*_cr_* is the TM-score of all RNA chains of a complex superposed together as one multimer without the protein (US-align multimer mode), which equals TM*_r_* for a single RNA chain. LDDT*_c_*, LDDT*_r_* and LDDT*_p_* are the corresponding all-atom LDDT values. The fold classes, the reliability score and the RNA structural classes use the backbone LDDT instead, 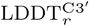 computed on the C3*^′^* atom of each nucleotide and 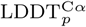 on the *C^α^* atom of each amino acid (Appendix A.10). It decides whether a chain is well folded (LDDT *>* 0.85). Unlike TM*_r_* it does not depend on chain length, and most RNAs in the protein–RNA complexes are short. It also ignores base and side-chain detail, which the all-atom LDDT penalises so strongly that too few RNAs would count as well folded to calibrate the score. The fold of the individual RNA and protein chains in the placement analysis is read from their all-atom LDDT. RMSD*_c_* and RMSD*_r_* are the root-mean-square deviations of the complex and of the RNA under the same convention. AF3’s confidence metrics follow the same convention. pTM*_p_* and pTM*_r_* are the predicted TM-scores of protein and RNA, ipTM*_p_* and ipTM*_r_* their interface scores, and pLDDT*_p_* and pLDDT*_r_* their mean per-atom confidence. The predicted aligned error PAE(*i, j*) (pae in AF3’s full-data JSON) is AF3’s expected error in the position of token *j* when the prediction is aligned on the frame of token *i*. The global PAE is its mean over all token pairs of the complex and the intra-RNA PAE its mean over the token pairs within one RNA chain. We compute both from the pae matrix as AF3 reports neither. The minimum chain-pair PAE is its lowest value between the tokens of two chains and is AF3’s own `chain_pair_pae_min` [12]. Global accuracy is read primarily from the length-normalised TM-score and from LDDT (Appendices A.9.3 and A.10), and interface accuracy from DockQ [40]. Each protein–RNA chain pair is scored separately (per-interface DockQ, DockQ*_pr_*), which defines the placement classes below. Complex-level analyses use DockQ*_Nr_*, the mean DockQ*_pr_* over all protein–RNA chain pairs of a complex, because in large assemblies the far more numerous protein–protein interfaces dominate the whole-complex DockQ (DockQ*_c_*) (Appendix M). DockQ*_pp_* is the lowest DockQ over a protein’s own protein–protein interfaces and measures how that protein is placed in the assembly. Interface LDDT, RMSD, QS-score, ICS and the base-pair interaction network fidelity (INF) are also used and are defined in Appendices A.9–B.

Several analyses group predictions by a two-stage scheme. A prediction is first assigned a fold class from the accuracy of its two components, namely (i) RNA and protein well folded, (ii) RNA only, (iii) protein only and (iv) neither. The protein counts as well folded when 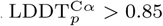 and TM*_p_ >* 0.85. The RNA counts as well folded when 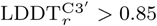 and, once the longest RNA chain reaches 60 nt, also TM*_r_ >* 0.5, because TM*_r_* is length-dependent and neither metric alone suffices. Within its fold class, the interface is graded by DockQ*_Nr_* into good (*≥* 0.8), intermediate (0.23–0.8) and poor (*<* 0.23), and poor interfaces are split by RMSD*_c_* into co-located (*<* 10 Å) and misplaced (*≥* 10 Å). The 0.8 and 0.23 boundaries are the DockQ values that the CAPRI classification assigns to high- and acceptable-quality docking models [40].

### 2.4. Controlled input ablations

Besides the sequences, AF3 accepts a multiple sequence alignment for the RNA (MSA*_r_*) and for each protein (MSA*_p_*), and structural templates for protein chains only. Tmpl*_p_* denotes a template taken from a protein-only structure, and Tmpl*_c_* a template taken from a protein deposited bound to RNA (an RNA-bound protein template). To separate the contribution of these inputs, we predicted 224 protein–RNA complexes (74 1v1 and 150 multi-chain) under all 16 combinations of MSA*_r_*, MSA*_p_*, Tmpl*_p_* and Tmpl*_c_* (3,584 predictions). The run with the input present is paired with the otherwise identical run without it, the difference is averaged over the eight combinations of the other three inputs, and only complexes that carry the input are included. Effects are reported as mean paired differences with bootstrap 95% confidence intervals and as paired standardised effects. Single-chain RNAs, which have no protein partner and hence no template, were predicted with and without MSA*_r_* (*n* = 81). In a second intervention, structures were predicted with and without the ions of the deposited entry as input (41 single-chain RNAs, 13 protein–RNA complexes and 13 multi-chain RNA complexes, paired). Full ablation results are given in Appendix I and the ion case studies in Appendix K.1.

### 2.5. Interface tracking, placement classes and RNA motifs

For each experimental and predicted protein–RNA chain pair, we identified amino acid–nucleotide contacts from hydrogen bonds and van der Waals interactions and recorded the residue identities and positions of each pair (Appendix A.5). Each native protein–RNA interface was followed into the prediction in three steps. Firstly, exact residue mapping carries the native contact residues onto the predicted chains and gives the fraction of native protein residues and RNA nucleotides that AF3 places in contact (protein and RNA contact match, Appendix A.7). Secondly, the predicted RNA is located relative to its native position after superposing the protein chain (Appendix A.7). Finally, the interface is assigned to the first of the following placement classes that applies (Appendices A.7 and N). A correct interface has a DockQ*_pr_ ≥* 0.23. A failed interface is a correct site with an incorrect RNA if AF3 still contacts native protein residues (protein contact match *≥* 0.1), in which case the RNA sits at the native site but with the wrong nucleotides or in the wrong pose. Failed interfaces without native protein contacts are first checked for the protein. The RNA-binding pocket is the set of native protein residues that contact the RNA, and its RMSD is computed on their *C^α^* atoms after superposing the protein chain. An incorrect binding site has a pocket RMSD above 4 Å. Where the pocket is in place, the class follows from the position of the RNA. A rotated RNA lies in place (centroid within 10 Å) but is turned by more than 45*^◦^*, a shifted site lies 10–20 Å away and a distal site more than 20 Å away. Where the native RNA is one strand of a duplex, AF3 can place the duplex correctly but exchange the two strands. Scored on that strand alone the RNA then looks turned by about 180*^◦^*, the dyad of the duplex, or displaced, although the duplex itself has barely moved. Where the predicted duplex covers at least half of the native duplex while the native strand does not occupy its own position, the pose is therefore read from the whole duplex. A duplex moved by at most 10 Å with its helix axis tilted by at most 45*^◦^* is a correct site with a slid duplex. AF3 then keeps the duplex on its native track but slides it by a few base pairs and exchanges the strands. A protein partner counts as misplaced in the assembly when DockQ*_pp_* is below 0.23. An RNA chain that binds several proteins is also described by its main partner, the protein with the largest native interface (most protein residues in contact with the RNA in the deposited structure; between two equally large interfaces, the one with the higher DockQ*_pr_*). Its placement class, contact matches and position are then those of this one interface, without merging the interfaces. A protein chain counts as accurately folded when its LDDT*_p_* and TM*_p_* exceed 0.85, as in fold classes i and iii, and an RNA chain as well folded when LDDT*_r_* exceeds 0.85 (with TM*_r_ >* 0.5 from 60 nt) and as misfolded when LDDT*_r_* is at most 0.75. Since short RNAs offer fewer possible contact positions, we report RNA contact match separately for RNAs of at most 15 nucleotides and those longer than 15 nucleotides. We assessed the sensitivity of class frequencies across 1,280 combinations of the five classification thresholds and checked whether a partner strand of a duplex or an identical protein copy occupied the native position (Appendix N). To ask whether failed interfaces reflect model ranking or model generation, 49 post-cutoff targets were predicted with three seeds and five models each. An interface that fails in the default model counts as recoverable if any of the 15 models reaches DockQ*_pr_ ≥* 0.23.

For the comparison with experimental binding preferences, every run of at least four consecutive protein-contacted nucleotides of each experimental and predicted interface (its RNA motifs) was scored against the position probability matrices of a merged atlas of CLIP-seq-, SELEX- and RNAcompete-derived motifs, converted to log-odds at scoring time against a uniform background, with the matrices of other RNA-binding proteins (RBPs) as a null distribution. The position of the predicted RNA motif within the RNA was compared with the native one (159 entries from 57 genes; Appendix B.3). A separate RNAcompete benchmark, in which proteins were predicted with their strongest- and weakest-binding RNA probe sequences, is described in Appendix B.2 and reported in Appendix P.

### 2.6. Regression framework and variance partition

To attribute the variation in AF3 accuracy to sources of information, the descriptors extracted for every structure, prediction and interface (Appendix B.4) were organised into six mutually exclusive groups fixed a priori. These are AF3 confidence (G0, with pLDDT, pTM, ipTM, global and chain-pair PAE), evolutionary signal (G1, with RNA and protein alignment depth, RNA family and the counts of protein-only and RNA-bound protein templates), structural complexity and composition (G2, chain numbers and lengths), RNA-intrinsic properties (G3, sequence complexity and folding energetics), predicted-interface features (G4, computed from the predicted structure only and hence not circular with the accuracy target) and experimental context (G5, ions, ligands, buffer, PEG, pH and temperature). Properties known only once the experimental structure exists, such as the determination method and resolution, were excluded from G5, because a user predicting an unknown structure does not have them. Grouping attributes explained variance to interpretable categories of information rather than to individual, often collinear, descriptors. The descriptors of each group are listed in Appendix S.

Targets were modelled with ridge regression (*α* = 10, fixed a priori) under grouped five-fold cross-validation with folds defined by PDB entry (scikit-learn GroupKFold, seed 0), so that all chains and models of one complex fall into the same fold. The primary target was TM*_c_* at the complex level and TM*_r_* at the per-chain level, with DockQ*_Nr_* as the interface-focused secondary target. Imputation, scaling and one-hot encoding were fitted inside each training fold, and no target-based feature selection was applied to the models reported in the main text. For each dataset we fitted the six single-group models, the confidence-only model with each other group added in turn, and the full model, and report the coefficient of determination (*R*^2^) on pooled out-of-fold predictions. To isolate the error that AF3 confidence does not capture, we modelled the residual of the confidence-only model, 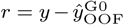, with the non-confidence groups under the same cross-validation. The credit assigned to correlated groups depends on the order in which they enter a model, so group rankings rest on a Shapley decomposition of the cross-validated *R*^2^. It averages each group’s marginal contribution over all orderings and sums to the full-model *R*^2^ (Appendix R). All *R*^2^ values measure cross-validated explained variance within the analysed PDB distribution. They are not presented as an out-of-sample predictor or as evidence of causation. A leakage-controlled stability selection re-run inside every fold confirms that the full-set results are not driven by redundant descriptors (Appendix S), and the variance partition is insensitive to *α* over two orders of magnitude (Appendix S, Supplementary Table S12).

### 2.7. Reliability score and its test

The reliability score gives calibrated probabilities for the RNA fold and the protein–RNA interface of a prediction, from information that requires neither an RNA alignment nor an experimental structure. We define the intra-RNA PAE of an RNA chain as the mean of AF3’s predicted aligned error matrix (pae in the full-data JSON) over the token pairs within that chain. A prediction has a familiar sequence when one of its RNA chains has at least 80% identity to an RNA chain of a PDB entry released before 30 September 2021, and a familiar fold when the longest RNA chain of the AF3 model reaches a TM*_r_* of at least 0.5 to an RNA chain of the pre-cutoff structures of this benchmark. Both are computed without the experimental structure. P_fold_, the probability that the longest RNA chain of a protein–RNA complex is well folded 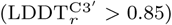, is a logistic regression on its intra-RNA PAE and four yes/no questions, namely whether the sequence is familiar, whether the fold is familiar, whether AF3 used an RNA-bound protein template for any protein chain and whether the model has two or more RNA chains. P_interface_, the probability that an RNA chain is placed correctly, is a logistic regression on AF3’s minimum chain-pair PAE between that chain and a protein chain (`chain_pair_pae_min` in the summary JSON), a familiar RNA (sequence or fold) and a well-folded partner (pTM*_p_ ≥* 0.84 and pLDDT*_r_ ≥* 0.78). P_usable_, the probability that an RNA chain is both well folded and placed correctly, joins the P_fold_ and P_interface_. RNA fold and interface are not independent, because a well-folded RNA is also more often placed correctly, so their product would tend to underestimate this probability and the smaller of the two would overestimate it. The two probabilities are therefore joined with a Gaussian copula, a joint distribution between these two limits. Its single correlation of 0.66 was fitted out of fold on post-cutoff targets. Every probability is reported at three levels, WEAK below 25%, MODERATE from 25% to 60% and STRONG from 60%. The fold questions were selected on post-cutoff targets by cross-validated greedy forward selection among 25 candidates, and the interface questions were chosen on pre-cutoff targets (Appendix T). pLDDT*_r_* and ipTM*_r_* were to be added only if they raised the cross-validated AUC by at least 0.01, which neither did. The weights and the copula correlation were fitted on post-cutoff targets, and the score was then frozen. RNA-only predictions have their own logistic models, on pLDDT*_r_* or the intra-RNA PAE depending on whether the RNA alignment was used and, for RNA complexes, on whether the AF3 model forms a duplex, chosen and fitted on post-cutoff RNAs, namely the post-cutoff single-chain RNAs and RNA complexes of this benchmark (Appendix T.4). The frozen scores were tested on 99 protein–RNA complexes (released between May and September 2026), 41 single-chain RNAs (released between May 2025 and August 2026) and 10 RNA complexes without protein (released between June 2025 and May 2026), of which all were released after this benchmark was assembled. They were predicted with the AlphaFold Server and processed with the same pipeline. The score is released as the decision_tree package of the code repository.

## 3. Results and Discussion

We analyse AF3 predictions for 1,307 protein–RNA complexes (344 single protein–RNA and 963 multi-chain assemblies), 165 single-chain RNAs and 106 multi-chain RNA complexes, of which 384, 66 and 20 were released after the AF3 training cutoff. On 49 post-cutoff protein–RNA complexes predicted with three seeds and five models each, accuracy hardly changed with the seed or the model (Appendix C, Supplementary Fig. S3), so we use one seed and the top-ranked model throughout unless stated otherwise.

### 3.1. RNA novelty causes a generalisation failure that AF3’s confidence does not register

To separate generalisation to novel RNAs from recall of training-era folds, we compared targets released before (b2021) and after (a2021) the AF3 training cutoff. Protein- and complex-level accuracy shift comparatively little with novelty. The protein fold is essentially preserved on unseen complexes (mean LDDT*_p_* 0.892 *→* 0.836, *d* = 0.64), while complex accuracy declines modestly (mean LDDT*_c_* 0.850 *→* 0.750, *d* = 0.86). In contrast, all RNA-dependent metrics fall steeply (Fig. 1). Within the protein–RNA complexes, TM*_r_* drops from 0.587 to 0.388 (*d* = 0.97), LDDT*_r_* from 0.854 to 0.686 (*d* = 1.02) and the protein–RNA interface score DockQ*_Nr_* from a median of 0.79 to 0.30 (*d* = 1.11). Single-chain RNA folds decline in LDDT*_r_* from 0.823 to 0.701 (*d* = 0.87) and in TM*_r_* from 0.705 to 0.594. Multi-chain RNA complexes decline in LDDT*_r_* (0.904 *→* 0.719, *d* = 1.32) but not significantly in TM*_r_* (0.629 *→* 0.504, *d* = 0.46, *P* = 0.08). Their loss is largest at the RNA–RNA interface, where the median nucleotide contact match falls from 0.95 to 0.26 (Appendix D, Supplementary Fig. S4).

This RNA accuracy gap is not explained by differences between the two sets. The pre- and post-cutoff protein–RNA sets do not differ appreciably in RNA length, sequence entropy, folding free energy or base composition (Cohen’s *|d| ≤* 0.25), and the drop persists within the shared tRNA family, within fixed RNA-length bands and within X-ray and cryo-EM structures analysed separately (Appendix E, Supplementary Fig. S5).

Instead, the gap follows the novelty of the RNA. When a post-cutoff RNA has a near-identical pre-cutoff twin (*≥* 80% sequence identity) with the same protein, AF3 predicts it about as well as the twin (ΔLDDT*_r_* = *−*0.020). With another protein it loses 0.100, half the loss of a novel RNA. Release after the cutoff therefore lowers accuracy little by itself, and the pooled gap of Fig. 1 is one of generalisation rather than of recall. Where the closest twin is only 50–80% similar, LDDT*_r_* falls by 0.20. The loss is largest when the novel RNA is bound to a protein that also occurs before the cutoff (*−*0.33, *n* = 12). Thus, a familiar protein partner does not rescue a novel RNA. Measured by structure, 68% of the post-cutoff protein-bound RNAs have no pre-cutoff RNA of similar fold (TM*_r_ ≥* 0.5), and these novel folds lose most against their closest structural twin (ΔLDDT*_r_* = *−*0.19, against *−*0.04 for structurally redundant RNAs) (Appendix E, Supplementary Figs. S6 and S7). Template leakage does not explain the retained prediction accuracy either, as AF3 used the matched pre-cutoff complex as a template in only 15 of 384 pairs (Appendix I).

The loss is concentrated in particular RNA architectures. After the cutoff, the median LDDT*_r_* of pseudoknots falls from 0.91 to 0.63 and that of internal loops from 0.92 to 0.66, whereas hairpins decline less (0.92 *→* 0.82; Appendix G, Supplementary Fig. S8; examples in Supplementary Fig. S12). AF3 reproduces the number of secondary-structure elements. However, it builds more compact folds than the targets, with longer stems and shorter loops.

**Fig. 1:**
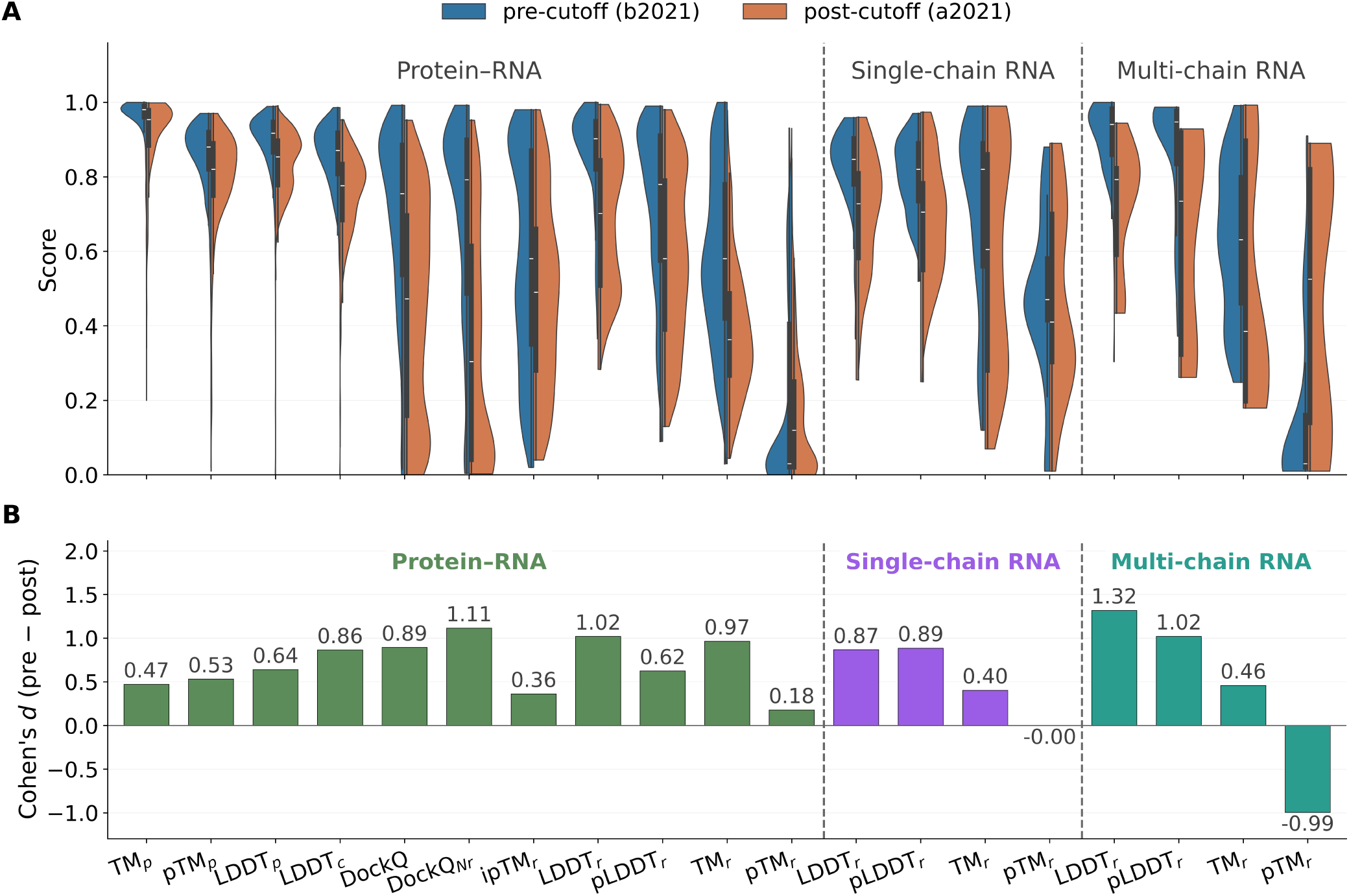
The pre-versus post-cutoff accuracy gap is large and specific to the RNA. Structures released before (b2021, blue) and after (a2021, orange) the AF3 training cutoff, for protein–RNA complexes, single-chain RNAs and multi-chain RNA complexes (dashed dividers). **A**, Distributions of accuracy and confidence (inner box, median and interquartile range). **B**, Cohen’s *d* (pre *−* post) beneath each distribution. RNA accuracy drops most, while pTM*_r_* barely moves.

AF3’s confidence does not reflect this loss. The RNA interface confidence shifts only modestly across the cutoff (ipTM*_r_*, *d* = 0.36), and the predicted RNA TM-score barely moves (pTM*_r_*, *d* = 0.18; single-chain RNAs 0.474 *→* 0.475, *d* = 0.00). In the multi-chain RNAs, pTM*_r_* even rises after the cutoff (*d* = *−*0.99), because the post-cutoff chains are much longer and pTM*_r_* grows with length (Appendix D). Unsurprisingly, AF3’s reliable confidence metrics are protein-centric. Protein ipTM*_p_* tracks TM*_c_* (Spearman *ρ ≈* 0.68), pLDDT*_p_* tracks LDDT*_c_* (*ρ ≈* 0.76), and pTM*_p_* separates accurately from poorly folded proteins (median 0.80 against 0.74, AUC 0.73). The global PAE rises from 7.5 in correct to 16.9 in incorrect single-RNA complexes (*ρ ≈* 0.76 with RMSD*_c_*; Appendix H, Supplementary Fig. S14B), and the minimum chain-pair PAE separates good from poor interfaces (*≈* 2.3 against *≈* 10). The RNA interface confidence ipTM*_r_* decreases monotonically from well-modelled to misplaced interfaces (*ρ* = 0.48; Fig. 2B). The RNA fold confidence, however, is informative only when evolutionary information is available. pTM*_r_* tracks TM*_r_* when the RNA has an alignment (*r* = 0.80, *n* = 102) and is uninformative without MSA*_r_* (*r* = 0.03, *n* = 550; Fig. 2A). Its strongest correlate is the RNA folding free energy (*ρ* = *−*0.76), ahead of alignment depth (*ρ* = 0.57) and any accuracy score (Appendix H, Supplementary Fig. S14C).

The loss therefore follows the novelty of the RNA rather than its release date, it lies in the RNA fold and the protein–RNA interface rather than in the protein. Mixing structures released before and after the training cutoff would therefore overestimate the accuracy that AF3 reaches on novel RNAs. This concern applies to learned predictors in general [41, 42] and to RNA in particular, given the few solved structures and the repeated representation of a few families [24, 43].

### 3.2. RNA folding and complex assembly depend on different information channels

To establish which input governs which part of the accuracy, we predicted 224 protein–RNA complexes under all 16 combinations of RNA alignment (MSA*_r_*), protein alignment (MSA*_p_*), protein-only template (Tmpl*_p_*) and RNA-bound protein template (Tmpl*_c_*), estimating each effect within target (Section 2.4, Appendix I, Supplementary Fig. S16). We also predicted 81 single-chain RNAs with and without their alignment.

**Fig. 2:**
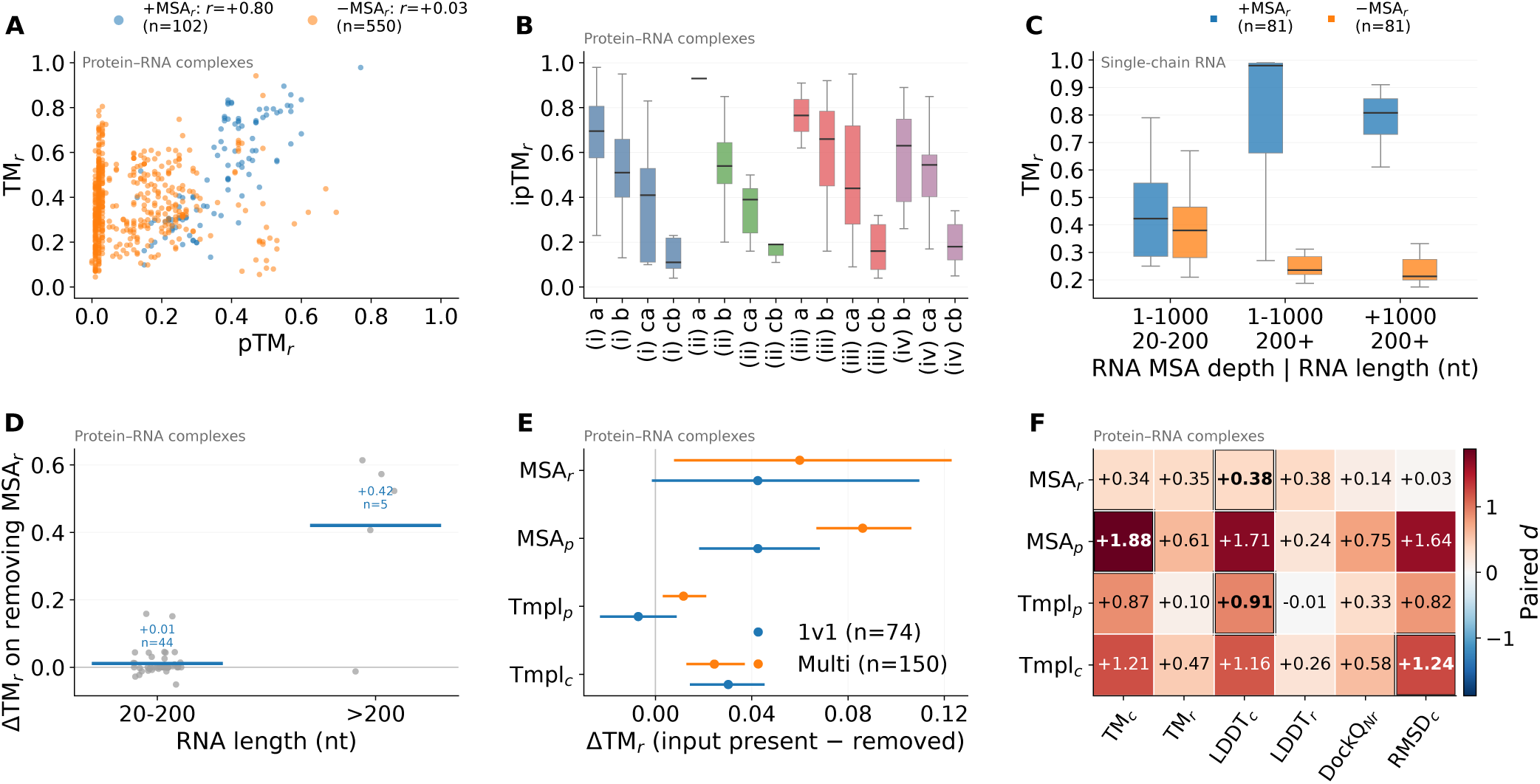
AF3’s RNA confidence and the contribution of each input. **A**, pTM*_r_* against TM*_r_* per RNA chain, with and without an RNA alignment. **B**, ipTM*_r_* by fold class (i–iv) and interface subclass (a, good; b, intermediate; ca, poor and co-located; cb, poor and misplaced). **C**–**F**, Controlled input ablation over all 16 combinations of MSA*_r_*, MSA*_p_*, Tmpl*_p_* and Tmpl*_c_*, with each effect estimated within target. **C**, Single-chain RNAs with and without their alignment, by alignment depth and length. **D**, Change in TM*_r_* on removing the RNA alignment from protein–RNA complexes, by RNA length. **E**, Effect of each input on TM*_r_* (95% bootstrap CI). **F**, Paired standardised effect of each input on each metric.

Since single-chain RNAs have no protein partner and hence no structural template, the RNA sequence alignment is their only external source of evolutionary information. Its benefit grows with alignment depth and is negligible for the shortest RNAs (Fig. 2C), in agreement with the rise of TM-score with RNA alignment depth reported by Ludaic and Elofsson [27]. Length and depth are strongly confounded (Spearman *ρ* = +0.71 post-cutoff and +0.85 pre-cutoff), so the apparent length dependence largely reflects alignment availability. Stratifying by RNA family shows the same dependence. Conserved, deeply aligned families such as tRNA and SSU rRNA are predicted well above the dataset mean (ΔTM*_r_* = +0.33 and +0.55), whereas unannotated synthetic RNAs sit at or below it (Appendix J, Supplementary Fig. S19).

In protein–RNA complexes, accuracy is governed by protein evolutionary information and by templates. The protein alignment dominates every complex- and interface-level metric (paired *d* = 1.88 for TM*_c_*; Fig. 2F). Removing MSA*_p_* lowers mean TM*_c_* from 0.86 to 0.55 among targets that carry one (paired difference +0.31, 95% CI 0.29–0.33). Templates add a smaller gain, and the RNA-bound protein template contributes more than the protein-only one (ΔTM*_c_* = +0.15, 95% CI 0.13–0.17, against +0.10, 0.08–0.11). It also raises the accuracy of the RNA itself, more than the protein-only template does (Fig. 2E). Template identity matters more than template count. Complexes without any template are weakest (TM*_c_* 0.64), protein-only templates give moderate (0.75), RNA-bound protein templates the largest (0.82) and both together the highest gains (0.87, *n* = 195, *P* = 1.1 *×* 10*^−^*^3^ against the baseline; Appendix I). AF3 rarely selects its own PDB-deposited structure as a template. Only 12.6% of pre-cutoff complexes list their own entry, because AF3 reuses one representative structure per family for pre- and post-cutoff members alike (Appendix I). The RNA alignment matters little in complexes with short RNAs. It raises TM*_r_* by only 0.01 in the 44 alignment-carrying complexes with RNAs of 20–200 nt, but by 0.42 (95% CI 0.20–0.58) in the five with RNAs longer than 200 nt (Fig. 2D).

A second intervention concerns the ions that stabilise RNA tertiary structure [7, 44, 45]. Supplying the deposited ions as input improves single-chain RNA folds (*n* = 41, LDDT*_r_* 0.699 *→* 0.749, *P ≤* 0.002), most for weakly aligned RNAs. It brings no gain for protein–RNA complexes (*n* = 13) or multi-chain RNA complexes (*n* = 13), and in complexes AF3 tends to place the added ions away from their coordinating sites (Appendix K.1, Supplementary Figs. S20 and S21).

In summary, the protein alignment and the RNA-bound protein template act on the assembly, the RNA-bound template also improves the RNA, and the RNA sequence alignment acts on the RNA fold. For the protein–RNA interface, the benefit from an RNA-bound template most likely comes from a protein conformation suited to RNA binding. For single-chain RNAs, which have no other external signal, the alignment can decide whether the fold is correct at all (Appendix I, Supplementary Fig. S18). Its useful content appears to be base-pairing topology, since synthetic alignments, which encode only a predicted secondary structure, approach the accuracy of natural ones [46]. Ions substitute in part for missing evolutionary signal in single-chain RNAs.

**Fig. 3:**
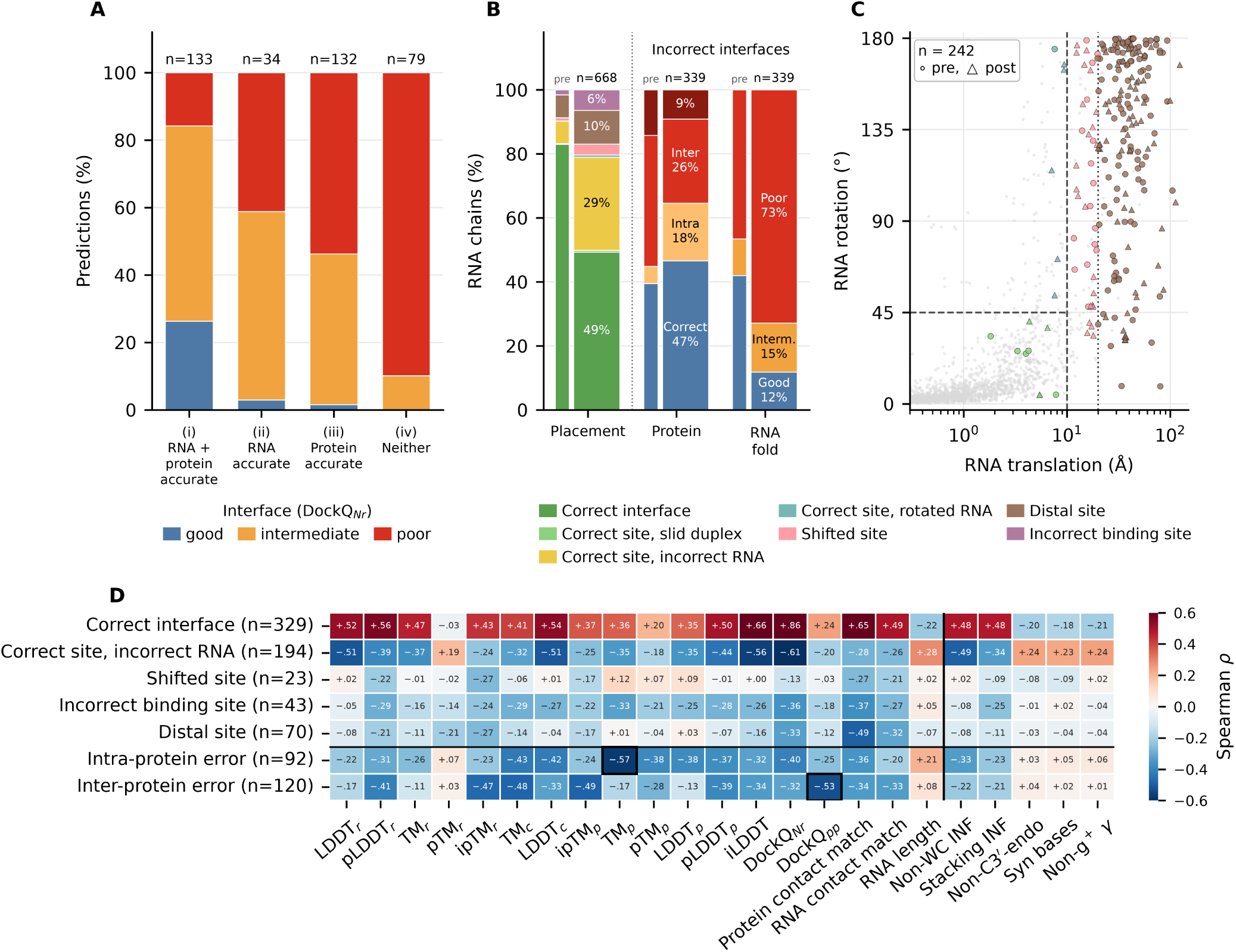
How AF3 protein–RNA interfaces fail. **A**, Interface quality (DockQ*_Nr_*) within each fold class (i, RNA and protein accurate; ii, RNA accurate; iii, protein accurate; iv, neither; post-cutoff). **B**, Placement class of each RNA chain with its main protein partner before (thin bars) and after the cutoff (Section 2.5), and, for the RNA chains whose interface is incorrect, whether the main partner is misplaced against its protein partners (inter, DockQ*_pp_ <* 0.23), less accurately folded itself (intra, TM*_p_ ≤* 0.85) or both, and the RNA fold. **C**, Whole-RNA translation and rotation in the protein frame for the failed main-partner interfaces without native protein contacts and with the pocket in place (circles, pre-cutoff; triangles, post-cutoff; grey, correct interfaces). Correct sites with an incorrect RNA are assigned by protein contact and are not shown, apart from one RNA that lies in place without contact. Dashed lines mark the 10 Å and 45*^◦^* cut-offs. Slid duplexes are placed by the shift and axis tilt of the whole duplex. **D**, Spearman *ρ* of each class’s share of a complex’s RNA chains with fold, confidence, interface and RNA-geometry descriptors (post-cutoff complexes; the last three columns are the shares of *syn* nucleotides, non-C3’-*endo* sugar puckers and non-*g+ γ* torsions in the experimental RNA structure). The two bottom rows are the shares of RNA chains that fail with an intra-protein error (main partner TM*_p_ ≤* 0.85) or an inter-protein error (main partner misplaced in the assembly, DockQ*_pp_ <* 0.23), as in B.

### 3.3. AF3 often finds the protein site but fails to pose the RNA

Protein–RNA recognition combines base-specific sequence readout with recognition of RNA shape, mediated by reusable RNA-binding domains [4, 47–49]. To locate where AF3 fails at the interface, we assigned each post-cutoff prediction a fold class: (i) RNA and protein well folded, (ii) RNA only, (iii) protein only or (iv) neither, and graded its interface by DockQ*_Nr_* within each class (Section 2.3, Appendix M, Supplementary Fig. S23). Both chains folded correctly is close to necessary but not sufficient. Of predictions with a good interface, 92% belong to class i, in which both RNA and protein are well folded. Median DockQ*_Nr_* is 0.66 for class i, compared with 0.29, 0.20 and 0.03 for classes ii, iii and iv, respectively. Yet only 26% of class-i predictions reach a good interface and 16% fail (Fig. 3A). TM*_r_* explains only about a quarter and LDDT*_r_* about two fifths of the variance in DockQ*_Nr_* (*ρ*^2^ *≈* 0.26 and 0.41; Supplementary Fig. S25), so a globally accurate RNA can still fail at the interface.

To resolve how interfaces fail, we classified each protein–RNA interface by where AF3 places the RNA relative to its protein partner (Section 2.5, Fig. 3B,C; two examples per class in Appendix N, Supplementary Fig. S28). After the cutoff, 46% of the interfaces are correct (DockQ*_pr_ ≥* 0.23), against 77% before it. As many RNA chains bind several proteins, we also describe each RNA chain by its main partner that is the protein with the largest native interface. Counted this way, 49% of the 668 post-cutoff RNA chains are placed correctly, against 83% of the 1,646 before the cutoff (Fig. 3B). Of the post-cutoff RNA chains with an incorrect interface, 73% carry a misfolded RNA (LDDT*_r_ ≤* 0.75), and their main partner is misplaced against its own protein partners in 35% or less accurately folded in 27%. The interfaces of an RNA with several proteins mostly succeed or fail together (6% of the other interfaces are correct when the main one fails, against 79% when the main interface is correct).

The dominant failure is a correct site with an incorrect RNA. AF3 contacts native protein residues with the binding pocket in place (median protein contact match 0.46, against 0.66 for correct interfaces; Appendix M, Supplementary Fig. S26), but a different part of the RNA occupies the site (nucleotide contact match 0.29 against 0.80, no native nucleotide at all in 30% of these interfaces). The RNA as a whole is turned (median rotation 97*^◦^*) and displaced (median 17 Å), and most of these RNAs are themselves poorly folded (86% in fold classes iii and iv). This class accounts for 54% of the post-cutoff failures, ahead of distal sites (23%) and incorrect binding sites (13%; Appendix N, Supplementary Tables S5 and S8).

Conversely, a correct placement does not require both chains to be accurately folded. Only 29% of the correctly placed post-cutoff RNA chains (62% before the cutoff) combine a well-folded RNA with an accurately folded main partner, and these reach the highest DockQ*_pr_* (median 0.82). A further 36% have a misfolded RNA (LDDT*_r_ ≤* 0.75), most of them short RNAs of at most 15 nt (73 of 120), and 14% combine a misfolded RNA with an inaccurately folded main partner yet still form an acceptable interface (median DockQ*_pr_* 0.43). The protein-contacting segment of the RNA can be placed correctly while the rest of the RNA diverges (Appendix N, Supplementary Table S7).

Duplex RNAs fail in a further way, as AF3 can place the duplex roughly at the native site but with the wrong strand in the native position. Of the 339 post-cutoff RNA chains with an incorrect interface, 153 are one strand of a duplex. In 28 of these, the strand at the native position has a different sequence, so the protein faces the wrong RNA. In 4 of them, the duplex stays on its native helical track, slid by 4–7 Å with the strands exchanged, in otherwise accurate complexes (Appendix N).

Distal and incorrect binding sites are mostly protein errors, of two kinds. At a distal site, the protein, including its pocket, is folded correctly (median pocket RMSD 1.2 Å, as for correct interfaces) but the RNA is docked elsewhere (median displacement 43 Å), and the protein chain is misplaced against its protein partners at 58% of these sites (lowest DockQ*_pp_ <* 0.23), against 21% of correct interfaces. At an incorrect binding site, the RNA-binding pocket itself is mispredicted (median pocket RMSD 11 Å) and the protein chain is less accurately folded (median TM*_p_* 0.69 against 0.96). In the other failure classes the protein chain is folded as accurately as at a correct interface, so most failures misplace a correctly folded protein rather than misfold it.

An incorrect RNA at the correct site is associated with errors in non-canonical RNA geometry. Complexes with a larger share of RNA chains in this class recover fewer non-Watson–Crick pairs and stacking interactions (*ρ* = *−*0.49 and *−*0.34), while their experimental RNAs contain more non-C3*^′^*-*endo* puckers (*ρ* = +0.24), *syn* nucleotides and non-canonical *γ* torsions, which AF3 often replaces with canonical ones (Appendix G, Supplementary Fig. S13). Mapped onto annotated protein domains, the predicted RNA-contacting residues locate the correct region but rarely respect domain boundaries, and compact single-domain RNA-binding folds are docked far better than large repeat-containing architectures (Piwi DockQ*_Nr_* 0.55, against 0.02 and 0.16 for the Dicer dimerisation and RNase III domains, respectively; Appendix O, Supplementary Fig. S29).

AF3’s confidence reflects interface failures only partly (Fig. 3D). The RNA interface confidence ipTM*_r_* falls about as much with a rising share of correct sites with an incorrect RNA as with the other failures. Per RNA chain, however, it separates this most frequent failure least well from correct placements (AUC 0.71, against 0.82–0.97). The protein-side confidence does flag it (ipTM*_p_ ρ* = *−*0.25, pLDDT*_p_ −*0.44, pLDDT*_r_ −*0.39), largely because most of these RNAs are misfolded. Distal sites lower the RNA interface confidence more than the protein one (ipTM*_r_ ρ* = *−*0.27, ipTM*_p_ −*0.17), and incorrect binding sites go with a lower pLDDT*_p_* (*−*0.28) and TM*_p_* (*−*0.33). When the main protein partner is misplaced in the assembly, both ipTM*_p_* and ipTM*_r_* decrease (*ρ* = *−*0.49 and *−*0.47). A poorly folded partner is reflected mainly in the protein confidence (pTM*_p_ −*0.38, pLDDT*_p_ −*0.37). Sampling rarely resolves the problem. Across the 49 post-cutoff targets predicted with 15 models each (134 interfaces), 71 of the 80 interfaces that fail in the default model have no acceptable RNA pose in any of the 15 models. Even the best of 15 raises the share of acceptable interfaces only from 40% to 47%. Distal sites are never recovered (0 of 29), and misposed RNAs at the correct site are only rarely recovered (4 of 26; Appendix N, Supplementary Fig. S27). Per failed interface, the correct pose is usually absent from all samples.

This failure is associated with poor recovery of non-canonical base pairs, stacking interactions and nucleotide conformations. These features help determine RNA tertiary geometry [50, 51] and remain difficult for current RNA 3D predictors [24, 52]. Since the correct pose is usually not generated at all, the limitation lies in the modelling itself rather than in ranking or sampling.

### 3.4. AF3 localises RNA motifs better than the interface geometry

On a controlled RNAcompete set, in which RNA-binding proteins were predicted with their strongest- and weakest-binding RNA probe sequences, AF3 captured relative binding preference only weakly (Appendix P, Supplementary Fig. S30). ipTM*_r_* reached an AUC of 0.54, below the 0.65 achieved by a sequence-derived RNA motif score, and it did not track measured binding intensity. Discrimination was strongest for Pumilio proteins, which recognise well-defined RNA sequence motifs through their modular PUF domains, and weakest for proteins containing RRM or KH domains, which recognise shorter and more variable motifs [53, 54] (Supplementary Fig. S31). We therefore posed a different question. As in a deposited complex the protein contacts particular stretches of the RNA and the motif atlas tells us which of those stretches carries the RBP’s known binding motif, does AF3 contact that same stretch? To test this, we scored each experimental and predicted RNA motif against curated CLIP-seq-, SELEX- and RNAcompete-derived motifs, with the motifs of other RBPs as a null, across 159 entries from 57 genes spanning both the pre- and post-cutoff sets (Section 2.5). Since CLIP-seq reports binding in cells, where accessibility and local folding shape the signal, CLIP-derived and *in vitro* motifs are scored separately.

**Fig. 4:**
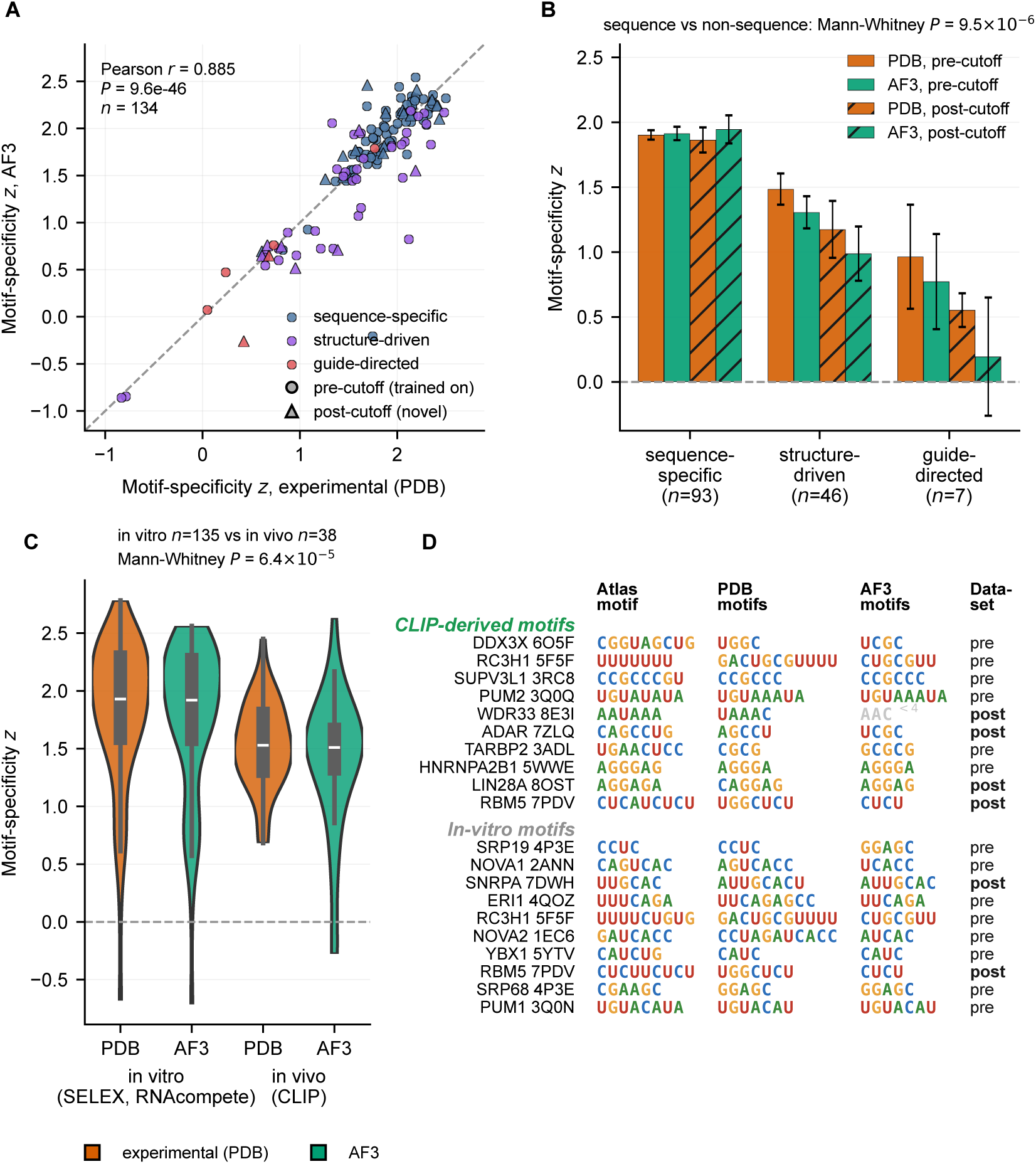
AF3 places the native RNA motif where experiments see it. **A**, Motif-specificity *z* of the AF3 RNA motif against that of the deposited structure, by RBP recognition class (circles, pre-cutoff; triangles, post-cutoff). **B**, Specificity by recognition class and PDB structure release period (orange, deposited; green, AF3; hatched, post-cutoff; mean *±* s.e.m.). **C**, Specificity against *in vitro* (SELEX, RNAcompete) and *in vivo* (CLIP) motifs. **D**, Best-matching atlas motif against the experimental and AF3 RNA motifs for the ten top-scoring genes per motif source (dimmed, contact runs shorter than four nucleotides).

Experimental interfaces match the RBP’s own motif (mean *z* = +1.70, *P* = 1.5 *×* 10*^−^*^72^, *n* = 146). For U1A/SNRPA, PUM1/PUM2, NOVA, hnRNPA1 and WDR33, the contacted nucleotides reproduce the known motif. They are read from the deposited structure alone, so the atlas plays no part in defining them. AF3 reproduces this closely (*z*_pred_ = +1.64, paired *P* = 0.17, *r* = 0.885, *n* = 134; Fig. 4A), and pre- and post-cutoff entries do not differ detectably in experimental specificity (*P* = 0.385) or in AF3’s deviation from it (*P* = 0.671). This score is sequence-based, so an AF3 motif could match the atlas equally well on a different, motif-like stretch of the same RNA, such as a repeat or a U-tract. We therefore followed the experimental stretch into the prediction by exact residue mapping. AF3’s predicted interface contacts at least half of its nucleotides in 88% of entries (*n* = 145), more than expected from its overall contact coverage (median 100% of them against 86% of all RNA positions, Wilcoxon *P* = 8.6 *×* 10*^−^*^13^; Appendix P, Supplementary Fig. S32A), and its best-scoring RNA motif lies on the native site in 80% of entries. Of the AF3 RNA motifs that match the RBP’s atlas motif (*z ≥* 1.5, *n* = 93), 88% sit on the native positions and 6% elsewhere. This also holds where the interface fails, as for entries with DockQ*_Nr_ <* 0.23 (*n* = 25) the best-scoring motif still lies on the native site in 17 of 21 (post-cutoff 11 of 13; Appendix M, Supplementary Fig. S26).

**Fig. 5:**
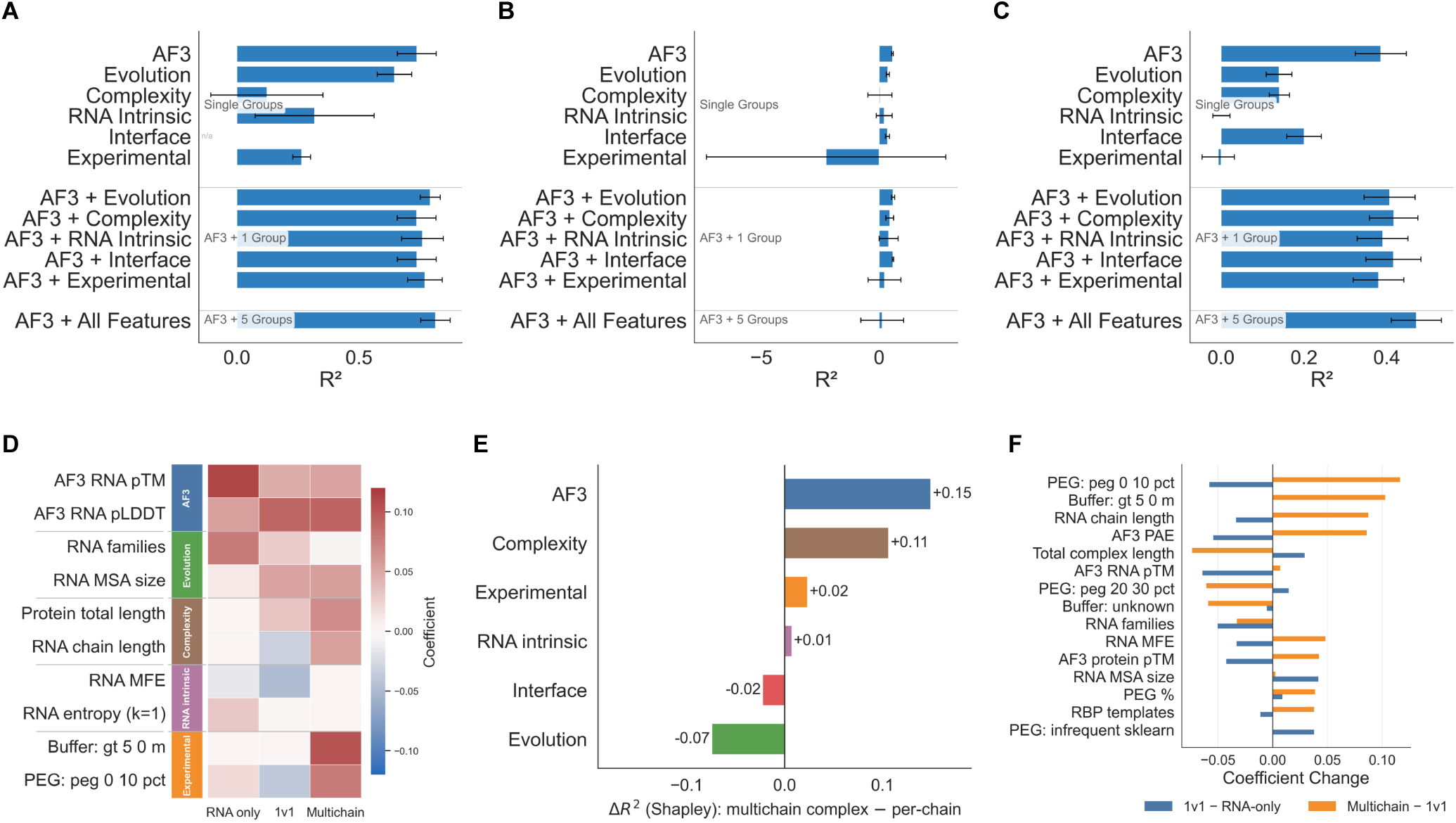
Determinants of AF3 accuracy beyond its own confidence (per-chain level). **A**–**C**, Cross-validated *R*^2^ of each feature group, of its combination with AF3 confidence and of the full model, for the TM*_r_* of single-chain RNAs (A), 1v1 complexes (B) and chains of multi-chain complexes (C). **D**, Standardised ridge coefficients of representative features across system classes. **E**, Difference in Shapley *R*^2^ between the complex-level and the per-chain model. **F**, Stepwise coefficient changes from single-chain RNAs to multi-chain complexes. Panels D and F are descriptive, and group rankings rest on the Shapley partition (Supplementary Table S11).

Entries of RBPs with predominantly sequence-driven recognition score highest, and the ordering replicates in both the pre- and post-cutoff sets (*z*_exp_ = +1.89 for sequence-specific RBPs, *n* = 93; +1.44 for structure-driven, *n* = 46; +0.85 for the guide-directed AGO2 and PIWIL1, *n* = 7; Mann–Whitney *P* = 9.5 *×* 10*^−^*^6^; Fig. 4B). Averaged per gene the classes no longer differ, because a few heavily represented genes carry the difference, whereas the motif-match and positional results do hold per gene (159 entries of 57 genes; Appendix B.3). Predicted RNA motifs agree more closely with the *in vitro* SELEX and RNAcompete motifs than with the *in vivo* CLIP motifs (Fig. 4C). This is expected, since a crystal structure and its prediction both show an isolated protein–RNA complex, whereas CLIP reports binding in living cells across the whole transcriptome, where co-binding factors and neighbouring regulatory elements shape the signal. A CLIP-derived motif may therefore reflect sequences associated with binding in cells rather than the nucleotides directly contacted by the RBP in the deposited complex. Even where the interface fails and the RNA is not in the correct pose, AF3 mostly still contacts the native motif-bearing segment, whereas on RNAcompete it barely distinguishes a preferred RNA from a weaker one.

### 3.5. What AF3 confidence misses differs across system classes

Beyond AF3’s own confidence, we asked how much of the variation in its prediction accuracy can be explained, and by which sources of information, using grouped cross-validated ridge regression over the six feature groups and an order-independent Shapley partition of the explained variance (Section 2.6). At the complex level, confidence alone explained about half of the variance in global fold quality (TM*_c_*, *R*^2^ *≈* 0.53) and adding the other groups raised it to *≈* 0.71, with 56% of the explained variance attributed to confidence and most of the remainder to structural complexity (28%) and predicted-interface features (10%). For the interface target (DockQ*_Nr_*), predicted-interface features carried the largest non-confidence share (33%). The part of TM*_c_* that confidence misses is itself explainable (residual *R*^2^ *≈* 0.26), whereas the residual of DockQ*_Nr_* is small (*≈* 0.08; Appendix R, Supplementary Fig. S35, Supplementary Table S11).

Pooling all systems masks class-specific behaviour, so we repeated the analysis per RNA chain within each system class (Fig. 5). Single-chain RNA accuracy was the most explainable setting (full model *R*^2^ *≈* 0.82, confidence alone 0.75), and evolutionary information was by far the largest complement to confidence (32% of the explained variance, against 43% for confidence). For chains of multi-chain complexes the confidence baseline was weakest (*R*^2^ *≈* 0.39) and the gains were spread over predicted-interface (19%), evolutionary (17%) and complexity (12%) features (full *R*^2^ *≈* 0.48). For single-chain protein–RNA 1v1 complexes, predictability was low and unstable, and the full model did not exceed the confidence-only baseline out of fold (Appendix S). Multi-chain RNA complexes without protein behave like the protein–RNA multi-chain class (Supplementary Fig. S38). The experimental-context group contributes little explained variance, although ligand-bound, acidic and high-PEG strata differ in accuracy (Appendix L.1, Appendix L.2, Supplementary Fig. S22).

These regressions quantify a structured signal beyond confidence without attributing it to a mechanism. AF3 confidence is the single most informative source in every setting, but it leaves a residual that is not noise, and the drivers of that residual depend on class and metric. Evolutionary signal dominates for single-chain RNAs, and predicted-interface and complexity features for multi-chain assemblies and for interface accuracy. Therefore, large, compositionally complex assemblies are systematically harder than AF3’s confidence anticipates. The cross-validated *R*^2^ values measure explained variance within the analysed PDB distribution, not out-of-sample prediction.

### 3.6. A reliability score ranks well-folded RNAs better than AF3’s pLDDT*_r_* on structures released in 2026

**Fig. 6:**
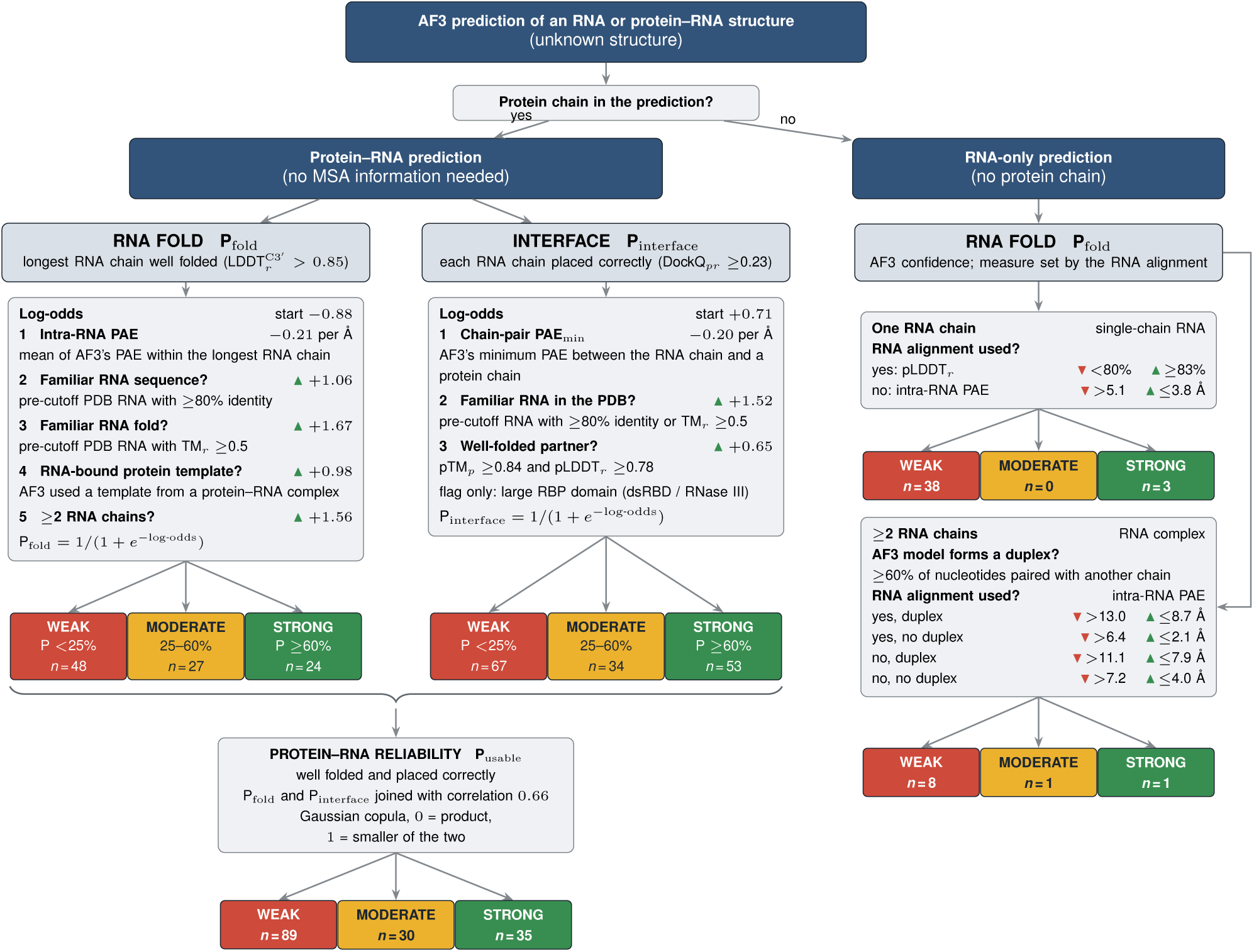
A reliability score for AF3 RNA and protein–RNA predictions. For a protein–RNA prediction (left), the probability of a well-folded RNA and the probability of a correct interface follow from AF3’s PAE and a few yes/no questions that need no experimental structure, and the two are joined into the probability that the protein–RNA model is reliable. For an RNA-only prediction (right), the probability follows from AF3’s per-residue confidence, chosen by whether an RNA alignment was used, and for RNA complexes also from whether the model forms a duplex. Each probability is WEAK below 25%, MODERATE up to 60% and STRONG above. Leaves give the number of 2026 AlphaFold Server predictions at each level, and their accuracy is shown in Appendix T (Supplementary Fig. S39 and Table S16). The coefficients (Supplementary Table S15) and the RNA-only cut-offs (Supplementary Table S16) are given in Appendix T.

The preceding sections identify signals that AF3’s confidence does not use. These are whether the RNA sequence was represented in the PDB before the training cutoff, whether AF3 used an RNA-bound protein template, how many RNA chains the assembly contains and whether the partner chains are themselves well folded. We combined them with AF3’s confidence into calibrated probabilities for the RNA fold and the protein–RNA interface (Fig. 6, Section 2.7). These probabilities can be read against the observed accuracy by system class and RNA length (Appendix T, Supplementary Table S14). That accuracy is lower throughout after the cutoff.

On 99 protein–RNA complexes released between May and September 2026, after this benchmark was assembled, the probabilities ranked outcomes better than AF3’s own measures (Appendix T.3, Supplementary Fig. S39). P_fold_ identified well-folded RNAs with an AUC of 0.89, against 0.82 for the intra-RNA PAE and 0.69 for pLDDT*_r_* (paired difference against pLDDT*_r_* +0.20, 95% CI 0.08–0.33). P_interface_ reached 0.92 against 0.89 for AF3’s minimum chain-pair PAE, a smaller gain, so at the interface our questions add little to AF3’s PAE. P_usable_ ranked the RNA chains that are both well folded and correctly placed better than AF3’s ranking score (AUC 0.91 against 0.80; paired difference +0.11, 95% CI 0.03–0.20). The levels behave as probabilities should. At STRONG, 71% of RNAs are well folded, 96% of interfaces are correct and 66% of RNA chains are usable, against 4%, 10% and 3% at WEAK, respectively. RNA-only predictions are scored from AF3’s per-residue confidence alone, chosen by whether an RNA alignment was used, and for RNA complexes also from whether the AF3 model forms a duplex (Appendix T.4, Supplementary Table S16).

The levels grade structural reliability, not binding affinity or preference among RNA sequences. The interface level barely separates preferred from weaker RNAcompete probes (AUC 0.55), and almost all of these short synthetic probes fall below the STRONG fold level, whereas 88% of the CLIP-atlas entries are graded STRONG. Within the motif-annotated set, interface quality rises with the interface level (median DockQ*_Nr_* 0.07 at WEAK, 0.27 at MODERATE and 0.83 at STRONG). AF3’s best-scoring RNA motif lies on the native site in 88% of STRONG predictions (*n* = 121), against 41–50% at the lower levels (*n* = 20 and 4; Appendix T). AF3 is less accurate on RNAs whose folds have no close match among pre-cutoff PDB structures, while its pTM*_r_* changes little. For a genuinely novel RNA, every leaf should therefore be read as an upper bound.

### 3.7. Limitations

Several limitations follow from the design and available data. The pre- and post-cutoff protein–RNA sets differ little in their RNA properties but do differ in assembly size. Post-cutoff complexes are larger, and two thirds exceed the 768-token maximum crop reported for AF3 training. Their size may therefore contribute to the complex-level accuracy gap alongside RNA novelty. The RNA-level gap, however, persists within fixed length bands (Appendix E). RNAs shorter than 20 nt could be matched to pre-cutoff RNAs by sequence only. The 2026 test cohort is dominated by cryo-EM structures and is small in the STRONG class (*n* = 53 interfaces and 35 usable-chain predictions) of the decision tree. The cross-validated *R*^2^ values quantify explained variance within the analysed PDB distribution and are not out-of-sample guarantees. The pooled regressions combine training-era and novel structures, so they identify in-distribution determinants rather than temporal generalisation. Several context analyses are sampling-limited (ions in complexes, *n* = 13; pH, *n* = 111; the no-template baseline, *n* = 6) and are read as indicative. Finally, a static single-complex prediction cannot represent crowding, concentration-dependent behaviour such as dimerisation or phase separation, or pH-dependent protonation, for which AF3 has no input channel [55].

## 4. Conclusions

The dominant post-cutoff interface failure retains contact with the native protein binding region but places an often-misfolded RNA in the wrong pose. It accounts for 54% of failed interfaces, and the RNA nucleotides contacting the protein often differ from those in the experimental structure. In the separate motif-annotated set, AF3 can still contact the native motif-bearing segment when the interface geometry is wrong. AF3 rarely generates the correct pose, and its RNA interface confidence reflects the failure only weakly. This failure and the loss of RNA fold accuracy both grow for RNAs without a pre-cutoff counterpart, while the protein fold is preserved and AF3’s RNA fold confidence does not register the loss. Poorly folded RNAs carry non-canonical pairs, stacking, C2*^′^*-*endo* sugars and *syn* glycosidic conformations that AF3 does not recover, even among structures released before its training cutoff. Protein alignments and RNA-bound protein templates act on the complex assembly, RNA alignments on the fold of single-chain and long RNAs, and supplied ions improve single-chain RNA folds. Since RNA familiarity, template use and the number of RNA chains predict these errors but are not part of AF3’s confidence, a calibrated reliability score built on them ranks well-folded RNAs better than confidence alone on complexes released after this benchmark was assembled.

These results point to different limits for different RNAs. For long RNAs and for RNAs without a protein partner, evolutionary signal is the binding constraint. RNA-native alignments and cross-family language models are the most direct lever there, together with explicit terms for the stacking, non-canonical pairs and backbone conformers that a single-structure target smooths away [16, 23, 50, 56, 57]. For short protein-bound RNAs, whose fold the protein largely dictates, alignments add little, and the limit is the pose of the RNA at the protein site. Many RNAs populate dynamic ensembles of interconverting conformations rather than a single structure, so ensemble-aware prediction and evaluation, which AF3 does not offer, would also be needed to capture RNA functions such as catalysis, RNA splicing and protein binding. For protein–RNA complexes, the limit is placement rather than recognition. The levers are training objectives that score interface geometry directly, better use of templates from RNA-bound proteins, predicted ensembles rather than single states, and inputs for the ions and experimental conditions the model currently ignores [7, 58, 59]. Until then, the reliability of an AF3 model of a novel RNA should be read from the familiarity of the RNA and the information available to the model, not from its confidence alone.

We focused on AF3 because it ranks among the leading methods in recent benchmarks [24, 27] and is one of the most widely used models for RNA and protein–RNA structural studies. The evaluation pipeline built for this study is, however, not specific to AF3. It takes predicted and deposited structures, maps every chain and residue exactly, and scores RNA folds, protein–RNA and RNA–RNA interfaces, placement classes and nucleotide-level conformations for protein–RNA complexes, single-chain RNAs and multi-chain RNA complexes. Other predictors of nucleic-acid complexes, such as Boltz-1, Chai-1, HelixFold3 and RoseTTAFoldNA [20, 60–62], can therefore be benchmarked in the same way, with only the reading of their confidence outputs to adapt. The databases already record DNA chains, so the framework can also be extended to protein–DNA complexes.

## Data and code availability

The evaluation pipeline, the regression and variance-partition analyses, the decision-tree package, the figure scripts and a notebook that reproduces the regression results with one command are available open-source at https://github.com/automl/RNA-Protein-3D-Structure-Evaluation under the Apache 2.0 license. The version of the code used for this manuscript is archived on Zenodo (DOI: 10.5281/zenodo.23890212). The SQLite databases of all evaluated structures and predictions, the curated per-chain and complex-level feature tables and the intermediate analysis tables are available on Zenodo (DOI: 10.5281/zenodo.23870599); with them, all figures and tables of the paper are regenerated by reproduce_all.sh. All cross-validation used a fixed seed, and all configurations are written to disk.

## Supplementary Data

Preprint. Supplementary data is attached to the main manuscript.

## Author Contributions

I.K.T: Conceptualization, Data curation, Formal analysis, Investigation, Methodology, Software, Visualization, Validation, Writing - original draft, Writing - review & editing. M.U.: Conceptualization, Data curation, Validation, Methodology, Writing - review & editing. J.K.H.F: Conceptualization, Writing - review & editing. R.B.: Funding acquisition, Writing - review & editing. F.R.: Project administration, Conceptualization, Supervision, Resources, Formal analysis, Investigation, Methodology, Validation, Writing - original draft, Writing - review & editing. All authors have read and approved the final manuscript.

## A. Extended experimental and computational methods

This appendix provides the full detail of the data-processing and feature-extraction pipeline summarised in the main-text Materials and Methods.

### A.1. Dataset composition

This appendix gives a descriptive overview of the benchmark introduced in Section 2.3 and the Datasets methods, so that the prediction accuracy results can be read against the composition of the data. Figure S1 summarises what the dataset contains. The protein–RNA systems divide into a minority of single (1v1) complexes and a majority of larger multi-chain assemblies, analysed alongside the single-chain RNA folds (panel A). Structures are predominantly X-ray crystallography with a substantial cryo-EM minority (panel B). The RNA chains are mostly short (median 14 nt for protein–RNA complexes), with a long tail of large ribosomal and intron RNAs, whereas the single-chain RNA folds are longer (median 71 nt; panel C). The bound proteins span a broad size range (median 690 residues; panel D). Ions and ligands recorded in the experimental solution are dominated by sodium and magnesium, with potassium, manganese and polyamines less frequent (panel E). Only a small fraction of RNA chains carry an Rfam family annotation, tRNA being by far the most common, followed by ribosomal and riboswitch families (panel F).

Figure S2 summarises the evolutionary and experimental inputs. Protein multiple sequence alignments are deep (median of order 10^3.5^ sequences), whereas the large majority of RNA chains have essentially no alignment, so RNA evolutionary signal is scarce across the benchmark (panel A). Most complexes have at least one protein or RNA-binding-protein structural template available (panel B). Recorded experimental temperatures cluster at room temperature with a distinct cold-room subset (panel C). Finally, splitting by the AF3 training cutoff shows that the pre- and post-cutoff sets are closely matched in these inputs, differing only modestly in protein MSA depth, with RNA alignments essentially absent in both eras (panel D). This complements the family-, length- and method-matched comparison in Fig. S5.

### A.2. Dataset attrition and variation in sample sizes

The RNA-containing corpus was assembled from the PDB in two date-split batches relative to AF3’s training cutoff and totalled approximately 5,700 downloaded complexes, 4,738 protein–RNA (*b2021* with 1,959 X-ray and 966 cryo-EM; *a2021* with 281 X-ray and 1,532 cryo-EM) and 1,010 single-chain RNA (*b2021* with 736 X-ray and 4 cryo-EM; *a2021* with 204 X-ray and 66 cryo-EM), from a PDB search already restricted to a resolution of *≤* 4.5 Å, each retained only if it contained at least one protein of *≥* 20 residues and one RNA of *≥* 4 nucleotides.

**Fig. S1:**
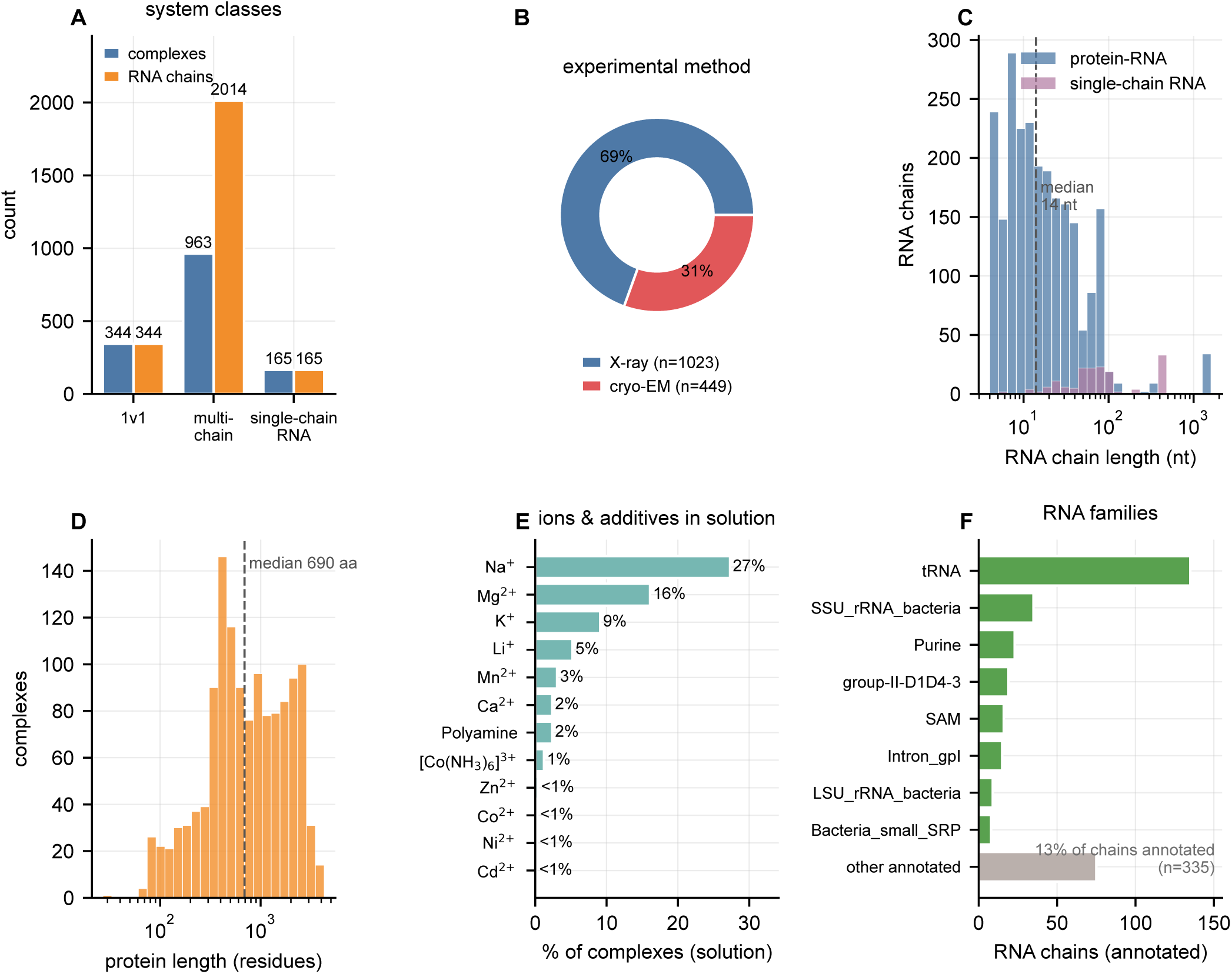
Composition of the benchmark. **A**, Complexes and RNA chains per system class. **B**, Experimental method. **C**, RNA chain length (log scale) for protein–RNA complexes and single-chain RNAs (dashed line, protein–RNA median). **D**, Protein length (log scale). **E**, Most common ions and additives recorded in the experimental solution, as a percentage of complexes (polyamine combines spermidine and spermine). **F**, RNA family composition among the annotated chains.

After the exclusions listed below, 1,307 protein–RNA complexes (923 of the 2,925 downloaded before and 384 of the 1,813 downloaded after the AF3 training cutoff), 165 single-chain RNAs (99 of 740 and 66 of 270) and 106 multi-chain RNA complexes without protein (86 and 20) were analysed. The independent 2026 test set followed the same filters. Of 301 cryo-EM protein–RNA entries released between May and September 2026, 120 were excluded for modified residues or ligands, 76 for discontinuous RNA chains and 9 for size, which left 99 complexes including 3 X-ray entries (Appendix T.3). Different analyses require different combinations of computed quantities, and a sample enters an analysis only if every input it requires was computed and passed quality control. Each figure and statistic is therefore reported on the *n* for which its inputs were available. No sequence-length cutoff and no redundancy filtering were applied.

The main sources of data attrition were the following.

1. **Broken or discontinuous chains.** RNA chain breaks in the deposited PDB entry are common and preclude reliable per-chain alignment and scoring. We flagged breaks geometrically from the distance between the phosphate atoms of consecutive nucleotides, a separation above *∼*10 Å indicating a chain break, with borderline cases (10–12 Å) confirmed by manual inspection, and excluded complexes whose RNA chains contained such breaks. The check was later automated, and the automated filter reproduces the selection made by manual inspection.
2. **Duplexes deposited as a single strand.** Where an RNA’s complex is an intermolecular duplex but only one strand is present in the deposited PDB coordinates. The AF3 input built from those coordinates is a lone strand. The model then folds a monomer against an experiment that resolved half of a dimer, satisfying its base pairing intramolecularly, and the two objects are no longer comparable. Reconstructing the partner would require submitting a duplex where the deposition provides only a single strand, so these entries were excluded.
3. **Irreconcilable chain identifiers or residue numbering.** In PDB deposited mmCIF files frequently use multi-character or non-standard chain labels and inconsistent residue indexing (mismatched author and label sequence identifiers, or insertion codes) that break the one-to-one mapping between a deposition and its AF3 prediction required for per-chain scoring. Most were remediated automatically by renaming chains to single-character identifiers and renumbering residues along the label sequence without insertion codes. Assemblies that still could not be reconciled, for example because they have more chains than single-character identifiers allow, were excluded.
4. **Resource limits for the largest assemblies.** AF3 could not be run within a four-day wall-clock limit for a subset of very large, ribosome-scale complexes, because MSA search, which can dominate AF3 runtime and reach hundreds of gigabytes of memory even for a single long (*∼*1500 nt) RNA, exceeded available compute even when the search was separated and run with 2 TB of memory. Long-RNA predictions are independently reported to require hours per target and to fail often enough that repeated runs are needed [24, 32]. These exclusions are an infrastructure limit of the MSA-search stage rather than a modelling failure, but they act only on the upper size tail of the distribution and fall predominantly on the largest multi-chain assemblies. The length- and class-dependent accuracy trends we report should be read as applying to the size range actually analysable at scale.
5. **A poly-nucleotide handling issue in OpenStructure.** OpenStructure leaves some chains unmapped, mostly short poly-U RNAs. The affected interfaces and complexes were rescored on the exact residue mapping, as described in Section B.
6. **Missing or unmappable experimental descriptors.** Experimental context (ions, ligands, buffer, pH, temperature, PEG) is recorded as free text of variable completeness. We parsed these fields into distinct descriptor groups where possible, but some entries lacked the relevant fields or could not be mapped to a defined category even after manual inspection, and were omitted from the analyses that depend on those descriptors.
7. **Evaluation-level failures.** The feature- and metric-extraction pipeline computes many quantities per complex, and individual computations can fail independently for structure-specific reasons. For example, when the residue numbering of a chain is not consecutive and contains a non-numerical character. Wherever a required metric could not be computed reliably, that value was dropped and the sample was excluded from the specific analysis using it. We consistently favoured exclusion over retaining values of uncertain quality. The small variation in the number of analysed targets (*n*) across analyses is therefore expected and is noted per result.

**Fig. S2:**
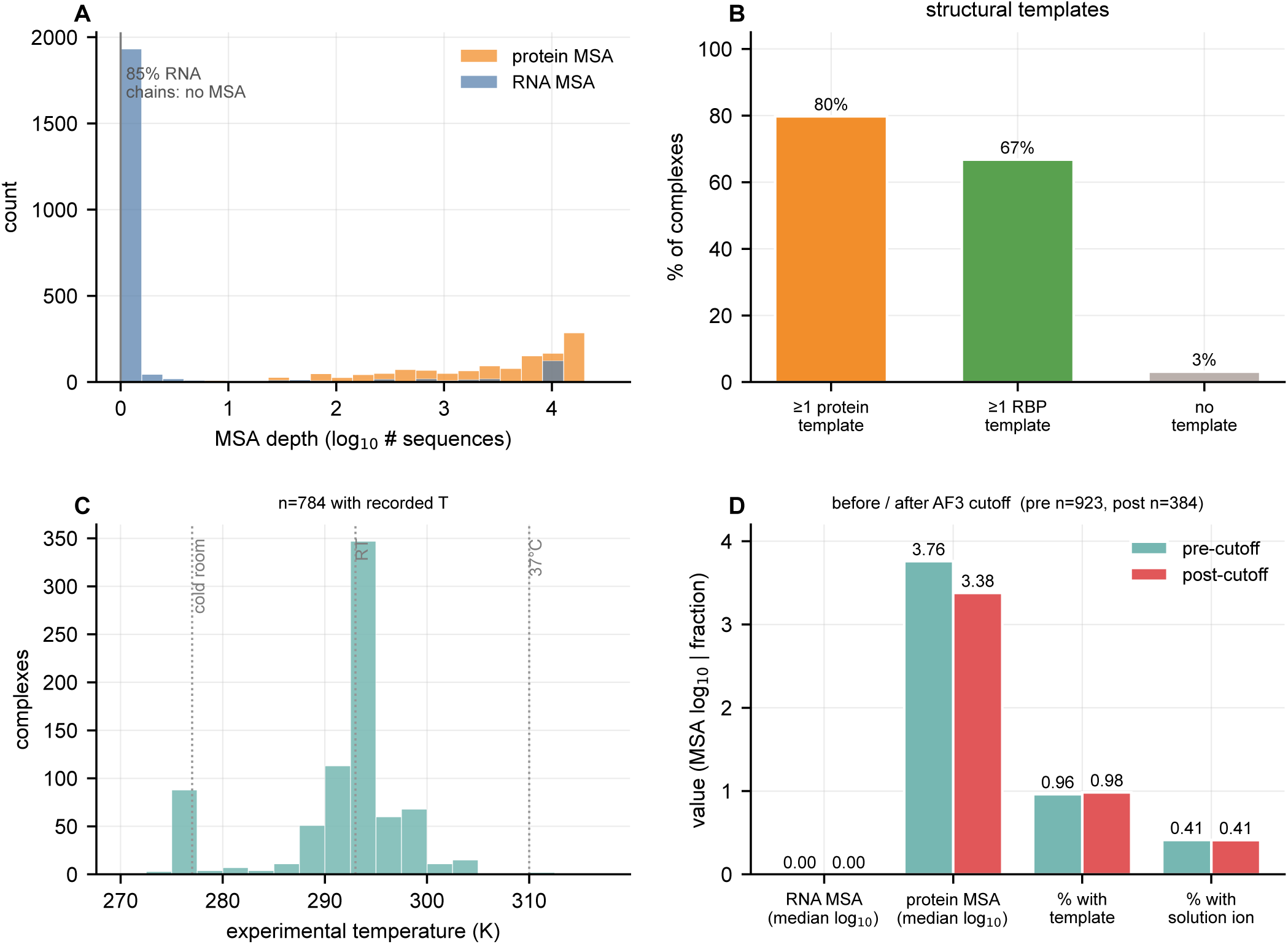
Evolutionary and experimental covariates. **A**, Depth of the protein and RNA alignments (log_10_sequences). Protein alignments are deep, whereas most RNA chains have essentially none. **B**, Complexes with a protein-only template, an RNA-bound protein template or none. **C**, Recorded experimental temperature. **D**, Alignment depth, template availability and recorded solution ions before (b2021) and after (a2021) the AF3 training cutoff.

### A.3. Data organisation, configuration, and database schema

All analyses were performed using a modular Python pipeline driven by a central configuration file (`database/config.json`). This configuration defines the dataset `classification` (e.g. `protein_rna`), base data directory and SQLite database path. The pipeline enforces a standardised directory layout in which experimentally determined structures (mmCIF/PDB) are stored under `parent_folder/pdb`, and predicted structures (e.g. AF3 outputs) under `parent_folder/af`.

All parsed metadata and computed features were stored in paired relational tables, `exp_“classification“` and `pred_“classification”`, within a single SQLite database. The experimental table contains one row per PDB entry, while the prediction table stores one row per model (indexed by `FileName`) and references the experimental entry via `exp_db_id`. This schema enables efficient joining of prediction metrics with ground truth annotations.

The execution driver (`mainExecution.py`) initialises a `StartConfig` object to resolve all paths, table names and runtime parameters, ensuring full reproducibility and eliminating hard-coded dependencies.

### A.4. Structure and polymer parsing

Experimental structures were parsed using `parsePDBCIF2DB.py`, which leverages `Bio.PDB` [63] and `MMCIF2Dict` to extract structured information from mmCIF files. Polymer entities were identified using the `_entity_poly.type` field and classified into protein, RNA or DNA categories.

Predicted structures were processed analogously using parseAFcif2DB.py, with additional parsing of AF3-specific metadata such as model indices, seeds and file naming conventions. Each prediction entry was linked to its corresponding experimental structure using `exp_db_id`, enabling direct pairwise evaluation.

### A.5. Intermolecular interaction detection

Intermolecular interactions were computed using `extractInteractions.py`, which integrates PyMOL [64] structural handling with Biopython-based parsing and explicit geometric rules. Prior to analysis, all structures (experimental and predicted) were preprocessed to remove solvent molecules, ions, and non-polymeric components, ensuring that only protein and nucleic acid atoms contributed to interaction detection. Atoms were classified into chemically meaningful categories, including protein backbone and side chain atoms, and RNA base, sugar, and phosphate groups. This classification enabled context-aware interaction assignment and downstream feature aggregation.

Hydrogen bonds were identified using geometric constraints inspired by HBPLUS [65]. Because mmCIF structures derived from both experimental data and AF3 predictions lack explicit hydrogen atoms, hydrogen bond detection relied on inferred donor–acceptor geometry. Candidate interactions were first screened using a permissive cutoff of 3.5 Å between donor and acceptor atoms. For final classification, stricter criteria were applied, requiring donor–acceptor distances *≤* 3.2 Å and an approximate donor–hydrogen–acceptor (D–H–A) angle *≥* 150*^◦^*, inferred from heavy-atom positioning. These thresholds were chosen to approximate energetically favourable hydrogen bonds and were validated against canonical HBPLUS criteria (H–A distance *<* 2.7 Å, D–A distance *<* 3.35 Å, D–H–A angle *>* 150*^◦^*), ensuring consistency with established structural biology standards.

Van der Waals contacts were defined as all non-hydrogen-bond interactions between atoms within a distance cutoff of 3.9 Å. These contacts capture weak, non-specific interactions that contribute to interface stability and complement hydrogen bonding patterns.

All detected interactions were aggregated at the chain-pair level, producing structured descriptors that encode both quantitative and spatial information. These include atom-level contact lists, total hydrogen bond counts, van der Waals contact counts, and residue-level interaction motifs representing contiguous regions of interaction across the interface. Interaction data were stored as JSON-encoded fields in the database, preserving detailed mappings between interacting residues and chains for downstream analysis.

### A.6. Chain and residue mapping

Correspondence between predicted and reference polymer chains was established with the OpenStructure chain mapper, which aligns chains by sequence rather than by chain label. Residues were indexed sequentially within each chain before scoring, so differences in numbering between experimental and predicted structures were not mistaken for structural differences. Short peptide and nucleotide chains were retained. Interface scoring used this sequence-based mapping without enforcing residue numbering, which maximised the number of mapped protein and RNA chains and hence of scorable interfaces. Sequence-based chain mapping is unreliable for complexes with several identical or near-identical RNA copies. The per-chain RNA LDDT was therefore computed independently of the global mapping. Each predicted RNA chain was compared with every reference RNA chain of matching length (and matching sequence where available) by single-chain LDDT*_r_*, and the best-scoring reference was kept for that chain (`RNA_LDDT_AllChains`). This gives one value per predicted RNA chain that does not depend on chain-mapping failures.

Residue-level analyses (contact match, interface placement and the fallback scores below) need the predicted counterpart of each experimental residue. This exact residue mapping follows the mapping OpenStructure used to score the complex. The chain mapping of `compare-structures` assigns each experimental chain to a predicted chain, and its reference–model sequence alignment assigns each modelled experimental residue to its position in the predicted chain. Experimental residues were located in the deposited mmCIF by chain and residue number and converted to the label chain identifiers used by OpenStructure. OpenStructure cannot map poly-U and U-repeat RNA chains and omits very short chains (Appendix A.2). Each chain it leaves unmapped was paired with the remaining predicted chain of highest sequence identity (*≥* 90%; among identical copies, the copy with the largest contact overlap), and its residues were mapped by global alignment to the full predicted sequence. This fallback covered 188 post- and 596 pre-cutoff interfaces, and only 5 and 17 interfaces remained unmappable. No offset or sliding tolerance is applied. As a check, 99.9% of the mapped experimental contact residues carry the same residue type as the predicted residue they are mapped to. The mapping is implemented in `evaluation/exact_residue_mapping.py`.

### A.7. AA and RNA motif evaluation

RNAMotif is defined per interface between a polynucleotide chain and its interaction partner and is obtained by first identifying all relevant chain pairs from the mmCIF structure. If proteins are present, every RNA chain is paired with every peptide chain (Cartesian product after expanding strand IDs), otherwise RNA–RNA pairs are formed. For each such pair, PyMOL is used to detect hydrogen-bond and van der Waals contacts, and the RNA motif is then constructed from all RNA nucleotides participating in these interactions. Specifically, the code collects the unique nucleotide identifiers (residue name and index), sorts them by residue index, and concatenates them into a comma-separated `RNAMotif` string. If the partner is not a protein (RNA–RNA case), a similar procedure is applied using base-to-base interaction extraction. The result is that `RNAMotif` represents the set of contacting nucleotides on the RNA side for each individual interface, and multiple motifs can exist per structure (one for each RNA–partner pair) rather than being limited to a single RNA chain.

RNAmotif_score is computed by first parsing the RNAMotif field into a list of motif strings, each corresponding to a specific protein–RNA (or RNA–RNA) interface aligned with the chain pairs. These motifs are then grouped by RNA chain, and for each chain, all motif strings associated with it are merged into a single “combined motif” by collecting the interacting nucleotides, removing duplicates, sorting them by residue index, and formatting them as a comma-separated sequence. For scoring, each reference RNA chain is paired with the corresponding predicted RNA chain (based on index order), and their combined motifs are compared using a global sequence alignment (PairwiseAligner of Biopython [63]) on gap-filled motif representations. The resulting alignment score is then normalised by the length of the longer motif, producing a per-chain similarity score. If multiple RNA chains are present, this process is repeated for each aligned chain pair, and the final RNAmotif_score is the average of these normalised scores across all chains.

AAmatch_score and RNAmatch_score measure whether AF3 contacts the same protein residues and RNA nucleotides as the experimental structure, residue by residue. Interface residues are the hydrogen-bond and van der Waals contacts defined above (`AAMotif, RNAMotif`), recorded separately for each protein–RNA chain pair listed in `ChainIDpairList_proteinRNA`. Experimental contact residues were carried to their predicted counterparts by the exact residue mapping (Section A.6). For each interface, the protein (nucleotide) score is the Jaccard index *|P ∩ E|/|P ∪ E|* of the predicted contact set *P* and the mapped experimental contact set *E*. The complex-level `AAmatch_score and RNAmatch_score` pool all protein–RNA interfaces of the complex, Σ*|P ∩ E|/*Σ *|P ∪ E|*. Native interfaces that AF3 does not form therefore count as unrecovered, and interfaces formed only by AF3 count as false contacts.

### A.8. Interface surface area and binding affinity estimation

Interface surface areas were computed using extractInterfaceArea.py, which implements a workflow analogous to classical tools such as NACCESS, but adapted for modern mmCIF-based structural pipelines. Structures were converted into temporary PDB representations, and atomic coordinates were processed using the FreeSASA library [66], which calculates solvent-accessible surface areas (SASA) using a rolling probe model (default probe radius of 1.4 Å and standard atomic radii).

For each complex, SASA values were computed for the intact assembly and for individual subunits (protein and RNA chains). The interface area (IA) was then defined as the solvent-accessible surface buried upon complex formation,

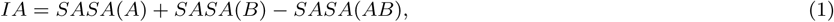

where *SASA*(*A*) and *SASA*(*B*) correspond to the isolated subunits and *SASA*(*AB*) to the complex. This formulation captures the total surface area buried at the interface and serves as a proxy for interaction strength and structural complementarity.

Interface atoms were defined as atoms that exhibit a reduction in solvent accessibility upon complex formation. Residues containing at least one such atom were classified as interface residues. For multi-chain systems, chain grouping strategies were applied to isolate specific interaction types (e.g., protein–RNA, RNA–RNA), ensuring accurate attribution of interface contributions.

Additional interface descriptors were computed to characterise the physicochemical properties of the interaction surface. These include the interface-to-surface ratio (*R_i/s_*), defined as the ratio of interface area to the total accessible surface area excluding the interface, as well as amino acid composition (AAC) and amino acid propensity (AAP) at the interface. These quantities were derived from atom-level coordinates extracted via PyMOL and provide insights into residue preferences and interaction specificity.

Binding affinity was estimated using `estimateBindingAffinity.py`, which integrates structural descriptors into a heuristic scoring model. The model combines hydrogen bond counts, van der Waals contacts and interface area using empirically derived weights informed by protein–RNA interaction studies [67, 68], with additional terms for electrostatic contributions and RNA-specific properties. The resulting free energy estimate (Δ*G*) was converted to a dissociation constant (*K_d_*),

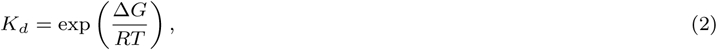

where *R* is the gas constant and *T* is temperature. This provides a relative measure of binding strength rather than an absolute thermodynamic quantity.

### A.9. Global structure alignment and evaluation

#### A.9.1. Structure preprocessing and alignment

For the 3D structural evaluation of RNA and protein–RNA complexes, we used PyMOL [64] (version 3.1.3) for structural alignment and coordinate extraction. All complexes were cleaned by removing solvent molecules, ions, ligands and other non-polymeric components (`remove solvent, organic`), and were subsequently separated. The predicted structures of the extracted protein–RNA and single-chain RNAs were then aligned to their corresponding crystallographic reference structures using *C^α^* atoms for proteins (`polymer.protein` `and name CA`) and phosphorus (P) atoms for RNA (`polymer.nucleic and name P`). Alignment was performed using PyMOL’s align function, which applies iterative least-squares fitting to minimise RMSD. The aligned structures were saved in separate PDB files. These files were then used to calculate both the RNA and protein–RNA complex RMSD and TM values.

#### A.9.2. RMSD computation

Following structural superposition in PyMOL, RMSD was computed as the Root-Mean-Square Deviation of aligned backbone atoms providing a direct measure of coordinate accuracy in Ångströms. In Cartesian coordinates, the root-mean-square deviation (RMSD) between predicted and reference conformations of a molecule is defined as

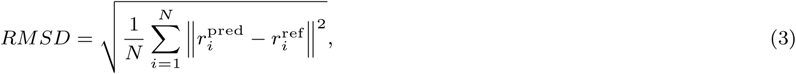

where (N) is the number of backbone atoms included in the calculation, and 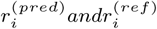 are the Cartesian coordinates of atom (i) in the predicted and reference conformations, respectively. The RMSD is minimised by optimal rigid-body superposition of the two structures.

#### A.9.3. TM-score calculation

The TM-score, which is based on global superposition and uses backbone coordinates (*C^α^*atoms for proteins and C3’ atoms for nucleic acids), was calculated using the USalign tool [38] (version 20241108) with RNA-specific parameters (`-mol RNA -mm 0 -ter 2`) for RNA structures and protein-specific parameters (`-mol prot -mm 0 -ter 2`) for protein moieties, providing a normalised, length-independent metric that, unlike RMSD, is less sensitive to local errors and focuses on the overall topology of the structures [69].

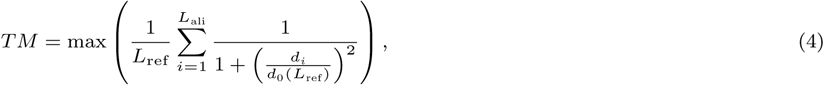

where *L*_ref_ is the length of the reference structure, *L*_ali_ is the number of aligned residues between the predicted and reference structures, *d_i_* is the distance between the *i*-th pair of aligned residues and *d*_0_(*L*_ref_) is a scaling factor to ensure the score of random RNA pairs is independent of RNA sequence length 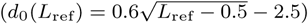.

### A.10. Approximate local accuracy metrics

All-atom LDDT scores were computed using OpenStructure [70] via its `compare-structures` interface, providing a superposition-independent measure of structural similarity based on heavy-atom distance differences. Scores were evaluated separately for the protein, the RNA and the complete complex, so that each molecular component of every ribonucleoprotein was assessed independently. The Local Distance Difference Test score [70] was computed by comparing inter-atomic distances between predicted and reference structures with a 15 Å inclusion radius and the standard LDDT *δ* thresholds (0.5, 1.0, 2.0 and 4.0 Å) to capture varying levels of distance accuracy.

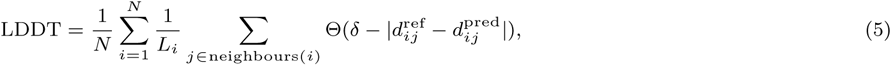

where *N* is the total number of atoms considered, *L_i_* is the number of neighbouring atoms of atom *i* within the distance cutoff, 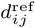 is the distance between atoms *i* and *j* in the reference structure, 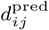 is the distance between atoms *i* and *j* in the predicted structure, Θ is a step function that returns 1 if 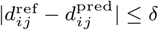, and 0 otherwise, and the four *δ* thresholds are averaged to yield the final score.

#### Backbone LDDT

The backbone LDDT is our own implementation (`calculate_complex_lddt in evaluation/extractEvalMetrics.py`) of the global LDDT score [70], as in AlphaFold [12]. It uses one atom per residue, the C3*^′^* atom of each nucleotide and the *C^α^* atom of each amino acid. For 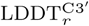, the longest RNA chain of the experimental structure and the corresponding chain of the model are written to separate files, and their atoms are paired in chain order. All atom pairs closer than 15 Å in the experimental chain are scored, and a pair counts as preserved at threshold *δ* when its distance in the model differs by less than *δ*. The score is the fraction of preserved pairs, pooled over all pairs rather than averaged per atom as in Eq. (5), and averaged over the thresholds of 0.5, 1, 2 and 4 Å. No superposition and no stereochemical checks are applied. 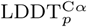 and the backbone LDDT*_c_* are computed in the same way on the protein and on the whole complex. For the reliability score, the same function is applied to each predicted RNA chain against the experimental RNA chain of the same sequence, the best of identical copies (Appendix T). The values are stored as `RNA_LDDT, Protein_LDDT and Complex_LDDT`.

#### Which LDDT is used where

The backbone LDDT is used for the fold classes i–iv (Section 2.3, Appendix F), the RNA structural classes (Appendix G) and the reliability score (Appendix T). The all-atom LDDT is used for the mean accuracies before and after the cutoff, the comparisons with similar pre-cutoff targets (Appendix E), the input ablation, the confidence analyses and the descriptor ranking, and for the fold of the individual RNA and protein chains in the placement analysis (Appendix N). There, an RNA chain is well folded at an all-atom LDDT*_r_ >* 0.85 (with TM*_r_ >* 0.5 from 60 nt), intermediate at 0.75–0.85 and misfolded at *≤* 0.75.

## B. Interface accuracy evaluation

Interface quality was evaluated using OpenStructure [70] through the compare-structures command-line interface. This workflow compares predicted and reference complexes at the atomic level and computes multiple complementary metrics.

DockQ [40] integrates interface RMSD, ligand RMSD and the fraction of native contacts into a single normalised score, providing a robust measure of docking accuracy. Interface LDDT (iLDDT) evaluates local structural agreement specifically at the interface, based on deviations in interatomic distances. Additional metrics include the QS-score (global interface similarity) and the Interface Contact Similarity (ICS), which quantifies the overlap between predicted and reference contact sets. Together these metrics capture different aspects of interface accuracy, geometry, contact recovery and topological agreement.

Interface metrics were resolved per protein–RNA interface rather than only at the level of the whole complex, so that each protein–RNA contact of a multi-chain assembly was scored separately and could be filtered to the RNA of interest. For every complex, a single compare-structures run over the full reference and predicted structures established the reference-to-model chain mapping and reported the per-interface Interface Contact Similarity. Each protein–RNA interface was then identified from the reference contact interfaces, mapped to the corresponding predicted chains, and indexed by the predicted protein–RNA chain pair. Because OpenStructure reports interface LDDT only as a single global value, per-interface iLDDT was obtained by isolating each protein–RNA interface, extracting the two participating chains and re-scoring them, so that the global iLDDT of the two-chain sub-structure corresponds to that individual interface. Where OpenStructure could not map all chains, the exact residue mapping was supplied instead (Section A.6). DockQ*_pr_* was computed with OpenStructure’s own DockQ implementation, and iLDDT, ICS and QS-score with its scorer, given the chain correspondence explicitly. The whole complex was rescored in the same way with all mapped chains, which gives its lDDT, iLDDT, ICS, QS-score and DockQ*_c_* (the interface-size-weighted mean that OpenStructure reports). Where OpenStructure maps all chains, the two agree. Per interface, DockQ*_pr_* agrees over 1,097 interfaces (Spearman *ρ* = 0.998, 99.9% on the same side of DockQ 0.23), iLDDT and ICS over 1,093 and 1,095 (*ρ* = 0.999) and QS-score over 1,095 (*ρ* = 0.988). Per complex, the five scores agree over 96 complexes (*ρ* = 0.95–0.998, 90th percentile of the absolute difference *≤* 0.007). The fallback scored 744 of the 748 missing interfaces and 545 of the 561 complexes with a missing or unmapped interface. For the other 16, OpenStructure’s scorer rejected the chain correspondence, and they keep OpenStructure’s values. The new values replace those of interfaces and complexes with an unmapped chain and fill the missing ones. All pre-cutoff per-interface iLDDT and ICS values were recomputed this way. The same per-interface DockQ was applied to the 15 seed and model samples (Appendix C) and to the 2026 test set (Appendix T.3).

Per-interface docking scores were additionally obtained through the SWISS-MODEL OpenStructure Actions API [70], which returns DockQ, QS-score, the fraction of native contacts (fnat) and the interface RMSD for each mapped chain pair. These per-interface values (indexed by chain pair, as for iLDDT and ICS) were retained alongside the local scores to provide an independent, contact-based assessment of protein–RNA docking accuracy.

### B.1. Interface placement

To locate the RNA of each native protein–RNA interface relative to its protein partner, the predicted protein chain was superposed on the experimental one using the *C^α^* atoms paired by the exact residue mapping, with iterative outlier rejection as in PyMOL align (five cycles, atoms deviating by more than twice the RMSD removed). In this frame we measured the displacement of the RNA centroid (translation; C4*^′^* atoms, P where C4*^′^* is missing) and the rotation still needed to superpose the RNA onto the native RNA. We also measured the RMSD of the native protein contact residues (pocket RMSD), which tests whether the protein’s binding region is itself predicted in place. The ligand RMSD (LRMSD) was taken from the OpenStructure calculation. Where OpenStructure provides none (unmapped chains, A.6), it was computed in the same frame. Where both exist, they agree (Spearman *ρ* = 0.95, median difference 0.25 Å). Each interface was then assigned to the first matching class. (i) Correct interface, DockQ*_pr_ ≥* 0.23. (ii) Correct site, incorrect RNA, if AF3 still contacts native protein residues (per-interface protein contact match *≥* 0.1). (iii) Incorrect binding site, if the protein’s binding region is mispredicted (pocket RMSD *>* 4 Å). (iv) Otherwise by the RNA translation, as correct site (*≤* 10 Å), as a rotated RNA if rotated by more than 45*^◦^*, shifted site (10-20 Å), or distal site (*>* 20 Å). (v) A failed duplex strand (at least four base pairs with another RNA chain) whose predicted duplex covers at least half of the native duplex (C4*^′^* atoms within 4 Å) while the strand itself covers less than half of its native position is placed by the whole duplex instead of step (iv). The duplex is the strand and the part of its partner within 20 Å, restricted to nucleotides present in both structures. A duplex moved by at most 10 Å with its helix axis tilted by at most 45*^◦^* is a correct site with a slid duplex, and otherwise the duplex shift decides between rotated RNA, shifted and distal site as in (iv) (`evaluation/apply_duplex_pose.py`).

The 10 Å and 4 Å cut-offs are the 95th percentiles of the translation and pocket RMSD of correct interfaces. Where OpenStructure reports no DockQ*_pr_* for an interface, the DockQ*_pr_* computed on the exact residue mapping is used (A.6). The analysis is implemented in `evaluation/interface_rna_pose.py`. The class of every protein–RNA interface is stored as RNA_interface_placement. The per-RNA-chain summaries by main partner are derived from it by `publication_figures/placement_per_rna_chain.py`, and the decision-tree labels take the best class over an RNA chain’s interfaces (`evaluation/placement_labels.py`). To test whether the classes depend on these cut-offs, each threshold was varied while the others were kept fixed (protein contact match 0.05-0.25, pocket RMSD 3-6 Å, correct-site translation 7.5-15 Å, shifted-site limit 15-30 Å, rotation 30-90*^◦^*), and all five were varied jointly over all 1,280 combinations of these classification thresholds (5 *×* 4^4^). DockQ*_pr_ ≥* 0.23 defines success and was not varied (`evaluation/placement_threshold_sensitivity.py`). The protein contact match threshold has the largest effect, as correct site, incorrect RNA falls from 60% to 42% of post-cutoff failures between 0.05 and 0.25. It nevertheless remains the largest failure class in all 1,280 settings.

### B.2. RNACompete Analysis

For RNACompete analyses, we first compute and write RNA motif overlap metrics (`RNAmotif_score, RNAMotif_inExp`, and the 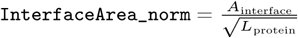) into the database. Next, maxprobe vs minprobe distributions (mean, median, Cohen’s d, AUC, Mann–Whitney) are compared and we test whether predicted scores rank maxprobes above minprobes using per-protein and global AUC, effect sizes and Wilcoxon tests, logistic regression of AF3/template deltas on non-AF3 outcomes, RNA-length-controlled subsets, margin-based correlations with experimental binding and agreement with RNAcompete intensity/z-score (the binding intensity of a probe standardised over all probes of the same protein). The script `domain_analysis.py` links amino-acid interface residues (AAMotif) to annotated RNA-binding domains (`domain_pos`; Appendix O), classifies each complex by domain coverage (within/partial/outside), and asks whether that coverage and domain family (RRM, KH, zinc finger, etc.) explain probe-class differences and which method separates maxprobes from minprobes best.

### B.3. RBP motif-atlas comparison of interface RNA motifs

This analysis tests whether the RNA nucleotides contacted at an interface match the sequence that the RNA-binding protein prefers in independent binding assays, both in the deposited structure and in its AF3 prediction. Code and derived tables are under clip/ in the repository, and clip/scripts/run_all.sh reproduces every number and both figures.

#### Interface RNA motifs

The `RNAMotif` column of `exp_protein_rna` and `pred_protein_rna` is not a sequence motif but the set of RNA residues contacting protein, stored as one list entry per chain pair of `ChainIDpairList_proteinRNA` with one-based positions into that pair’s RNA chain. An *interface RNA motif* is a maximal run of consecutive contacted positions, rendered back as sequence.

#### Motif atlas

Four RBP motif resources were merged into 2456 matrices over 359 gene symbols. Matrices and structure proteins are both keyed by gene symbol, so isoforms of a gene share one set of matrices. The resources are the RBPBench catalogue [71] (catRAPID_omics v2.1 extended [72], 598 matrices, distributed with `rbpbench` 1.0.6), cisBP-RNA human [73] (598 matrices; the two sets share no motif identifier and no identical matrix), ATtRACT [74] restricted to human and mouse (1253) and RBPDB human [75] (7). All matrices are kept as position probability matrices, the form in which the source atlases distribute them, with rows coerced to sum to one. They are not stored as log-odds matrices, so the background enters only at scoring time and can be chosen for the sequences being searched. Only RBPBench annotates assay provenance per motif, and from it we label as CLIP-derived the 294 matrices whose assay is eCLIP_ENCODE (206), PAR-CLIP (46), HITS-CLIP (26), eCLIP (15), iCLIP (14) or CLIP/PAR-iCLIP/dCLIP/FLASH, and treat the remainder (SELEX, RNAcompete, cisBP-RNA, RBPDB, RBNS) as *in vitro*. Proteins of the PDB entries were mapped to gene symbols through the UniProt REST API, and mouse symbols were upper-cased so that they match human orthologue motifs.

#### Scoring and null model

Each RNA motif is scored against every matrix of its gene by log_2_ odds against a uniform background with a 10*^−^*^3^ pseudocount. Because an RNA motif is frequently shorter than the matrix, alignments may overhang either end provided at least four columns overlap. Only overlapping columns contribute, and the score is rescaled to [0, 1] by the minimum and maximum attainable over exactly those columns. Because of that rescaling, a uniform background cancels exactly for a given alignment, so the choice affects only which overhang is selected. The observed and null scores are computed over the same RNA motifs under the same background. The entry score is the maximum over all (matrix, RNA motif) pairs. The relevant question is whether an RNA motif matches *its own* RBP’s atlas motif better than an arbitrary RBP’s, so for each entry we also score the same RNA motifs against 300 matrices drawn from other genes and report *z* = (*s − µ*_null_)*/σ*_null_ with the empirical 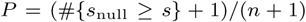. The null is drawn separately for the combined, CLIP-only and *in vitro*-only arms so that each is compared against a null of matching provenance and matrix-width distribution. Related RBPs often share similar motifs, so such motifs can enter the null and make it conservative. We therefore repeated the combined arm with nulls that also exclude same-family paralogs (gene symbol without its trailing number, e.g. ELAVL1–4), motifs similar to one of the RBP’s own (mean column Pearson correlation over the best ungapped overlap of at least four positions *≥* 0.75 or *≥* 0.60), or both (`clip/scripts/null_sensitivity.py`). Experimental interfaces became more strongly on-motif (mean *z* +1.71 to +1.96 against +1.70, 103–112 of 146 entries with *P <* 0.05 against 99). AF3’s RNA motifs stayed within 0.06 of the experimental *z* (*r ≥* 0.89), although this small shortfall reached nominal significance under three of the four stricter nulls (paired *P* = 0.014–0.039).

#### Recognition class

Genes are annotated by recognition mode in clip/scripts/rbp_class.tsv as sequence-specific, structure-driven (dsRBD, helicase and shape-reading proteins) or guide-directed (AGO2, PIWIL1). This is a manually curated assignment from the documented binding mode of each RBP family, drawing on the domain composition recorded in UniProt and InterPro and on the recognition-mode literature [4, 47, 76], fixed before any score was computed, since no motif atlas records *how* a protein achieves specificity. To make it verifiable, every gene was independently re-derived from its UniProt Pfam and InterPro domain annotation (clip/scripts/validate_classes.py). All 57 genes carry a domain record, and the two assignments agree for 54 (95%). The following three exceptions are the expected failure mode of a domain-based rule. WDR33 reads AAUAAA through a WD40 propeller and NUDT21 reads UGUA through a Nudix fold, neither of which the domain vocabulary marks as sequence-reading, while RBM8A carries an RRM but is deposited on mRNA by the spliceosome. In each case the measured specificity favours the manually curated assignment.

#### Domain families

Grouped by RBP InterPro domain, motif-annotated entries containing PUF, cold-shock or RRM domains had the highest experimental motif scores (group mean *z*_exp_ = +1.83 to +1.97; 80 entries from 24 genes), and AF3 gave similar scores (*z*_pred_ = +1.86 to +2.04). Experimental motif scores were lower for helicase-containing and Argonaute proteins (*z*_exp_ = +1.22 and +0.85, respectively), consistent with the often limited intrinsic sequence specificity of helicases and the guide-RNA-dependent target recognition of Argonaute proteins. [77, 78]. Among the 17 post-cutoff RRM and cold-shock entries, the mean predicted motif score remained above +2.0, although mean DockQ*_Nr_* was 0.30 for RRM and 0.14 for cold-shock proteins, and 7 of the 13 entries with a DockQ*_Nr_* fell below 0.23. The corresponding pre-cutoff means were 0.72 and 0.81. Agreement with the RBP motif can therefore persist when the interface geometry is poor. These domain summaries are weighted by entry. The broader recognition-class difference does not persist when genes are weighted equally. A domain of the complex is credited to the RBP when its InterPro accession is among the RBP’s UniProt InterPro entries or when all protein chains of the complex are copies of the RBP (clip/scripts/domain_breakdown.py).

#### Coverage limits

Of the 1,307 entries across both deposition sets, 159 (57 genes) map to a protein with a motif in the atlas. The CLIP-derived arm is the smaller one (38 scorable entries) because ATtRACT, cisBP-RNA and RBPDB are *in vitro* resources and only RBPBench annotates CLIP provenance. No available atlas contains viral RBP motifs, so viral entries are reported as uncovered rather than scored.

#### Gene-level sensitivity

The 159 entries cover 57 genes, and genes with many deposited structures would dominate tests over entries. Every test was therefore repeated on gene means, with each gene counted once. Experimental interfaces remain on-motif (mean *z* = +1.74 over 54 genes, *P* = 1.7 *×* 10*^−^*^31^, one-sample *t*-test; gene-cluster bootstrap 95% CI of the entry mean 1.45–1.92). AF3’s RNA motifs still match the experimental ones (*z*_pred_ = +1.64 against +1.71, paired *P* = 0.22, *r* = 0.77, 51 genes), and pre- and post-cutoff genes still do not differ detectably (*P* = 0.33 for experimental specificity and 0.30 for AF3’s deviation, Mann–Whitney). AF3 still contacts the motif-bearing positions more often than its overall contact coverage predicts (means 88% against 71%, Wilcoxon *P* = 1.4 *×* 10*^−^*^6^, 54 genes). The difference between recognition classes, by contrast, does not hold per gene (*z*_exp_ = +1.79 for 32 sequence-specific genes against +1.69 for 22 structure-driven or guide-directed genes, Mann–Whitney *P* = 0.65).

#### Positional test

The RNA-motif score is sequence-based, so an AF3 RNA motif could match the atlas motif while sitting on another, motif-like stretch of the RNA. The experimental RNA motif that best matches the RBP’s atlas motifs was therefore followed nucleotide by nucleotide into the prediction with the exact residue mapping (Section A.7). The recall is the share of its nucleotides that AF3 contacts. It is compared with the share of the whole RNA chain that AF3 contacts, the chance level for an RNA contacted at random positions (Wilcoxon signed-rank test). AF3’s best-scoring RNA motif is the RNA motif of the predicted interface with the highest score against the RBP’s atlas matrices. It lies on the native site when it sits on the native RNA chain and shares at least one nucleotide with the mapped experimental RNA motif (clip/scripts/positional.py).

### B.4. Feature extraction

#### B.4.1. AF3 confidence features

Confidence metrics were extracted from AF3 model outputs using the custom script af3Metrics_new.py, which parses per-model JSON files generated alongside predicted structures. For each prediction, two complementary JSON files were processed, (i) `*_summary_confidences_*.json`, containing model-level confidence summaries and chain-pair metrics, and (ii) `*_full_data_*.json`, containing atom-level predictions including per-atom pLDDT values and full predicted aligned error (PAE) matrices.

##### Polymer-resolved pLDDT aggregation

Atom-level predicted Local Distance Difference Test (pLDDT) scores were extracted from `the atom_plddts` field in the full data JSON. Each atom is associated with a chain identifier (`atom_chain_ids`), allowing grouping by polymer type. Chain types (protein, RNA, DNA) were determined using metadata from the summary JSON and cross-referenced with database-derived structural annotations. For each polymer class, all associated atomic pLDDT values were collected and averaged. The resulting means were normalised from the native AF3 scale (0–100) to a unit interval by division by 100 and rounded to two decimal places. These values were stored as `af3_protein_pLDDT_avg` and `af3_rna_pLDDT_avg`, providing a global measure of local structural confidence for each polymer type.

##### Global and pairwise PAE metrics

Predicted aligned error (PAE) values were obtained from the pae matrix in the full data JSON, which encodes pairwise alignment uncertainty between all residues or atoms. To derive a global uncertainty measure, the PAE matrix was flattened and the mean of all elements was computed, yielding `af3_global_pae_avg`. This metric reflects overall structural uncertainty, with lower values indicating higher confidence in relative positioning across the structure.

In addition, inter-chain confidence was captured using chain-pair minimum PAE values. These were obtained from the `chain_pair_pae_min` matrix in the summary JSON, which provides the minimum predicted alignment error for each chain pair. For protein–RNA complexes, a representative chain pair was selected either directly (for single protein–RNA chain pairs) or by choosing the pair corresponding to the longest interacting motif when multiple chains were present. Chain identifiers (e.g. A, B, C) were mapped to matrix indices, and the corresponding PAE value was extracted. This scalar value, representing the best-case inter-chain alignment confidence, was stored as `af3_chain_pair_pae_min` (serialised as a JSON-encoded value). Lower values indicate higher confidence in inter-chain spatial relationships.

##### Model-level confidence scores

Additional global confidence metrics were extracted directly from the summary JSON. These include predicted TM-score (pTM) and interface predicted TM-score (ipTM), which quantify model-wide and interface-specific structural confidence, respectively. These values were stored in polymer-resolved form (e.g. `af3_protein_pTM, af3_rna_ipTM`) as JSON-encoded entries. Further model descriptors included the fraction of disordered residues (`af3_fraction_disordered`), a binary clash indicator (`af3_has_clash`) reflecting steric inconsistencies in the predicted structure, and the AF3 ranking score (`af3_ranking_score`), which is used internally by the model to prioritise candidate predictions. These features provide complementary perspectives on model quality and stability.

#### B.4.2. Sequence complexity and thermodynamics

RNA sequence features were computed using `extractGCcomplexity.py`, operating on cleaned RNA sequences derived from database entries (typically the longest RNA chain per complex). Prior to analysis, modified nucleotides were mapped to their canonical parent bases (A, U, G, C), and non-standard characters were removed to ensure consistent downstream calculations.

Basic compositional features included GC content (#*G* +#*C*)*/L*, AU content (#*A* +#*U*)*/L*, and strand asymmetry measures such as GC skew (*G − C*)*/*(*G* + *C*) and AU skew (*A− U*)*/*(*A* + *U*), with safeguards for zero-denominator cases. These metrics capture nucleotide composition biases relevant to structural stability and interaction propensity.

Sequence complexity was quantified using a composite metric combining linguistic complexity and normalised Shannon entropy. Linguistic complexity was calculated as the average fraction of observed unique *k*-mers relative to the maximum possible, for *k* = 1 to 5, with the upper bound constrained by sequence length. Shannon entropy was computed as

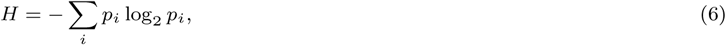

where *p_i_* represents nucleotide frequencies. Two entropy variants were included, mononucleotide entropy (*k* = 1, range 0-2 bits) and dinucleotide entropy (*k* = 2, capturing higher-order sequence dependencies). For the composite complexity score, entropy was normalised to [0, 1] by dividing by its theoretical maximum and combined with linguistic complexity using a weighted scheme (0.6 linguistic, 0.4 entropy).

Thermodynamic and structural propensity features were computed using the ViennaRNA package [79]. For each sequence, minimum free energy (MFE) folding was performed using the partition function framework, yielding the MFE value and corresponding dot-bracket structure. Additional ensemble-based descriptors included centroid energy (representing the most probable structure in the thermodynamic ensemble) and ensemble diversity, which reflects the structural variability of the folding landscape. These features provide sequence-intrinsic proxies for RNA stability, flexibility, and foldability, independent of AF3 structural confidence metrics.

#### B.4.3. Template and evolutionary features

Template usage and multiple sequence alignment (MSA) depth were extracted from AF3 metadata. MSA depth was estimated from sequence alignment data, and binary indicators were derived to classify predictions as MSA-supported or MSA-free. Template usage was inferred from AF3 template references and used to stratify predictions.

#### B.4.4. Biochemical and experimental context

Experimental metadata, including pH, temperature, ion composition, and ligand presence, were extracted from mmCIF files. Ion binding modes were classified based on proximity to RNA functional groups, and ligand interactions were categorised using geometric criteria. Buffer conditions were parsed using rule-based text analysis and classified by ionic strength.

### B.5. Secondary structure analysis

RNA secondary structures were derived from DSSR [80], which computes base pairing and structural annotations directly from 3D coordinates. Dot-bracket representations were converted into graph structures using forgi [81], enabling identification of stems, loops, junctions, and pseudoknots. Structural elements were stored as residue-level annotations and aggregated statistics.

### B.6. RNA secondary structure and conformational feature extraction

RNA secondary structure and nucleotide-level conformational features were derived from three-dimensional structures using DSSR [80], which computes base pairing, backbone geometry, and structural annotations directly from atomic coordinates. For each structure (experimental and predicted), DSSR was executed upstream to generate a corresponding JSON file (e.g. `*_pdb_dssr.json` for experimental structures and `*_af_dssr.json` for predicted models). These JSON outputs served as the primary input for downstream analysis.

#### Nucleotide-level conformational features

The script `dssrConformation.py` processes DSSR JSON outputs by iterating over the nucleotide entries, which encode residue-specific geometric descriptors derived from the 3D structure. For each nucleotide, key conformational parameters were extracted and discretised into biologically interpretable categories. Specifically, the glycosidic bond angle (*χ*) was classified into *anti*, *syn*, or *other* conformations. Ribose sugar puckering was categorised into standard DSSR classes such as C3*^′^*-*endo* and C2*^′^*-*endo*, and the backbone *γ* torsion angle was grouped into *g+*, *g-*, *trans*, or *other*.

Counts of these conformational states were aggregated across each RNA chain using `collections.Counter`, yielding compact frequency distributions that summarise global conformational preferences. These aggregated descriptors were serialised as string-encoded dictionaries and stored in the database as `RNA_GlycosidicBond, RNA_SugarPucker`, and `RNA_GammaAngle`.

#### Secondary structure representation and graph-based decomposition

Secondary structure elements were derived from DSSR-provided dot-bracket notation, obtained from the `dbn.all_chains` field in the JSON output. This representation includes both canonical base pairs and extended bracket types for pseudoknots. Importantly, the secondary structure is not predicted independently in this workflow but is instead directly inferred from the 3D coordinates via DSSR.

Dot-bracket strings were converted into graph representations using the forgi library [81], specifically via `BulgeGraph.from_dotbracket` with pseudoknot retention enabled (`remove_pseudoknots=False`). This transformation produces a bulge graph representation in which RNA secondary structure is decomposed into formal structural elements.

From this graph, the following elements were systematically identified.

*•* **Stems:** contiguous base-paired regions, including residue indices for both strands and base-pair connectivity.
*•* **Hairpin loops:** terminal loops defined by a single closing base pair, including loop length and residue composition.
*•* **Internal loops:** unpaired regions flanked by two stems, capturing asymmetric or symmetric loop configurations.
*•* **Multibranch loops (junctions):** regions connecting three or more stems, characterised by branch degree and connectivity.
*•* **Dangling ends:** single-stranded regions at the 5*^′^* and 3*^′^* termini, merged into contiguous residue intervals.
*•* **Pseudoknots:** non-nested base-pairing interactions identified via pseudoknotted_basepairs().

#### Structural classes

To compare accuracy across RNA architectures, every target RNA was assigned a single structural class from its *experimental* structure (assigning it from the prediction would be circular), using the element counts above and testing the classes in order of structural complexity, so that the first match wins. The classes are **tRNA** (tRNA, by Rfam family, or an unannotated 70-95 nt RNA with a multiway junction and a 3*^′^*-CCA end), **pseudoknot**, **multiway junction** (*≥* 3-way), **internal loop** (including bulges), **hairpin**, **duplex** (base-paired without a loop) and **single-stranded** (no base pairs). Where the forgi decomposition failed (all element lists empty, including dangling ends; 2 post- and 14 pre-cutoff protein–RNA and 5 and 33 single-chain RNA targets), the class was taken directly from the experimental DSSR annotation (stems, hairpins, internal loops and bulges, *n*-way junctions, and pseudoknot brackets in the all-chain dot-bracket). Where both annotations exist, the two class assignments agree for 98-100% of targets. G-quadruplexes were not analysed as a class, because DSSR detects G-tetrads in only three experimental structures, all pre-cutoff. The classification is implemented in `evaluation/rna_structure_classes.py`.

Secondary structure evaluation was performed using RnaBench [82]. We use Matthews Correlation Coefficient (MCC) and F1-Score to assess performance of secondary structure predictions. For MCC, the value range is -1 to 1, where a value of 1 denotes an optimal prediction, a value of 0 indicates a random prediction, and a value of -1 means a completely complementary prediction. F1-Scores range from 0 to 1 (optimal prediction).

### B.7. Statistical reporting

Group differences are tested with the Mann–Whitney *U* test and summarised by Cohen’s *d* as a descriptive effect size. Since TM- and LDDT-type metrics are bounded and often non-normal, significance statements rest on the rank-based tests, not on *d*. Paired analyses report medians with bootstrap 95% confidence intervals. Small strata, notably the post-cutoff single-chain RNA set (*n* = 66) and the per-family RNAcompete values (5–6 proteins per family), are reported with their sample sizes and read as indicative. To control the false-discovery rate over the whole paper, all 71 *P* values were corrected together with the Benjamini–Hochberg procedure. Of the 50 nominally significant results, 45 remain significant at a false-discovery rate of 0.05. The five that do not (*P* = 0.034–0.049, *q* = 0.051–0.069) are marked as nominal where they are reported, and no conclusion rests on them.

### B.8. Use of generative AI

Generative AI tools were used as assistive tools during later stages of this work, primarily to support selected post-hoc analyses of already generated evaluation data, including statistical analysis code and visualisation, and to a limited extent for manuscript drafting and language refinement. The study design, AF3 prediction setup, ablation experiments, structural and 3D evaluation procedures, evaluation tools, data-processing pipeline and generation of the primary evaluation results were developed and carried out without generative AI. The authors take full responsibility for all analyses, results, interpretations and text.

## C. Prediction accuracy is invariant to seed and model choice

To establish whether AF3’s stochastic choices affect protein–RNA modelling, we predicted 49 post-cutoff targets across a fully crossed grid of three random seeds and five ranked models (49*×*3*×*5 = 735 predictions) and partitioned the variance of each quality metric among target, seed and model (Fig. S3A). The seed and model index together explained at most 0.5% of the variance of any metric, whereas the target identity accounted for 57.0-99.7%. Thus, the outcome is determined by which complex is being modelled, not by how it is sampled. Every pairwise contrast between models (model*_k_* vs. model_0_) and between seeds (seed*_j_* vs. s1) fell within the conventional negligible-effect band (Cohen’s *|d| <* 0.2, largest *|d|* = 0.142 for the RNA interface confidence ipTM*_r_*; Fig. S3B). At the level of individual targets the scores were essentially flat across both factors, the mean shifting by only 0.021 (TM*_r_*) across models and 0.018 across seeds (Fig. S3C,D). The only metric to show any seed/model sensitivity at all was AF3’s own confidence, and even there the effect remained negligible. We therefore treat the five models and three seeds as interchangeable and, unless stated otherwise, report the default top-ranked model (model 0).

**Fig. S3:**
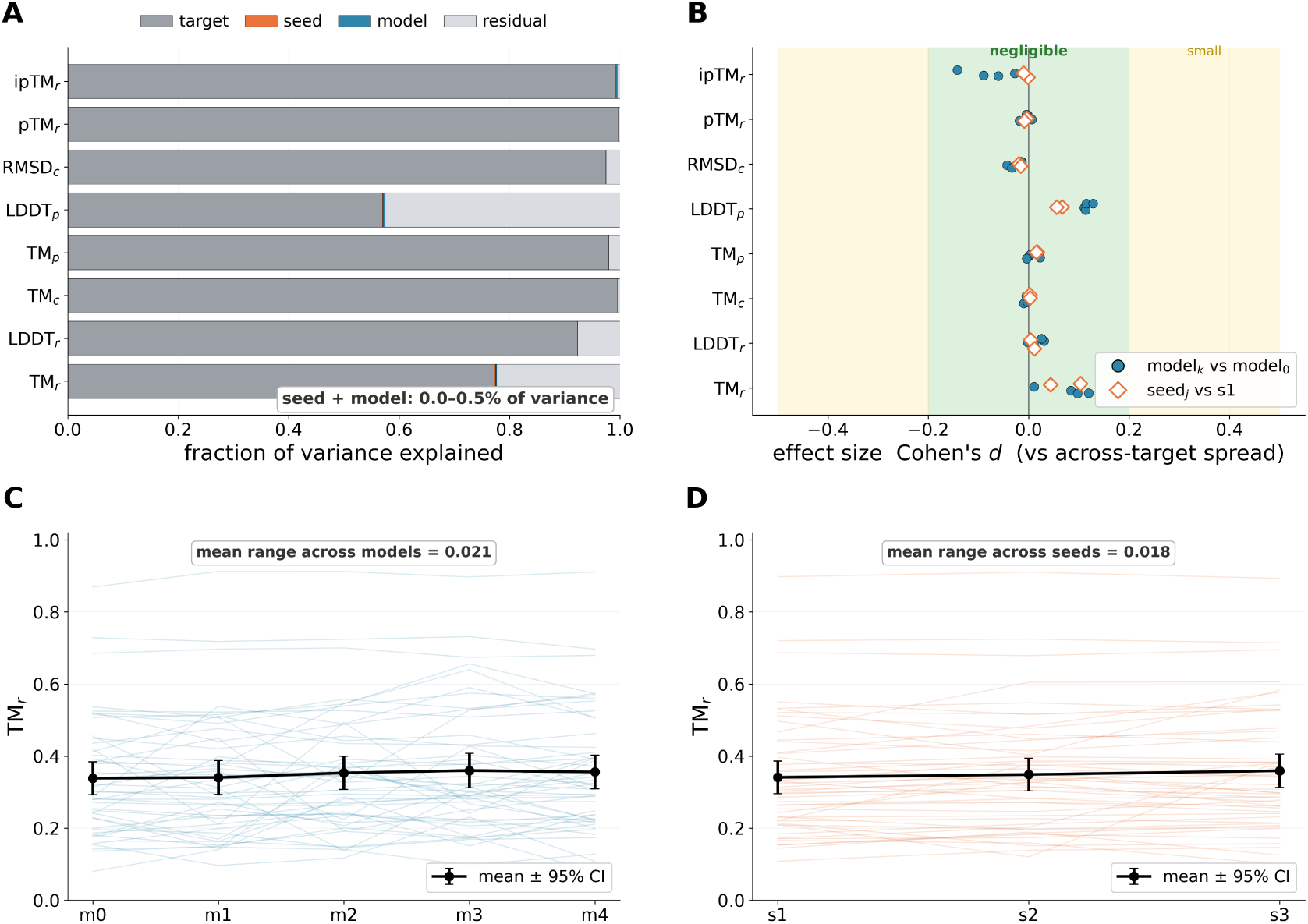
AF3 accuracy on post-cutoff protein–RNA complexes is invariant to the random seed and the model index. 49 post-cutoff targets were predicted with three seeds and five ranked models each (735 predictions). **A**, Share of the variance of each metric explained by target, seed, model and residual. Seed and model together explain at most 0.5%, the target 57–99.7%. **B**, Cohen’s *d* of each model (circles) and seed (diamonds) against model 0 and seed 1. All 48 contrasts are negligible (*|d| ≤* 0.14). **C**, **D**, Per-target TM*_r_* across models (C) and seeds (D), with the mean and 95% CI in black.

## D. Multi-chain RNA complexes

To test whether the generalisation gap also affects interfaces between RNA molecules, we evaluated AF3 on complexes of two or more RNA chains without protein. Twenty were released after the training cutoff (2023-2025; 18 with two RNA chains, 2 with six) and 86 before it (1995-2019; two to eight chains). Predictions were scored as for the protein–RNA set. Every pair of RNA chains in contact in the deposited structure was treated as an RNA–RNA interface (32 post- and 214 pre-cutoff interfaces), including contacts between copies in multi-copy asymmetric units. Contacts (hydrogen bonds and van der Waals contacts between nucleotides) were compared after exact residue mapping (Section A.6). For each interface, we computed the contact match, the Jaccard overlap of the predicted and experimental contacting nucleotides on both sides. After superposing the longer RNA chain on its native counterpart, we also measured the translation and rotation of the partner RNA. The analysis is implemented in `evaluation/rna_rna_interfaces.py`.

Global accuracy drops after the cutoff (Fig. S4A). The median LDDT*_r_* falls from 0.94 to 0.79 (Mann-Whitney *P* = 4 *×* 10*^−^*^7^) and the TM*_c_* from 0.77 to 0.27 (*P* = 0.006), whereas the decrease in TM*_r_* (0.63 to 0.38) is not significant (*P* = 0.08). The decline is clearer at the RNA–RNA interfaces. Before the cutoff, the median contact match is 0.95 and the partner RNA lies within a median 1.1 Å of its native position, rotated by 7*^◦^*. After the cutoff, the median contact match is 0.26 (*P* = 0.013), and the partner RNA is displaced by a median 29.5 Å and rotated by 80*^◦^* (Fig. S4B,C). Displaced partners are mostly also rotated (Fig. S4C). AF3’s chain-pair ipTM tracks the contact match (Spearman *ρ* = 0.58 pre- and 0.80 post-cutoff; Supplementary Fig. S4D). It is, however, low even for many correctly predicted pre-cutoff interfaces (median 0.21 over all pre-cutoff interfaces, 0.44 for those with contact match *≥* 0.5), so it ranks RNA–RNA interfaces rather than certifying them. With 20 post-cutoff complexes these estimates are descriptive.

Placed next to the protein–RNA and single-chain RNA sets, the multi-chain RNA set shows the same pre-to-post decline in LDDT*_r_* and TM*_r_* (Fig. 1). Its pTM*_r_*, in contrast, is higher after the cutoff (*d* = *−*0.99). This reflects RNA length rather than accuracy. The pre-cutoff multi-chain complexes are mostly short duplexes (longest chain a median 17 nt; 83% at most 30 nt), whereas the longest post-cutoff chain is a median 390 nt long, and pTM*_r_* rises with length (Spearman *ρ* = 0.49). Restricted to complexes whose longest chain exceeds 30 nt, pTM*_r_* is lower after the cutoff (median 0.55 before, 0.36 after). A family breakdown, as for the protein–RNA and single-chain RNA sets, is not possible here, because Rfam families are assigned to at least one chain in only 7 of the 20 post-cutoff complexes (mainly group I and group II introns) and 6 of the 86 pre-cutoff ones.

**Fig. S4:**
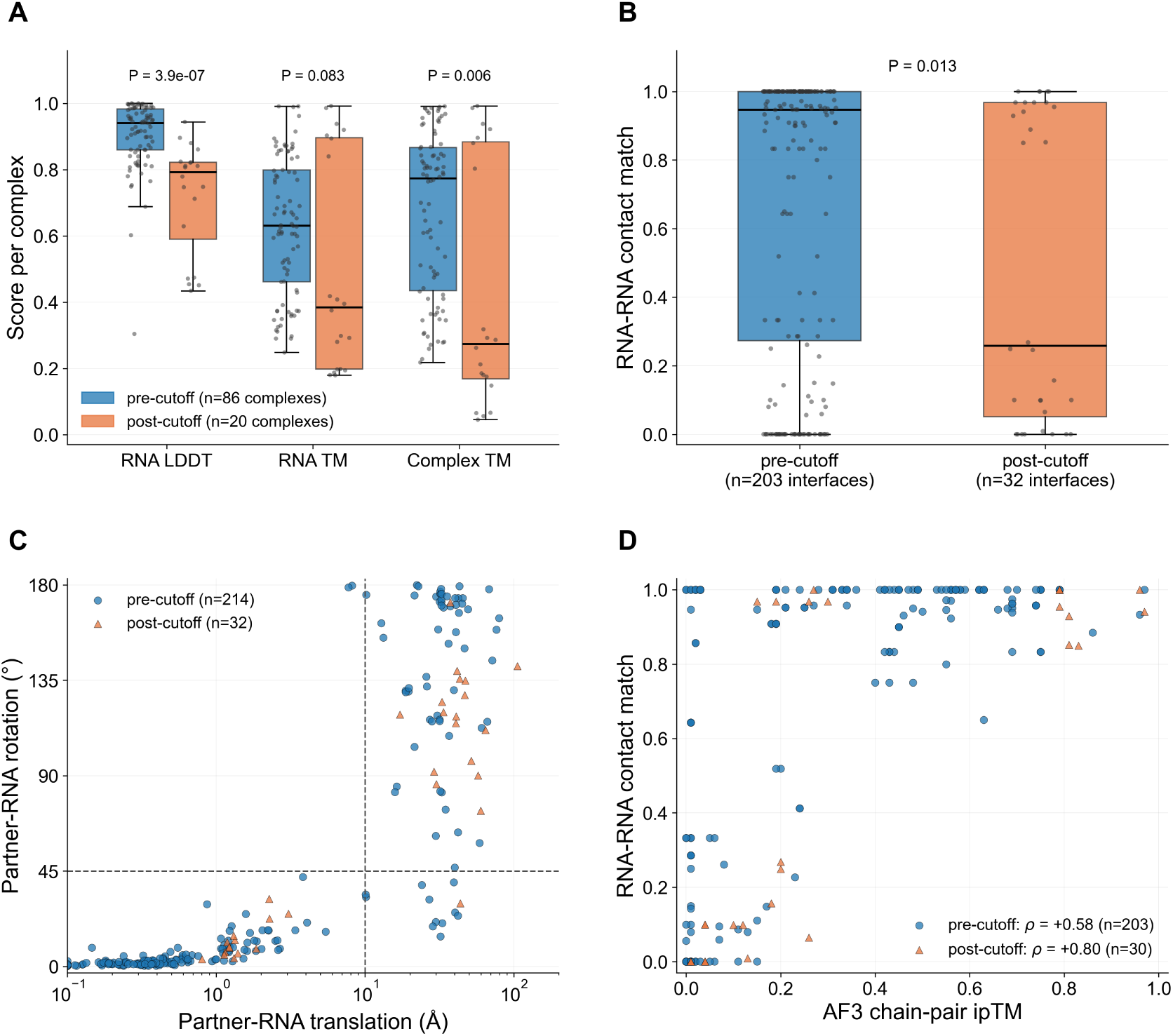
Multi-chain RNA complexes before and after the training cutoff. **A**, LDDT*_r_*, TM*_r_* and TM*_c_*, pre-cutoff (*n* = 86, blue) and post-cutoff (*n* = 20, orange), with Mann–Whitney *P* . **B**, Contact match per RNA–RNA interface after exact residue mapping (pre-cutoff *n* = 203, post-cutoff *n* = 32). **C**, Translation and rotation of the partner RNA after superposing the longer chain (dashed lines at 10 Å and 45*^◦^*). **D**, Chain-pair ipTM against contact match, with Spearman *ρ* per period.

## E. The pre- and post-cutoff sets are comparable, and the generalisation gap survives RNA family- and length-matching

A pre/post-cutoff accuracy gap is only evidence of poor generalisation if the two sets are otherwise comparable. For example, a larger gap could instead reflect the post-cutoff set simply containing intrinsically harder RNAs. We therefore compared the composition of the pre-cutoff (*b2021*) and post-cutoff (*a2021*) RNAs directly (Fig. S5). For the protein–RNA complexes the two sets are closely matched across every RNA sequence and structure descriptor. Length, dinucleotide (Shannon) entropy, folding free energy (MFE), GC content and base-composition skews all differ negligibly (Cohen’s *|d| ≤* 0.25; Fig. S5A,B), so the protein–RNA gap cannot be attributed to a mismatch in RNA difficulty. The protein content of the two sets, on the other hand, is not balanced. Post-cutoff complexes carry substantially more protein (median 553 *→* 1,226 summed residues; *d* = +0.42, *P* = 4.7 *×* 10*^−^*^15^), which places assembly size outside the negligible band that every RNA descriptor occupies. Because AF3 assigns one token per residue or nucleotide, the assemblies themselves are correspondingly larger (median 603 *→* 1,275 tokens), and 67% of post-cutoff complexes exceed the 768-token crop of AF3’s final training stage, against 41% pre-cutoff [19]. Post-cutoff protein–RNA targets are therefore larger as assemblies. This does not account for the gap, which persists within a fixed RNA-length band, within the shared tRNA family and within each experimental method (Fig. S5D,F), but it means the protein–RNA gap should be read as combining novelty with a shift towards assemblies larger than anything AF3 saw intact during training, rather than as isolating novelty alone.

The single-chain RNA sets are also less balanced. Post-cutoff single-chain RNA targets are longer (median 67 *→* 98 nt; *d* = *−*0.67 on log length) and correspondingly more thermodynamically stable (MFE *d* = +0.82), even though their intrinsic sequence complexity is unchanged (Shannon entropy *d* = *−*0.05). Annotated RNA families also turn over almost entirely between the two sets. Only three families recur in each context, with the protein–RNA set tRNA-dominated and the single-chain RNA set shifting from purine/SAM riboswitches to group-I/II introns (Fig. S5C). The accuracy drop persists after controlling for both confounds. Within the one well-populated shared family (tRNA) accuracy still falls post-cutoff (LDDT*_r_* 0.87 *→* 0.74), and it also falls within a fixed length band (TM*_r_* 0.79 *→* 0.54 for single-chain RNAs of 50-100 nt; LDDT*_r_* 0.85 *→* 0.71 for protein–RNA complexes with RNAs below 25 nt; Fig. S5D). The experimental-method mix also shifts strongly across the cutoff. Post-cutoff structures are predominantly cryo-EM, reflecting the growing role of that technique for large ribonucleoprotein assemblies (Fig. S5E), yet the RNA accuracy drop remains large within X-ray and within cryo-EM structures analysed separately (TM*_r_ d* = 1.05 and 0.83; Fig. S5F). The two methods can thus be pooled and the gap is not an artefact of the method shift either. In summary, the generalisation gap is intrinsic to novel RNAs rather than a by-product of family turnover, RNA size, or experimental method.

### Similarity-matched comparison

We compared each post-cutoff target with a pre-cutoff candidate selected by RNA sequence identity (Section 2.2; Fig. S6). A near-identical RNA (*≥* 80% identity) loses little accuracy when its twin carries the same protein (*−*0.020) and 0.100 when the twin carries another protein (Table S1). At 50–80% identity, median LDDT*_r_* falls by 0.20 in protein–RNA complexes and 0.13 in single-chain RNAs. Multi-chain RNA complexes show a similar decline. Matching by structure gives the same gradient as protein-bound RNAs with a redundant or similar pre-cutoff fold lose about 0.04 and 0.07, whereas those with a novel fold lose 0.19. Novel single-chain and multi-chain RNA folds lose 0.18 and 0.32, respectively.

Sequence similarity alone misses some familiar folds since 18 of the 29 structurally redundant post-cutoff protein-bound RNAs have no pre-cutoff RNA with *≥* 80% sequence identity. Conversely, a familiar protein does not rescue a novel RNA. In 12 such pairs, median LDDT*_r_* falls by 0.33 (Table S1). AF3 used the matched pre-cutoff complex as an RBP template in only 15 of the 384 protein–RNA pairs. RNA accuracy thus follows RNA familiarity more closely than protein familiarity.

### Structural similarity

The gap is not carried by sequence-novel RNAs that adopt familiar folds. Measured by structure (TM*_r_* to the closest pre-cutoff RNA; Section 2.2), 68% of the post-cutoff protein-bound RNAs of at least 20 nt have no pre-cutoff counterpart with TM*_r_ ≥* 0.5, and these novel folds carry most of the loss (median LDDT*_r_* 0.62, against 0.90 before the cutoff and 0.81 for structurally redundant post-cutoff RNAs). Removing the structurally redundant targets (TM*_r_ ≥* 0.7) leaves the gap intact (median 0.67 against 0.90).

### Alignment and matching details

For each pair of targets, sequence similarity was the highest identity among all their RNA-chain pairs. RNA sequences were globally aligned with Biopython PairwiseAligner [63] (match score 2, mismatch score *−*3, gap-open score *−*5 and gap-extension score *−*2), while protein sequences were aligned with BLOSUM62 (gap-open score *−*10, gap-extension score *−*0.5). Identity was calculated as identical aligned positions divided by the length of the longer sequence. This length normalisation, also used by Ludaic and Elofsson [27], prevents a short RNA from appearing fully identical to a longer sequence that contains it. To reduce the number of exact alignments, RNA sequences were pre-screened by alignment score normalised by the longer sequence, retaining the top 25 candidate sequences per query, whereas protein sequences were pre-screened by shared 3-mers, retaining the top 15. Exact identities were then calculated across all chain pairs for the retained target pairs. Each post-cutoff target was matched to the highest-identity RNA candidate retained by this prescreen, with ties broken by protein identity. Thus, sequence matching identifies the best candidate among the screened set and is not guaranteed to find the global nearest neighbour. Direct alignment allowed us to retain short protein-bound RNAs (9–12 nt) without CD-HIT-EST sequence-length or word-size filtering. For the structural comparison, the longest RNA chain of every post-cutoff target was aligned with US-align [38] (-mol RNA) against every RNA chain of the 1,061 pre-cutoff targets of all three sets. Similarity was the maximum TM*_r_* over pre-cutoff chains, normalised by the post-cutoff chain, and targets were classified as structurally redundant (TM*_r_ ≥* 0.7), similar fold (0.5–0.7) or novel fold (*<* 0.5). These cut-offs follow the use of TM*_r_* above 0.7 to identify redundant folds and 0.5 to separate familiar from divergent folds in a recent RNA benchmark [27]. Pre-cutoff chains shorter than half the post-cutoff chain were skipped because they cannot reach TM*_r_* = 0.5. RNAs shorter than 20 nt, whose TM*_r_* are dominated by small local deviations [39], were assessed by sequence similarity only (these targets were 149 protein–RNA, 3 single-chain and 7 multi-chain). Paired differences are summarised by the median with a bootstrap 95% confidence interval (5,000 resamples). The analyses are implemented in `evaluation/postpre_matched_similarity.py` and `evaluation/structural_similarity_to_pre.py`.

**Table S1.** RNA accuracy in similarity-matched pre- and post-cutoff pairs. Differences are post-cutoff minus the matched pre-cutoff target (median ΔLDDT*_r_* with bootstrap 95% confidence interval). Sequence matches are the highest-identity candidates retained by the prescreen; structural matches use the best RNA TM*_r_* against pre-cutoff chains. *Same protein* denotes *≥* 80% protein sequence identity. Well-folded percentages 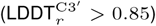 are for the pre-cutoff match and post-cutoff target, respectively. A dash indicates that the corresponding estimate was not reported. The last row is a subset of the protein–RNA novel-fold row.

| Stratum | *n* | Median ΔLDDT<sub>*r*</sub> | 95% CI | Well folded (%), pre → post |
|---|---|---|---|---|
| *Matched by RNA sequence* |  |  |  |  |
| Protein–RNA, ≥ 80%; same protein | 59 | −0.020 | [−0.045, −0.011] | – |
| Protein–RNA, ≥ 80%; other protein | 100 | −0.100 | [−0.152, −0.043] | – |
| Single-chain RNA, ≥ 80% | 17 | +0.004 | [−0.025, +0.022] | – |
| Protein–RNA, 50–80% | 213 | −0.20 | [−0.225, −0.168] | 79 → 31 |
| Single-chain RNA, 50–80% | 46 | −0.13 | [−0.180, −0.101] | 80 → 4 |
| Multi-chain RNA, 50–80% | 18 | −0.17 | [−0.359, −0.107] | 94 → 39 |
| Multi-chain RNA, ≥ 80% | 2 | −0.032 | – | – |
| *Matched by RNA structure* |  |  |  |  |
| Protein–RNA, redundant fold (TM<sub>*r*</sub> ≥ 0.7) | 27 | −0.04 | [−0.120, +0.011] | – |
| Protein–RNA, similar fold (0.5–0.7) | 38 | −0.07 | [−0.110, −0.038] | – |
| Protein–RNA, novel fold (TM<sub>*r*</sub> < 0.5) | 140 | −0.19 | [−0.281, −0.142] | 84 → 29 |
| Single-chain RNA, redundant fold | 13 | −0.005 | [−0.025, +0.007] | 100 → 92 |
| Single-chain RNA, novel fold | 43 | −0.18 | [−0.229, −0.139] | 79 → 2 |
| Multi-chain RNA, novel fold | 11 | −0.32 | [−0.494, −0.137] | – |
| Protein–RNA, novel fold; same protein | 12 | −0.33 | [−0.588, −0.166] | 83 → 17 |

## F. Failures on training-era structures

The pre-cutoff complexes were released before the AF3 training cutoff, so the failures among them occur on structures that were available for training, although whether AF3 was trained on any given entry cannot be verified. Table S2 compares, for each target property, the share of pre-cutoff complexes with a poorly folded RNA and with at least one failed interface. Of the 252 failed RNA-chain interfaces, 117 place the RNA at the correct protein site in a wrong pose, 97 at a distal site, 20 on a mispredicted protein binding region and 18 shifted or rotated at the site. After the cutoff, 313 of 657 RNA-chain interfaces fail, 202 of them at the correct site in a wrong pose.

## G. RNA secondary structure and conformational bias

This appendix details how AF3 reproduces RNA secondary structure and nucleotide-level conformations, summarised in Section 3.2.

INF as an accuracy measure.

INF separates our accuracy classes. Overall INF ranks a well-folded RNA (classes i and ii, 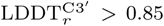) above a poorly folded one (classes iii and iv) with AUC = 0.83 (median 1.00 vs. 0.82), and the separation comes from non-canonical pairs (AUC = 0.79) and stacking (0.65) rather than Watson-Crick pairs (0.63). The INF *>* 0.75 success threshold used in recent RNA benchmarks [27] is, however, too lenient for protein–RNA complexes, because 62% of poorly folded RNAs still pass it, and INF is undefined for 65% of our complexes (249*/*384), whose short protein-bound RNAs form no base pairs. We therefore grade RNA accuracy by LDDT*_r_* and TM*_r_*, and the interface by DockQ*_Nr_*.

### G.0.1. RNA Structural Characteristics

The RNA-level analysis across 384 structures indicates that prediction quality metrics (F1-score, precision, recall, MCC, and Weisfeiler-Lehman graph kernel (WL) [83]) are consistently and significantly correlated with intrinsic RNA structural accuracy, particularly LDDT*_r_* and iLDDT (moderate positive correlations *r ∼* 0.16-0.28, *p ≪* 0.001), while showing weaker or inconsistent relationships with global complex quality (LDDT*_c_*) and only modest negative correlations with RMSD*_r_*, except for WL which exhibits a strong inverse relationship (*r ≈ −*0.45). Notably, all performance metrics increase with RNA length (especially MCC and F1), suggesting that longer RNAs provide more detectable signal for correct prediction.

At the secondary structure level, the model reproduces the overall topology well, with similar counts of structural elements (e.g., stems: 1.70 predicted vs. 1.73 experimental per structure; pseudoknots: 0.24 vs. 0.22), but systematically biases towards more compact conformations, as reflected by longer predicted stems (18.46 vs. 17.18 residues) and shorter loop regions, including hairpin loops (9.26 vs. 11.11), internal loops (13.22 vs. 17.20), and multibranch loops (21.61 vs. 28.35), indicating reduced flexibility and structural diversity. Motif distribution analysis reinforces this trend, showing overprediction of structured elements such as stems (35.9% predicted vs. 31.5% experimental) and hairpin loops (30.5% vs. 27.3%), alongside underprediction of single-stranded regions (62.8% vs. 67.2%), consistent with a bias towards more ordered and compact RNA structures.

**Fig. S5:**
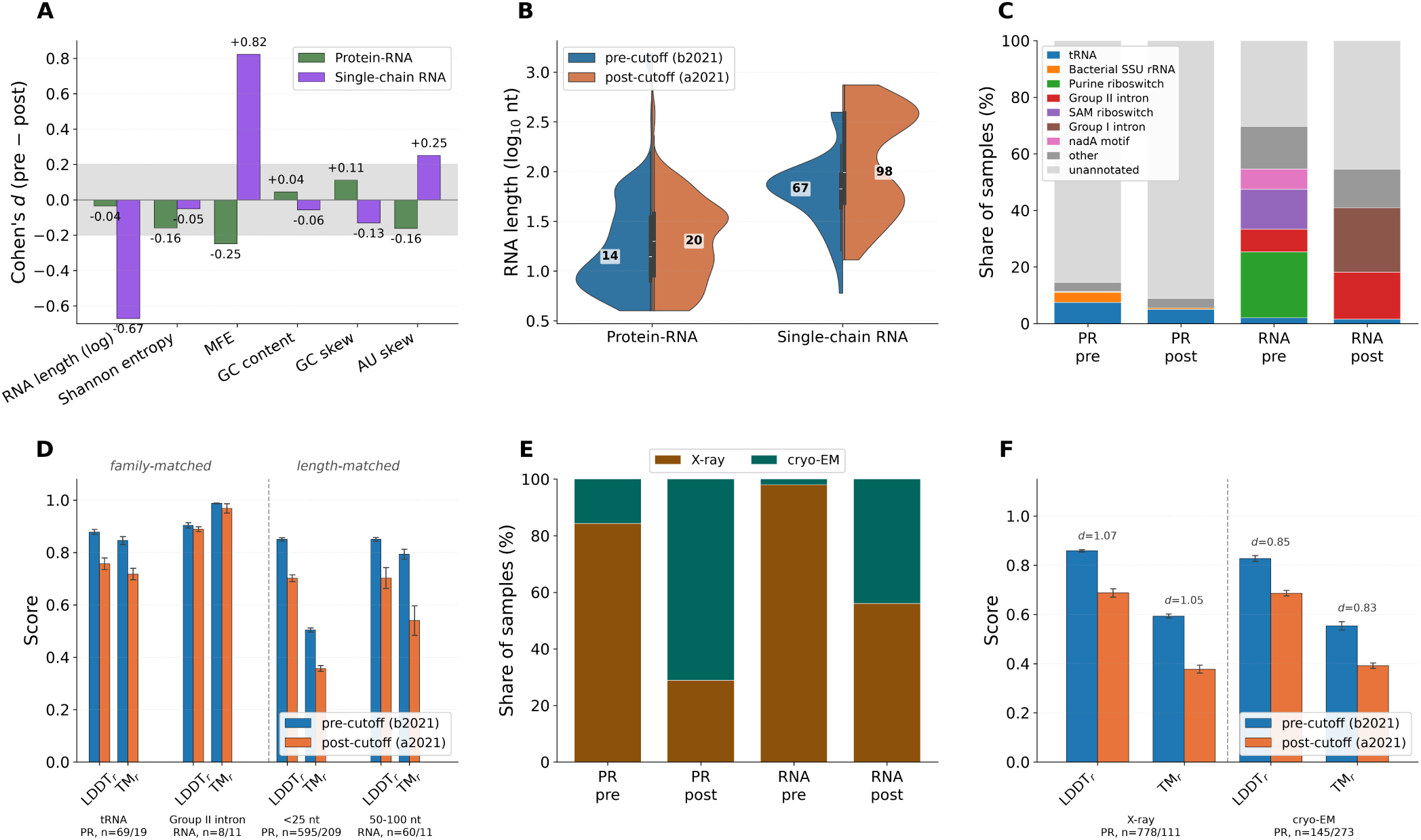
The pre- and post-cutoff sets are comparable, and the gap survives family- and length-matching. Pre-cutoff (b2021, blue) and post-cutoff (a2021, orange) RNAs of protein–RNA complexes and single-chain RNAs. **A**, RNA length. **B**, Cohen’s *d* of RNA complexity descriptors (shaded, *|d| <* 0.2). Protein–RNA descriptors are all negligible, whereas single-chain RNAs differ mainly in length. **C**, RNA family composition. **D**, Mean LDDT*_r_* and TM*_r_* within shared families (tRNA, group-II intron) and within the modal length bin. **E**, Share of X-ray and cryo-EM structures. **F**, RNA accuracy within X-ray and within cryo-EM structures (protein–RNA only).

**Fig. S6:**
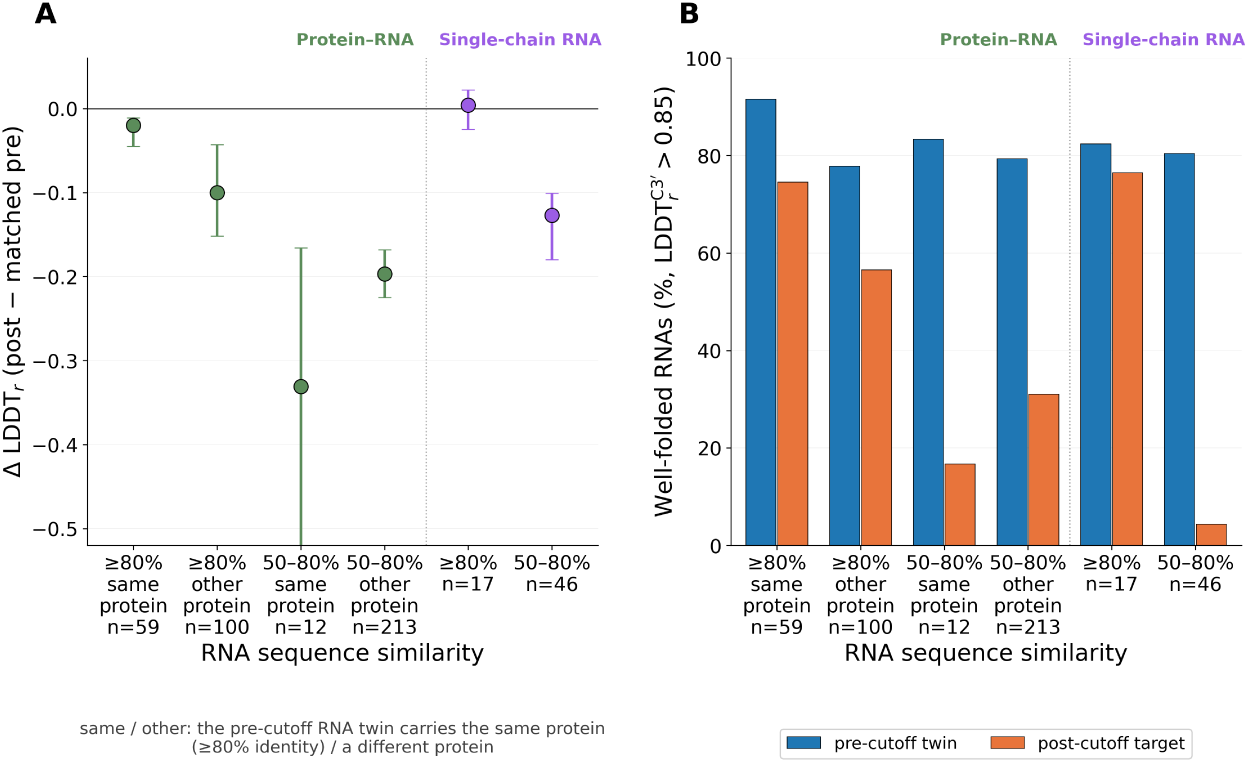
Within sequence-matched pairs, RNA accuracy follows the novelty of the RNA. Each post-cutoff target is paired with the pre-cutoff target whose RNA is most similar in sequence, grouped by that similarity (*≥* 80% or 50–80%) and, for protein–RNA pairs, by whether the twin carries the same or another protein (*n* under each group). **A**, Paired difference in LDDT*_r_* (post-cutoff target minus twin, median with bootstrap 95% CI). **B**, Share of well-folded RNAs among the twins (blue) and the post-cutoff targets (orange).

**Fig. S7:**
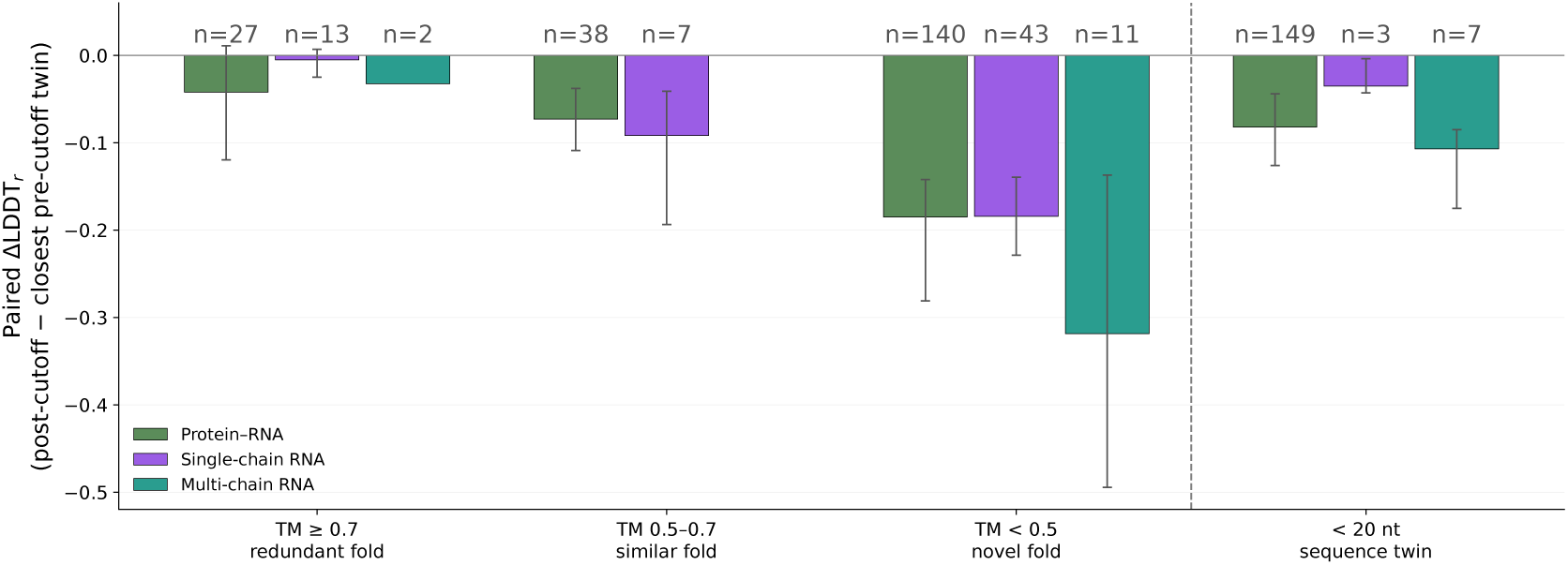
Accuracy against the closest pre-cutoff structural twin. Paired difference in LDDT*_r_* between every post-cutoff target and its closest pre-cutoff twin of the same system class (median, bootstrap 95% CI, *n* per bar), by the structural similarity of its longest RNA chain to the pre-cutoff RNAs (TM*_r_*: *≥* 0.7 redundant, 0.5–0.7 similar, *<* 0.5 novel fold) and, for RNAs shorter than 20 nt, against the closest sequence twin. Bars with *n ≤* 3 are indicative only.

**Table S2.**
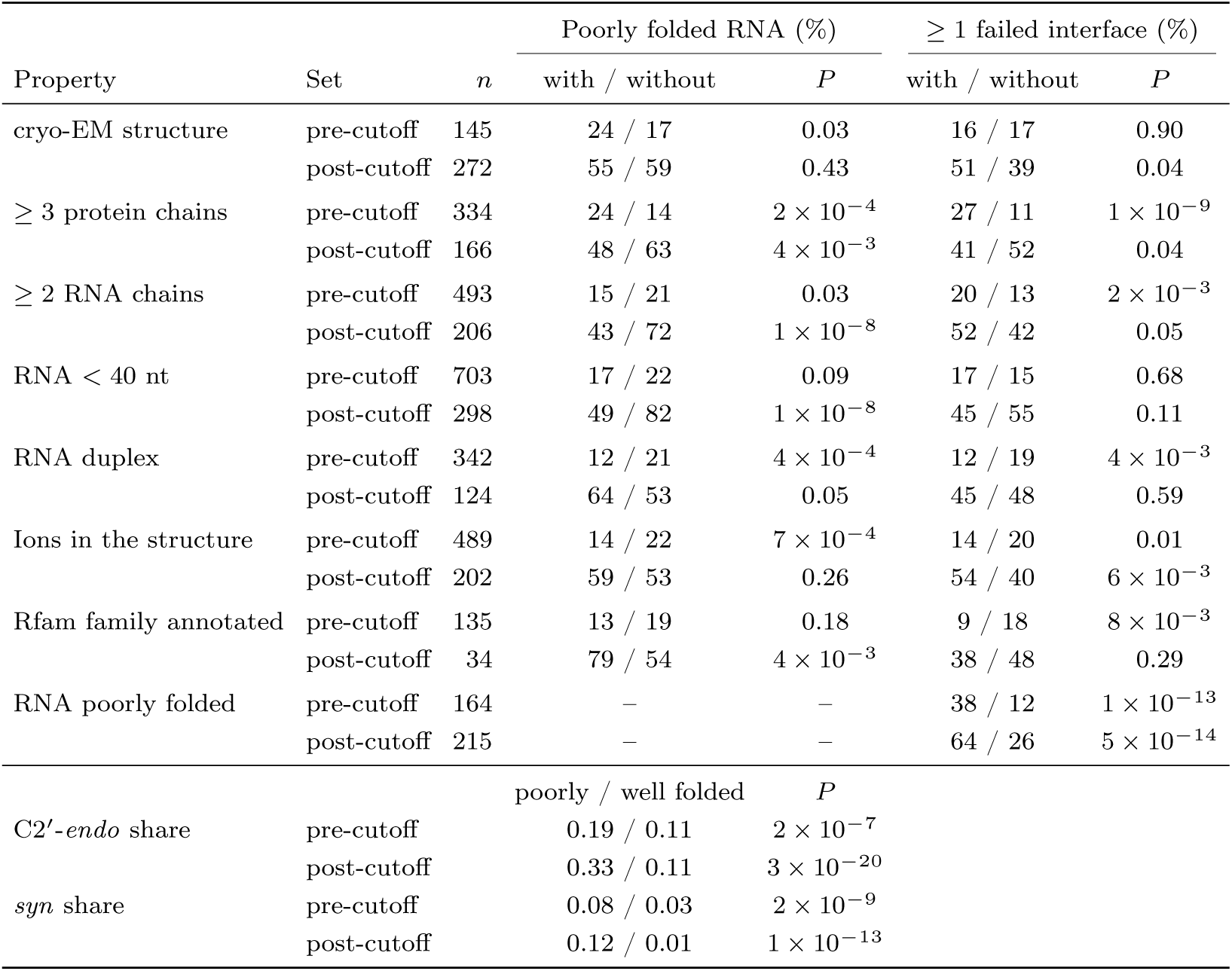
Failures on pre-cutoff protein–RNA complexes, which were released before the AF3 training cutoff and were therefore available for training, next to the post-cutoff complexes. A poorly folded RNA has 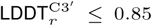 (fold classes iii and iv). A complex has a failed interface when at least one of its RNA chains falls outside the correct-interface placement class. For each property, the share of complexes that fail is given with and without the property (*n*, complexes with the property), with Fisher’s exact test. The lower rows compare the deposited RNA of poorly and well-folded predictions (median, Mann–Whitney test).

**Fig. S8:**
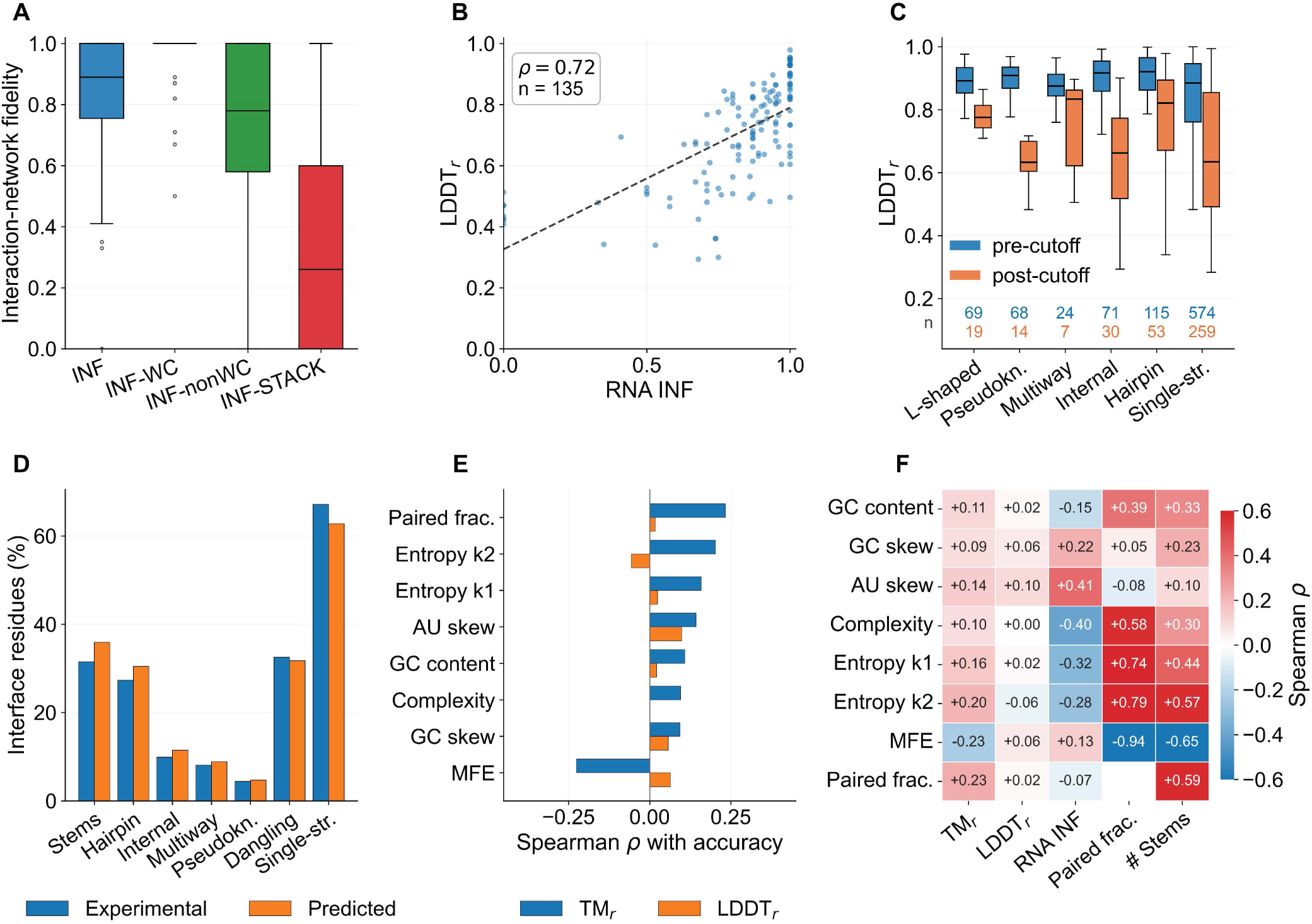
RNA accuracy depends on secondary structure rather than on sequence composition. Post-cutoff protein–RNA complexes (*n* = 384) and, in C, pre-cutoff ones (*n* = 923). **A**, Interaction-network fidelity (INF) for Watson–Crick pairs, non-canonical pairs and stacking. **B**, LDDT*_r_* against INF (Spearman *ρ* = 0.72, *n* = 135). **C**, LDDT*_r_* by the structural class of the target RNA, before (blue) and after (orange) the cutoff (dashed line, well-folded threshold). **D**, Secondary-structure context of the protein-contacting nucleotides, experimental and predicted. **E**, Spearman *ρ* of each sequence and structure descriptor with TM*_r_* and LDDT*_r_*. **F**, Correlation of the descriptors with accuracy and with secondary-structure content.

#### Accuracy by structural class

Assigning each target a single structural class (Section B.6) shows that the generalisation gap is concentrated in specific RNA architectures (Fig. S8C). Before the training cutoff, every class of protein-bound RNA is predicted well (median LDDT*_r_* 0.88-0.92; 79-93% of RNAs well folded). After it, accuracy collapses for the classes that depend most on tertiary and non-canonical contacts, namely pseudoknots (median LDDT*_r_* 0.91 *→* 0.63; well folded 85% *→* 7%; *n* = 68*/*14 pre/post) and internal loops (0.92 *→* 0.66; 93% *→* 13%; *n* = 71*/*30). Hairpins (0.92 *→* 0.82) and multiway junctions (0.88 *→* 0.83) lose least, and even tRNA, the best-represented fold, drops from 88% to 32% well folded (0.89 *→* 0.78). Within each period the classes differ significantly in both 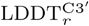 and TM (Kruskal-Wallis, post-cutoff *P* = 5 *×* 10*^−^*^5^ and 3 *×* 10*^−^*^13^). The TM*_r_* differences are, however, inflated by length. Single-stranded protein-bound RNAs are short (median 9-16 nt) and TM*_r_* penalises short chains [39], so 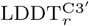 is the fairer comparison between classes. Single-chain RNA folds show the same pattern where classes are populated. Internal loops collapse (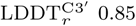 *→* 0.58; *n* = 12*/*13, and pseudoknots retain their global topology (TM*_r_* 0.87 *→* 0.85) yet fall from 73% to 25% well folded *n* = 70*/*40), although here the families also change across the cutoff (purine and SAM riboswitches before, group I and II introns after). The class dependence mirrors the base-pairing breakdown (Fig. S8A). AF3 transfers the canonical helices of hairpins to novel RNAs, but not the non-canonical and stacking interactions on which pseudoknots and internal loops often depend.

The bond-level analysis reveals substantial and highly significant discrepancies between predicted and experimental RNA conformational features, indicating systematic structural biases in AF3. Glycosidic bond conformations are strongly skewed towards the *anti* state in predictions (95.9% vs. 89.1% experimentally), with severe underrepresentation of *syn* conformations, supported by an extremely significant chi-square result (*p ∼* 10*^−^*^102^). Similarly, sugar pucker distributions are biased towards C3’-*endo*, while rarer puckers (e.g., C2’-*exo*) are markedly underpredicted, again with strong statistical significance (*p ∼* 10*^−^*^127^). Gamma torsion states show overrepresentation of the canonical *g*^+^ conformation and underrepresentation of *g^−^* and *trans* states (*p ∼* 10*^−^*^217^). This pattern mirrors its bias toward common sugar-pucker and base conformations. This bias is further reflected in torsion angle means, where large deviations are observed for *α*, *β*, and *χ*, while *γ* and *δ* are comparatively well reproduced. Accuracy metrics confirm this pattern as backbone angles like *δ* (*∼* 90%) and *ɛ* are well captured, whereas *α* and *β* remain poorly predicted (*∼* 30-35%). The presence of RNA MSA information improves classification accuracy for dominant states (e.g., glycosidic *anti*), but does not resolve the poor performance on minority or flexible conformations such as *syn*. AF3 exhibits a strong bias towards energetically favourable, canonical RNA conformations, which limit its ability to accurately capture structural diversity and non-canonical states.

**Fig. S9:**
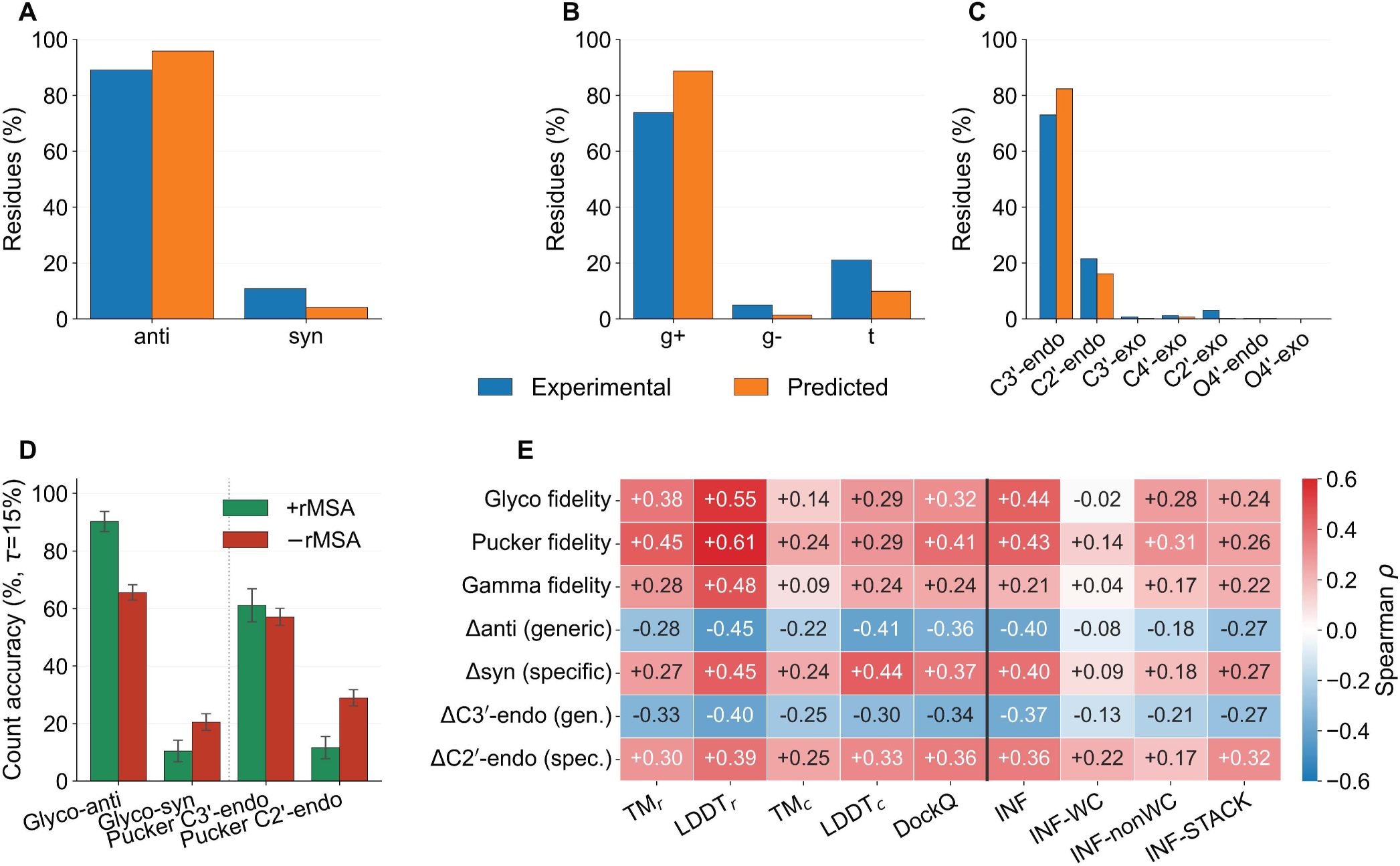
AF3 smooths RNA backbone conformers towards the generic state. Experimental (blue) and predicted (orange) conformers in protein–RNA complexes (*n* = 384). **A**–**C**, Glycosidic bond (A), *γ* rotamer (B) and sugar pucker (C). AF3 over-predicts *anti* and C3*^′^*-*endo* and under-predicts *syn* and C2*^′^*-*endo*. **D**, Share of structures whose conformer counts are within 15% of the experiment, with (green) and without (red) the RNA alignment. **E**, Spearman *ρ* of conformer fidelity and of the over-or under-prediction of each conformer with accuracy (left) and with base-pairing fidelity (right).

#### Structural examples

Figures S10–S12 show two examples of each RNA-fold failure mode, grouped by structural level. At the level of single nucleotides, AF3 replaces rare sugar puckers and glycosidic conformers with the generic C3*^′^*-*endo*, *anti* state (Fig. S10). Between nucleotides, it loses non-canonical pairs and replaces native stacks with others while keeping most Watson–Crick pairs (Fig. S11). AF3 does not simply lose stacking contacts. It swaps them, losing 9 of 21 experimental stacks and adding 13 new ones in 8S9V, and losing 11 of 23 while adding 15 in 8FUK. At the level of secondary and tertiary elements, it drops pseudoknots, rebuilds internal loops as canonical helices, misplaces duplex strands and over-compacts extended RNAs (Fig. S12). Throughout, the experimental structure is orange and the AF3 prediction green.

**Fig. S10:**
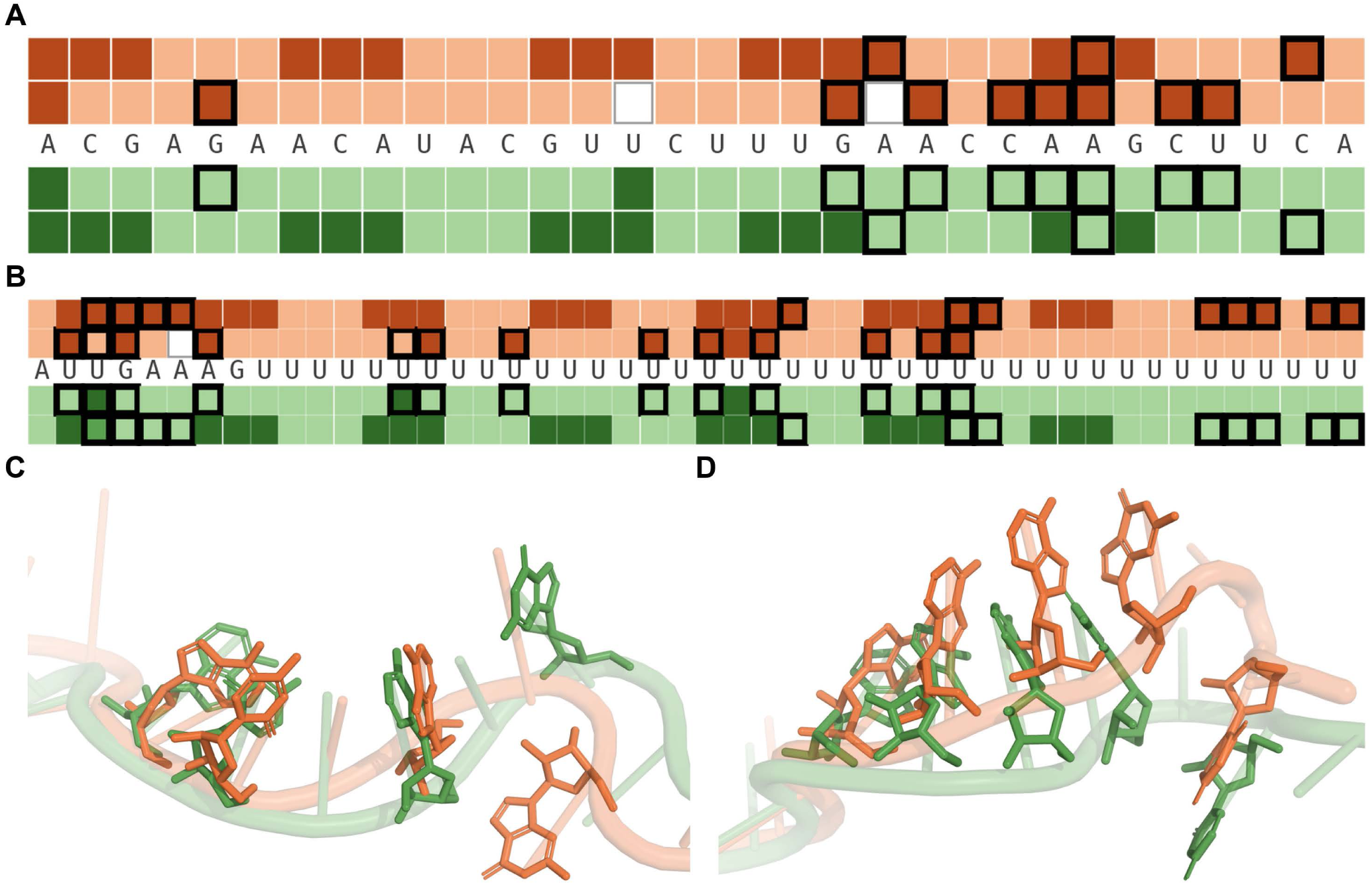
AF3 replaces rare nucleotide conformers with the generic one. Both RNAs come from protein–RNA complexes and are shown without the protein. **A**, **B**, Conformer of every nucleotide in the experimental structure (above the sequence, orange) and in AF3 (below, green). The row next to the sequence is the glycosidic bond and the outer row the sugar pucker. Pale tiles mark the generic *anti* and C3*^′^*-*endo* states, dark tiles *syn* and C2*^′^*-*endo*, and mid-tone tiles other puckers. Black outlines mark nucleotides whose conformer differs between the two structures. **A**, PDB 6XN5, where AF3 keeps 2 of 9 *syn* nucleotides. **B**, PDB 8BMW, where AF3 keeps 3 of 12 *syn* nucleotides and predicts C3*^′^*-*endo* for 12 C2*^′^*-*endo* ones. **C**, **D**, Differing nucleotides as sticks, superposed on the RNA (C, 6XN5; D, 8BMW).

#### Lost and added interactions across the post-cutoff sets

The following examples are representative. Comparing the DSSR annotations of every post-cutoff structure with its AF3 model, nucleotide by nucleotide, shows that AF3 rearranges the nucleotide-level interaction network rather than simply losing native interactions (Fig. S13). In protein–RNA complexes (*n* = 384), it keeps 92% of the Watson–Crick pairs, yet forms new ones equal to 30% of the experimental count. It loses 56% of the non-canonical pairs and adds 35%, and it loses 32% of the stacking contacts while adding 40%. It keeps only 11% of the *syn* nucleotides and 46% of the C2*^′^*-*endo* sugars. In 187 of the 354 complexes with stacking, AF3 adds more stacks than it loses. Single-chain RNAs (*n* = 66) show the same pattern at a lower level, with 6% of Watson–Crick pairs lost and 10% added, 43% of non-canonical pairs lost and 37% added, and 25% of stacks lost and 24% added.

## H. AF3 confidence against accuracy

Figure S14 summarises how AF3’s confidence metrics relate to accuracy on the post-cutoff protein–RNA set (Section 3.1). The ipTM*_r_* distributions by fold class and interface subclass are shown in Fig. 2B.

### Per-nucleotide confidence

AF3’s per-nucleotide confidence registers its RNA-fold errors only partly (Fig. S15). For every element of the experimental RNA (nucleotide conformer, base pair, stacking contact, pseudoknot pair or duplex strand) we compared the pLDDT*_r_* of the AF3 nucleotides it maps to between elements that AF3 misses and elements it reproduces. After the cutoff, missed elements carry lower pLDDT*_r_*, but the separation is weak for the rare conformers and the non-canonical interactions (AUC 0.73 for *syn* bases, 0.68 for C2*^′^*-*endo* sugars, 0.61 for non-g^+^ *γ* torsions, 0.66 for non-WC pairs and 0.58 for base stacks). It is clear only for misplaced duplex strands (AUC 0.93). Most of this separation comes from differences between structures. Compared only within a structure, the AUC falls to 0.53–0.61 for the conformers and to 0.61 for duplex strands (15 structures), so pLDDT*_r_* flags a poorly modelled RNA rather than the wrong nucleotide in it. Watson–Crick pairs are the exception, as the few that AF3 misses also sit in less confident regions within a structure (AUC 0.80). Pre-cutoff structures show the same pattern at higher accuracy. A rare state that AF3 does model is reliable mainly at high confidence. At pLDDT*_r_ ≥* 90 the experiment confirms 88% of AF3’s *syn* bases and 92% of its C2*^′^*-*endo* sugars after the cutoff, against 20% and 59% below 70 (`publication_figures/per_nucleotide_confidence.py`).

**Fig. S11:**
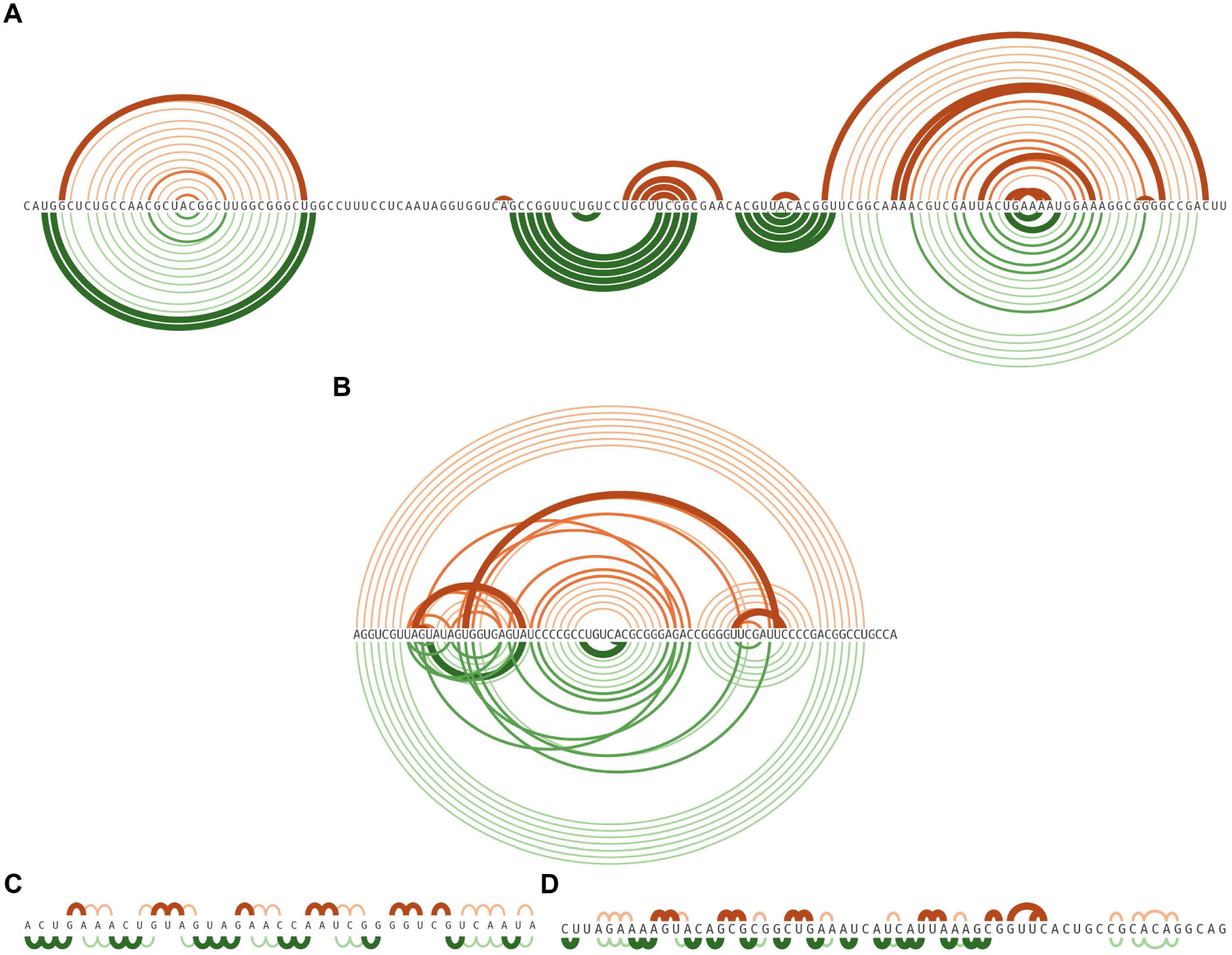
AF3 loses non-canonical pairs and rearranges base stacking. All four RNAs come from protein–RNA complexes. **A**, **B**, Base pairs of the experimental structure (above, orange) and of AF3 (below, green) along the sequence. Pale arcs are Watson–Crick pairs, mid-tone arcs non-canonical pairs, and dark thick arcs pairs present in only one of the two structures. **A**, PDB 8UBE. **B**, PDB 8OMR. **C**, **D**, Stacking contacts drawn in the same way, with pale arcs for stacks in both structures and dark thick arcs for stacks in only one. **C**, PDB 8S9V, where AF3 loses 9 of 21 experimental stacks and adds 13 of its own. **D**, PDB 8FUK, where it loses 11 of 23 and adds 15.

## I. RNA alignments and templates: additional analyses

This appendix details the MSA and template analyses summarised in Section 3.2.

The MSA*_r_* adds little once a protein is present, at least at the RNA lengths this dataset can test. Only *∼*20*/*75 of the 1v1 complexes carry an MSA*_r_* at all, and removing it has essentially no complex-level effect (Fig. 2D,F). The RNA chain is likewise far less template-sensitive than the assembly around it, with LDDT*_r_* improving under RBP templates while TM*_r_* barely changes. This generalises to the broader set of 52 protein–RNA complexes carrying an MSA*_r_*, which include multi-protein and multi-RNA assemblies (of 310 such multi-chain complexes, 82 RNA chains have alignment depth *>* 1; median depth *∼*240, with 27 deep (*>* 1000) and 33 shallow (*≤* 10)). Across the 46 paired complexes for which both conditions were run, removing MSA*_r_* changes complex accuracy negligibly (ΔTM*_c_* = +0.015, median 0; Wilcoxon *P* = 0.345) as most of these are shorter than 200 nt. Only five 1v1 MSA*_r_*-carrying complexes in this set exceed 200 nt.

**Fig. S12:**
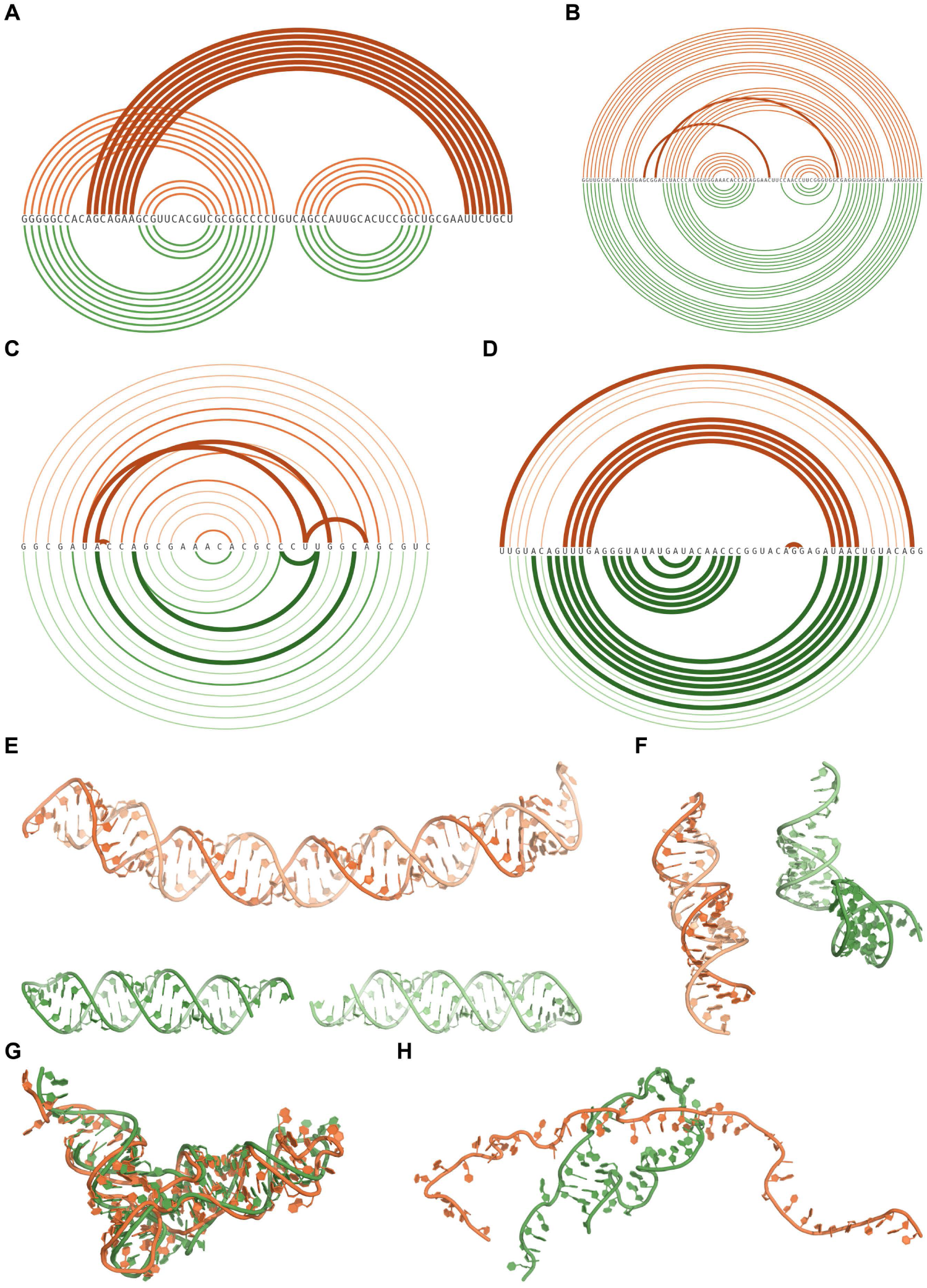
AF3 drops pseudoknots, closes internal loops, misplaces duplex strands and over-compacts RNA. All eight RNAs come from protein–RNA complexes and are shown without the protein. **A**–**D**, Base-pair arcs as in Fig. S11. **A**, **B**, Pseudoknots, with pseudoknotted pairs dark and thick. **A**, PDB 7QR3, where AF3 keeps every nested pair but none of the seven pseudoknotted ones. **B**, PDB 8T2B, where AF3 misses both single-pair pseudoknot contacts. **C**, **D**, Internal loops. **C**, PDB 8DK7, where AF3 replaces four of the eleven pairs inside the loops, mostly non-canonical, with three pairs of its own. **D**, PDB 8OPT, where AF3 shifts the flanking helix by one register and folds the large internal loop into a hairpin. **E**, **F**, Experimental (orange) and predicted (green) RNA strands, one shade per strand. **E**, PDB 7W0D, where AF3 splits the continuous experimental duplex into two. **F**, PDB 8SSW, where AF3 does not reproduce the experimental strand pairing. **G**, **H**, Experimental and predicted RNA superposed. **G**, PDB 8FZR. **H**, PDB 8D97, where AF3 folds an extended RNA into a compact one.

**Fig. S13:**
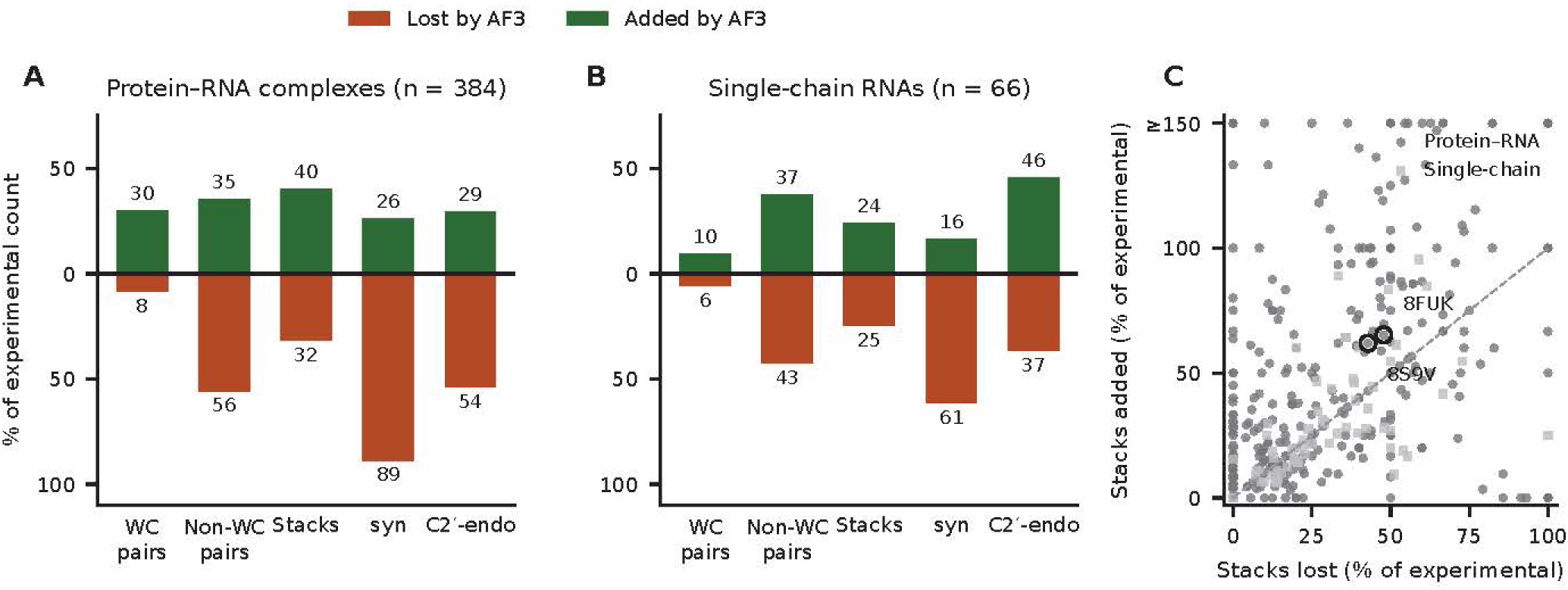
AF3 rearranges the nucleotide-level network of post-cutoff RNAs. Every experimental structure is compared with its AF3 model, nucleotide by nucleotide, using DSSR annotations. **A**, **B**, Interactions and conformers lost by AF3 (present only in the experiment, down) and added by AF3 (present only in the model, up), as percentages of the experimental count. **A**, Protein–RNA complexes (*n* = 384). **B**, Single-chain RNAs from the RNA-only set (*n* = 66). **C**, Stacking contacts lost against added per structure, as percentages of the experimental stacks, with the examples of Fig. S11 circled. Points above the diagonal gain more stacks than they lose.

**Fig. S14:**
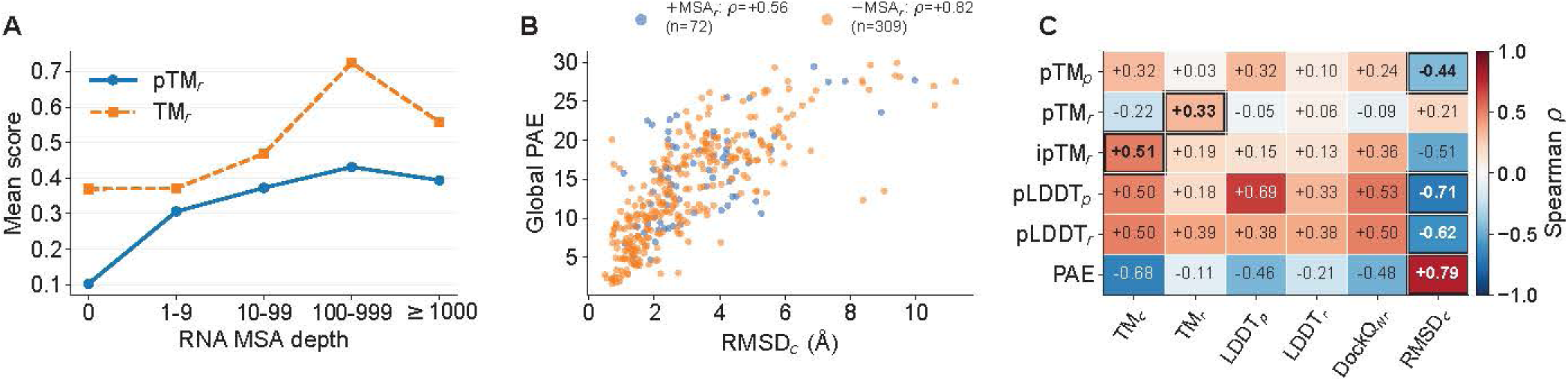
AF3 confidence against accuracy on the post-cutoff protein–RNA set. **A**, Mean pTM*_r_* and TM*_r_* by RNA alignment depth (*n* per bin). **B**, Global PAE against complex RMSD, split by RNA-alignment availability. **C**, Spearman correlation of every AF3 confidence metric (protein and RNA pTM, RNA ipTM, protein and RNA pLDDT, global PAE) with each accuracy axis. The boxed cell marks each metric’s strongest correlate. ipTM*_r_* by fold class and interface subclass is in Fig. 2B.

Interestingly, AF3 rarely templates onto the very structure it is predicting. On the pre-cutoff protein–RNA dataset (*b2021*), where each target’s own PDB entry predates the training cutoff and is therefore in principle available as a template, only 12.6% (112*/*888) of complexes list their own identifier among the selected templates, and always as a protein–RNA complex template rather than a protein-only one. This self-templating rate falls with family size, from 28% for singletons (targets that share no template with any other complex, so their own entry is often the only close template) to 6% for large families (*≥* 21 members), because AF3 collapses each family onto one representative structure that covers most of its members. To illustrate, for the protein-only template the Pumilio PUF domain (3GVO, 91%) and for the complex template the Pan2–Pan3 deadenylase (6R5K, 84%), which post-cutoff members reuse just as readily [19, 21]. Template usage is correspondingly uneven, with a few templates serving many targets (Gini *≈* 0.5 in both datasets). This preference for family representatives over the identical deposited structure is independent of assembly complexity (13.0% self-templating for single-protein–single-RNA versus 12.5% for multi-chain assemblies; *|r| <* 0.05 with either chain count).

Fig. S16 reports the ablation in full, namely which inputs each complex actually carries, the span between all four present and all four removed, the effect of each input on TM*_c_*, and the mean of every one of the 16 input combinations on the complexes that carry all four.

Fig. S17 shows one complex for each input, together with its score under all 16 combinations. Without the RNA alignment, the 373-nt RNA of 7UO5 loses its fold and its interface (TM*_r_* 0.79 *→* 0.28). Removing the RNA alignment alone lowers TM*_r_* by 0.25 or more in only 3 of the 49 complexes that carry one. The protein alignment matters far more often. Without it, the PIWI complex 7YFQ loses its protein fold and its interface (TM*_c_* 0.95 *→* 0.30, DockQ*_Nr_* 0.57 *→* 0.03), and 72 of the 223 complexes that carry it lose at least 0.25 in TM*_c_* from this input alone. Protein templates rarely matter on their own (in 1 of 183 complexes) but stand in for a missing MSA*_p_* (42 complexes). Without the protein alignment, the respiratory syncytial virus polymerase 8SNX keeps its fold with protein templates and loses it without them (TM*_c_* 0.91 *→* 0.20). The RNA-bound template mattered on its own for only one of its 141 carriers, the nucleoprotein–RNA complex 8PDS, whose RNA AF3 places correctly with the template and away from the binding site without it (DockQ*_Nr_* 0.90 *→* 0.03).

**Fig. S15:**
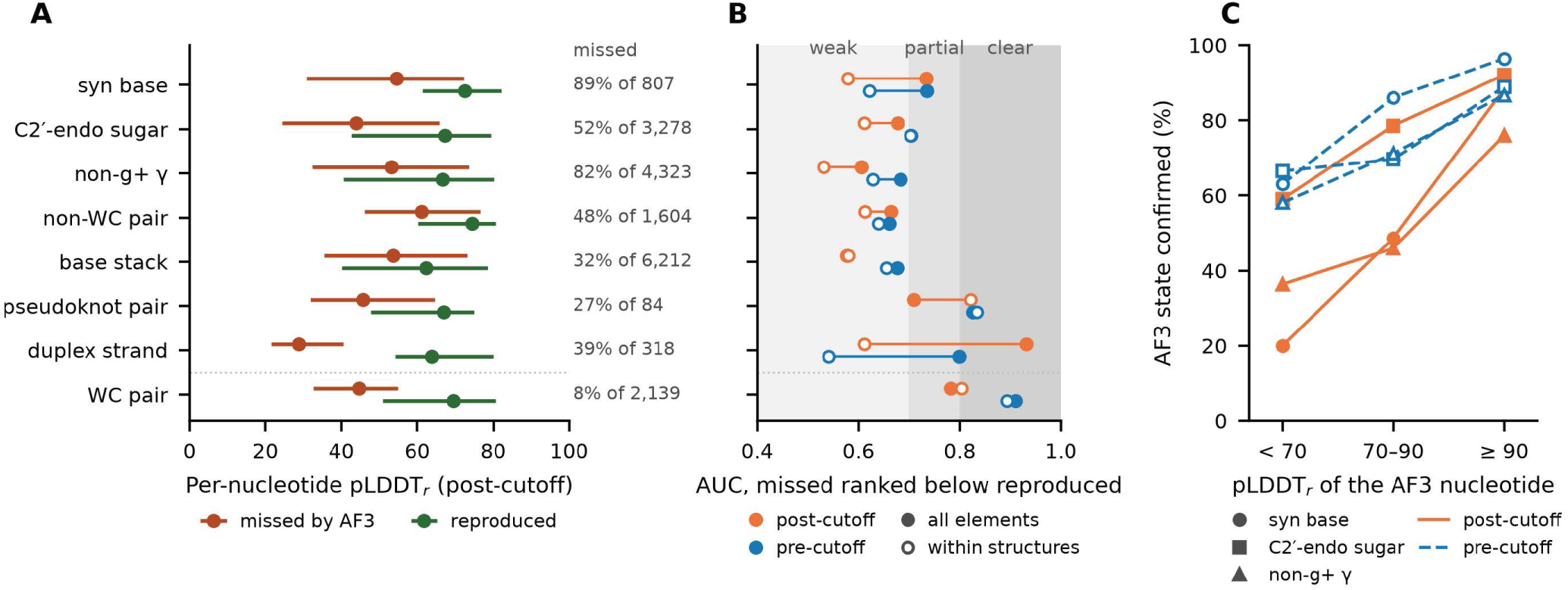
AF3’s per-nucleotide confidence flags poorly modelled RNAs more than the wrong nucleotides. Elements of the experimental RNA in protein–RNA complexes, compared with the AF3 model. Conformers are classed by the glycosidic torsion (*syn* when *|χ| <* 90*^◦^*), the sugar pucker (*δ ≥* 115*^◦^* for C2*^′^*-*endo*) and the *γ* rotamer. **A**, pLDDT*_r_* of the AF3 nucleotides of elements that AF3 misses (red) and reproduces (green), median and interquartile range, for post-cutoff complexes (*n* = 384, share missed on the right). Watson–Crick pairs are shown for reference. **B**, AUC of pLDDT*_r_* for ranking a missed element below a reproduced one, over all elements (filled) and within structures only (open), post-cutoff (orange) and pre-cutoff (*n* = 923, blue). Shading marks weak (*<* 0.7), partial and clear (*≥* 0.8) separation, as in the graphical abstract. **C**, Share of AF3’s *syn* bases, C2*^′^*-*endo* sugars and non-g^+^ *γ* torsions that the experimental structure confirms, by pLDDT*_r_* of the AF3 nucleotide.

**Fig. S16:**
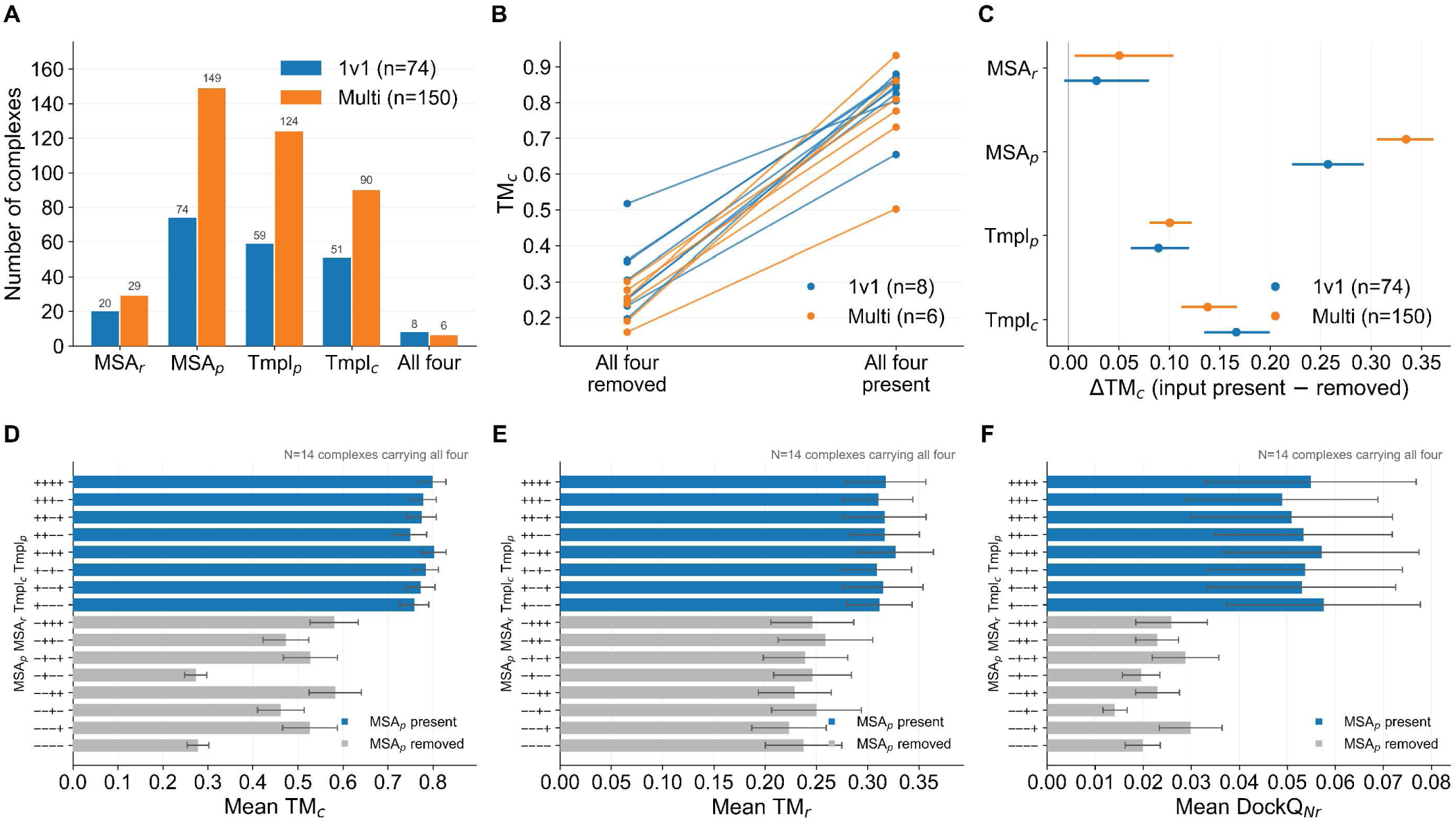
Controlled input ablation in full. 224 protein–RNA complexes predicted under all 16 combinations of MSA*_r_*, MSA*_p_*, Tmpl*_p_* and Tmpl*_c_*. **A**, Complexes that carry each input, for 1v1 and multi-chain complexes. Effects are computed on the carriers only. **B**, TM*_c_* with all four inputs removed against all four present, for the 14 complexes that carry all four. **C**, Effect of each input on TM*_c_* (paired within target, 95% bootstrap CI). **D**–**F**, TM*_c_*, TM*_r_* and DockQ*_Nr_* (mean *±* s.e.m.) for each of the 16 combinations over these 14 complexes, grouped by MSA*_p_*. These complexes carry short RNAs, so the long-RNA regime of Fig. 2D is not represented.

For single-chain RNAs, the RNA alignment can decide the whole fold. Fig. S18 shows one such RNA. With the alignment, AF3 reproduces it almost exactly (TM*_r_* 0.99). Without it, the same sequence folds into an unrelated structure (TM*_r_* 0.22). In protein–RNA complexes the effect is confined to the few complexes whose RNA carries a deep alignment. One such complex from the controlled ablation is shown with every other input present. The 94-nt RNA of 7X34 loses part of its fold without the alignment, and the 373-nt RNA of 7UO5, which loses its fold and its interface, is shown in Fig. S17.

## J. RNA-family and alignment-depth stratification

To test whether RNA accuracy tracks intrinsic foldability or proximity to well-represented, training-familiar folds, we stratified predictions by RNA family. Prediction accuracy in protein–RNA complexes varied systematically with RNA family and MSA*_r_* depth, and the two are strongly confounded because each family occupies essentially a single MSA*_r_*-size bin (Fig. S19). Canonical, evolutionarily conserved folds were predicted well above the dataset mean (e.g. tRNA ΔTM*_r_* = +0.33, SSU rRNA ΔTM*_r_* = +0.55), whereas the unannotated “None” category (*n ≈* 350; synthetic or family-less RNAs) and small, flexible RNAs (e.g. RT-10) sat at or below it (None vs. tRNA *P <* 0.001). Deep alignments (*>*1000 sequences) improved global fold metrics (ΔTM*_r_* = +0.15) whereas shallow or absent MSA*_r_*, common for synthetic and novel RNAs, degraded them. The dependence was much stronger for single-chain RNA predictions. Without a protein partner the single-chain RNA “None” group fell an order of magnitude further below the mean than its protein–RNA counterpart (ΔTM*_r_* = *−*0.26 vs. *−*0.03; Fig. S19a,b), so that lacking an interface the prediction relies almost entirely on intrinsic RNA regularity and evolutionary signal.

## K. Ions supplied as input

This appendix collects the ion case studies summarised in Section 3.2.

### K.1. Ion-addition case studies

For single-chain RNA structures, supplying the chelated ions as input improves accuracy. In 41 cases, LDDT*_r_* rose from 0.699 to 0.749, TM*_r_* from 0.626 to 0.655 and RMSD*_r_* fell by 2.24 Å (all *P ≤* 0.002, two-sided Wilcoxon; Fig. S20a). The gain is largest for weakly-aligned, structurally challenging RNAs (LDDT*_r_* gain versus predicted MFE *ρ* = +0.73; versus MSA depth *ρ* = *−*0.52) and diminishes as more ions are supplied (*ρ* = *−*0.54 with deposited ion count; Fig. S20b,c), so the ions largely substitute for missing evolutionary signal rather than adding independent information. Even when withheld from the prediction, ion-rich RNAs are modelled better (deposited ion count versus RMSD*_r_ ρ* = *−*0.41, *n* = 63; Fig. S20d), so ion load also marks intrinsically well-ordered, predictable RNA architecture. Restricting the count to ions that actually contact the RNA gives the same picture (ribose chelation *ρ* = *−*0.44, phosphate chelation *ρ* = *−*0.36, multidentate *ρ* = *−*0.37). The gain appears even though AF3 often places the supplied ions incorrectly [84], so the screening they provide collectively seems to matter more for the RNA fold than the coordination geometry of any individual ion. Two single-chain examples are shown in Fig. S21A,B.

In protein–RNA complexes, the benefit disappears. Supplying ions to 13 paired predictions left the interface and the RNA fold unchanged (DockQ*_Nr_* 0.457 *→* 0.434, *P* = 0.193; LDDT*_r_* 0.752 *→* 0.740, *P* = 0.635; RMSD*_r_* 4.82 *→* 4.14 Å, *P* = 0.635; Fig. S20f), because the protein partner and its template already supply the electrostatic and geometric scaffold that ions provide [76]. Only the RMSD*_c_* tended to get worse, rising from 13.45 to 16.77 Å (Δ = +3.32 Å, *P* = 0.068), so the added ions may perturb the placement of the assembly without improving either chain. Without an RBP template the interface iLDDT correlates negatively with ion count (Zn^2+^ *ρ* = *−*0.29, *P* = 0.001, *n* = 118; Mn^2+^ *ρ* = *−*0.24, *P* = 0.004, *n* = 9), a dependence that vanishes once a template is present (Fig. S20e). A reason might be that AF3 does not place the added ions correctly. In the paired complex examples the supplied ions are located 11–19 Å from their deposited sites and leave the misplaced RNA where it was (Fig. S21C,D). Ion distributions, and the counterion release that accompanies binding, are also highly dynamic and hard for a single-structure model to capture [85]. Explicit ion input therefore does not help the predicted complex, and can displace it.

Multi-chain RNA complexes without protein behave like the protein–RNA complexes. For 13 post-cutoff complexes predicted with their deposited ions and without them, neither the RNA fold nor the complex changed (LDDT*_r_* 0.683 *→* 0.691, *P* = 0.27; TM*_r_* 0.594 *→* 0.598, *P* = 0.64; RMSD*_r_* 19.6 *→* 17.3 Å, *P* = 0.27; TM*_cr_* 0.585 *→* 0.589, *P* = 0.59; two-sided Wilcoxon signed-rank test). Well-predicted complexes stayed accurate and poorly predicted large ones stayed poor (TM*_r_ ≈* 0.2).

## L. Ligands and solution conditions

This appendix collects the ligand and solution-condition analyses summarised in Section 3.5.

**Fig. S17:**
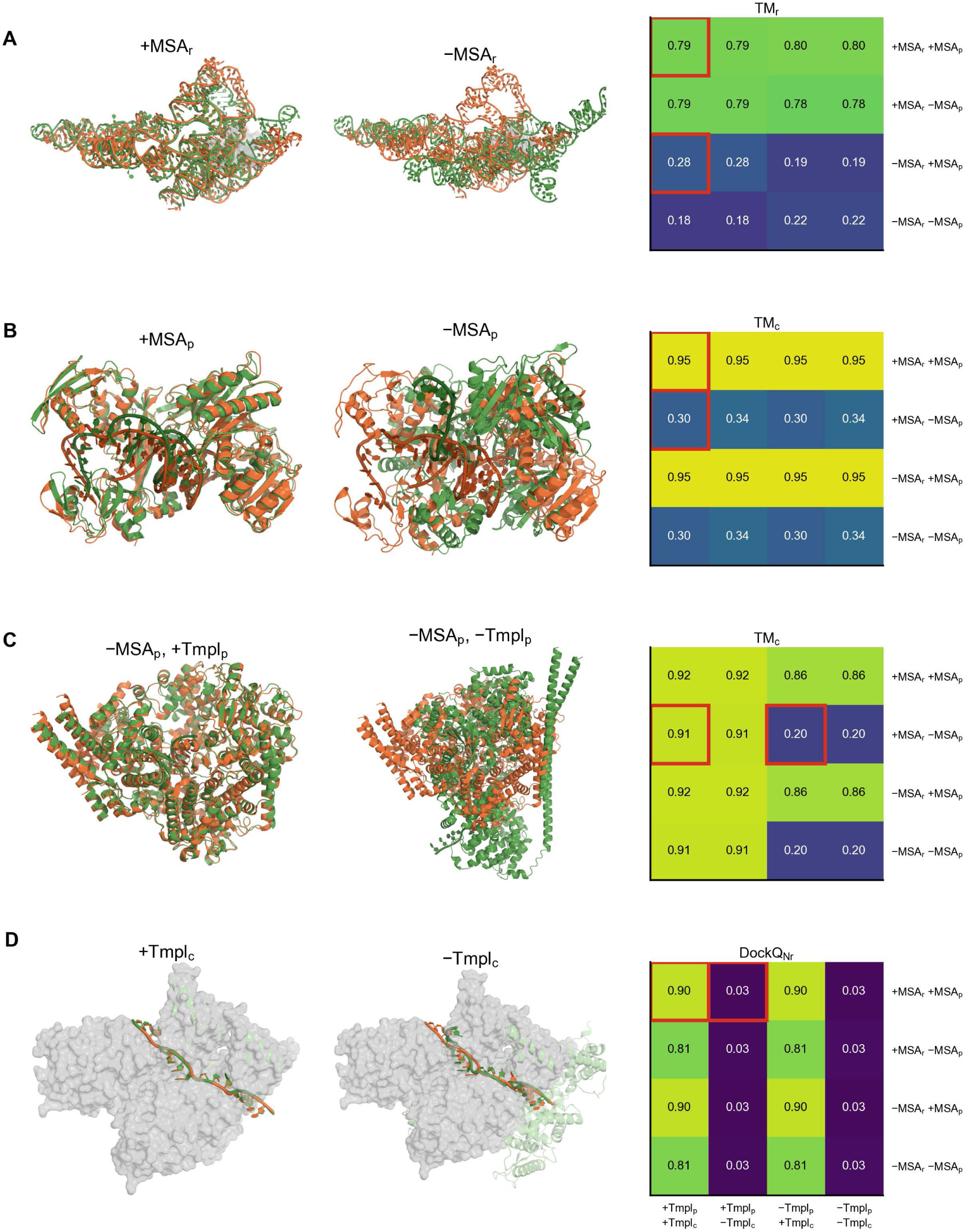
One example per input of the controlled ablation. Experimental structure (orange) and AF3 prediction (green, RNA in the darker shade in **B** and **C**) with and without the named input. Every other input is present unless labelled. Right, the score of the example under all 16 input combinations (rows, alignments; columns, templates), with the two conditions shown boxed. **A**, PDB 7UO5 (373 nt), superposed on the RNA, protein as grey surface (TM*_r_*). **B**, PDB 7YFQ, superposed on the complex (TM*_c_*). **C**, PDB 8SNX without the protein alignment, with and without protein templates (TM*_c_*). **D**, PDB 8PDS, superposed on the RNA phosphates, deposited protein as grey surface and AF3 protein as light-green cartoon (DockQ*_Nr_*).

**Fig. S18:**
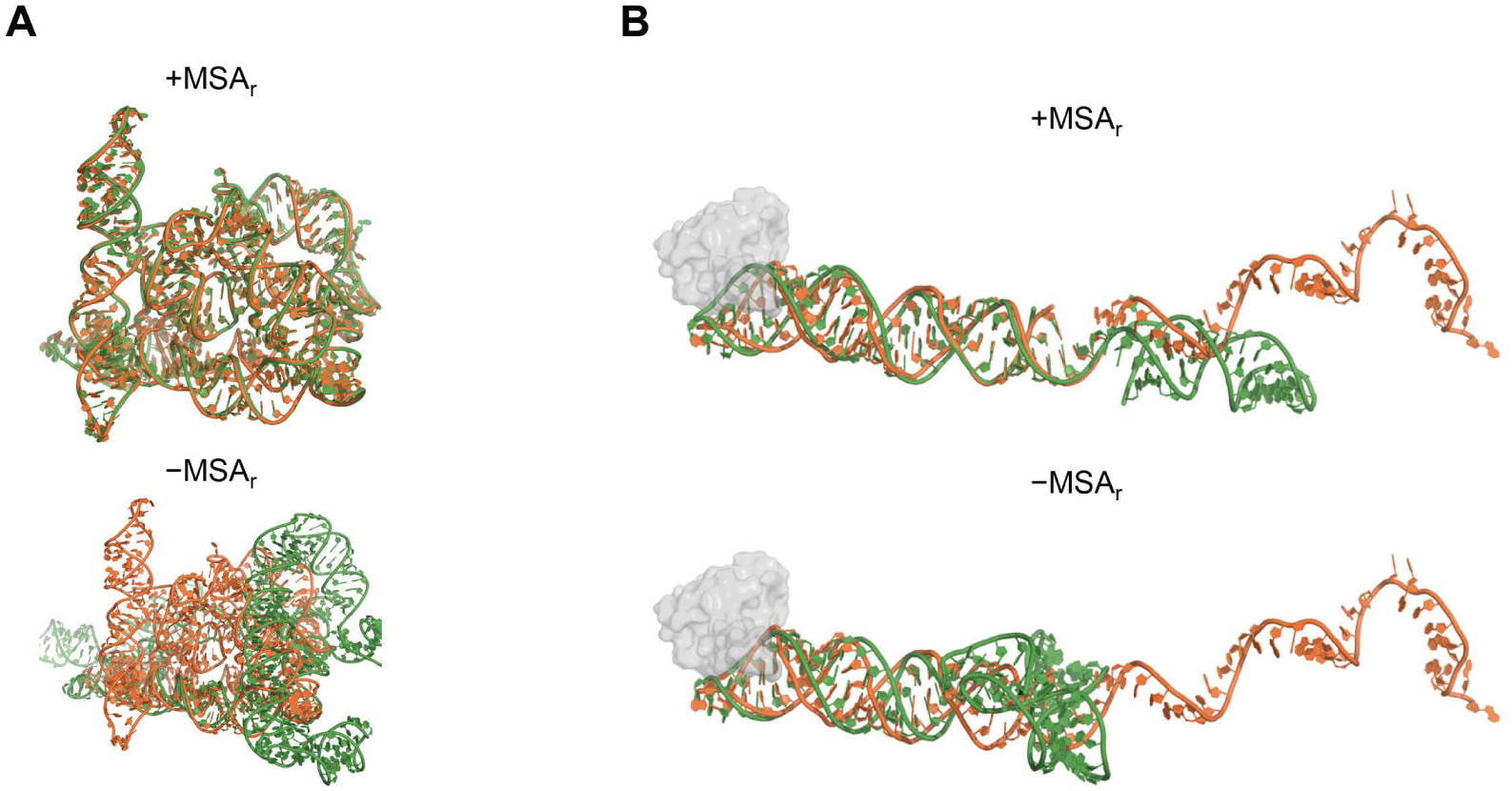
The RNA alignment decides the fold of single-chain and long protein-bound RNAs. Experimental structure (orange) and AF3 prediction (green) made with (+MSA*_r_*, top) and without (*−*MSA*_r_*, bottom) the RNA alignment, superposed on the RNA. The protein is a grey surface. **A**, Single-chain RNA from the RNA-only set, PDB 8OLS (LDDT*_r_* 0.94 *→* 0.49; TM*_r_* 0.99 *→* 0.22). B, Protein–RNA complex from the controlled ablation, with protein alignment and both templates present, PDB 7X34, 94 nt (LDDT*_r_* 0.85 *→* 0.71; TM*_r_* 0.41 *→* 0.27).

**Fig. S19:**
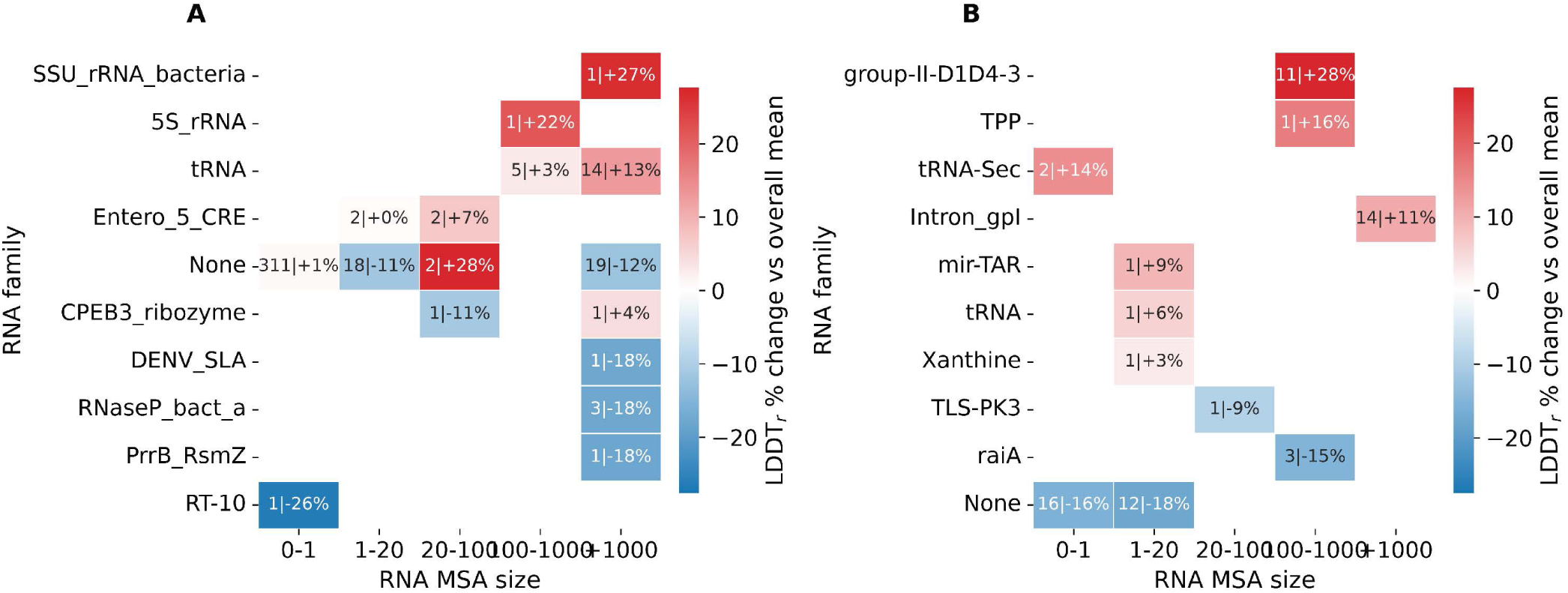
RNA-family dependence of LDDT*_r_*, stratified by alignment depth. Mean LDDT*_r_* per RNA family (rows) and RNA alignment-depth bin (columns), as the percentage change from the dataset mean (blue, above; red, below), annotated as count and change. **A**, Protein–RNA complexes (*n* = 384, mean 0.69). **B**, Single-chain RNAs (*n* = 63, mean 0.70). Each family occupies essentially one depth bin, so family and alignment depth are confounded.

**Fig. S20:**
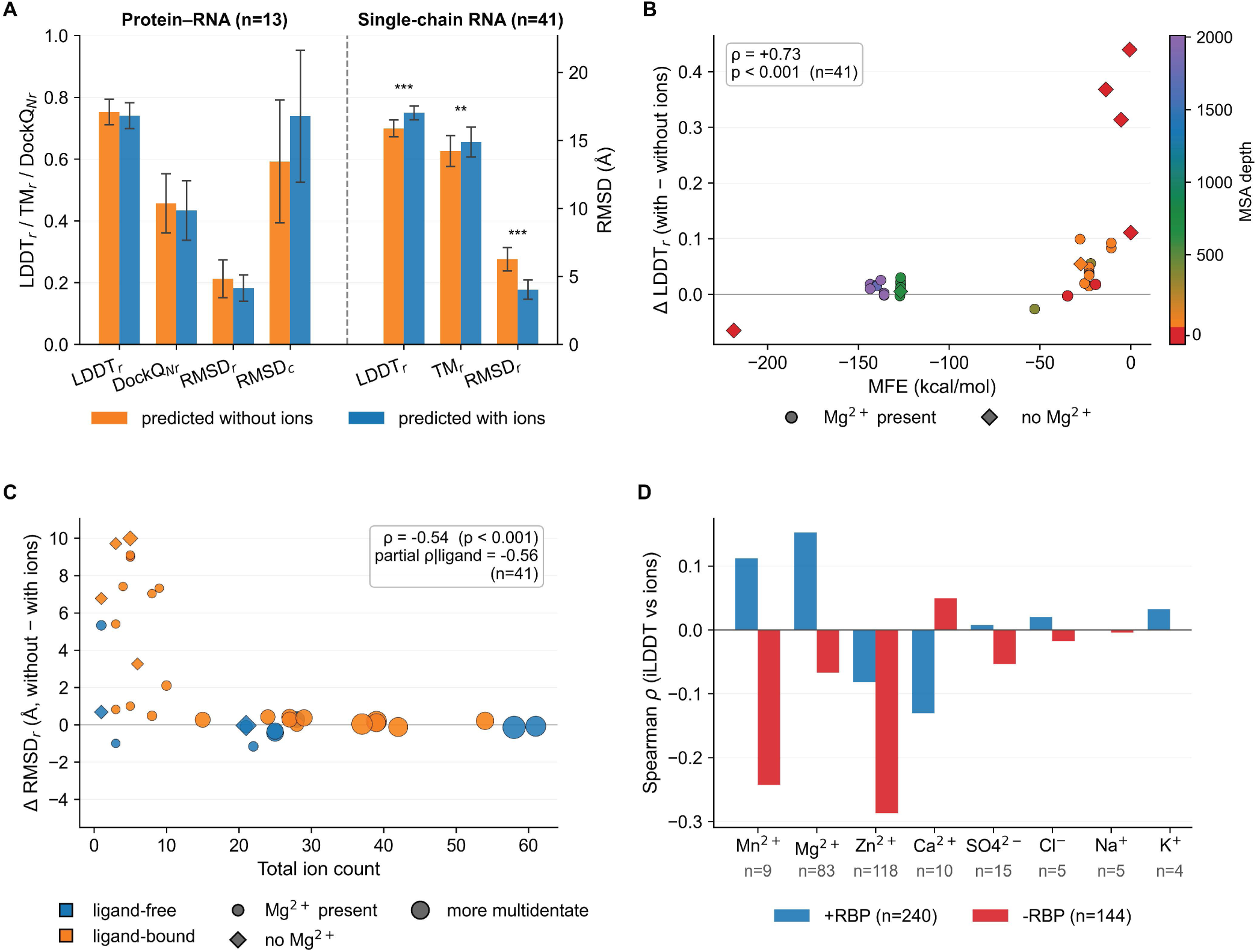
Ions improve single-chain RNA folds but not protein–RNA interfaces. **A**, Paired predictions without and with the deposited ions, for protein–RNA complexes (*n* = 13) and single-chain RNAs (*n* = 41). Bars show mean LDDT*_r_*, TM*_r_* and DockQ*_Nr_* (left axis) and RMSD (right axis) with s.e.m. (Wilcoxon signed-rank test). **B**, LDDT*_r_* gain against the predicted folding free energy, coloured by alignment depth (Spearman *ρ* = +0.73). **C**, RMSD*_r_* gain against the number of ions (Spearman *ρ* = *−*0.54). **D**, Spearman *ρ* between deposited ion count and interface LDDT per ion species, with and without an RNA-bound protein template.

### L.1. Ligand-bound ground truths

RNA is intrinsically unstable in aqueous solution because its phosphodiester backbone is susceptible to non-enzymatic hydrolysis, and bound ligands provide localised entropic and enthalpic stabilisation [86]. Consistently, a bound ligand marks an easier target rather than a harder one. Among the 384 protein–RNA complexes, those whose ground truth contains a ligand are predicted markedly better at the interface (DockQ*_Nr_* 0.451, *n* = 106, versus 0.309, *n* = 273; Mann-Whitney *P* = 9.2 *×* 10*^−^*^5^) and in the RNA chain (RMSD*_r_* 3.48 versus 5.83 Å, *P* = 2.6 *×* 10*^−^*^5^), while whole-complex fold quality is unchanged (TM*_c_* 0.817 versus 0.814, *P* = 0.071). The 66 single-chain RNA structures point the same way, though without reaching significance (RMSD*_r_* 6.42 Å with a ligand versus 8.50 Å without, *P* = 0.69). As with the ion result above, a bound cofactor seems to present a conformationally restrained, well-ordered pocket, which AF3 exploits.

### L.2. Solution conditions: pH, crowding and buffer

Beyond the deposited ions and ligands, each structure carries a record of the solution it was prepared in. Figure S22 collects those conditions, namely the pH-dependent bias in which charged residues AF3 places at the protein–RNA contact (**A**), buffer type and concentration (**B**), and PEG concentration and molecular weight (**C**, **D**). The pH analysis (**A**) uses the 111 crystallographic entries, since a standardised crystallisation pH is recorded for X-ray depositions alone, and PEG (**C**-**D**) is by construction a crystallisation additive. They are summarised in Section 3.5 and detailed below.

**Fig. S21:**
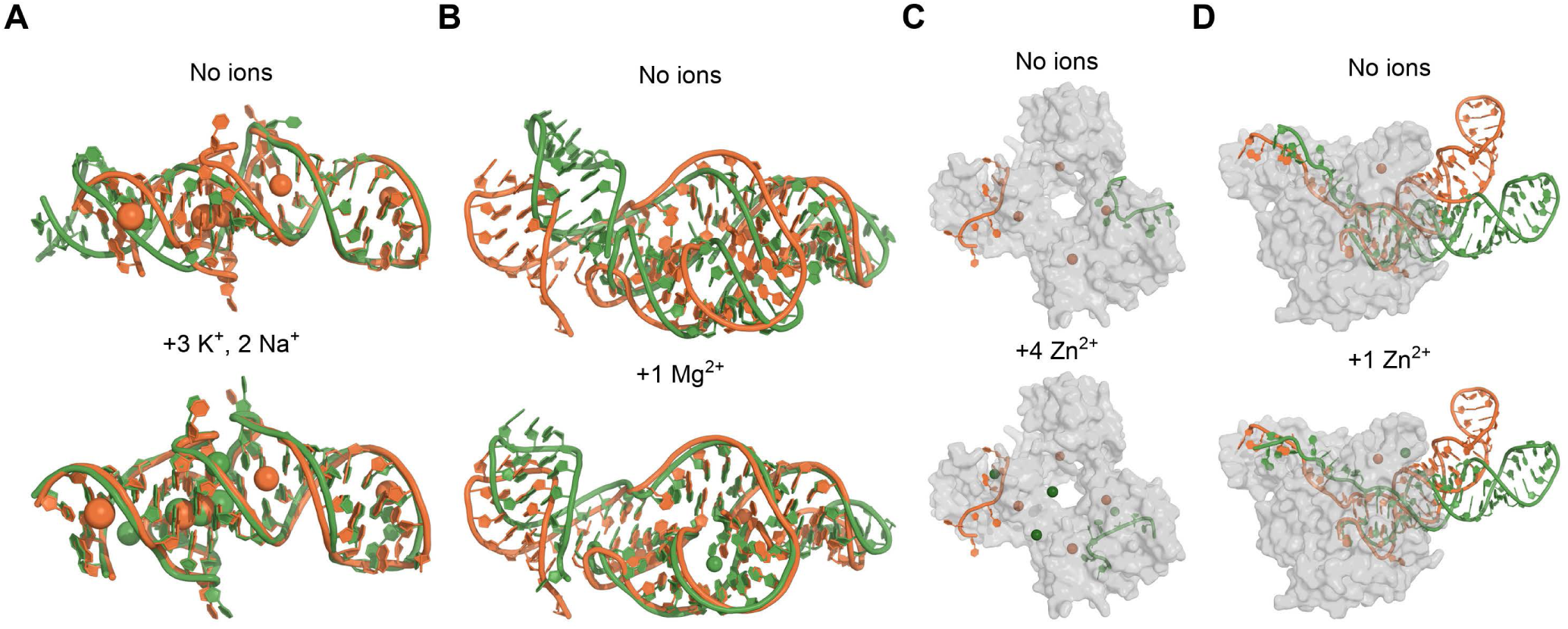
Ions rescue single-chain RNA folds but not protein–RNA complexes. Experimental structure (orange) and AF3 prediction (green) made without (top) and with (bottom) the deposited ions, whose number and species are given above each structure. Ions are spheres in the darker shade, and the protein is a grey surface. **A**, **B**, Single-chain RNAs from the RNA-only set, superposed on the RNA. **A**, PDB 8K7W (LDDT*_r_* 0.54 *→* 0.92). **B**, PDB 7U4A (LDDT*_r_* 0.69 *→* 0.78). **C**, **D**, Protein–RNA complexes superposed on the protein. **C**, PDB 7PDV, where AF3 binds the RNA to the wrong subunit with or without ions, and the added Zn^2+^ lie 11–19 Å from the deposited ones (LDDT*_r_* 0.69 *→* 0.63). **D**, PDB 8IDF, where the added Zn^2+^ lies 12 Å from its coordinating site and the RNA is unchanged (LDDT*_r_* 0.75 in both).

#### Buffer composition is confounded with the experimental method

Buffer information is populated for 109 of the 111 X-ray entries and for 104 of the 273 cryo-EM entries, but it does not record the same quantity in the two cases. For cryo-EM, it is the buffer the complex was kept in (typically 20-25 mM Tris or HEPES with 150 mM NaCl), whereas for X-ray it is the condition that drove crystallisation (precipitants, PEG and 0.1-2 M salts).

Among the 109 crystallographic structures neither buffer identity nor concentration is associated with accuracy at any level (*P ≥* 0.07 for TM*_c_*, TM*_cr_* and DockQ*_Nr_*), and the low-concentration group contains only nine structures. Among the 104 cryo-EM structures, where identity varies at essentially fixed concentration, buffer identity does track RNA placement within the assembly (TM*_cr_*, Kruskal-Wallis *P* = 0.002), with HEPES at 0.509 (*n* = 66) versus Tris 0.378 (*n* = 32), *P* = 0.0019. This is consistent with HEPES being relatively inert while Tris can engage charged side chains and perturb the electrostatic environment of RNA binding [87, 88]. The effect appears only in TM*_cr_* and not in TM*_c_* (*P* = 0.69) or DockQ*_Nr_* (*P* = 0.31), so it concerns RNA accuracy rather than assembly or interface quality, and Tris-buffered entries are on average slightly larger assemblies (4.2 versus 3.5 protein chains), so system size may contribute. We therefore treat buffer identity as a suggestive cryo-EM-only association rather than an established determinant.

#### Molecular crowding by PEG

The influence of polyethylene glycol (PEG) in crystallographic structures is dominated by a strong, non-linear coupling between concentration and molecular weight rather than by simple linear trends. Complex accuracy declines as PEG concentration rises (TM*_cr_* from *∼* 0.45 PEG-free to *∼* 0.26 at 30-50%; Spearman *ρ* = *−*0.29, *P <* 0.0001; Fig. S22C). A similar but less pronounced effect is seen with increasing PEG molecular weight (*ρ* = *−*0.26, *P* = 0.0001; the largest-MW bins are sparse, *n ≤* 4; Fig. S22D). This decline is amplified by the MW *×* concentration interaction, which explains *∼* 27-38% of the variance in global metrics (*η*^2^ up to 0.38 for LDDT*_c_*, *p* = 1.7 *×* 10*^−^*^6^). Mid-MW PEG (2-10 kDa) at high concentration (20-50%) markedly degrades accuracy (TM*_c_ ∼* 0.85 *→∼* 0.70), whereas low-MW PEG (*≤* 1 kDa) can improve it, consistent with a competition between crowding-induced distortion and mild stabilisation. Direct PEG–RNA association is consistent with this. It has been reported that PEG of 3350 and 6000 Da binds tRNA through its bases and backbone phosphates, more strongly at the higher molecular weight, and drives aggregation into multi-tRNA particles at high polymer concentration, while the RNA itself keeps its A-family fold [89]. The interface shows the same, but weaker and non-significant, crowding effect-driven decline. After the two anomalous two-sample bins run against the trend (PEG 0-10%, *n* = 2, DockQ*_Nr_* = 0.08; PEG *≥* 10 kDa, *n* = 2, 0.94) were removed, the binned DockQ*_Nr_* means trend downward with both PEG concentration (PEG-free 0.41 *→* 0.36 *→* 0.34 *→* 0.30 from 0 to 30-50%; Spearman *ρ* = *−*0.12, *P* = 0.20, *n* = 122) and PEG molecular weight (PEG-free 0.41 *→* 0.38 *→* 0.36 *→* 0.31 up to 2-10 kDa; *ρ* = *−*0.16, *P* = 0.07, *n* = 127), but neither association is significant. Global structure metrics (LDDT*_c_*, TM*_c_*) are far more sensitive to crowding than interface metrics are, so PEG acts on the conformational landscape of the assembly rather than on the contacts within it.

#### pH dependence

For the pH correlation analysis, 111 X-ray structures were evaluated, of which 66, 26 and 19 samples were neutral, acidic and basic, respectively, with pH values spanning 3.8-9.5. Higher pH levels correlate with improved structural accuracy and increased model confidence. LDDT*_c_* (*ρ* = +0.328*, p <* 0.001) and TM*_c_* (*ρ* = +0.223*, p* = 0.019) exhibit strong positive Spearman trend with pH. This is mirrored by a decline in model uncertainty as evidenced by a negative correlation between pH and chain-pair PAE (*ρ* = *−*0.305*, p* = 0.001). The interface characterised by DockQ*_Nr_* shows no dip in acidic conditions (*≈* 0.35 below pH 6 versus *≈* 0.36 at pH 6-8). While RMSD*_r_* (*r* = *−*0.203*, p* = 0.033) decreases as pH increases, TM*_r_* and LDDT*_r_* show negligible correlation with pH.

At the interface prediction errors show clear pH-dependent biases. Key charged and polar residues (ARG, LYS, ASP) shift from underprediction in acidic conditions (e.g., ARG: *−*1.667, LYS: *−*1.267, ASP: *−*0.467 at pH 5–6) to slight overprediction in neutral-to-basic conditions (LYS: +0.519, ASP: +0.222 at pH 7–8; ARG: +0.895 at pH *>* 8; Fig. S22A). HIS contacts also increase with pH (Spearman *ρ* = +0.196, *p* = 0.040) and are underpredicted in acidic conditions (*−*0.400 at pH 5–6). This suggests that AF3 does not fully capture the stabilising role of these residues, most likely because it does not model protonation states

**Fig. S22:**
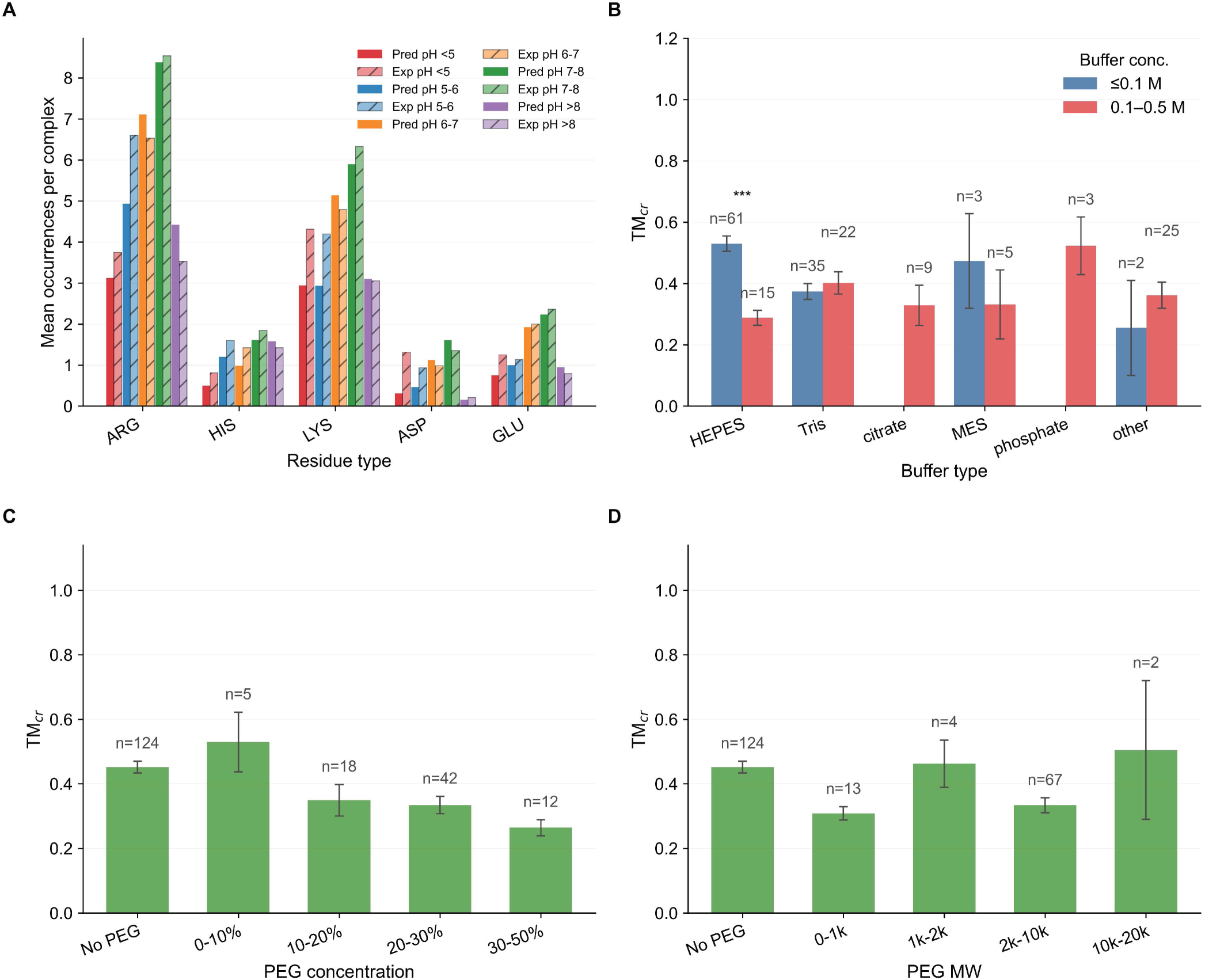
Solution conditions shape AF3 protein–RNA predictions. **A**, Charged residues per complex in predicted (solid) and experimental (hatched) structures, by experimental pH. **B**, TM*_cr_* by buffer and buffer concentration (mean *±* s.e.m.). Only HEPES shows a concentration effect (*P <* 0.0001). **C**, TM*_cr_* against PEG concentration (*n* = 201, Spearman *ρ* = *−*0.29). **D**, TM*_cr_* against PEG molecular weight (*n* = 210, *ρ* = *−*0.26).

## M. Protein–RNA structure and interface classification

Every predicted complex was classified in two stages, first by *structure* (global fold quality) into four quadrants, then, within each quadrant, by *interface* quality. The scheme is implemented in get_sample_audit_global_fold_interface_lists in evaluation/evaluationMetrics.py. Each exp_db_id sample contributes a single model, the one maximising 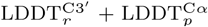. Models with a non-finite value for either are dropped. Global accuracy is read from the selected model’s whole-RNA 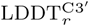 and TM , so the longest RNA chain enters only through the chain length used by the class-**(i)** TM*_r_* gate. The interface is graded on DockQ*_Nr_* , the mean of the per-pair RBP_DockQ values over *all* protein–RNA chain pairs listed in ChainIDpairList_proteinRNA.

Replacing the mean by the median leaves DockQ*_Nr_* essentially unchanged across RNA chains (Spearman *ρ* = 0.95, with 81% of chains keeping the same a/b/c grade), and the fold-interface decoupling holds under mean, median or max aggregation alike, with 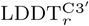 explaining 20-26% of DockQ variance in all three. These checks are quantified by `regression_analysis/analysis_dockq_aggregation_robustness.py`.

*Structure.* The fold quadrants **(i)** both components well folded, **(ii)** RNA only, **(iii)** protein only and **(iv)** neither use the thresholds defined in Methods, Section 2.3. The implementation additionally requires TM*_p_* to be finite, and requires TM*_r_* to be finite as well as above 0.5 for the class-**(i)** long-RNA gate, so that a missing global score cannot silently promote a model into a higher class.

*Interface.* Within each quadrant the interface is graded by DockQ*_Nr_* into *(a)* good, *(b)* intermediate and *(c)* poor at the Methods thresholds (Section 2.3), with models having no finite DockQ*_Nr_* set aside. The poor interfaces *(c)* are split by whole-complex RMSD into *(ca)* RMSD*_c_<* 10 Å, where the assembly is approximately co-located but the RNA is locally misplaced, and *(cb)* RMSD*_c_≥* 10 Å, where the RNA lies far from its native position.

These classifications are visualised for the full data set (*n* = 384) in Fig. S23. Its panels show **(A)** the 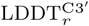 versus 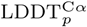 quadrant scatter, **(B)** the DockQ_pr_ distribution per quadrant, and **(C)** the longest-chain 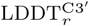 and TM against the interface class. Fig. S25 compares the per-RNA-chain and per-protein–RNA-interface accuracy with the single whole-complex metrics. The longest-chain 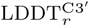 and TM already capture complex-level RNA accuracy, whereas the whole-complex interface metrics are inflated by protein–protein interfaces (Fig. S24), so the protein–RNA-only means should be preferred.

When a complex contains several RNA chains, the LDDT and TM-score of the single largest RNA chain already summarise the RNA accuracy of the whole complex well, as panels **A** and **B** lie tightly on the diagonal (*r* = 0.95, *r* = 0.96). Notably, a *correct* protein–RNA interface (green) tracks the local RNA geometry far more than the global RNA fold. Correct-interface complexes have a high median LDDT*_r_* (0.82 versus 0.51 for the rest) but only a moderate TM*_r_* (0.44 versus 0.28), so LDDT*_r_* separates correct-interface complexes better than TM*_r_* (AUC 0.82 versus 0.77). This is consistent with interface correctness being a local-contact rather than a global-fold property. In contrast, the whole-complex interface metrics (iLDDT, ICS, DockQ, QS) are poor descriptors of the protein–RNA interfaces, because they are dominated by the additional protein–protein interfaces present when multiple protein chains contact each other. The protein-single-chain RNA means (e.g. DockQ*_Nr_*) should therefore be preferred when reporting protein–RNA interface quality.

### M.1. RNA motif recovery from the interface

With exact residue mapping (Section A.6) the contact-recovery scores measure, residue by residue, whether AF3 contacts the same protein residues and RNA nucleotides as the experimental structure. Both therefore track interface quality (Spearman *ρ* over post-cutoff complexes). Protein sequence match correlates with DockQ*_Nr_* at *ρ* = +0.71, RNA sequence match at +0.59 and the alignment-based

RNA motif score at +0.43, so the RNA sequence match is the more informative RNA-side readout. Protein contact recovery also goes with the protein fold (LDDT*_p_* +0.37, TM*_p_* +0.38, pLDDT*_p_* +0.46) and with interface LDDT (+0.51). RNA contact recovery is only weakly related to the global RNA fold (LDDT*_r_* +0.20, TM*_r_* +0.15), falls with RNA length (*−*0.35) and RNA MSA depth (*−*0.23), and is best anticipated by AF3’s interface confidence (ipTM*_r_* +0.65, ipTM*_p_* +0.61, pLDDT*_r_* +0.56). A high protein contact match with a failed interface is rare, as only 3 complexes reach a protein match *>* 0.7 with DockQ*_Nr_ <* 0.23, against 43 on the RNA side. Where interfaces fail, the protein contact residues are therefore largely lost as well. How the RNA is then placed is analysed per interface in Fig. 3. Per interface, the contact match separates the placement classes (Fig. S26C,D). Correct interfaces recover most native contacts on both sides (median protein contact match 0.78 pre-and 0.66 post-cutoff; RNA contact match 0.85 and 0.80). When AF3 binds the correct site with an incorrect RNA, it keeps about half of the native protein contacts (median 0.50 and 0.46; at least 0.1 by definition) but only a fraction of the native RNA contacts (0.20 and 0.29), so the correct protein site is contacted largely by other RNA nucleotides than in the experimental structure. In all other failure classes AF3 contacts essentially none of the native residues on either side (median 0), consistent with the RNA sitting on a shifted or distal site or on a mispredicted protein binding region. Slid duplexes are the exception on the RNA side after the cutoff (protein 0.04, RNA 0.62).

**Fig. S23:**
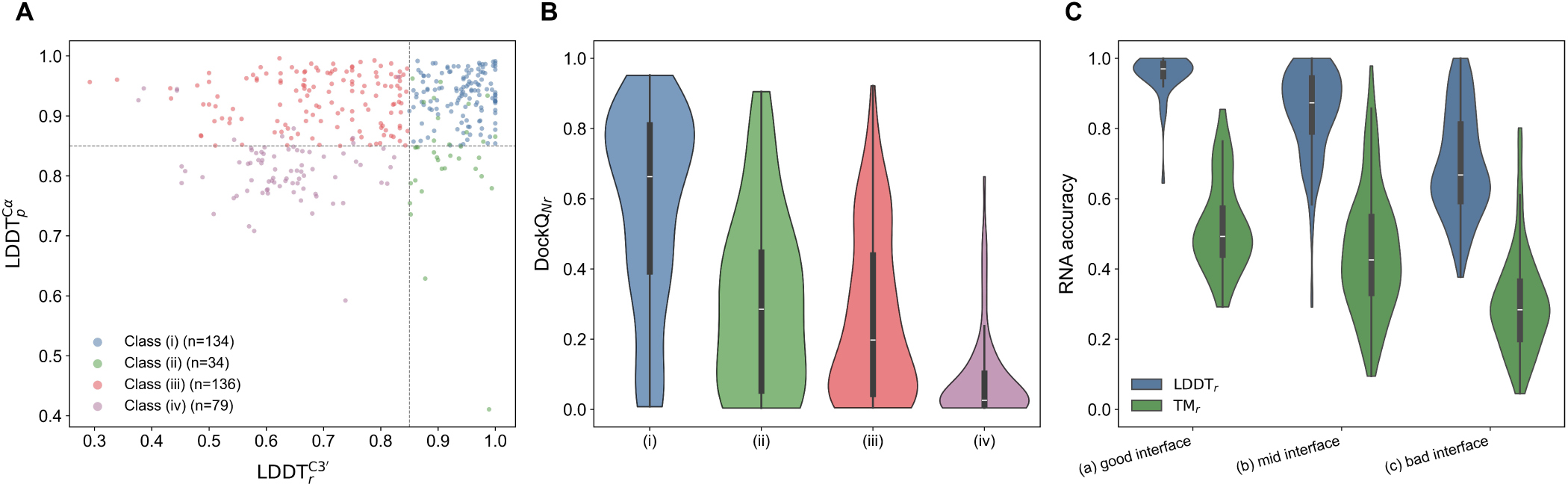
Classification of AF3 protein–RNA predictions by global accuracy and interface quality. Post-cutoff predictions (*n* = 384) fall into four fold classes (i, RNA and protein accurate; ii, RNA only; iii, protein only; iv, neither), and the interface is graded by DockQ_Nr_ into good (a, *≥* 0.8), intermediate (b) and poor (c, *<* 0.23). **A**, 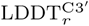 against 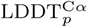, coloured by class (dashed lines, 0.85). **B**, DockQ*_Nr_* by class. **C**, 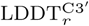 and TM by interface subclass. The subclass composition within each class is shown in Fig. 3A.

**Fig. S24:**
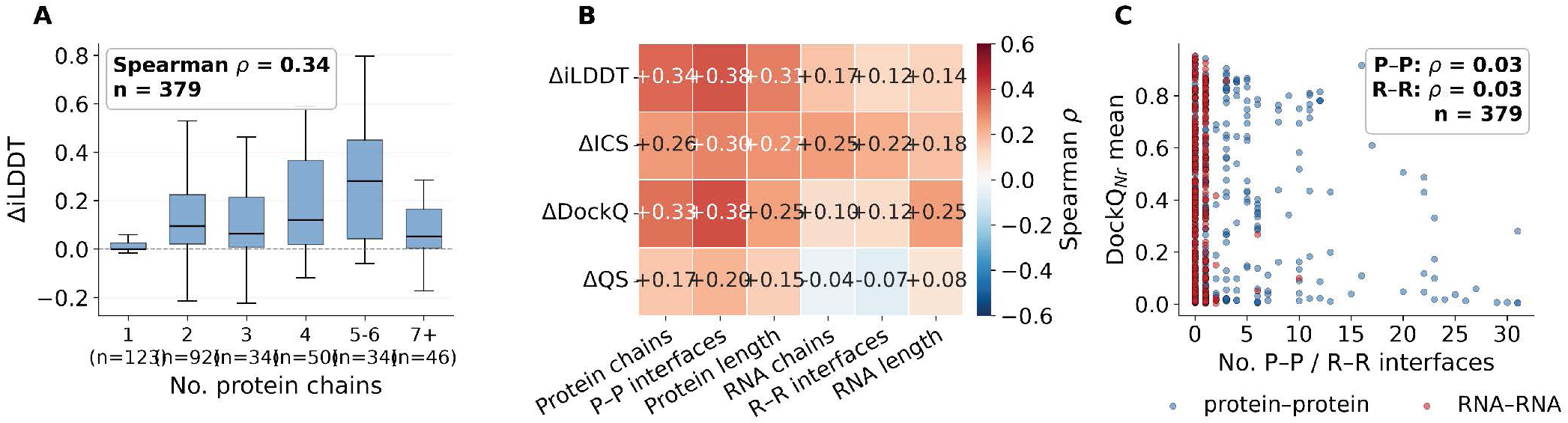
Protein–protein interfaces inflate whole-complex interface scores but do not lower protein–RNA interface quality. The gap Δ is the whole-complex value of a metric minus its mean over the protein–RNA interfaces. **A**, ΔiLDDT against the number of protein chains (Spearman *ρ* = 0.34). **B**, Spearman *ρ* of each gap with six composition drivers. Every gap tracks the number of protein–protein interfaces most closely. **C**, DockQ*_Nr_* against the number of protein–protein (blue) and RNA–RNA (orange) interfaces. Both trends are flat (*ρ* = 0.03 each), so protein–RNA interface quality should be reported with the protein–RNA means.

**Fig. S25:**
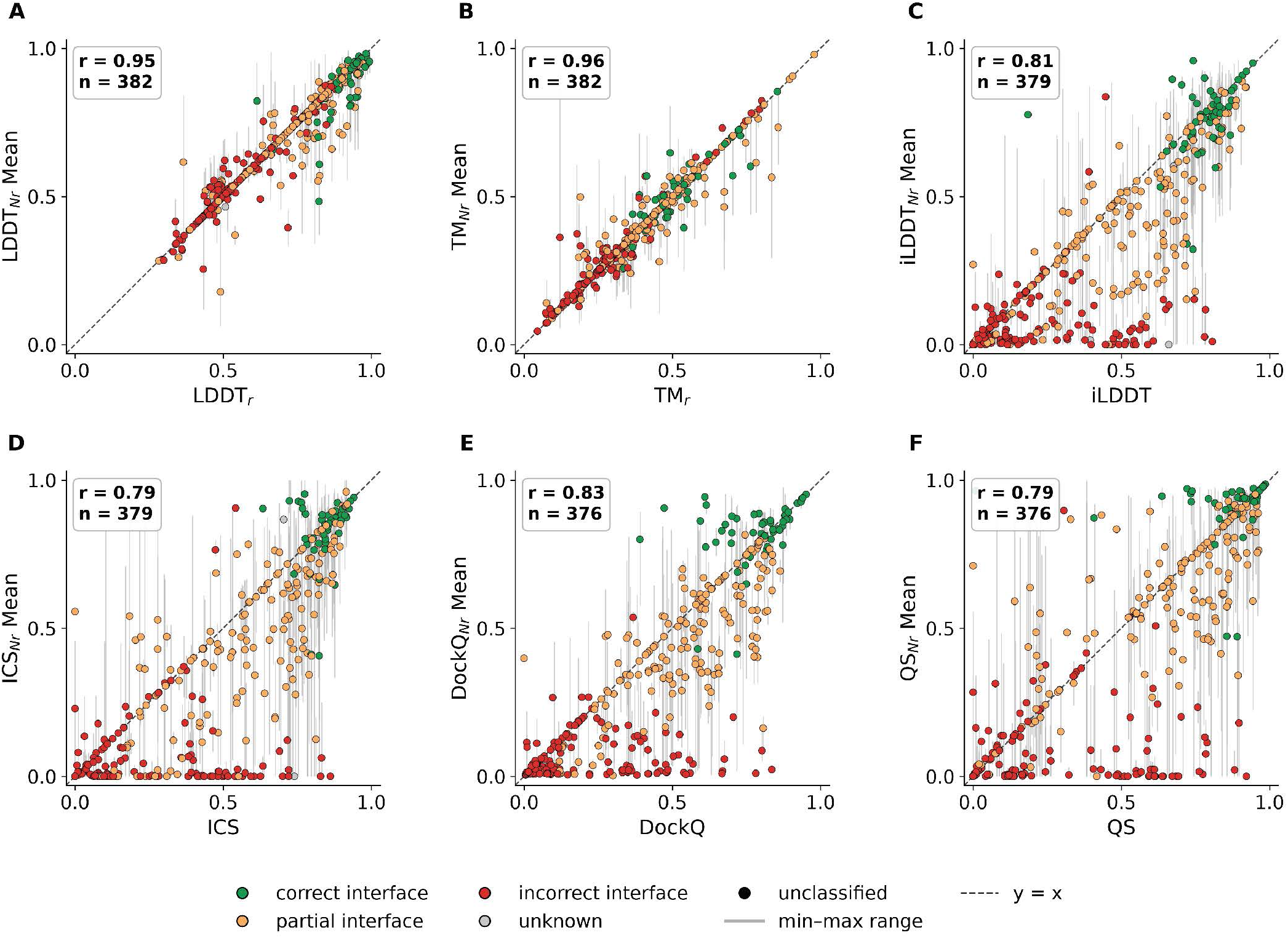
Per-chain and per-interface accuracy against the whole-complex scores. Each point is one complex. The *y* axis is the mean over all RNA chains (A, B) or over all protein–RNA interfaces (C–F), with grey bars for the range, and the *x* axis the whole-complex value (A, B, the longest RNA chain). **A**, LDDT. **B**, TM-score. **C**, iLDDT. **D**, ICS. **E**, DockQ. **F**, QS-score. Complexes with one RNA chain or one protein–RNA interface lie on the dashed line *y* = *x* (*r*, Pearson correlation).

**Fig. S26:**
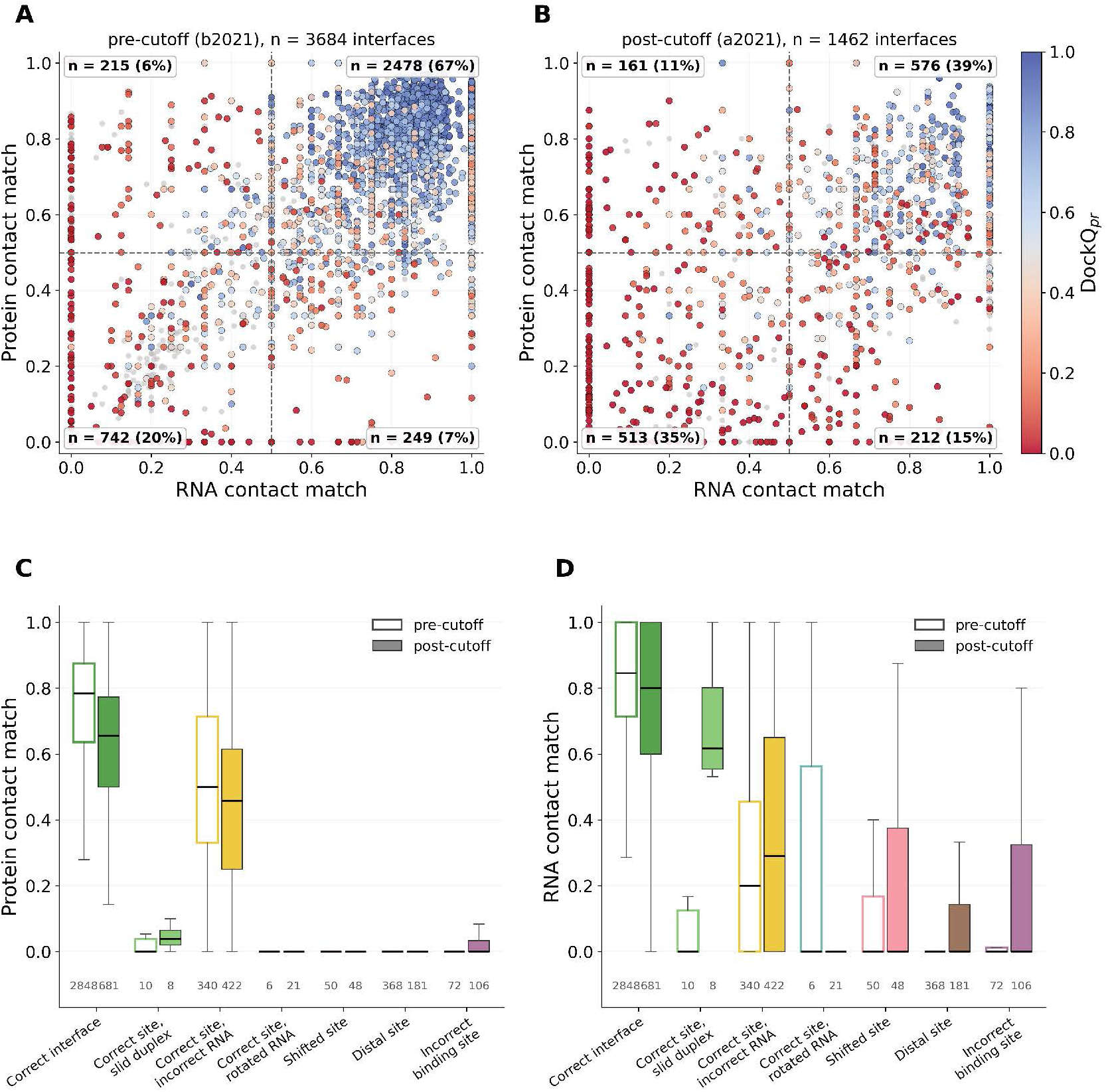
Per-interface contact recovery and RNA placement. **A**, **B**, RNA contact match against protein contact match per native protein–RNA interface, coloured by its DockQ from red (low) to blue (high), before (A, *n* = 3,684) and after (B, *n* = 1,462) the cutoff. Interfaces that AF3 does not form score 0. **C**, **D**, Protein (C) and RNA (D) contact match per placement class of Fig. 3B (open, pre-cutoff; filled, post-cutoff). A protein contact match *≥* 0.1 defines the correct-site class, so panel C separates the classes partly by construction.

**Fig. S27:**
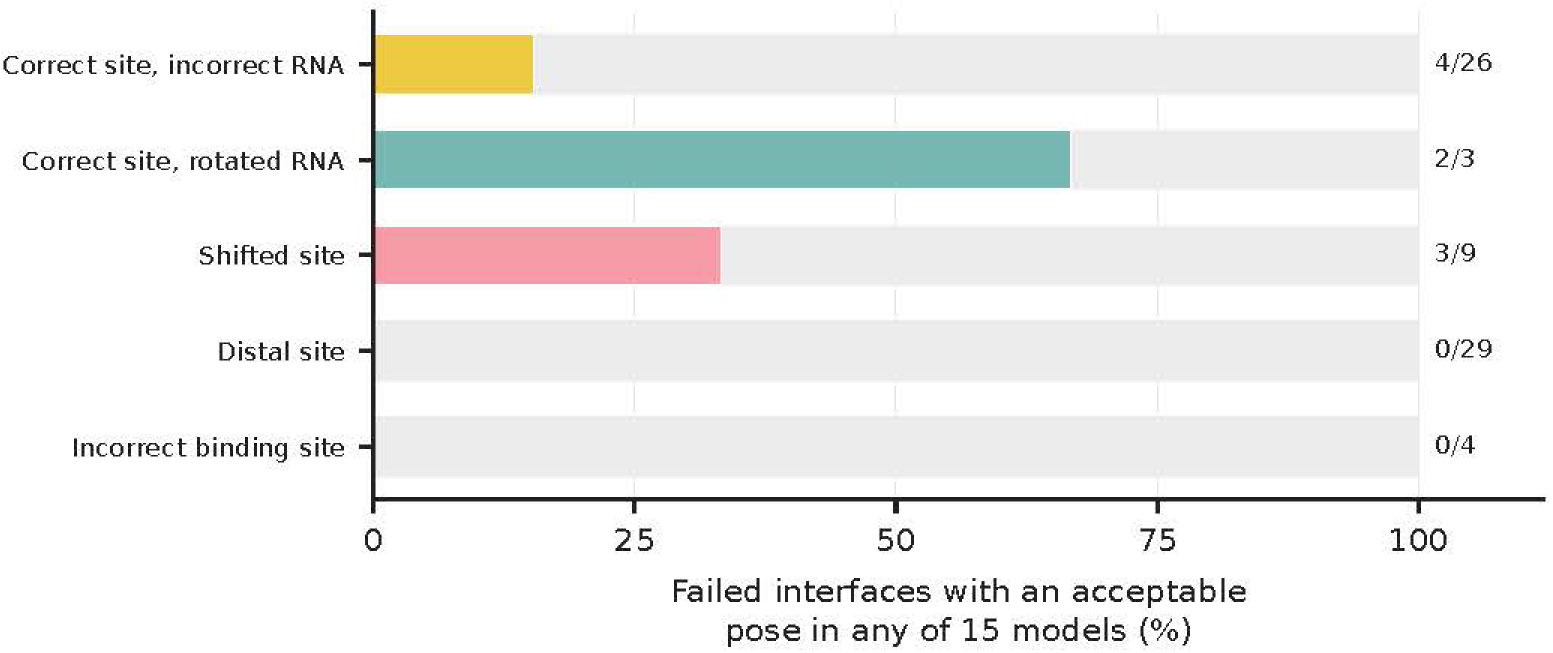
Would another sample have recovered the interface? For the 49 post-cutoff targets predicted with three seeds and five models each (134 protein–RNA interfaces), the share of the interfaces that fail in the default model (seed 1, model 0; DockQ*_pr_ <* 0.23) for which any of the 15 models reaches DockQ*_pr_ ≥* 0.23, by placement class (recovered/failed per bar). The nine failed interfaces without a placement class are not recovered either, and AF3’s ranking score selects an acceptable model in only 2 of the 9 recoverable cases.

This appendix details the interface analyses summarised in Section 3.3.

112 of the 133 gradable class-i predictions (84%) form a correct interface (DockQ*_Nr_ ≥* 0.23), but only 35 (26%) reach DockQ*_Nr_ ≥* 0.8, 21 (16%) fail, and the class-i median of 0.656 lies below the 0.8 threshold of a good interface. Accordingly, global fold accuracy and interface correctness are only loosely coupled. TM*_r_* explains only about a quarter and LDDT*_r_* about two fifths of the variance in DockQ*_Nr_* (Spearman *ρ*^2^ *≈* 0.26 and 0.41; Supplementary Fig. S23C), and a globally accurate RNA can still fail at the interface. In complex 7V6C, for instance, the RNA is folded perfectly (LDDT*_r_* = 1.0) yet its protein–RNA interfaces are almost entirely unrecovered (DockQ*_pr_* 0.003-0.005), even though protein–protein interfaces in the same assembly are well predicted (DockQ*_c_* 0.77-0.82). This is a case where the DockQ*_c_* would badly overstate protein–RNA quality. Conversely, as one of the few exceptions to this rule, 8VIV recovers a near-native protein–RNA interface (DockQ*_pr_* = 0.92, iRMSD = 0.55 Å) despite a poor RNA fold (LDDT*_r_* = 0.64, TM*_r_* = 0.29). LDDT*_r_* and TM*_r_* score the whole RNA, so a correctly folded binding segment can dock correctly while the rest of the RNA is misfolded. The error lies mainly in how the RNA is placed. In the most frequent failure, AF3 still contacts the native protein residues but places a different part of the RNA at the site (median protein contact match 0.46, nucleotide contact match 0.29; Section 3.3), so it identifies the protein side of the interface better than the RNA side. Most post-cutoff failures moreover preserve approximate global colocalisation (RMSD*_c_ <* 10 Å for 111 of the 177 poor interfaces) while misplacing the RNA locally. AF3’s characteristic interface failure after the cutoff is therefore incorrect intermolecular placement rather than loss of assembly.

To resolve how a failed interface fails, we followed every protein–RNA interface individually and carried the experimental contact residues onto the prediction by exact residue mapping (Section A.6). Recovery of the contacting protein residues and RNA nucleotides tracks the quality of each interface (DockQ*_pr_*, Spearman *ρ* = 0.64 and 0.71 post-cutoff; Supplementary Fig. S26) and falls on both sides after the cutoff. Both sides are recovered (contact match *≥* 0.5) for 67% of the pre-cutoff but only 39% of the post-cutoff interfaces, and both are lost for 20% and 35%. Contact recovery alone cannot, however, tell an RNA bound to another region of the protein from one bound at the right site in the wrong orientation. We therefore classified each interface by where AF3 places the RNA relative to its protein partner (Section B.1; Fig. 3B,C). The class frequencies are given in Section 3.3. After the cutoff 46% of the interfaces are correct (DockQ*_pr_ ≥* 0.23), against 77% before it. The dominant failure is a correct site with an incorrect RNA. AF3 contacts native protein residues, but the RNA is turned and displaced (median rotation 97*^◦^*, LRMSD 32.5 Å). The RNA is usually also misfolded in these cases, with a median LDDT*_r_* of 0.53 and only 4% of them well folded, against 0.82 and 43% at correct interfaces, so the wrong pose accompanies a wrong fold rather than a correctly built RNA placed badly. This accounts for 54% of the post-cutoff failures (29% of all post-cutoff interfaces, against 9.2% before the cutoff). It is also the only placement class that becomes more frequent as the RNA gets longer (Spearman *ρ* = +0.20, *P* = 3 *×* 10*^−^*^15^, median 37 nt against 25 nt at correct interfaces), while the other failure classes are flat or slightly commoner in short RNAs. Before the cutoff this class has no length dependence (*ρ* = *−*0.004) and longer RNAs are predicted better (*ρ* = +0.16 for correct interfaces). Thus, length becomes a liability only for novel targets, and only through this failure mode. In a further 13% of the failures the binding site is incorrect because the protein’s own binding region is mispredicted. A distal site, with the RNA more than 20 Å from its native position on the protein (median 43 Å), accounts for 23% of the post- and 43% of the pre-cutoff failures, whereas shifted sites (10-20 Å, 6%), rotated RNAs at the correct site (3%) and slid duplexes (1%) are rare. This ordering does not depend on the class thresholds. Over all 1,280 combinations of the five classification thresholds, a correct site with an incorrect RNA remains the largest failure class in every setting (42-61% of post-cutoff failures), ahead of distal sites (17-35%) and incorrect binding sites (8-21%; Section B.1). The rise in failures after the cutoff is thus driven by misposed RNAs at the correct protein site rather than by docking to a distal site. The two dominant failure modes are not equally visible to AF3 (Fig. 3D). At the complex level, a distal site goes with a lower interface confidence (ipTM*_r_ ρ* = *−*0.39, ipTM*_p_ −*0.31). An incorrect RNA at the correct site is only weakly reflected in ipTM*_r_* (*−*0.27), although it is strongly associated with the true local accuracy (LDDT*_r_ −*0.53, iLDDT *−*0.59) and becomes more frequent with RNA length (+0.34). AF3’s confidence therefore flags distally placed RNAs but largely misses its most frequent post-cutoff failure. Better ranking would not recover these interfaces either. For 49 post-cutoff targets predicted with three seeds and five models each (15 samples; Appendix C), 71 of the 80 failed default interfaces (89%) have no acceptable pose in any sample. The best of all 15 samples raises the share of acceptable interfaces only from 40% to 47%, so most failures are failures of modelling, not of ranking or sampling. Distal sites are never recovered (0*/*29) and misposed RNAs at the correct site rarely are (4*/*26).

#### Structural examples

Fig. S28 shows two failed interfaces of each placement class, together with two complexes whose protein partners are themselves misarranged. In the first five classes the protein is predicted in place, and only the RNA differs. In an incorrect binding site the protein’s binding region is itself misfolded.

**Fig. S28:**
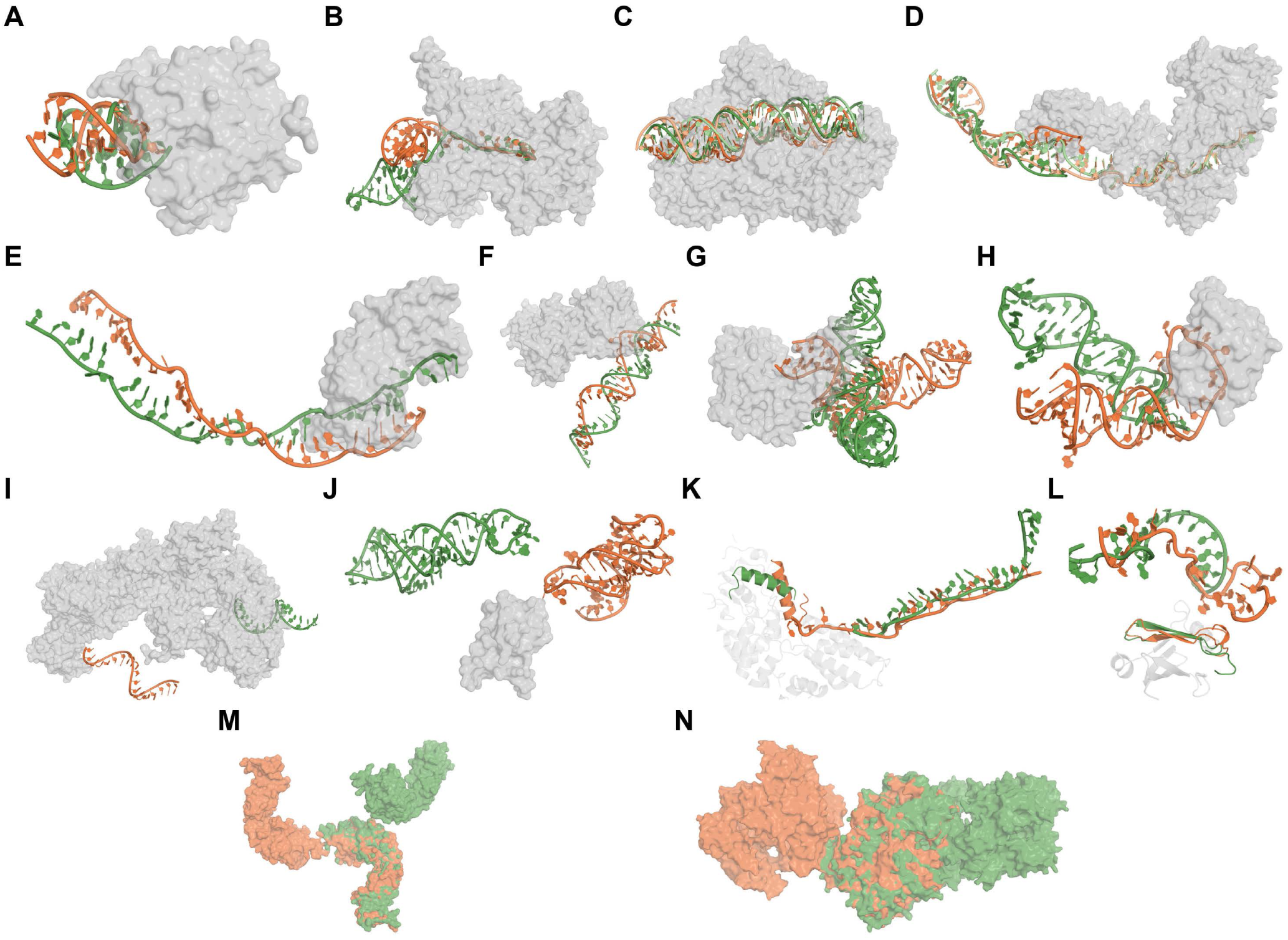
How failed protein–RNA interfaces fail. All fourteen examples are protein–RNA complexes. Experimental RNA (orange) and AF3 RNA (green), superposed on the experimental protein (grey surface). **A**, **B**, Correct site, incorrect RNA (PDB 8H1B and 8E29). **C**, **D**, Correct site, slid duplex, with both strands shown and the native strand of the interface and its predicted chain in the darker shade (PDB 7VG2 and 8S9X). **E**, **F**, Correct site, rotated RNA (PDB 8YHE and 7TJ2). **G**, **H**, Shifted site (PDB 8KDA and 7DLZ). **I**, **J**, Distal site (PDB 7V6C and 8JY0). **K**, **L**, Incorrect binding site, where the protein segment that binds the RNA (orange and green cartoon) is misfolded (PDB 7XSS and 8J62). **M**, **N**, Misarranged protein partners, shown as experimental (orange) and AF3 (green) protein surfaces superposed on one protein chain (PDB 8VMA and 7OS0).

#### Duplexes and swapped copies

Because many failed interfaces involve several RNA chains or identical protein copies, we tested whether they are mapping artefacts, in which AF3 builds the native arrangement on the partner strand of a duplex or on another copy (evaluation/interface_symmetry_check.py). With the predicted protein superposed on the native one, as for the placement classes, we measured the share of the native RNA’s C4*^′^* atoms that have a predicted C4*^′^* atom within 4 Å. This was done for the predicted chain of the RNA itself, for the partner strand of a duplex (at least four base pairs between the native strands, purine N1 to pyrimidine N3 *≤* 3.5 Å), for an identical RNA copy, and for the RNA with an identical protein copy superposed instead. The RNA itself occupies a median 84% of its native position at correct interfaces and 11% at failed interfaces. A partner strand or copy takes the native position (*≥* 50%) for only 3.8% of the post-cutoff correct sites with an incorrect RNA, 8% of distal sites, 6% of incorrect binding sites and 4% of shifted sites and rotated RNAs, so the placement classes are not mapping artefacts. Of the rotated RNAs, 8 of 21 are duplex strands, and for 6 of these 8 the predicted duplex still covers at least 30% of the native duplex (median 41%) while the strand itself lies elsewhere. AF3 places the duplex at the native site but turned about its axis, so the protein faces the other strand. Where the predicted duplex covers at least half of the native one, the duplex is placed as a whole (Section B.1). Eight post-cutoff interfaces then turn out to be duplexes slid by 2–6 Å along their own axis with the strands exchanged, which the strand alone had shown as rotated (5), shifted (2) or in place with the wrong contacts (1). They form the slid-duplex class, in which the strand at the native position recovers 32–88% of the native protein contacts (median 64%) but always has a different sequence. At correct sites with an incorrect RNA the predicted duplex covers a median 41% of the native one, whereas at distal sites (11%) and incorrect binding sites (5%) it lies mostly elsewhere.

#### RNA conformation and base pairing

Base-pair fidelity and rare backbone conformations separate the two dominant failure modes (Table S3; evaluation/interface_class_ rna_geometry.py). Across post-cutoff complexes, the share of interfaces that are correct sites with an incorrect RNA rises with the target’s share of *syn* nucleotides, sugar puckers other than C3*^′^*-*endo* and *γ* angles other than g^+^ (*ρ* = 0.25-0.29) and with AF3’s under-prediction of these conformations (*ρ* = *−*0.22 to *−*0.33), and falls with the fidelity of non-Watson-Crick pairs (*ρ* = *−*0.50) and stacking (*−*0.36) more steeply than with that of Watson-Crick pairs (*−*0.24). Complexes whose interfaces are mostly of this class have targets with 50% non-C3*^′^*-*endo* puckers (21% for complexes with mostly correct interfaces), of which AF3 predicts 13 percentage points fewer, and recover non-Watson-Crick pairs (median INF 0.58 against 0.92) and stacking (0.12 against 0.53) poorly. The shares of distal sites and incorrect binding sites are only weakly related to these measures (*|ρ| ≤* 0.23), consistent with the misplaced or misfolded proteins of Table S8. The measures are pooled over the RNA chains of a complex and describe associations. Flexible, single-stranded RNAs are both richer in non-canonical conformations and harder to dock.

**Table S3.** RNA base pairing and conformation by interface placement (post-cutoff complexes). Left, Spearman *ρ* between the share of a complex’s protein–RNA interfaces in a placement class and each RNA measure (*^∗^*, *P <* 0.001; *n* = 118-384 complexes with the measure). Right, medians for complexes whose interfaces are mostly correct or mostly correct sites with an incorrect RNA. INF, interaction network fidelity of the predicted base pairs or stacks (complexes whose target has none are excluded). Target shares come from the experimental structure, and AF3 *−* target is the predicted minus target share (negative, the rare conformation is under-predicted). Measures are pooled over the RNA chains of a complex.

| Measure | ρ with the share of |  |  |  | Median, complexes mostly |  |
|---|---|---|---|---|---|---|
|  | Correct interface | Correct site, incorrect RNA | Distal site | Incorrect binding site | correct | correct site, incorrect RNA |
| WC pair fidelity (INF) | +0.26 | -0.24 | -0.12 | -0.07 | 1.00 | 1.00 |
| Non-WC pair fidelity (INF) | +0.51<sup>*</sup> | -0.50<sup>*</sup> | -0.13 | +0.00 | 0.92 | 0.58 |
| Stacking fidelity (INF) | +0.49<sup>*</sup> | -0.36<sup>*</sup> | -0.15 | -0.23<sup>*</sup> | 0.53 | 0.12 |
| Target: *syn* share | -0.21<sup>*</sup> | +0.27<sup>*</sup> | +0.05 | +0.08 | 0.06 | 0.14 |
| Target: non-C3′-*endo* share | -0.23<sup>*</sup> | +0.25<sup>*</sup> | +0.05 | +0.06 | 0.21 | 0.50 |
| Target: non-g<sup>+</sup> γ share | -0.23<sup>*</sup> | +0.29<sup>*</sup> | -0.01 | +0.04 | 0.25 | 0.42 |
| AF3 − target: *syn* | +0.32<sup>*</sup> | -0.33<sup>*</sup> | -0.11 | -0.15 | 0.00 | -0.08 |
| AF3 − target: non-C3′-*endo* | +0.33<sup>*</sup> | -0.25<sup>*</sup> | -0.13 | -0.17<sup>*</sup> | 0.00 | -0.13 |
| AF3 − target: non-g<sup>+</sup> γ | +0.19<sup>*</sup> | -0.22<sup>*</sup> | +0.04 | -0.07 | -0.11 | -0.20 |

#### Which descriptors predict the failure classes

To weigh these associations against each other, we trained one gradient-boosting classifier per class (correct interface; correct site with an incorrect RNA; distal or shifted site; incorrect binding site; one against the rest) on six groups of descriptors. These are AF3’s confidence, experimental and environmental conditions (method, resolution, pH, temperature, ionic strength, PEG, buffer, Mg^2+^ and monovalent salt of the deposited solution, ions and ligands in the deposited structure), RNA sequence and structure of the target (length, GC content, sequence complexity, folding free energy and ensemble diversity, and the target’s shares of syn nucleotides, non-C3*^′^*-*endo* puckers and non-g^+^*γ* angles), evolutionary information (RNA and protein alignment depth, Rfam family, sequence identity and fold similarity to the closest pre-cutoff RNA), templates and composition (chain numbers, protein length, identical copies). Nothing that defines the classes and no accuracy of the AF3 model was used. Out-of-fold predictions came from cross-validation grouped by PDB entry (5 folds, 3 repeats), each complex weighted equally. The out-of-fold AUC above 0.5 was split among the groups by exact Shapley values, and features were ranked by the AUC drop when shuffled within the held-out fold (regression_analysis/interface). After the cutoff the classes are predicted with AUC 0.82-0.90. AF3’s confidence contributes most (0.08-0.12 of the AUC), followed by RNA sequence and structure (0.06-0.07), evolutionary information (0.05-0.08) and composition (0.05-0.07), whereas experimental conditions (0.03-0.06) and templates (0.00-0.02) add little. Correct interfaces are best predicted by a high RNA pLDDT (AUC drop 0.062), an RNA fold similar to a pre-cutoff fold (0.039) and a high protein pLDDT (0.016). Correct sites with an incorrect RNA correlate with RNA that is unlike pre-cutoff structures (fold similarity 0.033, sequence identity 0.007), with a lower protein pLDDT (0.029), and with targets rich in non-C3*^′^*-*endo* puckers and with a high ensemble diversity (0.008, 0.010). Distal and shifted sites correlate with a higher chain-pair PAE (0.038), a lower ipTM*_r_* (0.036), the target’s share of non-g^+^ *γ* angles (0.021), a lower sequence complexity (0.013) and a less negative folding free energy (0.010), whereas protein size and the total and longest RNA length contribute almost nothing. Incorrect binding sites correlate with a lower protein pTM (0.031) and RNA pLDDT (0.022), a high ensemble diversity (0.017) and a lower ipTM*_r_* (0.012), whereas an unfamiliar RNA fold (0.009), a lower protein pLDDT (0.007) and a higher global PAE (0.006) contribute less. No single experimental descriptor (pH, PEG, ions, buffer) exceeds an AUC drop of 0.008. Before the cutoff AF3’s confidence dominates every class (0.12-0.14). Correlated descriptors share their importance, so single values are lower bounds and describe associations.

#### Placement per RNA chain and by interface side

Each placement class mixes several kinds of error. Table S4 therefore lists, for the failed post-cutoff interfaces of each class, how often AF3 contacts the native site, keeps the pocket in place, leaves the whole RNA within 10 Å or moves it beyond 20 Å, turns it by more than 45*^◦^*, folds the RNA well and assembles the protein correctly. It also lists how often the protein and the RNA side of the interface are recovered (contact match *≥* 0.5). At a correct site with an incorrect RNA, only 4% of the RNAs are well folded, and about a third of these interfaces recover the native nucleotides while another third recover none. Distal, shifted and incorrect binding sites never contact the native protein residues. Apart from the incorrect binding sites, a mispredicted pocket occurs only at correct sites with an incorrect RNA (29%). Table S5 summarises the same classes per RNA chain, each RNA chain being described by its main protein partner. Correct binding sites with an incorrect RNA combine a misfolded RNA (84%) with a main protein partner that is folded correctly overall but less accurately in local detail (median TM*_p_* 0.89, LDDT*_p_* 0.79), and only 26% of these partners meet the accuracy threshold. Incorrect binding sites mostly have an inaccurate main protein partner (86%, median TM*_p_* 0.79), and shifted RNAs mostly sit on an accurate one (78%). Where AF3 makes no contact with the main protein partner, the distance separates the classes. Rotated RNAs lie within 10 Å, shifted RNAs 10–20 Å and distal RNAs more than 20 Å away. Table S6 crosses the protein and the RNA side of the failed post-cutoff interfaces. For RNAs longer than 15 nt, 51 of the 580 failed interfaces recover both sides but miss the geometry, 126 contact the native protein site with a different stretch of the RNA, 60 bind the native nucleotides with another protein surface and 172 recover neither side. Short RNAs reach a recovered RNA side far more easily (71 of 206 without the protein site), so the RNA side of short RNAs says little about the interface.

**Table S4.** Attributes of the failed post-cutoff protein–RNA interfaces in each placement class (% of the class). Site contact means that AF3 contacts at least 10% of the native protein contact residues (protein contact match *≥* 0.1). Pocket intact means that these residues superpose on the experimental ones within 4 Å (pocket RMSD). Whole RNA gives the displacement of the whole RNA from its native position after superposition of the protein, and Turned gives its rotation. RNA well folded is based on LDDT*_r_ >* 0.85 (and TM*_r_ >* 0.5 from 60 nt). Assembly correct means that the protein is placed correctly against its protein partners (DockQ*_pp_ ≥* 0.23, or TM*_p_ ≥* 0.5 for a protein without protein partners). Protein side correct and RNA side correct mean that AF3 contacts at least half of the native contact residues or nucleotides (contact match *≥* 0.5). RNA side none means that it contacts almost none of the native nucleotides (RNA contact match *<* 0.1).

| Placement class | *n* | Site contact | Pocket intact | Whole RNA ≤ 10 Å | Whole RNA > 20 Å | Turned > 45° | RNA well folded | Assembly correct | Protein side correct | RNA side correct | RNA side none |
|---|---|---|---|---|---|---|---|---|---|---|---|
| Correct site, slid duplex | 8 | 0 | 100 | 75 | 0 | 88 | 25 | 100 | 0 | 88 | 12 |
| Correct site, incorrect RNA | 422 | 99 | 71 | 29 | 46 | 74 | 4 | 56 | 46 | 36 | 34 |
| Correct site, rotated RNA | 21 | 0 | 100 | 100 | 0 | 100 | 19 | 29 | 0 | 10 | 90 |
| Shifted site | 48 | 0 | 100 | 0 | 0 | 79 | 12 | 69 | 0 | 23 | 71 |
| Distal site | 181 | 0 | 100 | 0 | 100 | 91 | 10 | 50 | 0 | 9 | 69 |
| Incorrect binding site | 106 | 0 | 0 | 6 | 73 | 91 | 11 | 43 | 0 | 15 | 52 |
| All failed interfaces | 786 | 53 | 71 | 20 | 57 | 81 | 8 | 53 | 25 | 26 | 48 |

**Table S5.** Placement of the post-cutoff RNA chains, each described by its main protein partner, the protein with the largest native interface (668 RNA chains). RNA good, LDDT*_r_ >* 0.85 (and TM*_r_ >* 0.5 from 60 nt), and RNA wrong, LDDT*_r_ ≤* 0.75. Partner accurate, LDDT*_p_* and TM*_p_* of the main partner *>* 0.85. Protein and RNA side, mean protein and RNA contact match of the main-partner interface, the RNA side separately for RNAs of more and of at most 15 nucleotides. No contact, RNA chains with both contact matches *<* 0.1.

| Placement (main partner) | *n* | RNA good | RNA wrong | Partner accurate | Protein side | RNA side | RNA side | No contact |
|---|---|---|---|---|---|---|---|---|
|  |  | (%) | (%) | (%) |  | (> 15 nt) | (≤ 15 nt) |  |
| Correct interface | 329 | 40 | 36 | 64 | 0.66 | 0.80 | 0.88 | 0 |
| Correct site, slid duplex | 4 | 50 | 25 | 50 | 0.03 | 0.55 | 0.95 | 0 |
| Correct site, incorrect RNA | 194 | 4 | 84 | 26 | 0.44 | 0.49 | 0.65 | 0 |
| Correct site, rotated RNA | 5 | 60 | 20 | 100 | 0.00 | 0.00 | 0.12 | 4 |
| Shifted site | 23 | 26 | 30 | 78 | 0.03 | 0.04 | 0.51 | 13 |
| Distal site | 70 | 16 | 69 | 54 | 0.01 | 0.16 | 0.38 | 27 |
| Incorrect binding site | 43 | 23 | 63 | 14 | 0.03 | 0.29 | 0.31 | 12 |

#### Protein fold and assembly placement by class

Table S8 lists, for each placement class of the post- and pre-cutoff interfaces, the fold class of the complex, the per-chain accuracy of the interface’s protein and RNA chains and the placement of the protein chain against its protein partners. Each predicted chain is scored against the reference chain assigned to it one-to-one by TM-score among chains of the same sequence. Except at incorrect binding sites, the protein chain of a failed interface is folded as accurately as at a correct interface (median TM*_p_* 0.93-0.97 against 0.96, LDDT*_p_* 0.81-0.93 against 0.87), so most failures misplace a correctly folded protein rather than misfold it. The misplacement often extends to the protein assembly. The protein chain has an incorrect protein–protein interface (DockQ*_pp_ <* 0.23) at 58% of distal sites and 54% of correct sites with an incorrect RNA, against 21% of correct interfaces. Rotated RNAs occur often in otherwise accurate complexes (50% class i after the cutoff, protein LDDT 0.93), all with two or more RNA chains. Most are duplex strands placed at the native site but turned. Slid duplexes occur only in class-i complexes, before and after the cutoff, and after the cutoff none has a misplaced protein partner. The pre-cutoff interfaces show the same pattern. Their protein chains are folded at a median TM*_p_* of 0.93-0.98 in every class except incorrect binding sites (0.78), and 71% of distal sites have an incorrect protein–protein interface, against 33% of correct interfaces.

**Table S6.** Protein side against RNA side of the failed post-cutoff protein–RNA interfaces, for RNAs of more and of at most 15 nucleotides. Correct, contact match *≥* 0.5. Partial, 0.1–0.5. None, *<* 0.1.

|  | RNA > 15 nt (*n* = 580) |  |  | RNA ≤ 15 nt (*n* = 206) |  |  |
|---|---|---|---|---|---|---|
| Protein side | RNA correct | partial | none | RNA correct | partial | none |
| Correct | 51 | 49 | 57 | 26 | 2 | 11 |
| Partial | 58 | 64 | 69 | 19 | 9 | 4 |
| None | 13 | 47 | 172 | 40 | 31 | 64 |

**Table S7.** Composition of the correctly placed RNA chains (DockQ*_pr_ ≥* 0.23 with the main partner) by the fold of the RNA and of the main partner, with the median DockQ*_pr_* of each group. RNA good, LDDT*_r_ >* 0.85 (and TM*_r_ >* 0.5 from 60 nt). Intermediate, 0.75–0.85. Poor, *≤* 0.75. Partner accurate, LDDT*_p_* and TM*_p_ >* 0.85 (publication_figures/placement_per_rna_chain.py).

|  | Post-cutoff (*n* = 329) |  | Pre-cutoff (*n* = 1,365) |  |
|---|---|---|---|---|
| RNA fold | Partner accurate | inaccurate | Partner accurate | inaccurate |
| Good | 95 (0.82) | 38 (0.72) | 844 (0.89) | 162 (0.71) |
| Intermediate | 39 (0.65) | 37 (0.60) | 108 (0.82) | 54 (0.60) |
| Poor | 75 (0.56) | 45 (0.43) | 142 (0.49) | 55 (0.46) |

**Table S8.** Composition of the interface placement classes (post- and pre-cutoff protein–RNA interfaces). Fold class of the complex (i, RNA and protein accurate; ii, RNA accurate; iii, protein accurate; iv, neither), median TM*_p_* and LDDT*_p_* of the interface’s protein chain and LDDT*_r_* of its RNA chain, share of interfaces whose protein chain is misplaced against its protein partners (lowest DockQ*_pp_ <* 0.23, complexes with several protein chains), and share with two or more RNA chains and with identical protein copies.

|  |  | Fold class (%) |  |  |  | Protein chain |  | RNA chain | Partner PP | ≥ 2 RNA | Identical |
|---|---|---|---|---|---|---|---|---|---|---|---|
| Placement | *n* | i | ii | iii | iv | TM<sub>*p*</sub> | LDDT<sub>*p*</sub> | LDDT<sub>*r*</sub> | wrong (%) | chains (%) | proteins (%) |
| *Post-cutoff* |  |  |  |  |  |  |  |  |  |  |  |
| Correct interface | 681 | 60 | 12 | 23 | 6 | 0.96 | 0.87 | 0.82 | 21 | 72 | 41 |
| Correct site, slid duplex | 8 | 100 | 0 | 0 | 0 | 0.97 | 0.87 | 0.74 | 0 | 100 | 0 |
| Correct site, incorrect RNA | 422 | 8 | 6 | 40 | 46 | 0.93 | 0.81 | 0.53 | 54 | 58 | 57 |
| Correct site, rotated RNA | 21 | 50 | 0 | 33 | 17 | 0.97 | 0.93 | 0.79 | 79 | 100 | 81 |
| Shifted site | 48 | 35 | 6 | 42 | 17 | 0.95 | 0.87 | 0.77 | 34 | 85 | 77 |
| Distal site | 181 | 18 | 7 | 60 | 15 | 0.95 | 0.86 | 0.61 | 58 | 85 | 60 |
| Incorrect binding site | 106 | 11 | 23 | 16 | 50 | 0.69 | 0.75 | 0.62 | 66 | 86 | 34 |
| *Pre-cutoff* |  |  |  |  |  |  |  |  |  |  |  |
| Correct interface | 2848 | 68 | 17 | 9 | 5 | 0.98 | 0.90 | 0.91 | 33 | 69 | 33 |
| Correct site, slid duplex | 10 | 100 | 0 | 0 | 0 | 0.95 | 0.90 | 0.71 | 22 | 100 | 20 |
| Correct site, incorrect RNA | 340 | 37 | 12 | 39 | 11 | 0.97 | 0.86 | 0.75 | 56 | 64 | 77 |
| Correct site, rotated RNA | 6 | 100 | 0 | 0 | 0 | 0.93 | 0.89 | 0.82 | 50 | 67 | 33 |
| Shifted site | 50 | 66 | 8 | 24 | 2 | 0.98 | 0.92 | 0.69 | 41 | 94 | 64 |
| Distal site | 368 | 53 | 5 | 39 | 4 | 0.98 | 0.92 | 0.74 | 71 | 87 | 66 |
| Incorrect binding site | 72 | 29 | 39 | 8 | 24 | 0.78 | 0.78 | 0.79 | 78 | 71 | 67 |

#### RNA chains with several protein partners

Half of the post-cutoff RNA chains (356 of 668, with 1,155 interfaces) contact two or more proteins in the deposited structure, and 688 of 1,646 before the cutoff. Their interfaces mostly share one outcome. After the cutoff, all interfaces of 156 of these RNA chains are correct, those of 144 all fail, and only 56 (16%) mix correct and failed interfaces (29% before the cutoff). All interfaces fall into one placement class for 60% of them. The other partners have smaller native interfaces than the main partner (median 7 against 22 protein residues) and are correct less often (44% against 55%). Their outcome follows the main partner. They are correct for 79% of the RNA chains placed correctly on the main partner but for only 6% of those misplaced on it, so a misplaced RNA is misplaced against all its proteins. Where another partner fails next to a correct main interface (85 interfaces), the protein itself is usually misplaced in the assembly (61%, against 14% of the other partners with a correct interface), as for 8YE6. The failure modes carry over between partners (Table S9). An RNA with an incorrect pose at the main partner’s site mostly has one at the other partners’ sites too, and a distal RNA is mostly distal from them as well. Before the cutoff the same holds (86% and 14%, and 77% of the failing other partners misplaced in the assembly; publication_figures/placement_multi_partner.py).

**Table S9.** Placement class of the other protein partners of post-cutoff RNA chains with two or more partners (799 interfaces of 356 RNA chains), against the class of the main partner, the protein with the largest native interface. Rotated or slid, correct site with a rotated RNA or a slid duplex.

|  | Other partner |  |  |  |  |  |
|---|---|---|---|---|---|---|
| Main partner | Correct | Correct site, incorrect RNA | Distal | Incorrect binding site | Shifted | Rotated or slid |
| Correct | 327 | 53 | 21 | 8 | 0 | 3 |
| Correct site, incorrect RNA | 18 | 127 | 22 | 28 | 13 | 14 |
| Distal | 7 | 23 | 53 | 9 | 4 | 0 |
| Incorrect binding site | 0 | 13 | 15 | 18 | 1 | 2 |
| Shifted | 0 | 11 | 0 | 0 | 7 | 0 |
| Rotated or slid | 0 | 1 | 0 | 0 | 0 | 1 |

## O. Protein domains of predicted interfaces

This appendix gives the per-domain analysis of binding affinity, interface-residue localisation and interface accuracy summarised in Section 3.3. RNA-binding domains were annotated with InterProScan 5.78-109.0 [90], run through the EMBL-EBI web service on the deposited sequence of each protein chain, with modified residues read as their parent amino acid (database/annotateRBDs.py). The rules follow the rbd-annotation pipeline (https://github.com/michauhl/rbd-annotation), adapted with its author’s permission. A hit is kept when its InterPro entry is one of the 1,000 RNA-binding domain entries of RBP2GO [91] and its extension, and it is assigned to the InterPro parent of that entry. Hits from member databases that report an E-value are kept at E *≤* 10*^−^*^5^. A hit contained in another hit of the same entry is dropped, and hits that overlap by at least 60% of the shorter one are merged. Copies of a domain in several chains of a complex are pooled and count once. A domain counts once per occurrence and a repeat array once. The same annotation covers the pre-cutoff, post-cutoff and 2026 protein–RNA sets and the RNAcompete proteins.

The number of annotated protein domains was associated with the predicted binding affinity (Fig. S29a). Complexes with more domains trended towards stronger (more negative) log_10_ *K_d_* (Spearman *ρ* = *−*0.35, *P* = 3 *×* 10*^−^*^5^, *n* = 138; median *−*51 for one- or two-domain proteins versus *−*97 for proteins with *≥* 3 domains), which is consistent with multi-domain RNA-binding proteins engaging RNA through larger interfaces. This architectural dependence did not, however, carry over to prediction quality. Domain multiplicity was at most weakly and negatively related to interface DockQ*_Nr_* (per-domain copy number *ρ* = *−*0.16, *P* = 0.03; total domain copies *ρ* = *−*0.19, *P* = 0.07), so accuracy declines slightly as domains accumulate, but the trend is weak, only nominally significant for the per-domain copy number and accounts for roughly 3% of the variance. Domain count therefore tracks annotated affinity far more than it does how accurately AF3 models the interface.

Mapping predicted RNA-contacting residues onto annotated domains (240 domain occurrences across 37 domains in 73 complexes with a single distinct protein, comprising 36 electron-microscopy and 37 X-ray structures, Fig. S29b) showed that the model rarely localised interactions precisely within domain boundaries. Only 17.5% (42) of occurrences fell entirely within the annotated domain, the majority were only partially captured (67.9%, 163), and a small fraction lay outside (14.6%, 35). AF3 therefore identifies the correct interaction region but systematically mis-places domain boundaries. The pattern was domain-dependent. To illustrate, a few compact folds were fully resolved (La_HTH, 7/7 within; CSD, 8/10), most were partial-dominated at 80-100% (e.g. Helicase_C-like, RLR_CTR, RIG-I-like_C, RNase_III, Helicase_ATP-bd, PAZ), and for a few domains the predicted contacts often fell outside the domain (Argonaute Mid, 4 of 5; pseudouridine synthase, 44%).

Interface accuracy itself was strongly domain-dependent (Fig. S29c). Since DockQ*_Nr_* scores only the protein–RNA contacts, its values run stricter than whole-complex DockQ. A handful of well-sampled domains reached medium interface quality (Pumilio PUM-HD DockQ*_Nr_* 0.65, *n* = 4; RLR_CTR 0.65, *n* = 5; Piwi 0.55, *n* = 10; La_HTH 0.57, *n* = 7), most sat in the low-acceptable range (0.27-0.36, PsdUridine_synth, PAZ, Helicase_C-like, Helicase_ATP-bd, CSD, WD40, RRM), and several were effectively unmodelled at the interface (dsRBD 0.19, *n* = 5; RNase_III and Metallo-*β*-lactamase *≈* 0.16, *n* = 4-7; Dicer dimerisation 0.02, *n* = 4). This suggests that compact single-domain RNA-binding folds are easier interface targets than the large, repeat-containing endonuclease architectures. Thus, while the *number* of domains a protein carries tracks binding affinity but not interface accuracy, it is the domain *identity* that governs how well AF3 places the RNA, with canonical RNA-processing folds such as the Dicer dimerisation domain placed little better than chance (DockQ*_Nr_ ≈* 0.02). DockQ*_Nr_* scores only the interface, so unlike global complex RMSD it is unaffected by assembly size or y how well the whole structure superimposes.

## P. RNAcompete binding-specificity benchmark

This appendix provides the full RNAcompete analysis summarised in Section 3.3.

244 RNA-binding proteins including their 3 most binding (maxprobe) and 3 least binding (minprobe) RNA sequences from RNAcompete experiment were assessed to determine whether AF3-based predictions capture experimentally measured RNA binding preferences. The RNA motif score achieved the strongest discrimination between experimentally strongest- and weakest-binding probes, with a pooled ROC-AUC of 0.645 and a moderate effect size (Cohen’s *d* = 0.48; Mann-Whitney *P* = 9.2 *×* 10*^−^*^22^). Maxprobes displayed significantly higher RNA motif scores than minprobes (Wilcoxon signed-rank test, *n* = 244, *P* = 4.7 *×* 10*^−^*^12^), with 68% of proteins exhibiting higher motif scores in their strongest-binding probes. In contrast, interface-based measures such as protein–RNA interface area and interface ratio exhibited only weak separation (AUC *≈* 0.52), and even protein-length normalisation using 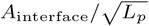 yielded only marginal improvement (AUC = 0.524). Predicted binding affinity performed similarly (mean rank accuracy = 0.546; AUC = 0.529).

**Fig. S29:**
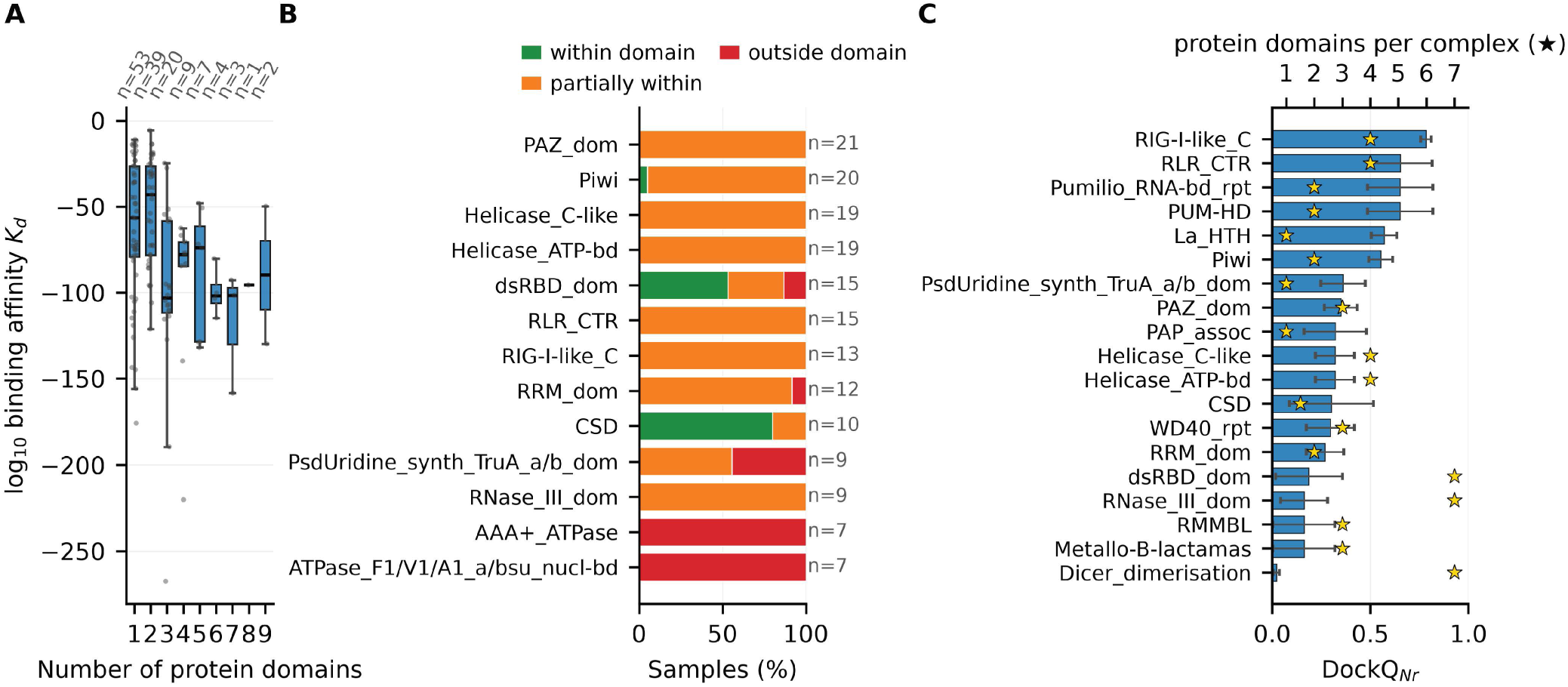
Protein-domain context of AF3 protein–RNA predictions. **A**, Binding affinity (log_10_ *K_d_*) by the number of annotated protein domains (*n* = 138 complexes). **B**, For the best-sampled domains, the share of protein–RNA interfaces (complexes with a single distinct protein) whose predicted interface residues lie within (green), partly within (orange) or outside (red) the domain. Most predicted interfaces overlap the annotated domain only partly. **c**, DockQ*_Nr_* per domain (mean *±* s.e.m.), with the median number of domains per complex (stars). The least accurately docked domains, such as the Dicer dimerisation and RNase III domains, sit in large multi-domain proteins. Several estimates rest on few complexes.

Among AF3-derived features, RNA interface confidence (ipTM*_r_*) provided the strongest structural predictor of binding preference, achieving a mean within-protein rank accuracy of 0.565 (median = 0.556) and a pooled AUC of 0.539. Logistic regression analyses further identified ipTM*_r_* as the dominant positive predictor of predicted affinity discrimination (standardised coefficient = +1.39). The minimum chain-pair PAE, which is a metric reported to correlate with whether or not two chains interact [92], behaved comparably. It only marginally exceeded ipTM*_r_* overall (pooled AUC = 0.547 versus 0.539; identical mean within-protein rank accuracy = 0.565) and separated maxprobe from minprobe only better at low false-positive rates (Fig. S30b). This distinction is subtle, however, because both maxprobe and minprobe form a complex with the protein. We test the ranking of relative binding preference (strong versus weak binder) rather than the interacting-versus-non-interacting discrimination for which the chain-pair PAE was originally proposed. Despite significant enrichment of stronger predicted affinities in maxprobes (Wilcoxon *P* = 1.9 *×* 10*^−^*^4^), neither ipTM*_r_* nor predicted binding affinity correlated with experimental binding intensity differences across proteins (Spearman *ρ* = 0.044, *P* = 0.49 and *ρ* = *−*0.005, *P* = 0.94, respectively). This indicates that AF3-derived metrics capture relative binding preference but fail to reproduce the quantitative experimental binding landscape. Restricting the analysis to proteins with identical RNA lengths (*n* = 12) substantially improved the discrimination by AF3’s confidence, increasing ipTM*_r_* AUC from 0.539 to 0.683 and mean rank accuracy from 0.565 to 0.694. AF3 pTM*_r_*, pLDDT*_r_* and ranking score, on the other hand, were only modestly improved in maxprobes. Protein confidence metrics, including pTM*_p_*, pLDDT*_p_* and disorder fraction, were essentially non-informative.

To investigate the impact of RNA-binding domains, we quantified the RNA motif overlap with InterPro domain annotations across all protein–RNA complexes. Most RNA motifs exhibited at least partial overlap with annotated RNA-binding domains, with 91.8% of samples classified as having residues within a single domain and only 2.2% lying entirely outside annotated domains. Canonical RNA-binding families showed strong enrichment, including RRM domains (98.2% partial or complete overlap), KH domains (85.0%), and Pumilio repeats (100%). Certain domain families displayed especially precise localisation, including S1 domains (50%), SAM domains (33.3%) and Pumilio repeats (25.0%), whereas PUM-HD domains overlapped mostly in part (5.6% complete overlap). However, regarding the maxprobe vs minprobe separation RNA motif-based methods consistently outperformed AF3 ipTM*_r_* across motifs localised within, partially within, or outside annotated domains. For example, RRM and KH proteins exhibited moderate separation by AF3 ipTM*_r_* (mean AUCs of 0.564 and 0.545, respectively), while the separation by RNA motif scores significantly outperformed those by ipTM*_r_* within RRMs (paired Wilcoxon *P* = 6.6 *×* 10*^−^*^3^). In contrast, PUM-HD proteins represented a notable exception, achieving exceptionally high separation by ipTM*_r_* (mean ipTM*_r_* AUC = 0.87), comparable to motif-based methods. This suggests that AF3 ipTM*_r_* can be used for maxprobe vs minprobe separation in modular, sequence-specific recognition systems such as Pumilio repeat proteins, whereas motif scores computed from the probe sequence remain the dominant separator in canonical RNA-binding domains such as RRMs and KH domains.

**Fig. S30:**
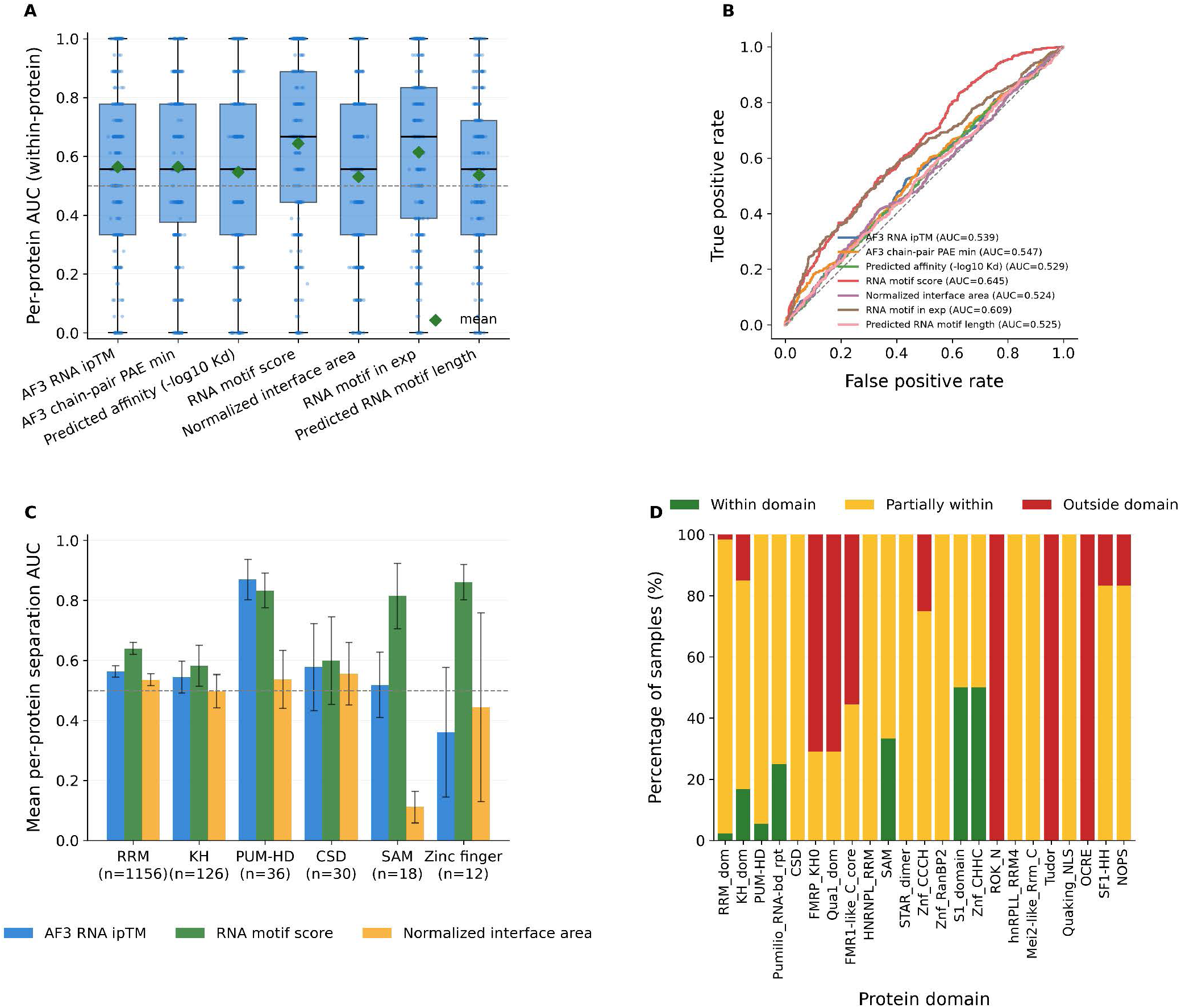
AF3 interface metrics and sequence features for RNA-binding specificity on RNAcompete. **A**, Within-protein ranking AUC of seven scores across 244 RBPs (boxes, interquartile range and median; diamonds, mean; dashed line, chance). **B**, Pooled ROC curves for maxprobe against minprobe sequences. The minimum chain-pair PAE (AUC 0.547) marginally exceeds ipTM*_r_* (0.539). **C**, Family-separation AUC of ipTM*_r_*, the RNA motif score and the normalised interface area by RNA-binding domain family (families with at least five samples, without proteins of several or no assigned families; *n*, number of samples; bars, mean *±* s.e.m. over proteins). **D**, Share of samples whose top-ranked motif lies within, partly within or outside the annotated RNA-binding domain.

### P.1. Competition between strong- and weak-binding probes

To evaluate whether structure-based predictions can identify experimentally preferred RNA targets in a competitive setting, we analysed 728 RNAcompete-derived complexes in which each protein was predicted together with one experimentally strongest-binding probe (maxprobe) and one to three weakest-binding probes (minprobes). If the model successfully captured RNA-binding specificity, the maxprobe would be expected to preferentially form the strongest interaction. Across all complexes, the maxprobe interacted with the protein in 83.5% of cases (608/728), compared with interaction frequencies of 78.3%, 69.9%, and 65.0% for the first, second, and third minprobe, respectively. However, interactions were frequently not exclusive. Both maxprobe and at least one minprobe bound in 560 of 728 complexes (76.9%), whereas only 48 complexes (6.6%) exhibited maxprobe-only binding and 120 (16.5%) showed binding exclusively by one or more minprobes. The tendency for minprobes to remain bound demonstrates that RNAcompete minprobes are generally lower-affinity binders rather than true non-binders due to a continuous affinity landscape rather than a binary binding/non-binding distinction.

Within-complex ranking analyses similarly indicated only modest discrimination of the experimentally preferred probe. RNA motif score was the strongest predictor, ranking the maxprobe first in 55.1% of complexes and achieving a competition ROC-AUC of 0.697. In contrast, ipTM*_r_* ranked the maxprobe first in only 47.2% of complexes (AUC = 0.619), while interface contact counts performed comparably (AUC = 0.597). The per-chain minimum chain-pair PAE, evaluated here per RNA chain from the full chain-pair PAE matrix, performed on par with ipTM*_r_*, ranking the maxprobe first in 42.9% of complexes (AUC = 0.605). The two AF3 confidence signals thus carry similar, and similarly limited, within-complex discriminative information. Mean score differences between maxprobes and competing minprobes were small for ipTM*_r_* (Δ*S* = *−*0.003) but consistently positive for RNA motif score (Δ*S* = 0.067). Length-controlled analyses produced similar conclusions. Restricting the dataset to complexes in which all competing probes had identical RNA lengths increased maxprobe interaction frequency to 88.2% and modestly improved ranking performance (ipTM*_r_* AUC = 0.623; RNA motif score AUC = 0.730).

These observations were largely conserved across RNA-binding protein domains. For the dominant RRM family (*n* = 578 complexes), RNA motif score substantially outperformed ipTM*_r_* (AUCs of 0.699 versus 0.618), with a similar pattern observed for KH domains (0.713 versus 0.609). PUM-HD proteins represented a notable exception, where maxprobe separation based on ipTM*_r_* was strongest (ipTM*_r_* AUC = 0.713) and approached motif-based accuracy (AUC = 0.771), consistent with the highly structured and modular mode of Pumilio-mediated RNA recognition. In contrast, ipTM*_r_* discriminated poorly for CSD and zinc-finger proteins (AUC 0.53 and 0.43, 15 and 6 complexes).

**Fig. S31:**
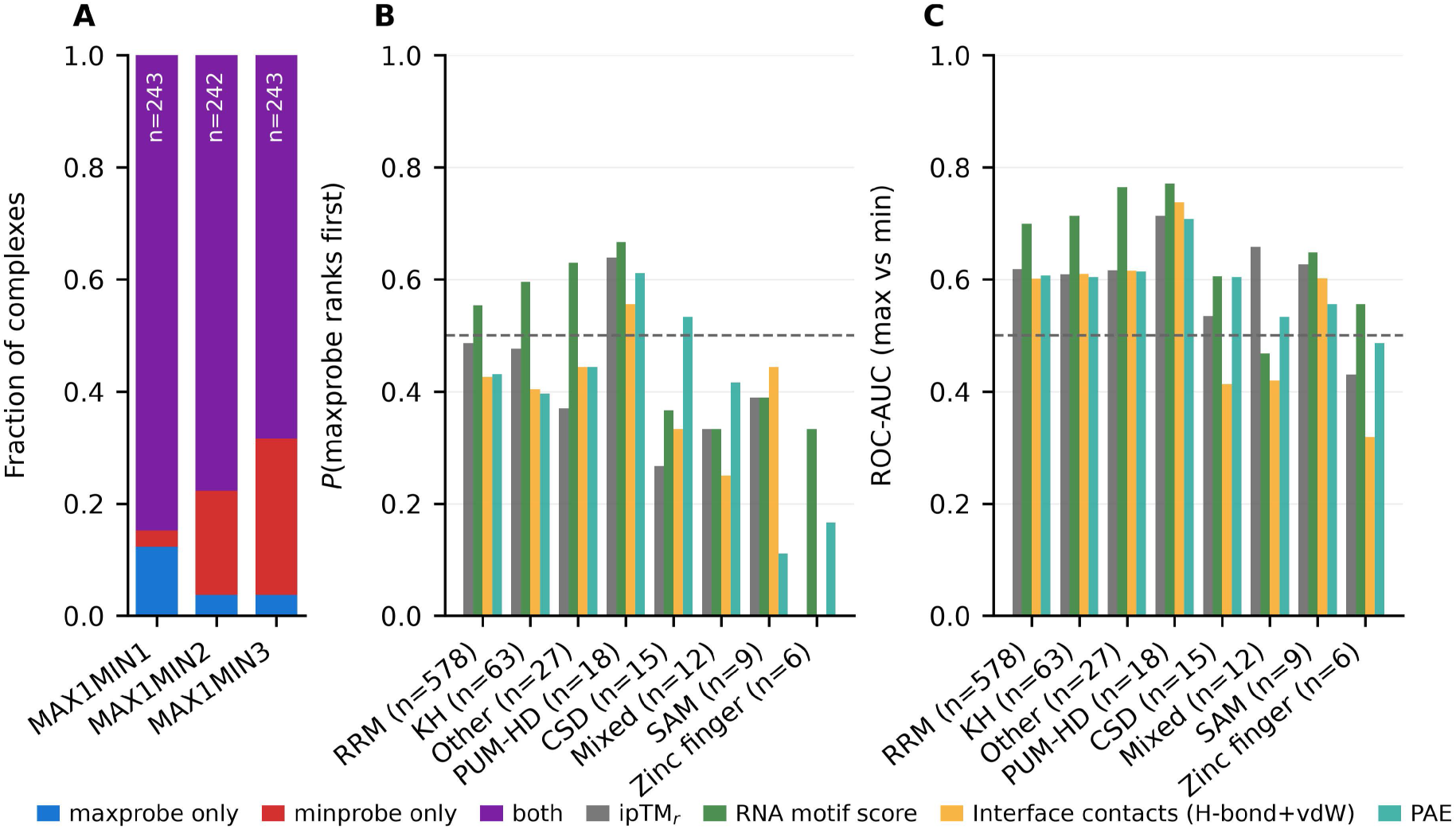
Predicted binding selectivity in RNAcompete competition complexes. **A**, Share of competition complexes in which AF3 binds the maxprobe only (blue), the minprobe only (red), both (purple) or neither (grey), per competition design. **B**, Probability that the maxprobe ranks first by ipTM*_r_*, RNA motif score, interface contact count and minimum chain-pair PAE, per RNA-binding domain family (at least five complexes). **C**, ROC-AUC for maxprobe against minprobe chains with the same metrics and families (dashed lines, chance).

### P.2. Positional test of the RNA motif

The positional test of Appendix B.3 follows the experimental RNA motif into the AF3 model (Fig. S32). AF3 contacts these positions more often than its overall contact coverage predicts, and its best-scoring RNA motif lies on the native site in 83% of pre-cutoff and 69% of post-cutoff entries.

**Fig. S32:**
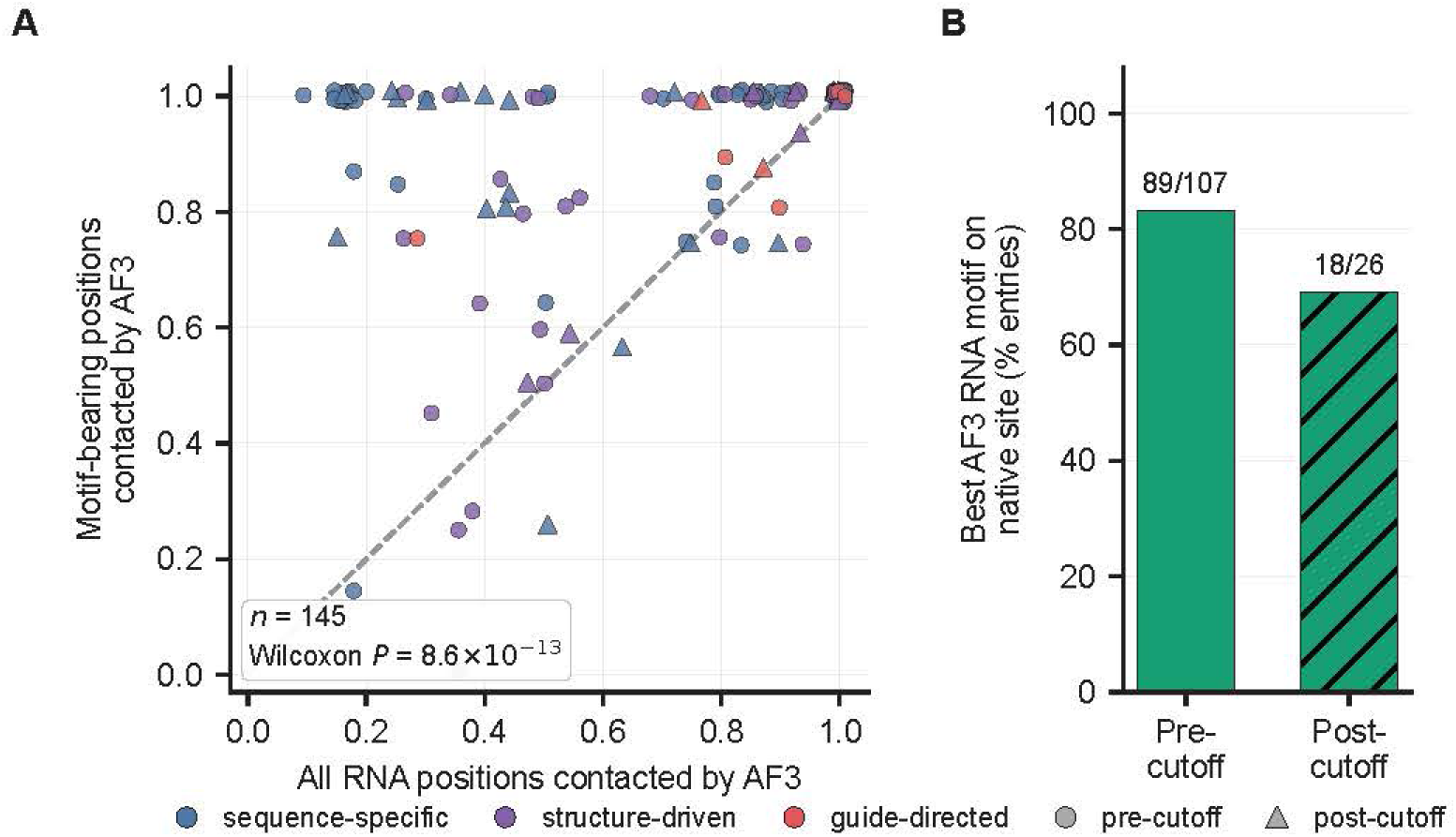
Positional test of the RNA motif. **A**, Per entry, the share of the experimentally motif-bearing positions that AF3 contacts against the share of all RNA positions it contacts (dashed line, chance level; Wilcoxon signed-rank test). Colours mark the RBP recognition class, circles pre-cutoff and triangles post-cutoff entries. **B**, Share of entries whose best-scoring AF3 RNA motif lies on the native site, per cohort (recovered/scored above each bar).

## Q. Important feature determinants of RNA accuracy

To consolidate the determinants above into a single quantitative picture, we ranked every available system descriptor against four RNA outcomes, global fold (TM*_r_*), local all-atom accuracy (LDDT*_r_*), global error (RMSD*_r_*) and the good-versus-bad RNA classification of the decision framework (quadrants i+ii vs. iii+iv, i.e. LDDT*_r_ >* 0.85), using two complementary readouts, the univariate Spearman correlation of each descriptor with each outcome, and its cross-validated random-forest permutation importance (Fig. S33). Descriptors that also exist for the predicted model (interface contacts, secondary-structure counts, backbone geometry, electrostatics) are taken from the *experimental* structure as they describe the target and not the prediction. The three interaction-network-fidelity components (Watson-Crick, non-Watson-Crick and stacking) are the only predicted-versus-experimental quantities retained, shown as a deliberate contrast. The interface agreement scores (AA match score, RNA match score, RNA motif score) are excluded for the same reason, because they compare the predicted with the experimental contact residues and are therefore interface-accuracy outcomes rather than determinants.

Two determinants dominate, and both recur on an independent single-chain RNA benchmark (Fig. S34). First, recovery of non-canonical pairing and base stacking co-varies most strongly with every outcome. Non-WC and stacking INF are among the top random-forest features (*ρ* up to 0.68 on protein–RNA and 0.85 on single-chain RNA) since these INF components are themselves predicted-versus-experimental agreement scores, they characterise, rather than predict, accuracy, whereas Watson-Crick fidelity, although positively correlated, carries essentially no random-forest importance (*≈* 0) as canonical pairing is recovered by default and so does not separate good from bad structures. Second, the target’s backbone conformation is the leading intrinsic determinant. The experimental C2*^′^*-*endo* sugar-pucker fraction *anti*-correlates with local accuracy (*ρ* = *−*0.62 with LDDT*_r_*, the strongest single descriptor) as does the glycosidic-*syn* fraction, while the canonical *g*^+^ *γ*-fraction correlates positively (Fig. S9). Beyond these, templates and structural elaboration contribute secondarily. The protein–RNA complex template count favours local accuracy (*ρ* = +0.26 with LDDT*_r_*, +0.30 with the RNA class), and multibranch, pseudoknot and stem counts favour the global fold (TM*_r_*, *ρ* = +0.32 to +0.38) but not local accuracy. Interface geometry (H-bond and van der Waals counts, buried area) and solution conditions (pH, buffer, ions, PEG) are weak throughout. The full random-forest models reach out-of-fold *R*^2^ *≈* 0.66 (protein–RNA) and *≈* 0.80 (single-chain RNA), driven almost entirely by the two dominant axes above rather than by these weak groups. On the single-chain RNA set the same backbone signal survives (C2*^′^*-*endo ρ ≈ −*0.5 to *−*0.6), but a bundle of co-varying descriptors, MSA*_r_* depth, sequence complexity, entropy, length and secondary-structure counts, turns strongly positive. This is due to the fact that in that set the long, complex RNAs are precisely the deeply conserved, MSA*_r_*-rich families (rRNA-, tRNA- and riboswitch-like), these are best read as a single well-characterised structured-RNA-family axis rather than independent effects (RNA length, which lowers accuracy on protein–RNA, reverses sign here). The single-chain RNA estimates rest on only *n* = 66 complexes (15 classified good) and are correspondingly noisier.

## R. Complex-level and per-class regression analysis

This appendix gives the full regression analysis summarised in Sections 3.5 and 3.5.

**Fig. S33:**
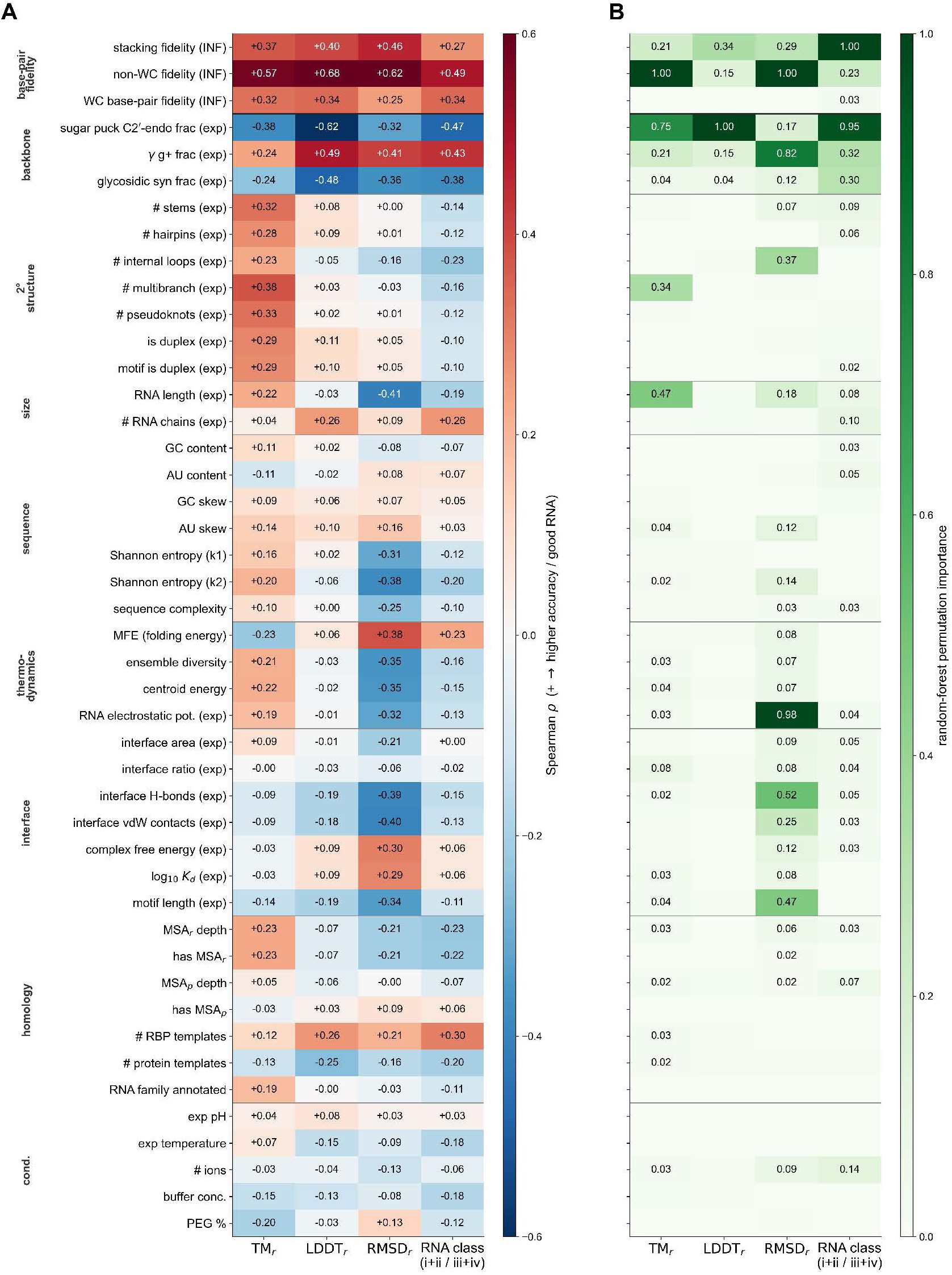
Non-canonical and stacking recovery and the target’s backbone conformation dominate RNA accuracy. Descriptors (rows) against TM*_r_*, LDDT*_r_*, RMSD*_r_* and the well-folded classification (columns) for protein–RNA complexes (*n* = 384). **A**, Spearman *ρ*, oriented so that red denotes higher accuracy. **B**, Cross-validated random-forest permutation importance, normalised per column. The INF components (top block) are the only predicted-versus-experimental quantities and contrast Watson–Crick fidelity, which carries no importance, with non-canonical and stacking fidelity.

**Fig. S34:**
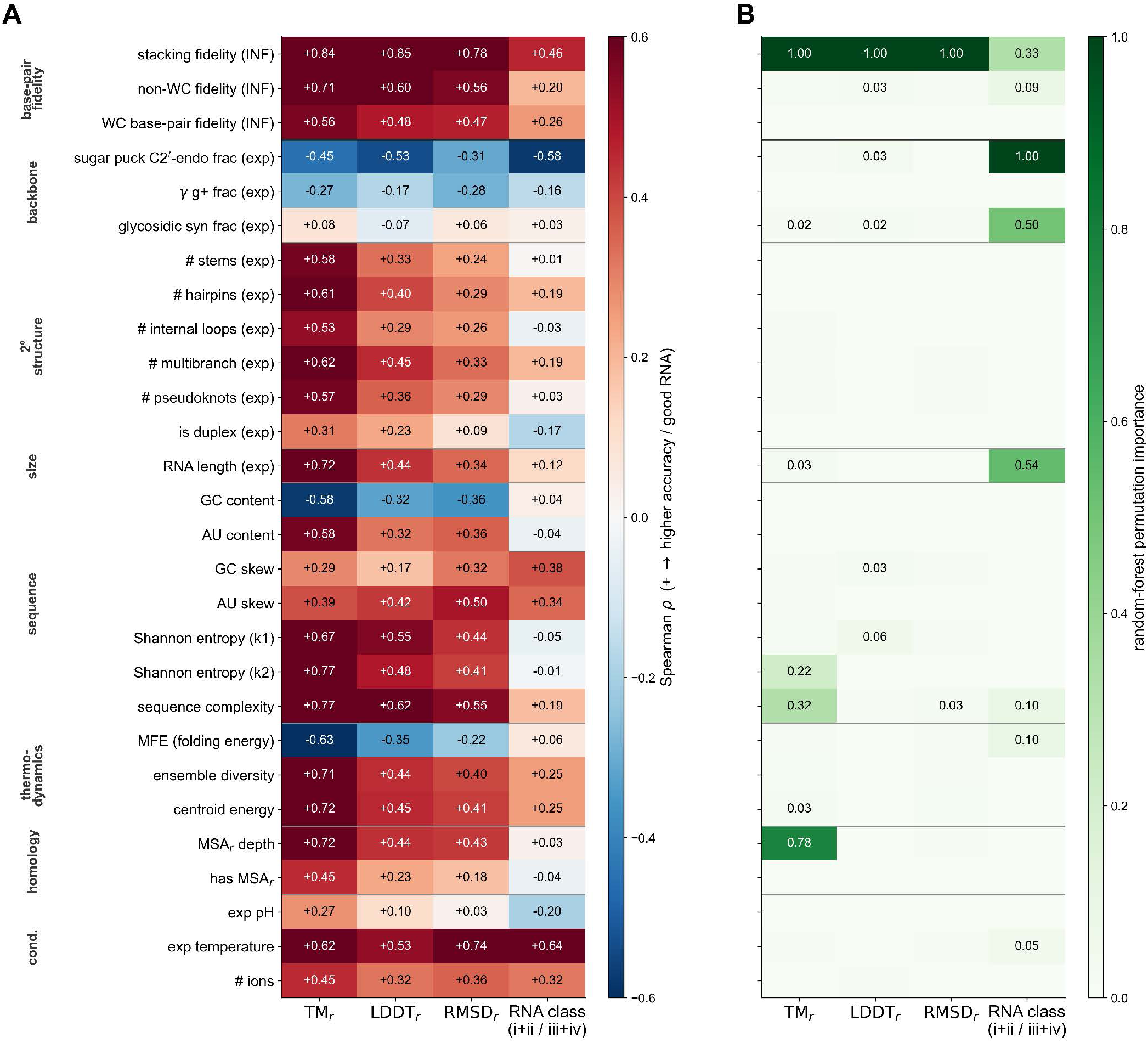
The backbone-conformation determinant of RNA accuracy replicates on single-chain RNA complexes. As Fig. S33, but for the single-chain RNA / RNA–RNA benchmark (*n* = 66, with RNA–RNA interface descriptors and without protein-specific features). The experimental C2*^′^*-*endo* sugar-pucker fraction again *anti*-correlates with accuracy and non-WC/stacking fidelity again lead, while a set of mutually correlated descriptors (MSA depth, sequence complexity, entropy, length, secondary-structure counts) is strongly positive, reflecting the predominance of deeply conserved, well-characterised structured RNA families in this set. Estimates are noisier (*n* = 66, 15 structures classified good).

Shapley decomposition of explained variance.

Attributing incremental *R*^2^ by adding groups in a single fixed order is order-dependent. Because the feature groups are partially correlated, the credit assigned to a group depends on that order. For every coalition *S* of feature groups we therefore evaluated the pooled out-of-fold *R*^2^, *v*(*S*), under the identical pipeline and grouped cross-validation as the main models (with *v*(*∅ø*) given by the per-fold mean predictor), and assigned each group *g* its Shapley value

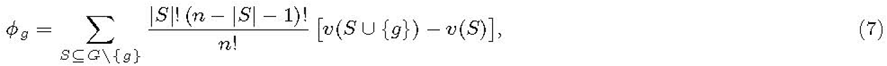

the average marginal contribution of *g* to explained variance over all orderings of the *n* groups. By construction the *ϕ_g_* are order-independent and sum to the full-model explained variance, *v*(*G*) *− v*(*∅ø*), so that each group can be expressed both as an absolute *R*^2^ contribution and as a share of the total (Table S11). For the residual target the partition was computed over the five non-AF3 groups. All analyses used Python with scikit-learn and pandas, a fixed random seed, and configurations written to disk (**variance_partitioning_shapley.py**).

### R.1. Complex level

Besides AF3’s own confidence measures, there might be other factors influencing performance that are not directly captured by confidence scores. Therefore, we asked how much of the variation in AF3 accuracy across protein–RNA complexes can be explained, and by which sources of information, using grouped cross-validated ridge regression at the complex level (primary target: TM*_c_*; Fig. S35). Throughout, we report cross-validated explained variance (*R*^2^) and interpret it as a measure of statistical association within the analysed PDB distribution rather than as a deployed predictor or a causal claim.

#### Predictive value of individual feature groups

AF3’s own confidence metrics (G0) were by far the strongest single source of information about its accuracy, explaining roughly half of the variance in global fold quality on their own (TM*_c_ R*^2^ *≈* 0.53 across all complexes; *≈* 0.47 for the multi-chain subset), with a moderate rank correlation (Spearman *ρ ≈* 0.68). Among the remaining groups, structural complexity (G2) was the strongest individual contributor for global accuracy (*R*^2^ *≈* 0.24-0.33), followed by predicted-interface features (G4) and experimental context (G5), whereas evolutionary signal (G1) and RNA-intrinsic properties (G3) were individually weak at the complex level (Fig. S35A; Table S10). This picture shifted for the interface-focused target. For DockQ*_Nr_* (and for the whole-complex DockQ), predicted-interface features (G4) rose to become the dominant non-confidence group, consistent with these targets measuring interface rather than global accuracy.

#### Gains from adding feature groups to confidence

Adding each non-confidence group to AF3 confidence individually produced consistent gains (Fig. S35A, “AF3 + one group”), and the full combined model raised the explained variance for TM*_c_* by roughly a third relative to confidence alone, from *R*^2^ *≈* 0.53 (AF3 only) to *R*^2^ *≈* 0.71 across all complexes (multi-chain: 0.47 *→* 0.67). These single-group increments are order-independent by design, since each simply asks what a group adds on top of AF3 confidence. They cannot, however, be used to rank the groups against one another. Because the feature groups are partially correlated, the increments overlap, do not sum to the full-model gain, and the credit assigned to any one group depends on the order in which the groups are entered. To rank the sources fairly we therefore partitioned the full-model explained variance with an order-independent Shapley-*R*^2^ decomposition, which averages each group’s contribution over all group coalitions rather than a single entry order (Section 2.6; Table S11). AF3 confidence accounted for the largest single share of explained variance (*≈*56% for TM*_c_* across all complexes), with the remainder attributable to other, largely orthogonal, information. Structural complexity contributed the largest non-confidence share (*≈*28%), followed by predicted-interface (*≈*10%) and experimental (*≈*1%) features, while evolutionary and RNA-intrinsic groups contributed little at the complex level. For DockQ*_Nr_* the non-confidence signal was instead dominated by predicted-interface features (*≈*33% of explained variance; *≈*29% for the whole-complex DockQ), reinforcing that the determinants of accuracy depend on whether global fold or interface geometry is assessed. Together these results show that AF3 confidence, though the single dominant signal, reports mainly the protein-carried complex. It captures global fold accuracy well but leaves a substantial, non-redundant share of the variance (comparable in aggregate to confidence itself, and concentrated at the interface for DockQ*_Nr_*) to complementary structural and contextual features.

#### Residual analyses reveal signal not captured by AF3 confidence

The residual analysis makes this explicit. Modelling the part of TM*_c_* that a confidence-only model fails to predict, we found the AF3 residual to be itself substantially explainable (residual *R*^2^ *≈* 0.26). The shortfall of AF3 confidence is therefore systematic rather than random noise (Fig. S35B). The Shapley partition of the residual was dominated by structural complexity (*≈*61% of explained residual variance) and predicted-interface features (*≈*27%), with experimental context contributing a further *≈*6% (Supplementary Fig. S36). At the interface the residual is small (DockQ*_Nr_* residual *R*^2^ *≈* 0.08), so the interface descriptors mostly add to, rather than correct, AF3’s confidence (Fig. S37). Notably, complexity and predicted-interface geometry contributed more strongly to explaining residual *error* than to overall accuracy, indicating that large, compositionally complex assemblies are systematically harder than AF3’s confidence anticipates. The standardised coefficients of the full model (Fig. S35C) are consistent with this interpretation. Global PAE is the strongest negative correlate of accuracy, while the dominant RNA-chain fraction, system size and template-related terms are positively associated. We interpret these coefficients as directional and descriptive, because ridge shrinkage and inter-group collinearity mean that individual weights should not be read as independent effect sizes, which is precisely why group-level Shapley partitioning is used for ranking.

**Fig. S35:**
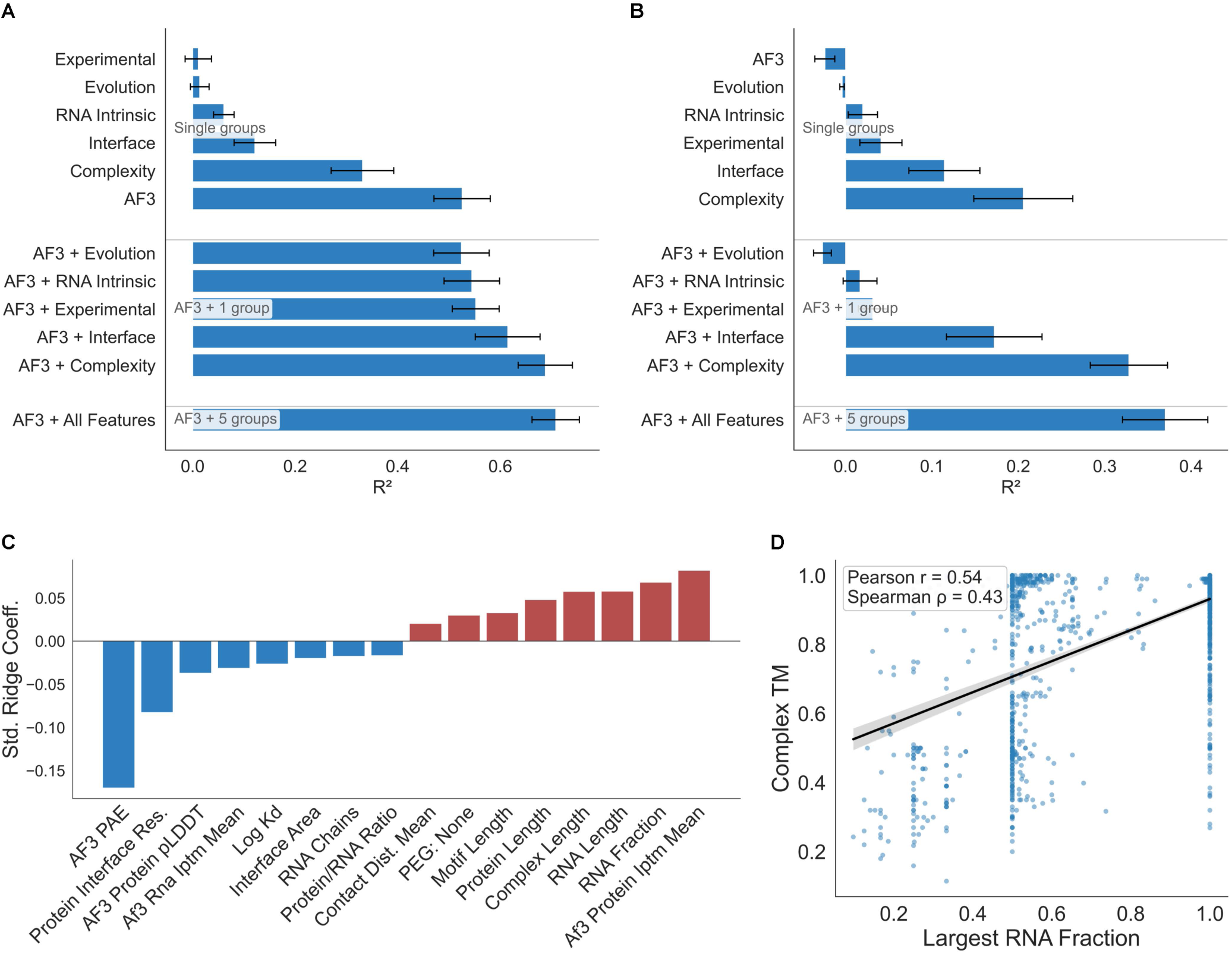
Determinants of AF3 accuracy at the complex level. Grouped cross-validated ridge regression of TM*_c_* for protein–RNA complexes. **A**, *R*^2^ of single feature groups, of AF3 confidence plus each group and of the full model. **B**, *R*^2^ for the residual of the confidence-only model. Complexity and predicted-interface geometry explain more of the residual than of overall accuracy. **C**, Standardised coefficients of the full model (blue, higher accuracy; red, lower). Global PAE is the strongest negative correlate. **D**, TM*_c_* against the dominant RNA-chain fraction.

### R.2. Per system class

#### Single-chain RNA systems

Single-chain RNA per-chain accuracy was the most explainable of all settings (full model *R*^2^ *≈* 0.82). AF3 confidence was again the strongest single group (*R*^2^ *≈* 0.75), but evolutionary information was a strong complementary signal. Adding it to confidence produced the single largest incremental gain, and in the order-independent partition evolution was by far the largest non-confidence contributor (*≈*32% of explained variance; AF3 *≈*43%; Table S11). Broader groups added only marginal gains, and the full model reached its maximum without changing this interpretation. The residual analysis confirmed real non-AF3 signal, arising primarily from evolutionary and RNA-intrinsic properties. This is consistent with single-chain RNA folding being governed by sequence conservation and intrinsic foldability, information only partially reflected in AF3’s confidence.

#### Protein–RNA 1v1 systems

For single protein–RNA complexes, overall per-chain predictability was lower and markedly less stable. AF3 confidence remained the strongest single group, and predicted-interface features were the key complementary signal (the best interpretable model added interface to confidence). However, performance did not improve monotonically as further groups were added. The full model did not exceed, and in our grouped cross-validation slightly underperformed, the confidence-only baseline (Table S10; full *R*^2^ below G0 alone). The Shapley partition exposed the cause. The experimental/chemistry group carried a large *negative* out-of-fold contribution in this small subset, i.e. its many sparse chemical descriptors injected noise rather than signal. Accordingly, the residual TM signal was near zero. We therefore interpret the 1v1 per-chain regime as one of genuinely reduced and heterogeneous predictability, and report it with the explicit caveat that the full model is unstable here. Group shares are not interpreted as a reliable ranking in this subset (and it is omitted from Table S11). Automated stability selection, which reselects features within each cross-validation fold, nonetheless recovers a stable predictive model for this subset (nested *R*^2^ *≈* 0.53, up from an unstable full-model *R*^2^ *≈* 0.18; the confidence-only G0 baseline is *≈* 0.57), indicating that the instability lies in the group-level attribution rather than in predictability itself. The interface-focused DockQ*_Nr_* target showed the same instability in this subset (confidence-only *R*^2^ *≈* 0.41, full model *≈* 0.28), so interface determinants of 1v1 complexes are read from the pooled analysis.

#### Multi-chain complexes

For multi-chain systems the AF3 confidence baseline was weaker than in the single-chain RNA and 1v1 classes (*R*^2^ *≈* 0.39), and the gains from additional information were distributed across several groups rather than dominated by any single factor (full *R*^2^ *≈* 0.48). Predicted-interface and complexity features were both important contributors (interface *≈*19% and complexity *≈*12% of explained variance, with evolution contributing a further *≈*17%; Table S11), reflecting the increased role of inter-chain interactions and global assembly in multi-chain complexes. RNA-intrinsic descriptors were largely unavailable or uninformative in this class. The residual analysis showed a modest but real non-AF3 signal driven by complexity and interface context.

#### Multi-chain RNA complexes

The 106 complexes of two or more RNA chains without protein were analysed at the per-chain level (*n* = 306 chains). Protein-side descriptors, alignment depth, thermodynamic descriptors and the experimental-context group are not available for this set, so G1 reduces to the RNA family, G3 to sequence-derived descriptors (GC content, entropy, complexity) and G4 to the predicted RNA–RNA interface (contacting residues, contact distances, partner chains, hydrogen bonds). AF3 confidence explained *R*^2^ *≈* 0.34 and the full model *≈* 0.41. Complexity (*≈*33%) and the predicted RNA–RNA interface (*≈*16%) were the main complements, the family label added little (*≈*7%), and the sequence-derived RNA descriptors did not improve held-out prediction (negative share; Table S11). The residual was barely explainable (*R*^2^ *≈* 0.09), and stability selection retained 10 of 14 candidates (0.414 *→* 0.392 nested; Table S10, Fig. S38).

#### Comparison of predictive determinants across system classes

Across classes a consistent two-part picture emerges. AF3 confidence is the single most informative source of information about its own accuracy everywhere, yet it is consistently incomplete, leaving a large and *structured* residual. What fills that gap is class- and metric-dependent in a mechanistically sensible way. It is evolutionary signal in single-chain RNA, predicted-interface and, for global fold, complexity features in multi-chain complexes, and predicted-interface features specifically for interface accuracy (DockQ). This shift is summarised directly by the order-independent contribution contrast between global and per-chain models (Fig. 5E), in which AF3 confidence and complexity weigh more on global-complex accuracy while evolution weighs more on per-chain accuracy. The residual signal was strongest and cleanest in single-chain RNA systems, weakest and least stable in 1v1 complexes, and moderate in multi-chain assemblies. The total explainability also varied widely (from *R*^2^ *≈* 0.82 for single-chain RNA to *≈* 0.28 for 1v1 DockQ*_Nr_*), itself an informative statement about where AF3 behaviour is, and is not, accountable to measurable structural and biochemical properties. The regressions therefore establish *that* a structured signal beyond confidence exists and quantify it, without attributing it to a specific mechanism.

## S. Feature Selection for Regression Analysis

### S.1. Descriptor groups

Group G0 comprises AF3’s own confidence signals, spanning both the RNA and the protein side of each prediction, namely RNA and protein pLDDT, RNA and protein interface predicted-TM (ipTM), protein pTM (and, at the per-chain level, RNA pTM), the global PAE and the minimum chain-pair PAE. They quantify AF3’s assessment of local, chain-level and inter-chain reliability and serve as the baseline against which all other information is measured. At the complex level the per-chain confidence scores enter as chain-averaged summaries, while the raw per-chain values are used in the per-chain analyses.

Group G1 (evolutionary signal) captures the depth of available homology information, namely the RNA and protein MSA depths (each as log_10_ of the alignment size), the number of unique RNA families and the RNA family annotation, together with template availability, summarised by the counts of protein templates and of RNA-binding-protein (RBP) complex templates, the structural-prior component of the evolutionary signal. The RNA family label was retained primarily for the stratified family-based and pre-/post-cutoff generalisation analyses. In the pooled regressions it enters only where it survives the cardinality and frequency filters of the preprocessing.

Group G2 describes system size and composition, namely the numbers of RNA and protein chains, the total RNA, protein and complex lengths, the protein-to-RNA length ratio, and the fraction of the complex’s total RNA length contributed by its longest RNA chain (equal to 1 for single-RNA systems). At the per-chain level the length of the individual RNA chain is included as well.

Group G3 (RNA intrinsic) collects sequence- and folding-derived descriptors computed independently of the predicted structure, namely sequence complexity and Shannon entropy, the ViennaRNA minimum free energy, ensemble diversity and centroid energy and, where available, the log_10_ dissociation constant (log_10_ *K_d_*). At the complex level they enter in their per-complex aggregated forms, and at the per-chain level the per-chain raw descriptors and DSSR-derived secondary-structure counts are used where present. Group G4 (interface) collects descriptors of the predicted protein–RNA interface, computed exclusively from the predicted structure, namely predicted interface area and ratio, numbers of unique RNA and protein interface residues, RNA interface coverage, contact-distance statistics (mean and median), the fraction of close (*<* 3.5 Å) contacts, the fraction of interface contacts made to cytosine and the interface RNA-motif extent (motif length and coverage), with chain-resolved counterparts at the per-chain level. Because they are derived from the prediction rather than the native complex, they are not circular with the accuracy target and measure the self-consistency of the predicted interface.

**Table S10.** Automated stability selection preserves predictive performance. For each subset and level, the number of candidate descriptors, the number retained by grouped stability selection (ElasticNet, 100 resamples of PDB groups, selection frequency *≥* 0.70), and the cross-validated *R*^2^ of the full and the nested stability-selected sets (mean over folds). Selection preserves *R*^2^ in every stable subset. For the unstable 1v1 per-chain subset, nested selection recovers a stable model (0.18 *→* 0.53).

| Dataset | Target | Candidate | Selected | Grouped CV *R*<sup>2</sup> |  |
|---|---|---|---|---|---|
|  |  | features | features | Full set | Selected (nested) |
| All-complex | TM<sub>*c*</sub> | 40 | 38 | 0.710 | 0.710 |
| All-complex | DockQ<sub>Nr</sub> | 40 | 15 | 0.552 | 0.547 |
| All-complex | DockQ (whole complex) | 40 | 37 | 0.459 | 0.459 |
| Multi-chain | TM<sub>*c*</sub> | 40 | 38 | 0.666 | 0.664 |
| Multi-chain | TM<sub>*r*</sub> (per-chain) | 35 | 34 | 0.478 | 0.476 |
| Single-chain RNA | TM<sub>*r*</sub> | 22 | 9 | 0.836 | 0.837 |
| Multi-chain RNA | TM<sub>*r*</sub> (per-chain) | 14 | 10 | 0.414 | 0.392 |
| 1v1 | TM<sub>*r*</sub> (per-chain) | 42 | 8 | 0.175 | 0.531 |

Group G5 captures the chemical and experimental context of each structure, namely the numbers of listed ions and ligands, ionic strength, buffer molarity and category, PEG presence, percentage and category, experimental pH and temperature. The X-ray resolution and the X-ray indicator, which are known only once the experimental structure exists, are excluded from the regressions reported here (they can be re-enabled for the dedicated X-ray-versus-EM analysis).

### S.2. Stability selection

To analyse determinants of AF3 performance, we initially compiled a set of 163 features, including direct descriptors (e.g., chain length, RNA/protein counts, MSA depth) and derived quantities (e.g., aggregated complex-level properties and interface ratios). All features were organised into the six groups defined above (G0-G5) and their union (G*_X_*).

Because the full feature set contains partially redundant and correlated variables, directly fitting all of them yields unstable coefficient estimates and reduced interpretability. We therefore reduced each dataset’s feature set with an automated, leakage-controlled *stability-selection* procedure, which replaces the earlier hand-curated pruning.

For each dataset subset (single-chain RNA, 1v1 and multi-chain protein–RNA complexes, and multi-chain RNA complexes) and analysis level (single-chain or complex-level), we drew 100 grouped resamples, subsampling complexes by pdb_id so that all chains of a complex stay together, and fit an ElasticNet (with the mixing and penalty strength chosen once by internal cross-validation) on each resample. A descriptor was retained if it was selected (assigned a non-zero coefficient) in at least 70% of the resamples. To obtain an honest estimate of the retained set’s performance, the whole selection was then re-run *inside* each grouped cross-validation training fold (nested selection), so the reported selected-set *R*^2^ never sees the held-out data.

#### Preprocessing and feature screening

All models share a single preprocessing pipeline that is fit within each training fold only, so that no test-fold information leaks into imputation or scaling. Numeric features were median-imputed and standardised (zero mean, unit variance). Categorical features were imputed with the most frequent level and one-hot encoded, with levels occurring fewer than ten times collapsed into an “infrequent” category. After encoding, a near-zero-variance filter (*σ*^2^ *<* 10*^−^*^8^) removed constant columns. Prior to modelling, and independently of the target, candidate features were screened on the pooled data to remove columns that were entirely missing, observed in fewer than ten samples, more than 80% missing, constant, or (for categoricals) of excessively high cardinality (*>* 25 levels). Because this screen is unsupervised, it does not bias the cross-validated estimates. The ridge penalty (*α* = 10), the number of folds (*k* = 5) and the random seed (0) were fixed a priori and held constant across all models, and the variance partition is insensitive to *α* over two orders of magnitude (Table S12).

Table S10 reports, for every subset, the number of candidate descriptors, the number retained, and the cross-validated *R*^2^ of the full candidate set versus the nested stability-selected set. Selection preserved performance almost exactly in every stable subset (e.g. all-complex TM*_c_* 0.710 *→* 0.710, single-chain RNA 0.836 *→* 0.837), confirming that the trimmed sets are parsimonious without loss.

Notably, for the 1v1 per-chain subset, where the full model does not exceed the confidence baseline out of fold, nested selection *improves* held-out *R*^2^ from 0.18 to 0.53, recovering a stable predictive model. The instability is therefore in the attribution, not the intrinsic predictability.

The descriptors retained at the complex level are listed by group in Table S13 (the per-chain subsets retain the raw per-chain descriptors in place of the aggregated complex-level ones). The selected set spans all six groups and includes, at G0, the protein-side confidence signals (protein ipTM/pTM/pLDDT together with RNA ipTM and the global and minimum chain-pair PAE) and, at G1, both the RNA and protein MSA depths alongside the RBP- and protein-template counts.

**Table S11.** Shapley partition of the cross-validated explained variance over the six feature groups. 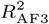 is the confidence-only and 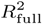 the full model. Group columns give each group’s share of 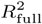 (%), averaged over all orderings. The 1v1 per-chain subset is omitted because its full model did not exceed the confidence-only baseline. DockQ denotes the whole-complex DockQ, reported as a sensitivity variant of DockQ*_Nr_*. A share rounding to 0 marks a group that does not improve held-out prediction. G0 is AF3 confidence, G1 evolution, G2 complexity, G3 RNA-intrinsic properties, G4 the predicted interface and G5 the experimental context.

| Dataset | Target | *n* | *R*<sup>2</sup><sub>AF3</sub> | *R*<sup>2</sup><sub>full</sub> | G0 | G1 | G2 | G3 | G4 | G5 |
|---|---|---|---|---|---|---|---|---|---|---|
| All complexes | TM<sub>*c*</sub> | 1306 | 0.53 | 0.71 | 56 | 2 | 28 | 3 | 10 | 1 |
| Multi-chain complexes | TM<sub>*c*</sub> | 962 | 0.47 | 0.67 | 59 | 1 | 25 | 2 | 11 | 3 |
| Protein–RNA 1v1 | TM<sub>*c*</sub> | 344 | 0.47 | 0.46 | 84 | 3 | 6 | 3 | 3 | 1 |
| All complexes | DockQ<sub>*Nr*</sub> | 1287 | 0.49 | 0.55 | 55 | 2 | 6 | 5 | 33 | 0 |
| Multi-chain complexes | DockQ<sub>*Nr*</sub> | 943 | 0.50 | 0.56 | 56 | 2 | 6 | 3 | 32 | 1 |
| All complexes | DockQ | 1301 | 0.36 | 0.46 | 55 | 3 | 9 | 4 | 29 | 0 |
| Multi-chain complexes | DockQ | 957 | 0.38 | 0.49 | 60 | 2 | 9 | 3 | 27 | -1 |
| Multi-chain (per-chain) | TM<sub>*r*</sub> | 1996 | 0.39 | 0.48 | 50 | 17 | 12 | 1 | 19 | 0 |
| Single-chain RNA (per-chain) | TM<sub>*r*</sub> | 165 | 0.75 | 0.82 | 43 | 32 | 4 | 12 | 0 | 10 |
| Multi-chain RNA (per-chain) | TM<sub>*r*</sub> | 306 | 0.34 | 0.41 | 52 | 7 | 33 | -8 | 16 | 0 |

**Table S12.** Ridge-penalty (*α*) sensitivity of the Shapley-*R*^2^ partition (all-complex TM*_c_*, full feature universe). Group columns give each group’s share (%) of the full-model *R*^2^, and the ridge penalty *α* = 10 used throughout was fixed a priori. The partition is essentially invariant over two orders of magnitude in *α*, confirming that the reported group ranking is not an artefact of the chosen penalty.

| α | G0 | G1 | G2 | G3 | G4 | G5 | *R*<sup>2</sup><sub>full</sub> |
|---|---|---|---|---|---|---|---|
| 1 | 56 | 2 | 28 | 3 | 10 | 1 | 0.709 |
| 10 | 56 | 2 | 28 | 3 | 10 | 1 | 0.710 |
| 100 | 56 | 2 | 28 | 3 | 9 | 2 | 0.689 |

### S.3. Expected accuracy by system class and RNA length

Table S14 lists AF3 accuracy as conditional means on the working set, conditioning only on properties known before prediction. **System class.** Expected per-chain TM*_r_* falls from 0.66 for single-chain RNA through 0.58 for single protein–RNA (1v1) complexes to 0.50 for chains of multi-chain assemblies, so whether a chain folds alone or as part of a protein complex shifts the expected fold accuracy by about 0.16 TM*_c_*. Global and interface accuracy diverge at the complex level. TM*_c_* is high for 1v1 complexes (0.95), but interface accuracy is only moderate (DockQ *≈* 0.68 for 1v1 and *≈* 0.59 for multi-chain complexes), so a high complex TM*_c_* is not evidence of a reliable interface.

#### RNA length

Longer RNA chains are modelled better, not worse. Single-chain RNA accuracy rises from 0.38 at *≤* 40 nt to 0.86 above 200 nt, and multi-chain RNA from 0.46 to 0.96 across the same range. This ordering reflects alignment availability and family composition rather than length itself. Of the 55 RNA chains longer than 200 nt, 54 fall into three deeply conserved and structurally redundant groups, small-subunit ribosomal RNA (*n* = 35), group II intron domains (*n* = 8) and RNase P (*n* = 11), so the upper length bins test the best-aligned and most heavily represented folds in the PDB rather than long RNA as such. Consistent with this, the advantage reverses when the RNA alignment is withheld (Section 3.2), and it is not removed by splitting on release date, since a post-cutoff ribosomal chain remains family-identical to training-era ones. Short (*≤* 40 nt) RNAs are the least reliable stratum in every class. The largest, ribosome-scale assemblies fall outside the analysable set for resource reasons (Appendix A.2), so the upper end of the length gradient reflects large-but-tractable rather than the very largest RNAs. Length is also where AF3 sits furthest from its training regime. It is trained on crops of 384 to 768 tokens and run at inference without cropping [19]. Tokens are shared between the RNA and its protein partners, so 67% of our post-cutoff complexes are larger than anything the model saw intact during training. Because most crops are selected by spatial rather than sequential proximity, local geometry survives cropping and global architecture does not, consistent with the locally sound but globally wrong long-RNA models reported by independent evaluations [24, 32].

**Fig. S36:**
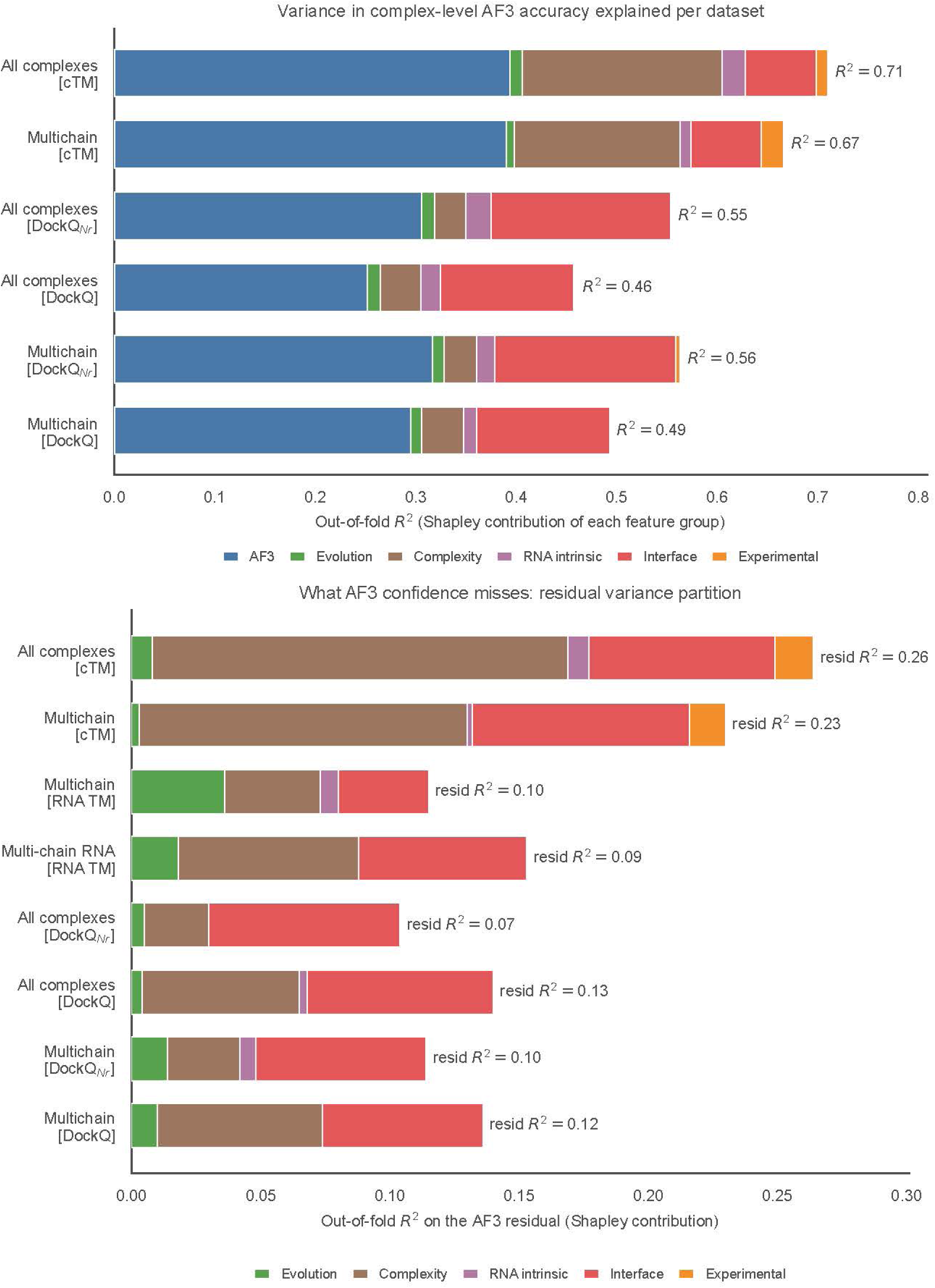
Order-independent (Shapley-*R*^2^) variance partition. **(top)** Direct-target partition at the complex level. Bar length is the full-model out-of-fold *R*^2^ and each coloured segment is a feature group’s Shapley contribution, for the all-complex and multi-chain datasets and for the global (TM*_c_*) and interface (DockQ*_Nr_*, with the whole-complex DockQ as sensitivity variant) targets. **(bottom)** Partition of the AF3 residual, the variance not captured by a confidence-only model, over the five non-AF3 groups, showing that complexity, experimental context and predicted-interface features explain the structured part of AF3 error. Full-model *R*^2^ values reproduce the full-set column of Table S10.

**Fig. S37:**
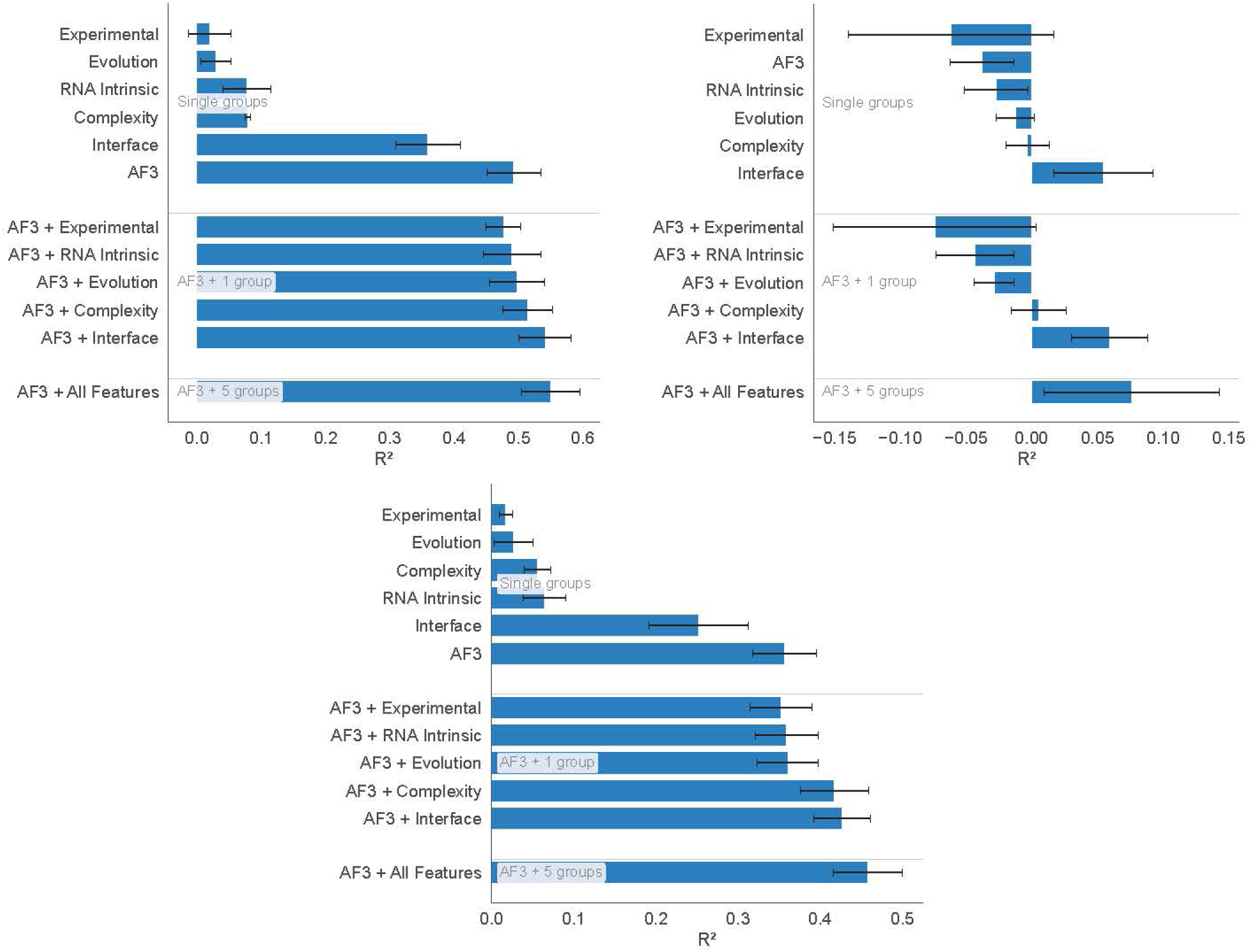
Determinants of interface accuracy at the complex level. Grouped cross-validated *R*^2^ of single feature groups, of AF3 confidence plus each group and of the full model, for all protein–RNA complexes. **Top left**, DockQ*_Nr_*. **Top right**, residual of the DockQ*_Nr_* confidence-only model. **Bottom**, whole-complex DockQ (sensitivity variant).

#### Training-era and novel targets

Splitting the same branches by release date shows that the combined values are inflated by training-era familiarity. Per-chain TM*_r_* drops by 0.11 for single-chain RNA, 0.14 for multi-chain and 0.30 for 1v1 chains on post-cutoff targets, the 1v1 expectation falling from 0.64 to 0.34. The post-cutoff single-chain RNA value (0.59) reproduces the independent family-based estimate of Section 3.1. For a novel target the post-cutoff column should therefore be read, which makes the lookup a generalisation-aware expectation rather than an in-sample fit.

## T. Decision tree: final model, development and independent test

This appendix defines the decision tree of Fig. 6, how its questions were chosen, and its test on PDB entries released in 2026. It ends with the expected-accuracy table of Section 3.6. The tree is implemented in the decision_tree package of the code repository, which reads raw AF3 output.

### T.1. Final model

The tree gives calibrated probabilities for the RNA fold and the protein–RNA interface of a prediction, and no input requires an RNA alignment depth or an experimental structure. The intra-RNA PAE of an RNA chain is the mean of AF3’s predicted aligned error matrix (pae in the full-data JSON, rows and columns selected by token_chain_ids) over the token pairs within that chain. It is our summary of AF3’s output, not a value that AF3 reports. For protein–RNA complexes, the log-odds that the longest RNA chain is well folded are *−*0.879 *−* 0.207 PAE + 1.063 *s* + 1.668 *f* + 0.978 *t* + 1.555 *m*, whereby PAE is the intra-RNA PAE of that chain in Å; *s* = 1 for a familiar sequence, an RNA chain with *≥* 80% identity (identical aligned columns over the longer sequence) to an RNA chain of a PDB entry released before 30 September 2021 (3,891 unique sequences from 5,639 entries); *f* = 1 for a familiar fold, where the longest RNA chain of the AF3 model reaches a TM*_r_ ≥* 0.5 (normalised by the model chain) to an RNA chain of the pre-cutoff structures of this benchmark, and chains shorter than 20 nt count as not familiar; *t* = 1 for an RBP template, i.e. a protein–RNA complex among AF3’s template hits for any protein chain; and *m* = 1 for two or more RNA chains in the model. The log-odds that an RNA chain forms a correct interface are 0.709 *−* 0.201 PAE_min_ + 1.524 *u* + 0.651 *w*, with PAE_min_ the minimum chain-pair PAE between the RNA chain and a protein chain (chain_pair_pae_min), *u* = 1 for a familiar RNA (sequence or fold) and *w* = 1 for a well-folded partner (pTM*_p_ ≥* 0.84 of the partner protein and pLDDT*_r_ ≥* 0.78). Each Å of intra-RNA PAE thus multiplies the odds of a well-folded RNA by 0.81, and the four questions multiply them by 2.9, 5.3, 2.7 and 4.7 (Table S15). Each log-odds *z* gives the probability 1*/*(1 + *e^−z^*). The probability that an RNA chain is usable, that is well folded and placed correctly, is Φ_2_ Φ*^−^*^1^(*P*_fold_), Φ*^−^*^1^(*P*_interface_); *ρ* with *ρ* = 0.66, where Φ is the standard normal distribution function and Φ_2_ the bivariate one. All probabilities are reported at three levels, WEAK below 25%, MODERATE from 25% to below 60% and STRONG from 60%. Single-chain RNAs and RNA complexes without protein are scored by the RNA-only subtrees of Appendix T.4. A large RBP domain (dsRBD, RNase III, Dicer) is reported but not scored. If an answer is unknown, the probability is computed over its possible values (both answers of a yes/no question, or the 5th to 95th percentile of the post-cutoff targets for a count or score), and the resulting range is reported.

**Fig. S38:**
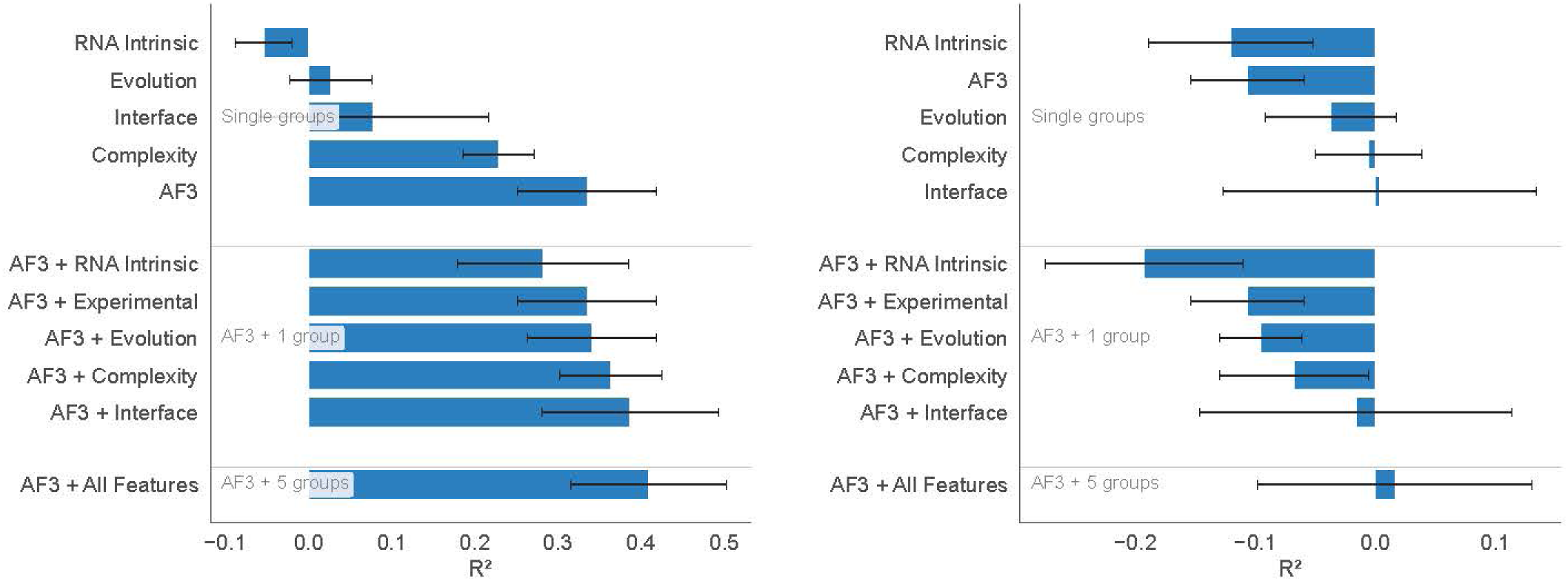
Determinants of per-chain accuracy in multi-chain RNA complexes. As Fig. 5A–C, for the 306 RNA chains of the 106 multi-chain RNA complexes without protein (TM*_r_*; **left**, direct target; **right**, residual of the confidence-only model). The experimental-context group has no descriptors in this set.

**Table S13.**
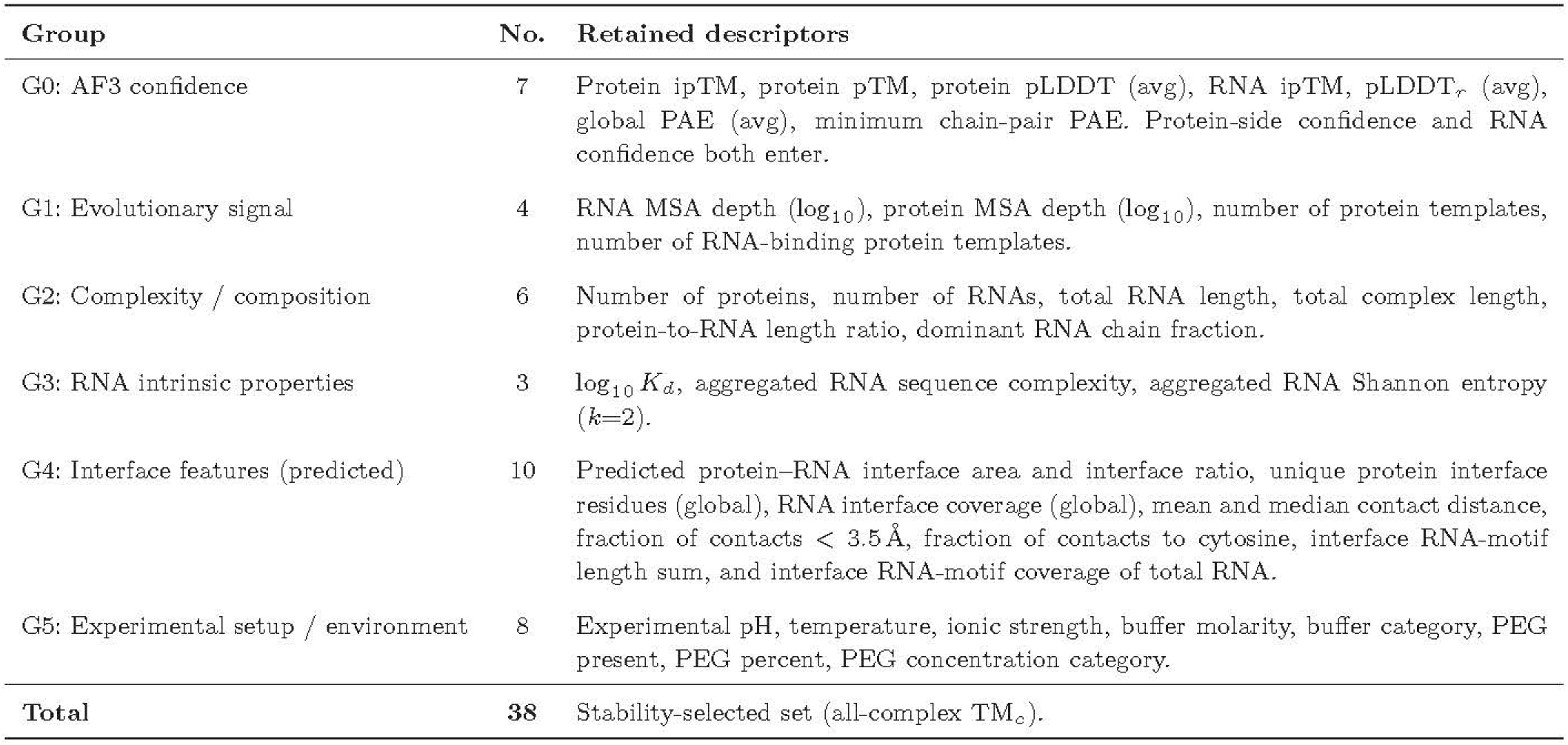
Descriptors retained by the automated stability selection, grouped by category. The set shown is the all-complex TM*_c_* model (38 of 40 candidates retained at selection frequency *≥* 0.70). The DockQ_Nr_ and per-chain subsets differ slightly (per-chain models substitute the raw per-chain descriptors for the aggregated complex-level ones, see Table S10 for retained counts). “No.” is the number of retained descriptors per group. Interface descriptors (G4) are computed from the *predicted* structure and are therefore not circular with the accuracy target.

| Group | No. | Retained descriptors |
|---|---|---|
| G0: AF3 confidence | 7 | Protein ipTM, protein pTM, protein pLDDT (avg), RNA ipTM, pLDDT<sub>*r*</sub> (avg), global PAE (avg), minimum chain-pair PAE. Protein-side confidence and RNA confidence both enter. |
| G1: Evolutionary signal | 4 | RNA MSA depth (log<sub>10</sub>), protein MSA depth (log<sub>10</sub>), number of protein templates, number of RNA-binding protein templates. |
| G2: Complexity / composition | 6 | Number of proteins, number of RNAs, total RNA length, total complex length, protein-to-RNA length ratio, dominant RNA chain fraction. |
| G3: RNA intrinsic properties | 3 | log<sub>10</sub> *K*<sub>*d*</sub>, aggregated RNA sequence complexity, aggregated RNA Shannon entropy (*k*=2). |
| G4: Interface features (predicted) | 10 | Predicted protein–RNA interface area and interface ratio, unique protein interface residues (global), RNA interface coverage (global), mean and median contact distance, fraction of contacts < 3.5 Å, fraction of contacts to cytosine, interface RNA-motif length sum, and interface RNA-motif coverage of total RNA. |
| G5: Experimental setup / environment | 8 | Experimental pH, temperature, ionic strength, buffer molarity, buffer category, PEG present, PEG percent, PEG concentration category. |
| **Total** | **38** | Stability-selected set (all-complex TM<sub>*c*</sub>). |

**Table S14.** Observed AF3 accuracy by system class and RNA length, as conditional means (per-chain TM*_r_*, mean and SD, *n* RNA chains) for all data and before and after the training cutoff. The largest ribosome-scale assemblies are absent (Appendix A.2). Bins with *n <* 15 (marked *^∗^*) are indicative only.

| Branch | Combined, *n* | Pre-cutoff, *n* | Post-cutoff, *n* |
|---|---|---|---|
| *System class* |  |  |  |
| single-chain RNA | 0.66 (0.27), 165 | 0.70, 99 | 0.59, 66 |
| Protein–RNA 1v1 | 0.58 (0.26), 344 | 0.64, 270 | 0.34, 74 |
| Protein–RNA multi-chain | 0.50 (0.21), 1996 | 0.54, 1416 | 0.40, 580 |
| *RNA chain length* |  |  |  |
| Single-chain RNA, ≤ 40 nt | 0.38, 35 |  | 0.35, 11<sup>*</sup> |
| Single-chain RNA, 41-100 nt | 0.69, 84 |  | 0.41, 22 |
| Single-chain RNA, 101-200 nt | 0.68, 8<sup>*</sup> |  | 0.52, 3<sup>*</sup> |
| Single-chain RNA, > 200 nt | 0.86, 38 |  | 0.82, 30 |
| Multi-chain, ≤ 40 nt | 0.46, 1672 |  | 0.38, 476 |
| Multi-chain, 41-100 nt | 0.61, 267 |  | 0.49, 97 |
| Multi-chain, 101-200 nt | 0.74, 12<sup>*</sup> |  | 0.39, 3<sup>*</sup> |
| Multi-chain, > 200 nt | 0.96, 45 |  | 0.88, 4<sup>*</sup> |

**Table S15.** Log-odds of the protein–RNA probabilities. Each probability is 1*/*(1 + *e^−z^*), where *z* is the intercept plus the sum of coefficient times input. Odds ratios are per unit of a continuous input or for a yes answer. Inputs were selected by cross-validated forward selection on post-cutoff targets, and the weights were fitted on the same targets. The usable-chain probability joins P_fold_ and P_interface_ by a Gaussian copula with correlation 0.66.

| Input | Coefficient | Odds ratio |
|---|---|---|
| *P*<sub>fold</sub> *(well-folded RNA)* |  |  |
| intercept | −0.879 |  |
| intra-RNA PAE of the longest RNA chain (Å) | −0.207 | 0.81 |
| familiar sequence: an RNA of a PDB entry released before 30 September 2021 with ≥ 80% identity (yes = 1) | +1.063 | 2.90 |
| familiar fold: AF3 model's longest RNA chain has a pre-cutoff RNA with TM<sub>*r*</sub> ≥ 0.5 (yes = 1) | +1.668 | 5.30 |
| AF3 used an RNA-bound protein template for any protein chain (yes = 1) | +0.978 | 2.66 |
| ≥ 2 RNA chains in the model (yes = 1) | +1.555 | 4.74 |
| *P*<sub>interface</sub> *(correct interface)* |  |  |
| intercept | +0.709 |  |
| minimum chain-pair PAE between the RNA chain and a protein chain (Å) | −0.201 | 0.82 |
| familiar RNA: familiar sequence or familiar fold (yes = 1) | +1.524 | 4.59 |
| well-folded partner: pTM<sub>*p*</sub> ≥ 0.84 and pLDDT<sub>*r*</sub> ≥ 0.78 (yes = 1) | +0.651 | 1.92 |

### T.2. Development

#### Interface side

The tier cut-offs were set on the DockQ*_nr_* classes of the working set. Among three warning signs (large RBP domain, no RBP template, poorly folded partner), the poorly folded partner and the large domain together improved the ranking of pre-cutoff targets most. Together they raised the AUC from 0.731 without a warning sign to 0.739. The poorly folded protein partner alone gave 0.737 and the large domain alone 0.733, carried by 21 pre-cutoff complexes. A missing template lowered the AUC to 0.698. The levels were calibrated on post-cutoff targets, with 39% [35-44%], 89% [80-94%] and 88% [80-94%] correct interfaces at UNRELIABLE, CAUTION and RELIABLE. On post-cutoff targets the bad interface tier captures most RNAs placed at a distal site but only 44% of those posed incorrectly at the correct site (Section 3.3), the failure that AF3’s RNA interface confidence largely misses.

#### Fold side

The confidence level, given by the mean intra-RNA PAE, was chosen on pre-cutoff targets among pLDDT*_r_* combined with the global PAE and the intra-RNA PAE (AUC 0.741, against 0.735 and 0.715-0.728). The further questions were selected on post-cutoff protein–RNA targets (383 RNAs, 168 well folded), the regime of new predictions, among 25 candidates known before prediction or read from the AF3 model (sequence- and structure-familiar RNA, Rfam family, pseudoknots, stems, hairpins, internal and multibranch loops of the model, RNA length, numbers of RNA and protein chains, GC content, folding energy per nucleotide, sequence complexity, RBP and protein-only templates used and their numbers, large RBP domain, RNA pLDDT, pTM*_r_*, global PAE, fraction disordered, predicted interface area). Each question was a boolean or a continuous feature at its 25th, 50th or 75th percentile, with cut-off and sign set on the training folds of a 5*×*-repeated 5-fold cross-validation grouped by PDB entry. Questions were added greedily while the cross-validated AUC rose by at least 0.01, at most four (python -m decision_tree.develop_v3). Two or more RNA chains (cross-validated AUC 0.674 *→* 0.745), an RBP template (0.775), a familiar fold (0.791) and a familiar sequence (0.804) were selected, and the levels are the post-cutoff tertiles of the score.

#### Earlier versions

Two earlier forms were superseded. Gated trees that ask the pre-prediction questions (Rfam family, RNA MSA depth, pseudoknot, RBP template, domain) in a fixed order ranked post-cutoff targets close to chance when AF3’s confidence was asked last (AUC 0.44 for the fold, 0.61 for the interface) and moderately when it was asked first (0.60 and 0.67), because the questions on RNA family and alignment depth point the wrong way on novel targets. A fold score of the confidence level plus one point for a familiar RNA, designed on pre-cutoff targets (post-cutoff AUC 0.75), did not exceed AF3’s confidence alone on the 2026 set. The same selection procedure applied to the interface side chose pTM*_p_*, a familiar fold, three or more protein chains and pLDDT*_r_* (cross-validated AUC 0.847 against 0.718). These point scores are superseded. The inputs of the final score were selected anew by the automatic procedure described next, on post-cutoff targets alone.

#### Probability form

Logistic regressions replaced the point scores, because a fixed number of points per answer cannot weigh the questions against AF3’s continuous confidence. Both familiarity questions are computed without the experimental structure (Final model). The weights were fitted on post-cutoff targets (382 RNAs, 167 well folded; 668 RNA chains, 344 correct interfaces), with the intra-RNA PAE and the chain-pair PAE in place of AF3’s confidence levels (python -m decision_tree.develop_probability). pLDDT*_r_* for the fold and ipTM*_r_* for the interface would have been added only if they raised the cross-validated AUC (5*×*-repeated 5-fold, grouped by PDB entry) by at least 0.01. Neither did (*−*0.002 and *−*0.003), and the cross-validated AUC is 0.853 for the fold and 0.845 for the interface. Two later candidates were not adopted either. The continuous pTM*_p_* raised the cross-validated interface AUC (0.844 *→* 0.868) but lowered it on the 2026 set, and secondary-structure features from ViennaRNA gained at most 0.008. Fold and interface are not independent. Out of fold, the product of the two probabilities underestimated the share of usable chains (31% predicted, 36% observed) and the smaller of the two overestimated it (39%). A Gaussian copula whose one correlation was fitted by maximum likelihood (*ρ* = 0.66) predicted 36% with the lowest log-loss (0.425, against 0.438 and 0.429). The level cut-offs of 25% and 60% were set in advance. On post-cutoff targets 10%, 41% and 83% of RNAs are well folded at WEAK, MODERATE and STRONG, 11%, 41% and 83% of interfaces are correct, and 7%, 45% and 78% of RNA chains are usable.

### T.3. Independent test on PDB entries released in 2026

#### Cohort

99 protein–RNA complexes released between 13 May and 30 September 2026 (96 cryo-EM, 3 X-ray), selected with the filters of the main data set (of 301 cryo-EM entries, 120 were excluded for modified residues or ligands, 76 for discontinuous RNA chains and 9 for size), predicted with the AlphaFold Server (seed 1, model 0) and processed with the same pipeline (python -m decision_tree.validate_2026). None is part of the pre- or post-cutoff sets. The final score was selected, fitted and frozen on post-cutoff targets before it was applied to the 2026 set, and no 2026 value entered its design. **Labels** follow the definitions used for the post-cutoff set. An RNA is well folded when the 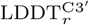 of its longest chain exceeds 0.85, computed as the backbone 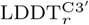 (Appendix A.10) from the aligned RNA chains or, for four complexes without them, directly from the structures (identical classification for all 95 complexes with both). An RNA chain forms a correct interface when its best protein–RNA DockQ is *≥* 0.23 (OpenStructure comparison of the SWISS-MODEL server; run locally with the same settings for two complexes, reproducing the server value), which reproduces the post-cutoff placement labels for 585 of 586 chains. Familiarity was computed as for post-cutoff targets, without the experimental structure (63 of 99 RNAs familiar by sequence, 12 by the fold of the AF3 model, 69 by either).

#### Results

On all 99 complexes (99 RNAs, 28 well folded; 154 RNA chains with a protein–RNA DockQ, 70 correct interfaces, 33 usable; Fig. S39), P_fold_ reaches AUC 0.886. The intra-RNA PAE reaches 0.819 (ΔAUC +0.067, 95% CI *−*0.001–0.142, paired bootstrap over complexes), pLDDT*_r_* 0.687 (+0.200, 0.078–0.326), the earlier point score 0.837 and the gated trees 0.554 and 0.351 (confidence first and as first drawn, without their alignment question, since the AlphaFold Server exports no RNA alignment). P_interface_ reaches 0.923, against 0.894 for the chain-pair PAE (+0.030, 0.001–0.066), 0.866 for ipTM*_r_* (+0.057, 0.002–0.123) and 0.877 for the earlier point score. An RNA chain is usable when it is well folded (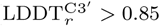 against the experimental chain of the same sequence) and placed correctly, and it carries P_fold_ of its complex. P_usable_ reaches 0.911, against 0.796 for AF3’s ranking score (+0.115, 0.031–0.200), 0.819 for the chain’s intra-RNA PAE (+0.092, 0.025–0.171) and 0.891 for the earlier sum of point scores (Fig. S39C, F, I; python -m decision_tree.develop_probability –test). At WEAK, MODERATE and STRONG, 4%, 33% and 71% of RNAs are well folded (*n* = 48, 27 and 24; mean predicted 7%, 42% and 78%), 10%, 35% and 96% of interfaces are correct (67, 34 and 53; predicted 9%, 44% and 80%) and 3%, 23% and 66% of RNA chains are usable (89, 30 and 35; predicted 7%, 40% and 73%). The interface probabilities are thus slightly low on the 2026 set (mean 41% against 45% correct), and the fold and usable probabilities slightly high (34% against 28%, and 28% against 21%). The Brier score is 0.127, 0.120 and 0.106, against 0.203, 0.248 and 0.168 for a constant at the observed rate. One RNA chain without any protein–RNA DockQ was left out. A conserved Rfam family and a pseudoknot in the model mark poorly folded RNAs (10% and 3% well folded, against 40% and 41% without).

#### T.4. Subtrees for RNA-only predictions

The fold probability above was designed on protein–RNA complexes. RNA-only predictions follow two further subtrees, one for single-chain RNAs and one for RNA complexes without protein, so the decision tree has three subtrees in all. Each RNA-only subtree has two branches, for predictions made with and without the RNA alignment, because the alignment changes both the accuracy and the confidence measure. Among the 46 post-cutoff single-chain RNAs with an alignment (depth *>* 1) predicted both ways, median TM*_r_* fell from 0.83 with the alignment to 0.26 without it (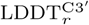 0.78 against 0.50). The per-residue measures, pLDDT and the intra-RNA PAE, track this accuracy, whereas pTM*_r_* does not. In both subtrees a prediction is well folded when 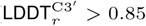, computed as in the main analysis, and a logistic regression on the chosen confidence measure, fitted with mild L2 regularisation because the development sets are nearly separable, gives the probability, with the same levels as for protein–RNA complexes (Table S16).

##### Single-chain RNAs

The development set is the post-cutoff single-chain RNAs, 66 predicted with the alignment (15 well folded) and 48 without it (5 well folded). The latter are 28 runs without the alignment and 21 RNAs without any alignment, whose single run serves both branches. Among pTM*_r_*, mean pLDDT*_r_* and the intra-RNA PAE, pLDDT*_r_* ranked well-folded RNAs best with the alignment (AUC 1.00, against 0.97 and 0.78) and the intra-RNA PAE best without it (0.99, as pLDDT*_r_*, against 0.53 for pTM*_r_*). Adding the familiarity points of the protein–RNA score lowered the development AUC in both branches (0.97 against 1.00, and 0.89 against 0.99), so the subtree uses AF3’s confidence alone. The 41 single-chain RNAs released after the benchmark download were predicted locally with and without the alignment and with the AlphaFold Server. The measures and weights of the subtree were chosen and fitted on the development RNAs, so these RNAs test it. The logistic models rank the development RNAs almost perfectly (AUC 0.997 on pLDDT*_r_* with the alignment and 0.995 on the intra-RNA PAE without it). Among the 2026 RNAs at WEAK, none of the local predictions and 3 of 38 server predictions were well folded, and 2 of 2, 2 of 2 and 3 of 3 were well folded at STRONG (local with and without the alignment, and server) (Table S16; python -m decision_tree.develop_probability –rna).

##### RNA complexes without protein

The development set combines 46 multi-chain RNA complexes released between January 2022 and March 2025, predicted locally with and without the alignment, with the 20 post-cutoff complexes of the main set, which were AlphaFold Server predictions without an alignment. This gives 46 predictions with the alignment (18 well folded) and 64 without it (26 well folded). Six complexes with multi-letter chain identifiers were excluded. The intra-RNA PAE of the longest RNA chain ranked well-folded RNAs best in both branches (AUC 0.86 and 0.91, against at most 0.65 and 0.78 for pTM, ipTM, pLDDT*_r_* and global PAE). Familiarity points were tested by 5-fold cross-validation grouped by sequence cluster and lowered the AUC (0.71 against 0.83 with the alignment, 0.78 against 0.88 without it), so confidence alone was kept again. A duplex question completes this subtree (below).

##### RNA length, alignment and duplexes

Both the alignment and pTM*_r_* matter mainly for RNAs of at least 200 nt. Among the single-chain development RNAs predicted with the alignment, pTM*_r_* ranked well-folded RNAs perfectly at *≥* 200 nt (AUC 1.00, *n* = 31) but below chance at *<* 200 nt (0.48, *n* = 35), where pLDDT*_r_* and the intra-RNA PAE still ranked them well (0.99 each). Without the alignment, none of the 30 single-chain RNAs of *≥* 200 nt was well folded. In RNA complexes without protein, removing the alignment changed accuracy little (median 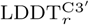 0.74 with and 0.73 without it, 46 complexes). Only the five complexes whose longest RNA has *≥* 200 nt got worse (median change *−*0.07). Many of these complexes are duplexes, in which at least 60% of the nucleotides pair with another chain in the deposited structure (22 of the 66 development complexes). The duplexes are short (median 46 nt in total), carried no RNA alignment beyond the query, and 37 of their 38 predictions were well folded, against 7 of the 72 predictions of the other complexes. The same call made from the AF3 model separates the groups almost as well (89% and 86% of predicted duplexes well folded with and without the alignment, against 7% and 19% of the remaining complexes). The subtree therefore adds whether the AF3 model forms a duplex to the intra-RNA PAE of the longest chain, in one logistic model per branch. A duplex multiplies the odds of a well-folded RNA by 10 with the alignment and by 6 without it, and each Å of intra-RNA PAE by 0.70 and 0.63. The development AUC is 0.98 and 0.95, against 0.86 and 0.91 for the intra-RNA PAE alone (python -m decision_tree.develop_probability –rna). All weights were set on the 66 development complexes, and no 2026 multi-chain RNA entered the design. The 2026 test set has 10 complexes released between June 2025 and May 2026, predicted with the AlphaFold Server. Over its three kinds of prediction, 1 of 16 predictions at WEAK was well folded, and all 3 at MODERATE and all 3 at STRONG were (AUC 1.00 in each; –rna –test). The test set is small, so it agrees with the design but cannot establish its accuracy.

#### T.5. Binding-specificity sets

The binding-specificity sets were scored with the same probabilities (python -m decision_tree.applications –probability). RNAcompete probes are synthetic, so they are never familiar. On the fold side all RNAcompete probes are WEAK, and 88% of the RNAcompete min/maxprobe competition complexes are WEAK, 12% MODERATE and fewer than 1% STRONG. On the interface side most RNAcompete probe complexes are MODERATE (64% of maxprobes and 67% of minprobes, with 19% and 12% STRONG), and the level barely separates maxprobes from minprobes (AUC 0.55). Most v5 probe chains are WEAK (81% of maxprobes and 90% of minprobes). Where the best chain of a RNAcompete min/maxprobe competition complex is STRONG (*n* = 26), ipTM*_r_* ranks the maxprobe first in 83% of complexes and the motif score in 87%, against 53% and 60% where it is MODERATE (*n* = 193) and 43% and 52% where it is WEAK (*n* = 509). The CLIP-atlas entries separate. From WEAK (*n* = 4) through MODERATE (20) to STRONG (121) interface probability of the best RNA chain, the RNA motif is recovered (recall *≥* 0.5) in 75%, 75% and 90% of entries, AF3’s best-scoring RNA motif lies on the native site in 50%, 41% and 88%, and the median DockQ*_Nr_* rises from 0.07 to 0.27 and 0.83. On the fold side 88% of the CLIP entries, mostly released before the cutoff, are STRONG, with a median DockQ*_Nr_* of 0.82 against 0.12 for WEAK entries (*n* = 9)

**Table S16.** RNA-only subtrees of the decision tree. The probability of a well-folded RNA (LDDT^C3^*^′^ >* 0.85) is a logistic regression on AF3’s confidence, fitted with mild L2 regularisation on post-cutoff development data separately for predictions with and without the RNA alignment. PAE is the intra-RNA PAE of the longest RNA chain in Å, pLDDT*_r_* is in %, and *d* = 1 when the AF3 model of an RNA complex forms a duplex (*≥* 60% of the nucleotides paired with another chain). Levels are WEAK below 25%, MODERATE from 25% to 60% and STRONG from 60%. For single-chain RNAs STRONG starts at pLDDT*_r_* 83.0% (WEAK below 79.9%) with the alignment and at an intra-RNA PAE of 3.8 Å (WEAK above 5.1 Å) without it. For RNA complexes STRONG starts at 2.1 Å (8.7 Å for a duplex) with the alignment and 4.0 Å (7.9 Å) without it. Entries give well-folded RNAs over all RNAs at each level.

| Subtree and branch | Log-odds of a well-folded RNA | Set | WEAK | MODERATE | STRONG |
|---|---|---|---|---|---|
| Single-chain, with alignment | −39.0 + 0.474 pLDDT<sub>*r*</sub> | development | 1/51 | – | 14/15 |
|  |  | 2026 local | 0/33 | 2/4 | 2/2 |
| Single-chain, without alignment | 4.71 − 1.14 PAE | development | 0/42 | 1/1 | 4/5 |
|  |  | 2026 local | 0/35 | 0/2 | 2/2 |
|  |  | 2026 server | 3/38 | – | 3/3 |
| Multi-chain, with alignment | 1.15 + 2.33 *d* − 0.352 PAE | development | 1/23 | 2/7 | 15/16 |
|  |  | 2026 local | 0/4 | 1/1 | 1/1 |
| Multi-chain, without alignment | 2.29 + 1.84 *d* − 0.469 PAE | development | 1/30 | 3/10 | 22/24 |
|  |  | 2026 local | 0/4 | 1/1 | 1/1 |
|  |  | 2026 server | 1/8 | 1/1 | 1/1 |

**Fig. S39:**
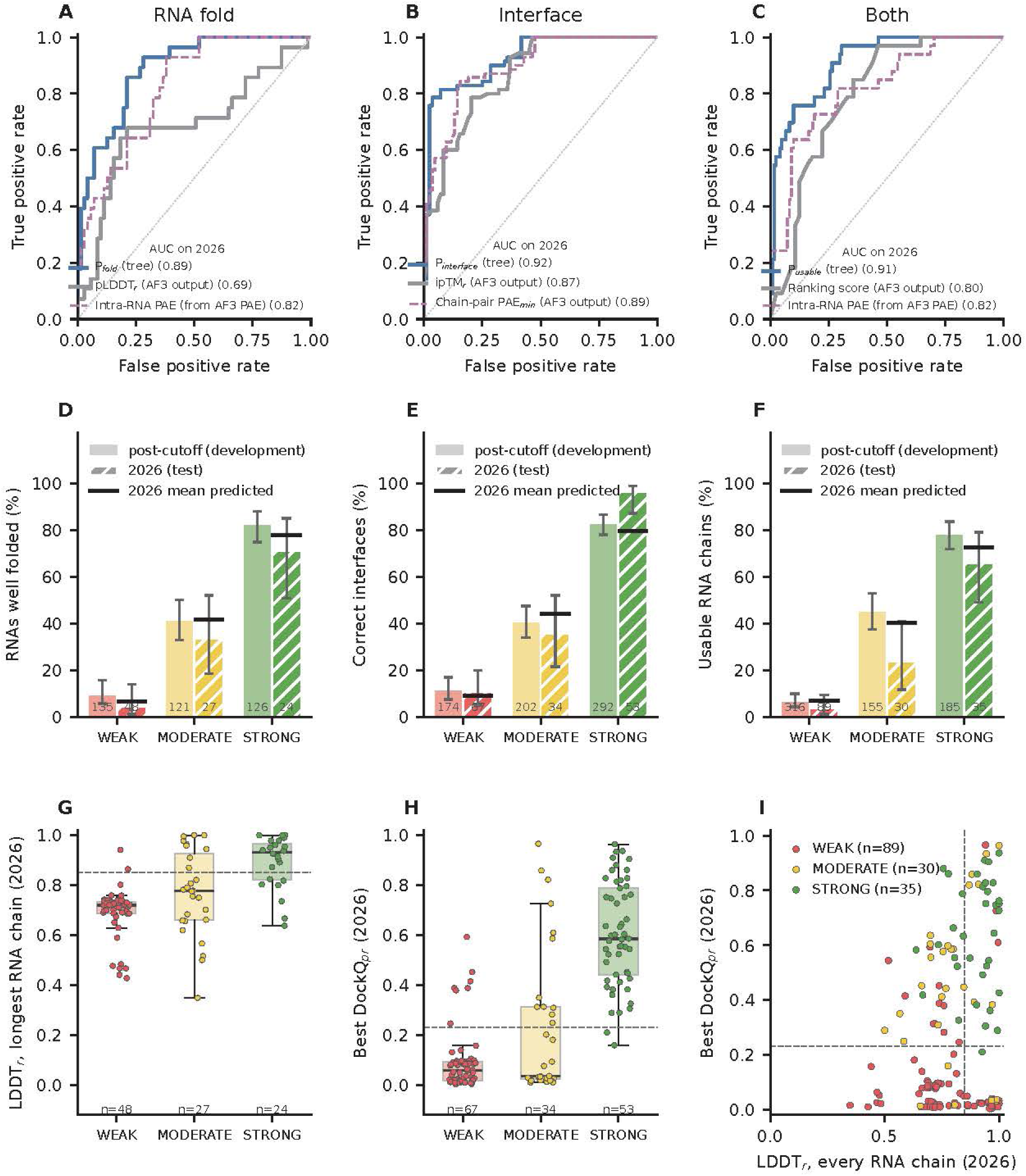
The reliability score on PDB entries released in 2026. 99 protein–RNA complexes released between May and September 2026 (99 RNAs for the fold, 154 RNA chains for the interface and for both), predicted with the AlphaFold Server. **A**–**C**, ROC curves for well-folded RNAs (A), correct interfaces (B) and usable RNA chains, well folded and placed correctly (C). The probabilities of the tree are compared with AF3 outputs (pLDDT*_r_*, ipTM*_r_*, the minimum chain-pair PAE and the ranking score) and with the intra-RNA PAE, which we derive from AF3’s PAE matrix. **D**–**F**, Share of well-folded RNAs (D), correct interfaces (E) and usable RNA chains (F) per level on the post-cutoff development set and on the 2026 set (95% Wilson CI). Black bars give the mean predicted probability of the 2026 targets at each level. **G**, LDDT*_r_* of the longest RNA chain of every 2026 target by fold level. **H**, Best protein–RNA DockQ*_pr_* of every 2026 RNA chain by interface level. **I**, LDDT*_r_* of every 2026 RNA chain against its DockQ*_pr_*, coloured by usable level.

